# Synthesis and structural optimization of 2,7,9-trisubstituted purin-8-ones as FLT3-ITD inhibitors

**DOI:** 10.1101/2022.12.10.519888

**Authors:** Monika Tomanová, Karolína Kozlanská, Radek Jorda, Lukáš Jedinák, Tereza Havlíková, Eva Řezníčková, Miroslav Peřina, Pavel Klener, Alexandra Dolníková, Petr Cankař, Vladimír Kryštof

## Abstract

Therapy of FLT3-positive acute myeloid leukemia still remains complicated, despite the availability of newly approved kinase inhibitors. Various strategies to avoid reduced efficacy of therapy are explored including the development of dual targeting compounds, which inhibit FLT3 and another kinase necessary for survival and proliferation of AML cells. We have designed new 2,7,9-trisubstituted 8-oxopurines as FLT3 inhibitors and report here structure-activity relationship studies. We demonstrated that substituents at positions 7 and 9 modulate activity between CDK4 and FLT3 kinase and the isopropyl group at position 7 increased substantially the selectivity toward FLT3 kinase, which led to the discovery of compound **15a** (9-cyclopentyl-7-isopropyl-2-((4-(piper-azin-1-yl)phenyl)amino)-7,9-dihydro-8*H*-purin-8-one). Cellular analyses in MV4-11 cells revealed inhibition of autophosphorylation of FLT3 kinase in nanomolar doses including suppression of downstream STAT5 and ERK1/2 phosphorylation. We also describe mechanistic studies in cell lines and activity in a mouse xenograft model in vivo.

## 1. Introduction

Acute myeloid leukemia (AML) is the most common type of acute leukemia in adults, who have unfortunately poor prognosis and low survival rate. Approximately 30% of patients bear mutations in FMS-like tyrosine kinase 3 (FLT3), a transmembrane receptor kinase, which plays a key role in normal hematopoietic cell maturation, but its oncogenic activation results in uncontrolled expansion of myeloid precursors. FLT3 can be activated by the internal tandem duplications (ITD) of the juxtamembrane domain or by point mutations in the kinase domain [1]. FLT3 mutation status relates to a prognosis, but it has been also validated as a drug target in AML and hematooncologists can now treat patients with newly approved drugs [2], including those based on small molecule kinase inhibitors, such as sorafenib, midostaurin, gilteritinib, crenolanib or quizartinib [1,3].

Despite available targeted drugs, therapy of FLT3-positive AML remains complicated by high rate of resistance to FLT3 inhibitors, often caused by changes in the kinase domain [4]. Various strategies to avoid reduced efficacy of therapy are explored, including combinations with conventional cytotoxics [2] or modern targeted drugs like BH3 mimetics [5,6], azacytidine [7], or kinase inhibitors [8-10]. Other approaches focus directly on the kinase realm by generating kinase inhibitors targeting resistance-causing FLT3 point mutations [11], or by developing dual targeting compounds, which inhibit FLT3 and another kinase necessary for survival and proliferation of AML cells, for example, JAK2, MEK, MER, or CDK4 [12].

In the current work, we have designed new 2,7,9-trisubstituted 8-oxopurines as FLT3 inhibitors and report here structure-activity relationship (SAR) studies, which led to the discovery of compound **15a**. We also describe mechanistic studies in cell lines and activity in a mouse xenograft model in vivo.

## 2. Results

### 2.1. Chemistry

The synthetic route to a variety of purines **7-10** is described in Scheme 1. Reductive amination was used as a convenient method for the synthesis of derivatives **2** and **3** from commercially available pyrimidine **1**. Thus, we changed the typical reaction sequence to *N*-methyl pyrimidine **2**, where alkylation usually precedes to chlorination step [21]. In the case of **3**, we optimized reaction conditions using 2,2-dimethoxypropane and sodium tri-acetoxyborohydride [22].

The further diverse step was the aromatic substitution of chloride at position 4 with diverse amines using *N,N*-diisopropylethylamine in butanol. Amines were subjected under our modified conditions [23], but some aromatic amines were more problematic. Noteworthy, only desired position was involved in the substitution and no side products or hydrolysis were observed. *N*-methyl intermediate **2** was the most reactive toward aromatic nucleophilic substitution compared to *N*-isopropyl one **3** or non-substituted amine **3**. In these cases, a longer reaction time or addition of base was needed.

Cyclization to purines **7** proceeded with trifluoroacetic anhydride and purinones **8-10** were isolated after reaction with a phosgene solution. Several aromatic and one styryl substituents were introduced by Chan-Lam cross-coupling reaction using conditions from the literature [24], since the conventional method of aromatic nucleophilic substitution failed. 2-Methoxy, 4-methoxyphenyl, 2-tolyl, naphthyl, and styryl boronic acid were used as coupling partners with purinone intermediates **9** with the NH group at position 9. This intermediate was prepared easily after the deprotection of 2,4-dimethoxybenzyl group with TFA.

**Scheme 1.**
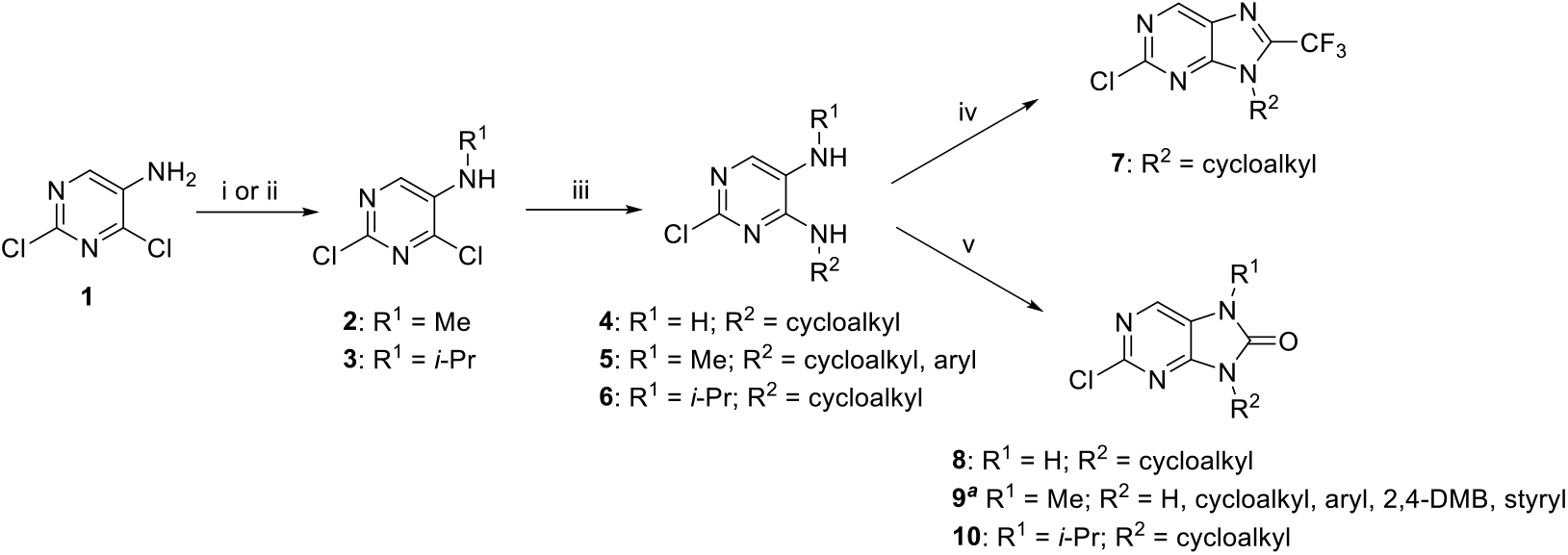
Synthesis of intermediates for Buchwald Hartwig amination. (i) for R^1^ = Me: 37% aq. HCHO, AcOH (conc.), NaBH_3_CN, MeOH, 0 – 5 °C, 20 h; (ii) for R^1^ = *i-*Pr: (CH_3_)_2_C(OCH_3_)_2_, AcOH (conc.), NaBH(OAc)_3_, CH_2_Cl_2_, 0 °C – rt, 20 h; (iii) amine, DIEA, 85 °C, 48 h; (iv) TFAA, toluene, 70 °C, 16 h; (v) for R^1^ = H, R^2^ = cycloalkyl: COCl_2_, 1M LiHMDS in hexane, THF, -10 °C – rt. ^a^Intermediate R^2^ = H was obtained after deprotection of 2,4-DMB group with TFA in anisole at 90 °C for 18 h. Several intermediates R^2^ = aryl, styryl were prepared by Chan-Lam coupling reaction with aryl or styryl boronic acid, Cu_2_S, TMEDA, in MeCN at rt for 24 h.

Buchwald-Hartwig amination was used with piperazineaniline **11** to substitute the chlorine atom at position 2 (Scheme 2) since no general and high yielded method for conventional aromatic nucleophilic substitution was found. The reaction was efficiently performed with precatalyst XPhos Pd G2 under microwave irradiation [25-27]. Only derivatives **13** with R^1^ = H were more problematic. Prolongation of the reaction time from 1 – 2 to 6 – 9 hours and an additional amount of the precatalyst solved the problem; however, deprotected purine **9** (R^2^ = H) was still resistant to the coupling reaction. Finally, the Boc protecting group was removed to give a library of purines **12** – **15**, only a cleavage of the 2,4-DMB group required harsh conditions to get purine **14a**.

**Scheme 2.**
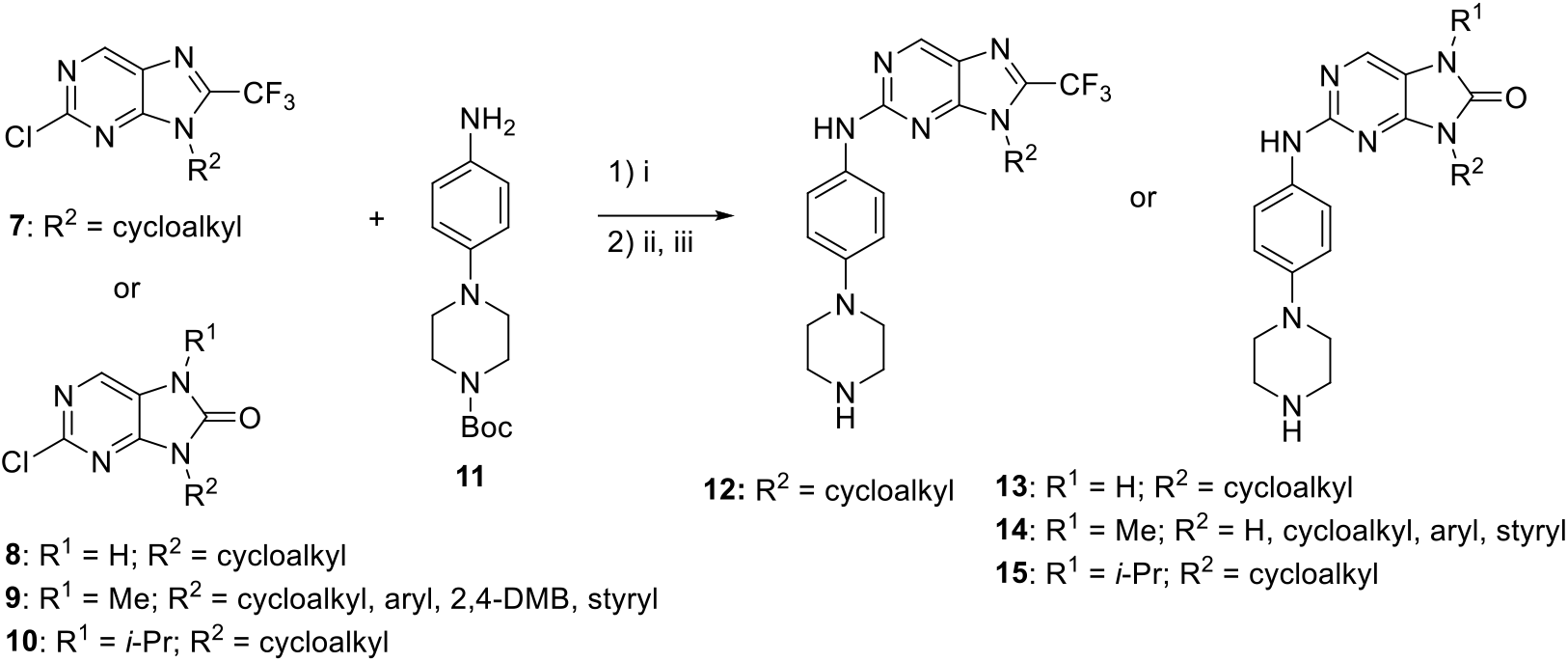
Synthesis of target inhibitors.(i) XPhos Pd G2, K_2_CO_3_, 1,4-dioxane/H2O (4:1), MW (150 W), 100 °C, 1 – 9 h; (ii) 36% HCl, CH_2_Cl_2_/MeOH (1:1), 48 h, (iii) for R^1^ = Me, R^2^ = H: TFA anisole, 90 °C, 5 days.

### 2.2 Kinase inhibitory activities – the SAR study

Newly prepared compounds **12-15** were assayed for inhibition of FLT3-ITD kinase as well as CDK4 and CDK2, known off-targets of structurally related compounds. We observed that CDK4 was sensitive to nearly all compounds with IC_50_ values in mid-nanomolar values. Slight improvement in the anti-kinase potencies was observed with the increasing size of cycloalkyl moieties at position 9 (R2 substituent), namely from cyclopentyl to cycloheptyl. The substitution of the oxo for trifluoromethyl group at position 8 (derivatives **12a-12d**) dramatically improved inhibitory activities to all tested kinases to nanomolar values with slight preferences for CDK4 and FLT3. Compounds **13a-13d** having no substitution at position 7 of the purine ring did not display potency to the tested kinases.

The introduction of a methyl group at position **7** of the purine ring resulted in series **14**, which were prepared with diverse cycloalkyl and aryl substituents as R2. Our results showed the ability to affect the selectivity and, more importantly, the inhibitory activities of compounds because all compounds exceeded the activities of unsubstituted derivative **14a**. An implementation of bulky substituents, namely cyclooctyl, cycloheptyl, 4-methylcyclohexyl, and cyclohexyl lead to derivatives (**14i, 14h, 14g** and **14e**, respectively) exhibiting all tested kinases with IC_50_ ≤ 300 nM. Other derivatives bearing dodeka (**14j**), cyclopentyl (**14d**), 4-hydroxycyclohexyl (**14f**), 4-methylphenyl (**14o**), 4-methoxyphenyl (**14q**) and 4-fluorostyryl (**14t**) lost the CDK2 inhibitory potency (IC_50_ > 1 μM), while inhibition of CDK4 and FLT3 was preserved in mid-nanomolar values.

The introduction of phenyl and cyclobutyl substituents (**14m, 14c**) showed to be important for FLT3 inhibition (IC_50_ = 50 – 60 nM), but lead to diminishing of activities against CDK4 (IC_50_ = 400 – 520 nM). Also, other derivatives with tetrahydropyranyl (**14k**), 2,4-dimethoxybenzyl (**14r**), cyclopropyl (**14b**), and naphthyl (**14s**) exhibited similar specificity for FLT3 kinase over CDK4 and CDK2 (IC_50_ > 1 μM), but the IC_50_ values for FLT3 dropped to mid-nanomolar values.

Finally, we explored the activities of 4 compounds that possess isopropyl substituent at position 7 and cyclopentyl, cyclohexyl, 4-methylcyclohexyl, or cycloheptyl groups (**15a-15d**) at position 9 of the purine ring. All these derivatives display interesting activities for FLT3 kinase (IC_50_ = 30 – 70 nM) and selectivity over tested CDKs. Comparison with counterparts differing in the methyl and isopropyl moieties (pairs **15d** and **14h, 15a** and **14d, 15c** and **14g, 15b** and **14e**) indicates that isopropyl markedly contributes to selectivity against FLT3.

### 2.3 Kinase selectivity

Three representative compounds (**14d, 14e, 15a**) were profiled in 1 μM concentration against 30 kinases that were selected as known off-targets of approved CDK4/6 inhibitors [28]. Compounds **14d** and **14e** differing only in cyclopentyl and cyclohexyl moiety at position **9** of the purine ring were confirmed to be potent CDK4 and FLT3 inhibitors (Figure S1). Additional sensitive kinases include STK16, EphA1, MEKK2, SRC and MER (≤ 10% of residual enzyme activities), which are not targets of structurally similar CDK4/6 inhibitor trilaciclib. We screened also compound **15a**, an analogue of **14d**, bearing isopropyl at position 7 of the purine instead of methyl group. This change rapidly diminished the potency against CDK4 (dropped to 90% of residual enzyme activity), but the preference for FLT3 remained (1^st^ rank) (Figure 2). Further four kinases, namely Src, TTK, Lyn and Abl were inhibited also potently (≤ 10% of residual activities).

**Figure 1.**
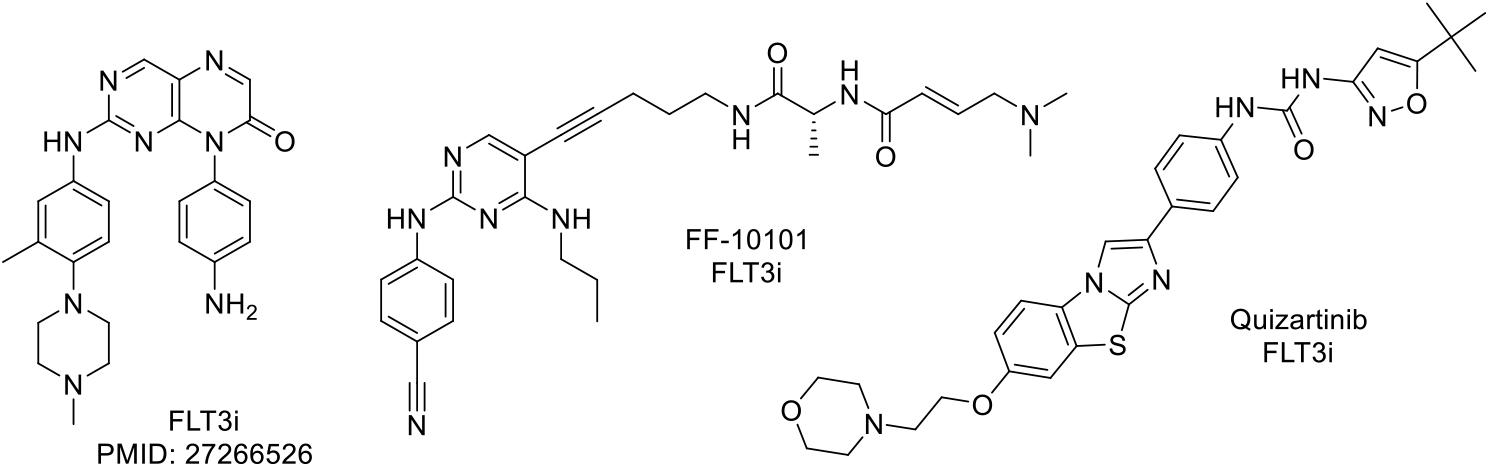
Example of inhibiting FLT3 kinase inhibitors.

**Figure 2.**
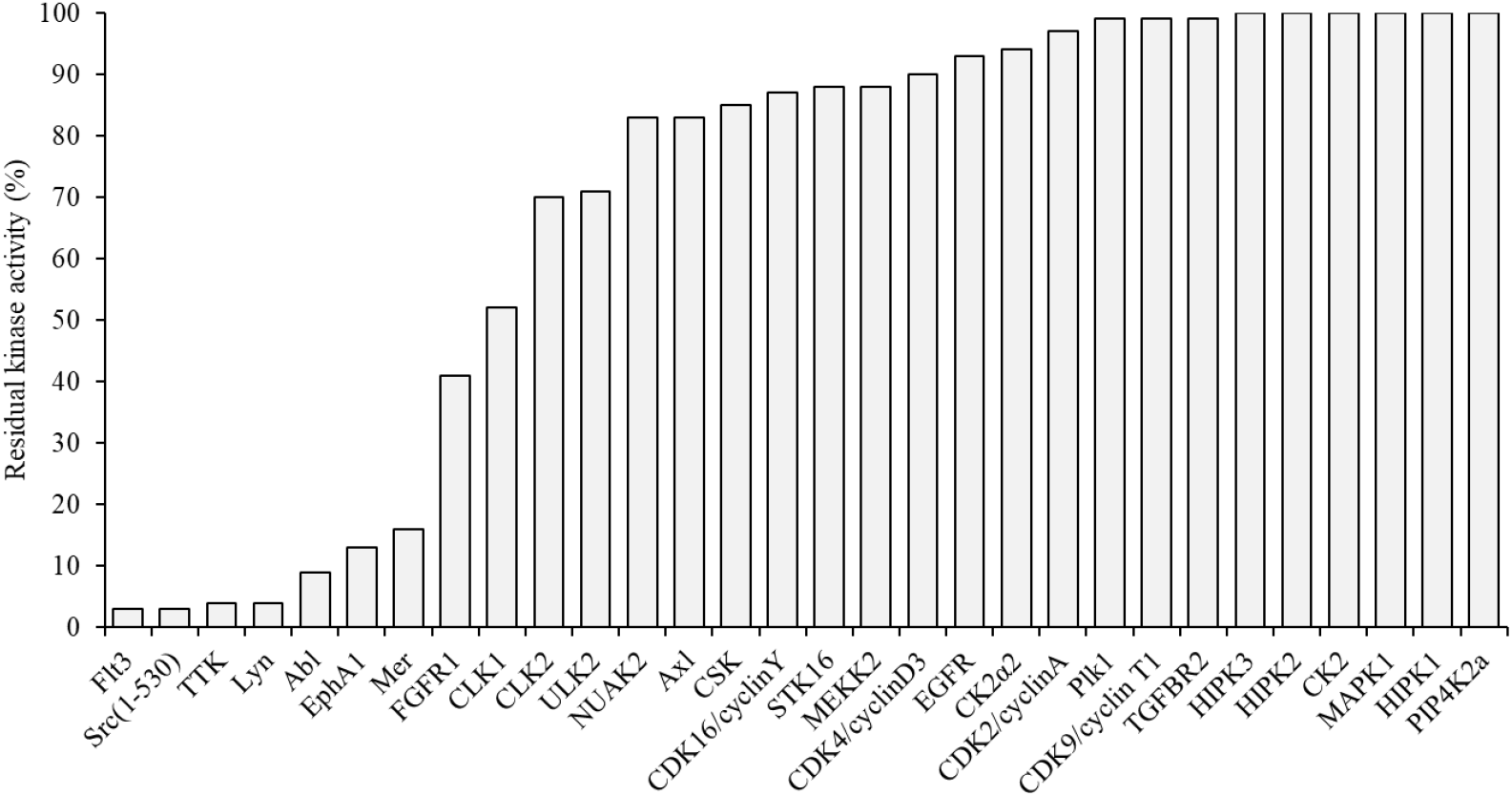
Kinase selectivity profiles of **15a** assayed at 1 μM concentration.

### 2.4 Binding of **15a** in the FLT3 active site

Molecular docking of **15a** into the FLT3 in active conformation (DFG-in) [29] proposed plausible binding in the active site in a manner corresponding to known inhibitors of CDK4, CDK6, and FLT3 [30,31] (Figure 3). The orientation of the compound is consistent with type I binding. In the hinge region, two putative conserved hydrogen bonds are formed between the backbone of Cys694 and **15a**. The isopropyl in position 7 of purine core can make extensive hydrophobic interactions with Val624, Lys644, Val675, Leu 767, as well as the gatekeeper Phe691. The cyclopentyl ring forms hydrophobic contacts with Asp698 and Leu767.

**Figure 3.**
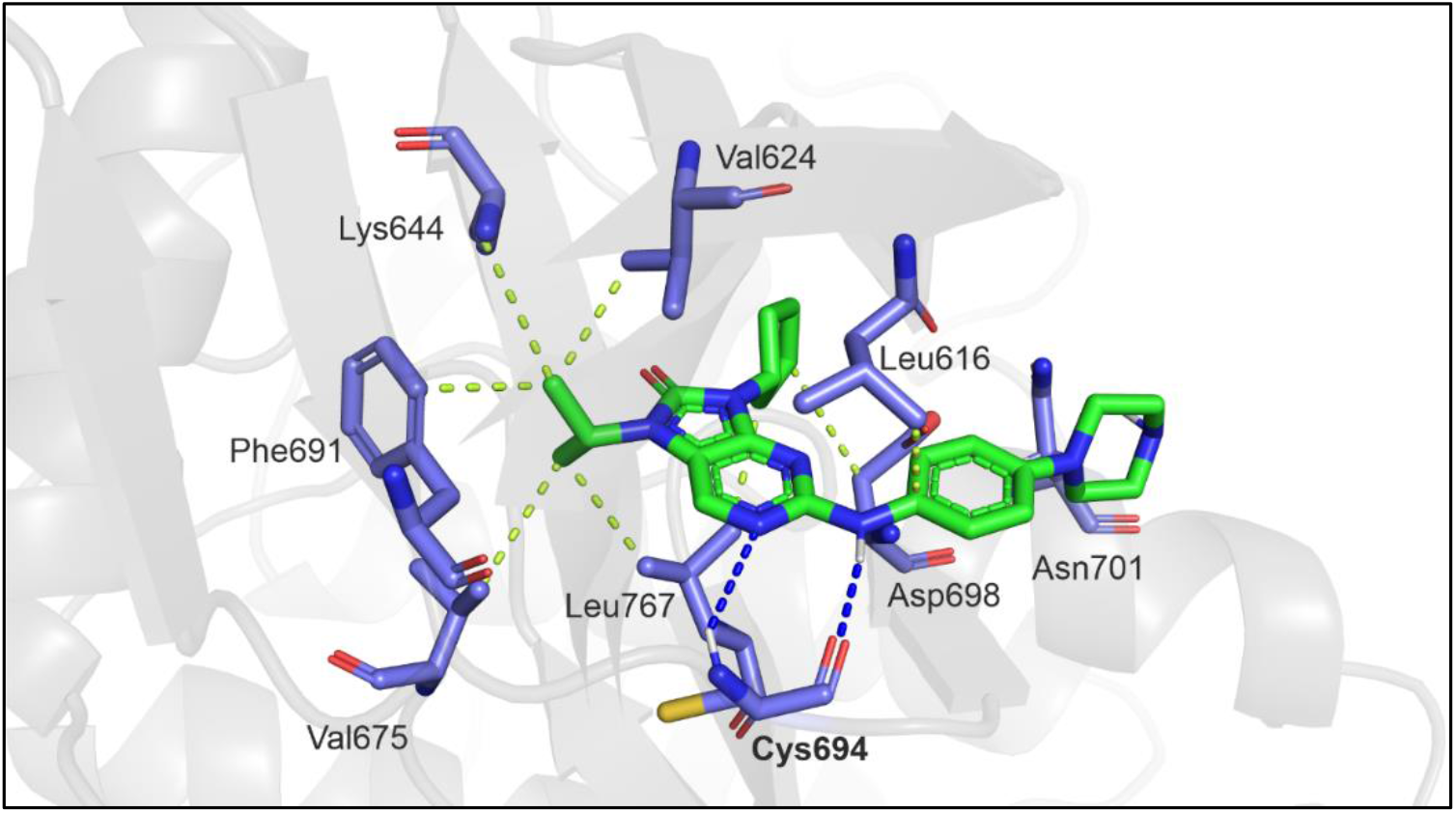
Binding pose of **15a** (green sticks) in the model of FLT3. The kinase is shown in gray with interacting residues shown as blue sticks. Heteroatoms are blue (nitrogens), red (oxygens), yellow (sulfur), and white (hydrogens).

### 2.5 Antiproliferative properties of prepared compounds

In parallel to kinase assays, we measured the effect of compounds on viability of cancer cell lines MV4-11 and K562 using a resazurin-based assay. MV4-11 is an AML cell line dependent on a constitutively activated FLT3-ITD kinase with high sensitivity to FLT3 inhibition, K562 cells (chronic myelogenous leukemia) are independent on FLT3. As shown in Table 1, most of compounds have GI_50_ values in a mid-nanomolar range in MV4-11 cells, while K562 cells were substantially less sensitive (most of GI_50_ > 1 μM). Eleven compounds have displayed GI_50_ below 200 nM in MV4-11 and from these, compounds **14i** and **14e** exerted selectivity index > 20 (ratio for GI_50_’s of K562 and MV4-11 cells). Compound **15a** was the most potent in MV4-11 cells expressing GI_50_ = 50 nM.

**Table 1.**
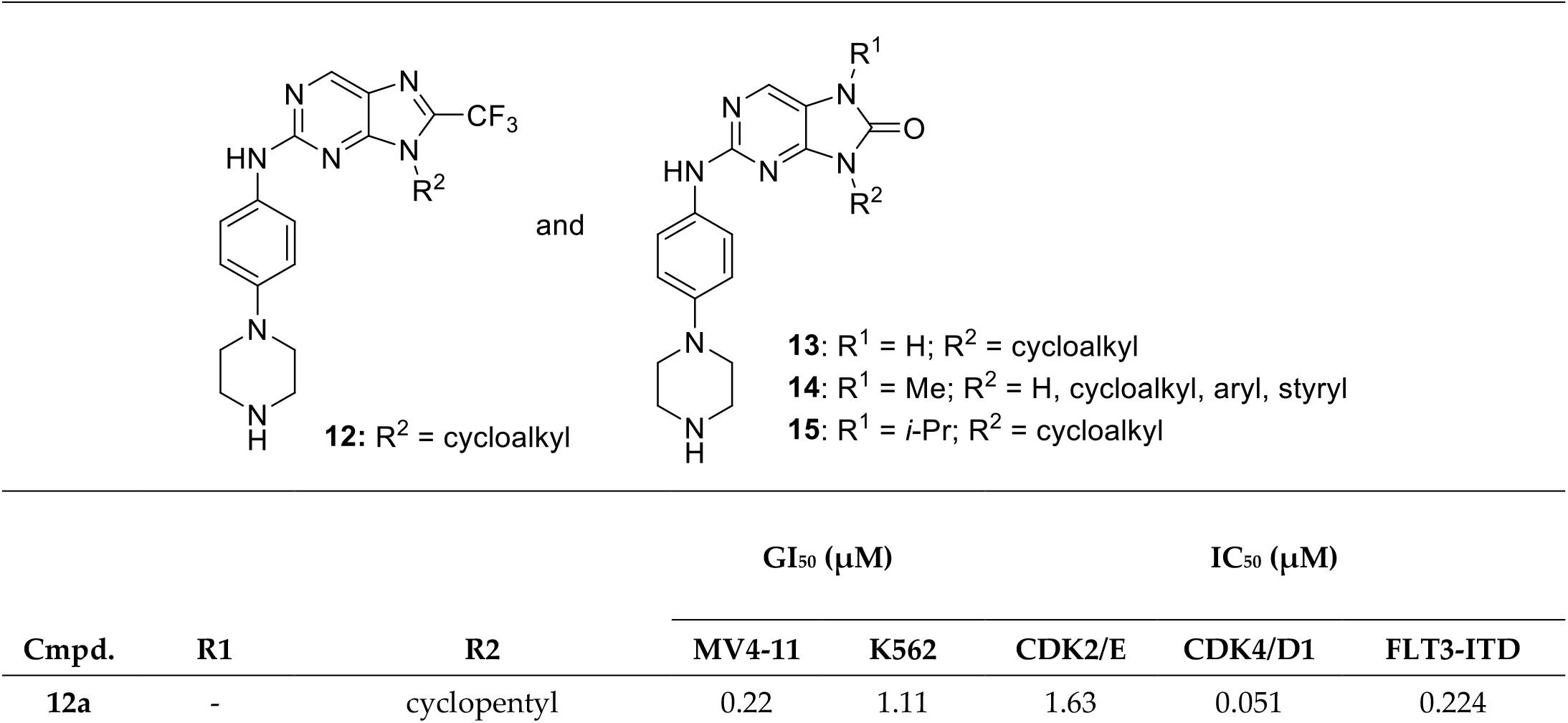
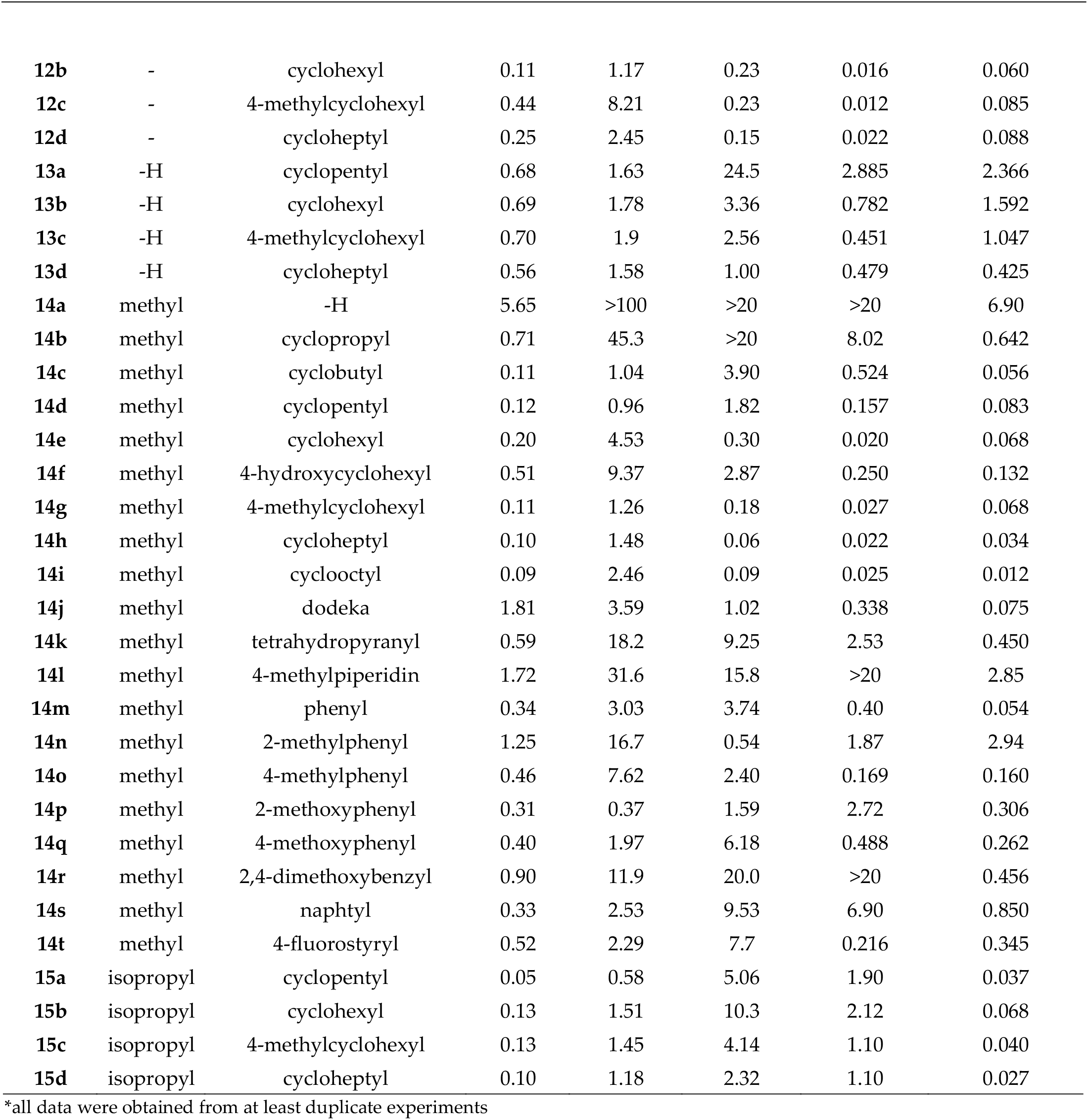
Antiproliferative and kinase inhibitory potencies of novel compounds.

| **12:** R<sup>2</sup> = cycloalkyl<br>and<br>**13:** R<sup>1</sup> = H; R<sup>2</sup> = cycloalkyl<br>**14:** R<sup>1</sup> = Me; R<sup>2</sup> = H, cycloalkyl, aryl, styryl<br>**15:** R<sup>1</sup> = *i*-Pr; R<sup>2</sup> = cycloalkyl |  |  |  |  |  |  |  |
| --- | --- | --- | --- | --- | --- | --- | --- |
| Cmpd. | R1 | R2 | GI<sub>50</sub> (μM) |  | IC<sub>50</sub> (μM) |  |  |
|  |  |  | MV4-11 | K562 | CDK2/E | CDK4/D1 | FLT3-ITD |
| **12a** | - | cyclopentyl | 0.22 | 1.11 | 1.63 | 0.051 | 0.224 |
| **12b** | - | cyclohexyl | 0.11 | 1.17 | 0.23 | 0.016 | 0.060 |
| **12c** | - | 4-methylcyclohexyl | 0.44 | 8.21 | 0.23 | 0.012 | 0.085 |
| **12d** | - | cycloheptyl | 0.25 | 2.45 | 0.15 | 0.022 | 0.088 |
| **13a** | -H | cyclopentyl | 0.68 | 1.63 | 24.5 | 2.885 | 2.366 |
| **13b** | -H | cyclohexyl | 0.69 | 1.78 | 3.36 | 0.782 | 1.592 |
| **13c** | -H | 4-methylcyclohexyl | 0.70 | 1.9 | 2.56 | 0.451 | 1.047 |
| **13d** | -H | cycloheptyl | 0.56 | 1.58 | 1.00 | 0.479 | 0.425 |
| **14a** | methyl | -H | 5.65 | >100 | >20 | >20 | 6.90 |
| **14b** | methyl | cyclopropyl | 0.71 | 45.3 | >20 | 8.02 | 0.642 |
| **14c** | methyl | cyclobutyl | 0.11 | 1.04 | 3.90 | 0.524 | 0.056 |
| **14d** | methyl | cyclopentyl | 0.12 | 0.96 | 1.82 | 0.157 | 0.083 |
| **14e** | methyl | cyclohexyl | 0.20 | 4.53 | 0.30 | 0.020 | 0.068 |
| **14f** | methyl | 4-hydroxycyclohexyl | 0.51 | 9.37 | 2.87 | 0.250 | 0.132 |
| **14g** | methyl | 4-methylcyclohexyl | 0.11 | 1.26 | 0.18 | 0.027 | 0.068 |
| **14h** | methyl | cycloheptyl | 0.10 | 1.48 | 0.06 | 0.022 | 0.034 |
| **14i** | methyl | cyclooctyl | 0.09 | 2.46 | 0.09 | 0.025 | 0.012 |
| **14j** | methyl | dodeka | 1.81 | 3.59 | 1.02 | 0.338 | 0.075 |
| **14k** | methyl | tetrahydropyranyl | 0.59 | 18.2 | 9.25 | 2.53 | 0.450 |
| **14l** | methyl | 4-methylpiperidin | 1.72 | 31.6 | 15.8 | >20 | 2.85 |
| **14m** | methyl | phenyl | 0.34 | 3.03 | 3.74 | 0.40 | 0.054 |
| **14n** | methyl | 2-methylphenyl | 1.25 | 16.7 | 0.54 | 1.87 | 2.94 |
| **14o** | methyl | 4-methylphenyl | 0.46 | 7.62 | 2.40 | 0.169 | 0.160 |
| **14p** | methyl | 2-methoxyphenyl | 0.31 | 0.37 | 1.59 | 2.72 | 0.306 |
| **14q** | methyl | 4-methoxyphenyl | 0.40 | 1.97 | 6.18 | 0.488 | 0.262 |
| **14r** | methyl | 2,4-dimethoxybenzyl | 0.90 | 11.9 | 20.0 | >20 | 0.456 |
| **14s** | methyl | naphtyl | 0.33 | 2.53 | 9.53 | 6.90 | 0.850 |
| **14t** | methyl | 4-fluorostyryl | 0.52 | 2.29 | 7.7 | 0.216 | 0.345 |
| **15a** | isopropyl | cyclopentyl | 0.05 | 0.58 | 5.06 | 1.90 | 0.037 |
| **15b** | isopropyl | cyclohexyl | 0.13 | 1.51 | 10.3 | 2.12 | 0.068 |
| **15c** | isopropyl | 4-methylcyclohexyl | 0.13 | 1.45 | 4.14 | 1.10 | 0.040 |
| **15d** | isopropyl | cycloheptyl | 0.10 | 1.18 | 2.32 | 1.10 | 0.027 |

\*all data were obtained from at least duplicate experiments

FLT3 inhibition causes G1-cell cycle arrest in FLT3-dependent cell lines such as MV4-11; we, therefore, performed flow cytometric analysis to evaluate cell cycle changes in MV4-11 and K562 cell lines treated with 200 nM compounds for 24 h. Data show that most compounds block the cell cycle in the G1 phase in MV4-11 cells. Specifically, 18 compounds arrested > 85% cells in the G1 phase, which is > 30% cells in the G1 phase of the cell cycle than in control cells (Figure 4A). Observed results confirmed the data obtained from viability assays because all non-toxic compounds (e.g. **14a, 14j, 14l, 14n**) did not significantly affect the cell cycle in MV4-11 cells. No changes in cell cycle distribution upon treatment were observed in K562 cells, confirming the specificity of compounds (Figure 4B). Only a few compounds such as **12c, 14e, 14g** – **14i** caused moderate G1 block in K562 that probably stems from strong inhibition of CDK4 (see Table 1). The control experiments were performed with palbociclib and quizartinib; palbociclib was able to significantly arrest both cell lines in G1 phase, while quizartinib arrested selectively only MV4-11.

**Figure 4.**
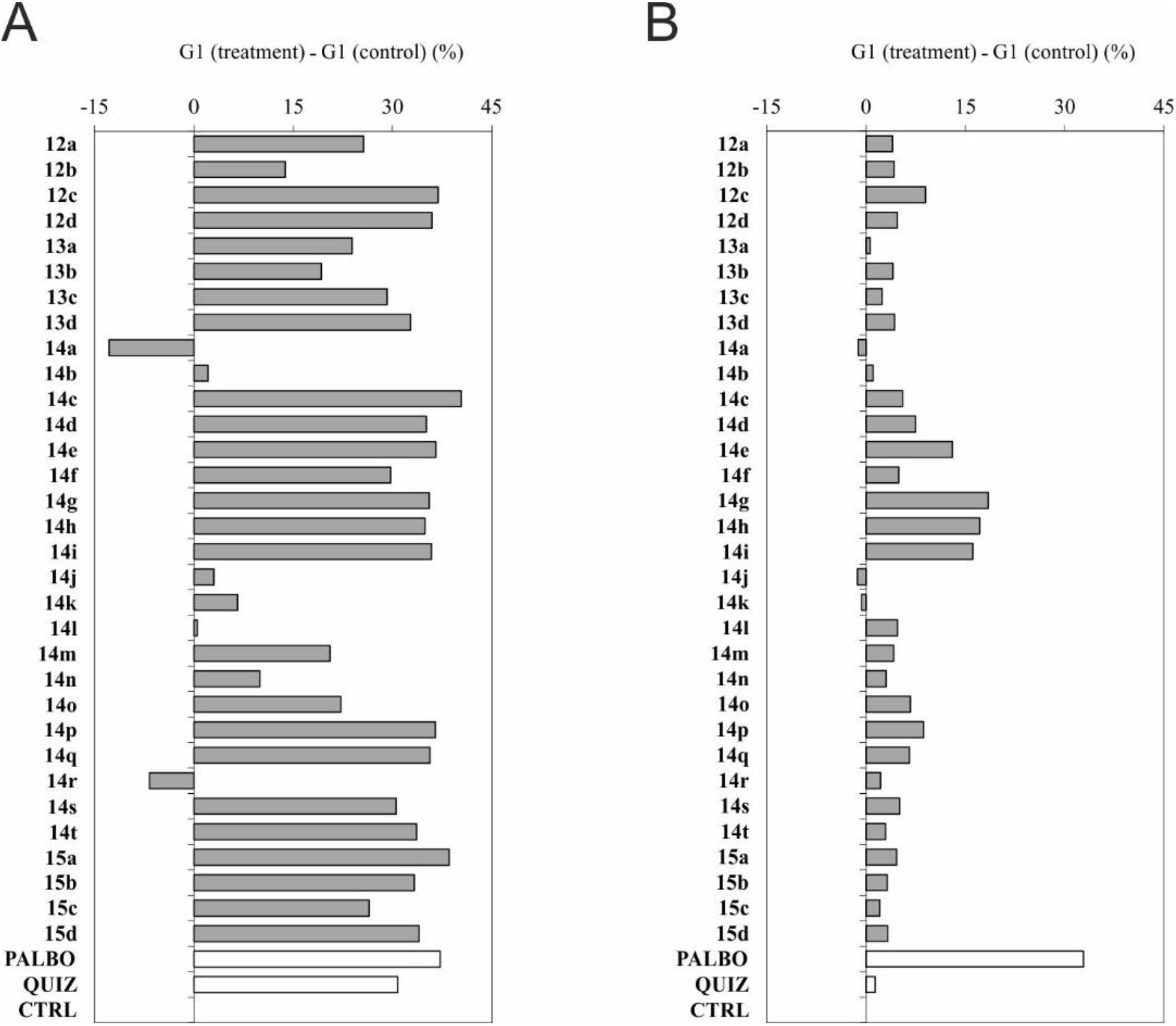
Cell cycle G1 phase population upon treatment of MV4-11 (A) and K562 (B) cells with 200 nM concentration for 24 h. Data (in %) are expressed as difference between G1 populations in treated and control cells. PALBO, palbociclib; QUIZ, quizartinib.

Finally, we performed flow cytometric analysis with different concentrations of **15a** in MV4-11 and MOLM13 (another FLT-ITD positive cell line) and K562 cell lines treated for 24 h (Figure 5). Increasing doses of **15a** arrested cells in the G1 phase of the cell cycle up to 90 % as also observed for FLT3 inhibitor quizartinib. Oppositely, cell line K562 did not show any changes upon treatment with **15a** and quizartinib.

**Figure 5.**
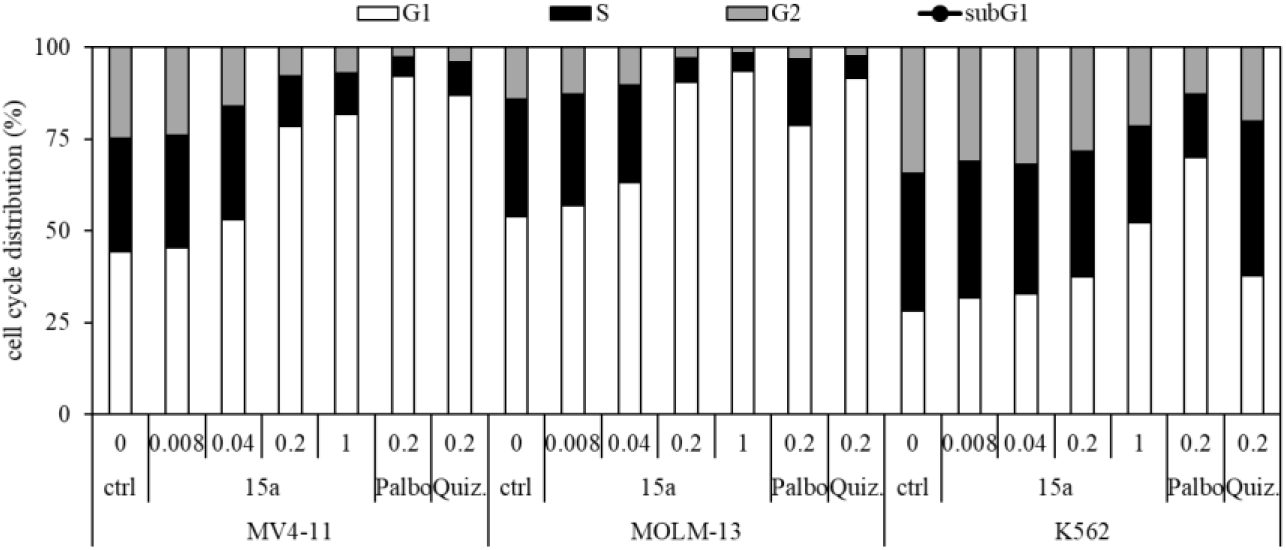
Cell cycle distribution of MV4-11, MOLM-13 and K562 cells treated with different concentrations of **15a** for 24 h. Quiz., quizartinib; Palbo, palbociclib.

### 2.6 Inhibition of FLT3 signaling in vivo

Next, we investigated the compounds for their ability to inhibit FLT3 and its downstream signaling in MV4-11 cell line. Western blot analysis after 1 h treatment of MV4-11 cells revealed that **14e** inhibits autophosphorylation of FLT3 kinase at Y589/591 and subsequently diminishes the phosphorylation of its downstream proteins STAT5 and ERK1/2 in nanomolar doses (Figure 6). A similar inhibitory pattern was observed in quizartinib treated cells.

**Figure 6.**
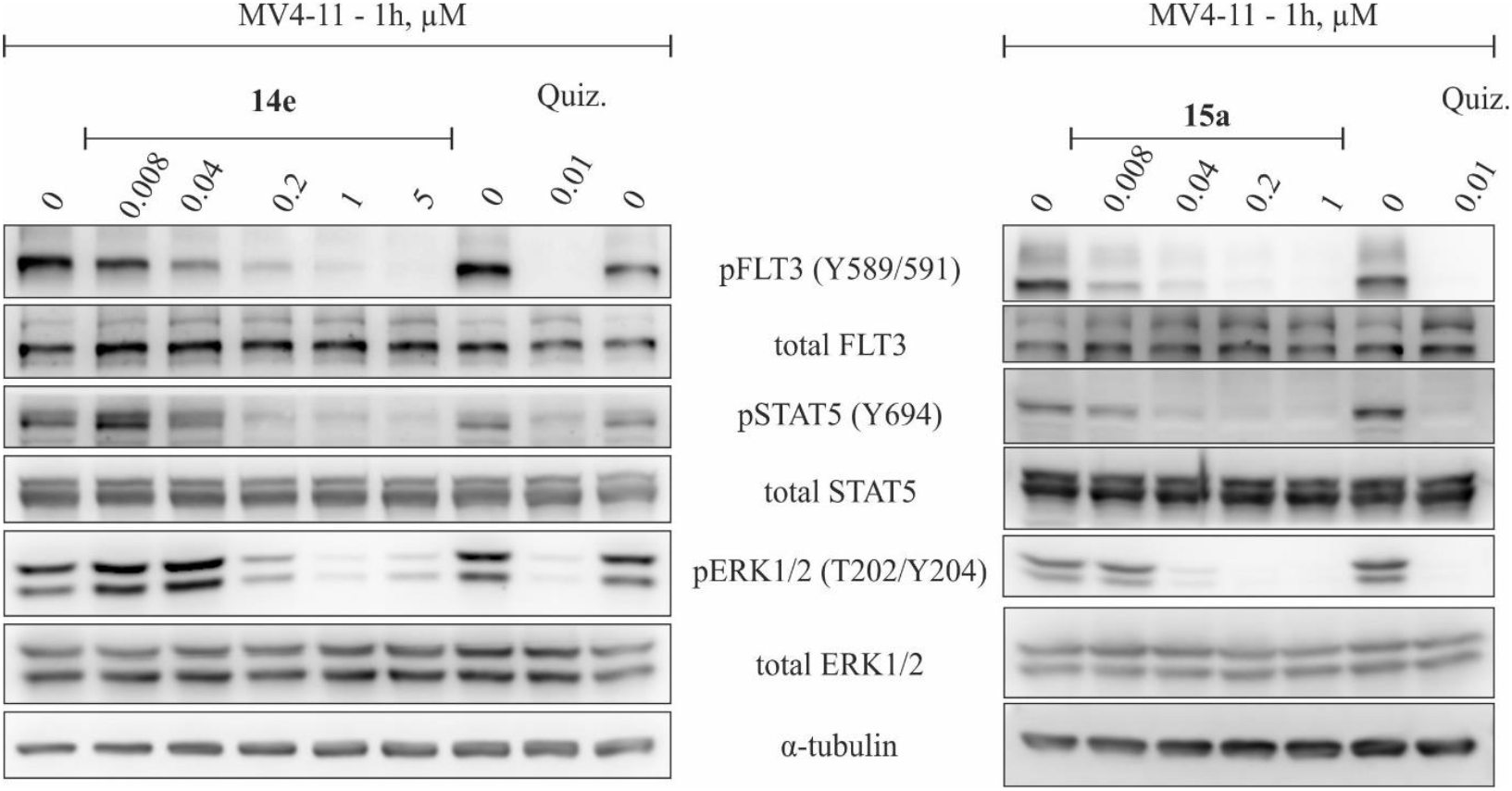
Effect of **14e** and **15a** on phosphorylation of FLT3 and its downstream signaling pathways. MV4-11 cells were treated for 1 h with indicated concentrations of the compounds; quizartinib (Quiz.) was used as a standard. α-tubulin was used for equal protein loading.

In parallel, we examined the effect of the most potent and selective FLT3 inhibitor **15a**, which showed a similar inhibitory pattern of signalling inhibition. The dephosphorylation of FLT3 was observed already at 8 nM concentration and nearly complete suppression of ERK1/2 at 40 nM of **15a**.

### 2.7 In vivo activity of **15a** in MV4-11 xenograft model

Finally, we tested *in vivo* activity of **15a** in a model of AML based on subcutaneous xenotransplantation of MV4-11 cells in immunodeficient mice. As shown in Figure 7A, daily i.p. treatment with **15a** (20 mg/kg) was associated with a significant reduction in the growth of xenografted MV4-11 cells compared to vehicle-dosed control mice. Importantly, no significant reduction of mice weight was observed during the therapy Figure 7B.

**Figure 7.**
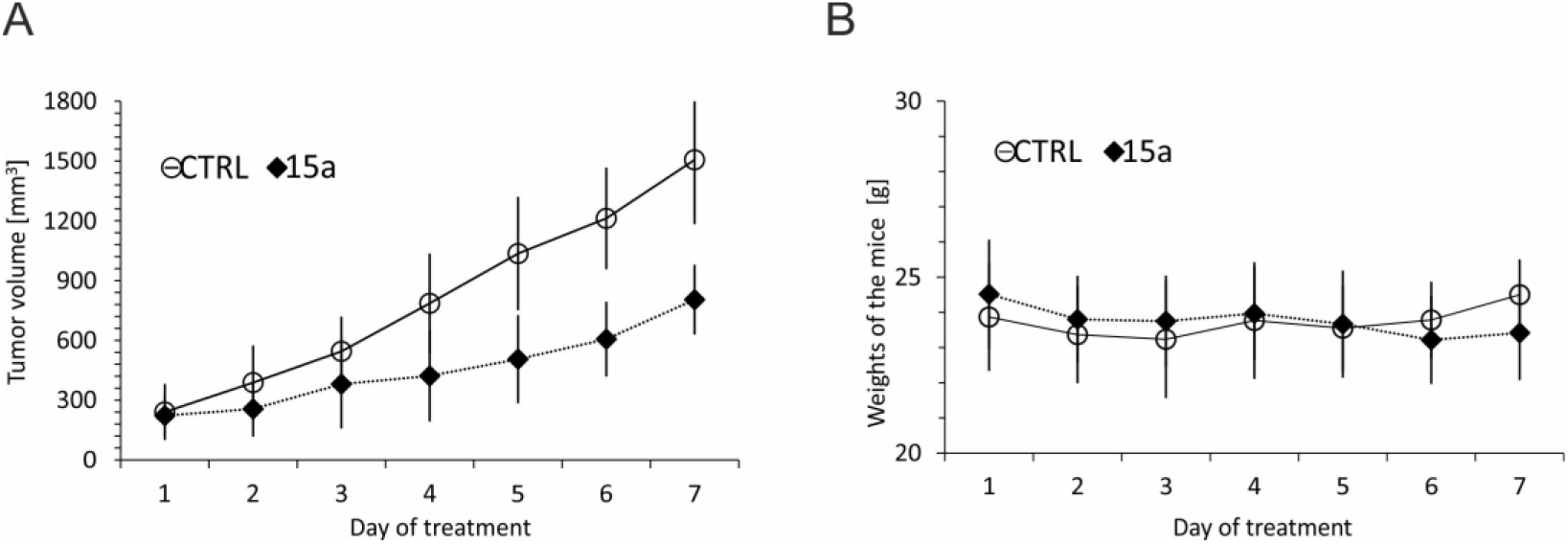
Calculated tumor growth curves (A) and mice weight (B) in MV4-11 xenografted mice treated with **15a**.

## 3. Materials and methods

### 3.1 General information

Starting materials and reagents were purchased from various commercial sources (VWR, Merck, Fluorochem, Acros Organics) and used as received. All reactions were carried out under air. Reaction workup and column chromatography were performed with commercial grade solvents without further purification. All reactions were monitored by LC/MS analysis or by thin-layer chromatography (TLC) using aluminium plates precoated with silica gel (silica gel 60 F254, Merck, US) impregnated with a fluorescent indicator. TLC plates were visualized by exposure to ultraviolet light (λ = 254 nm). Flash chromatography was performed using silica gel (35 – 70 μm partical size) column chromatography. The cross-coupling reactions under microwave irradiation were performed in a 10 mL glass tube sealed with a PTFE coated septa. Reaction mixture was pre-mixed for 1 min with rapid stirring and then irradiated at a maximum power of 150 W with simultaneous cooling using compressed air (24 psi). When the desired temperature was reached (ramping time ∼1 min), the power was automatically adjusted to maintain the set reaction temperature for a specified time. Finally, the vessel was cooled to approximately 40 °C with com-pressed air (cooling time ∼1 min).

### 3.2 Instrumentation

The LC−MS analyses were carried out using UPLC Waters Acquity equipped with PDA and QDa detectors. The system comprised XSelect HSS T3 (Waters) 3 mm×50 mm C18 reverse phase column XP, 2.5μm particles. Mobile phases: 10 mM ammonium acetate in HPLC grade water (A) and gradient grade acetonitrile for HPLC (B). A gradient was mainly formed from 20% to 80% of B in 4.5 min, kept for 1min, with a flow rate of 0.6 mL/min. The MS ESI operated at a 25 V cone voltage, 600°C probe temperature, and 120°C source temperature.

All ^1^H and ^13^C NMR experiments were performed at magnetic field strengths of 11.75 T (with operating frequencies 500.16 MHz for ^1^H and 125.77 MHz for ^13^C) and 9.39 T (with operating frequencies 399.78 MHz for ^1^H and 100.53 MHz for ^13^C) at an ambient temperature (20°C). ^1^H and ^13^C spectra were referenced relative to the signal of DMSO-*d*_*6*_ (^1^H: *δ* = 2.50 ppm, ^13^C: *δ* = 39.51 ppm) or CDCl_3_ (^1^H: *δ* = 7.260 ppm, ^13^C: *δ* = 77.160 ppm). HRMS analyses were performed using UPLC Dionex Ultimate 3000 equipped with an Orbitrap Elite high-resolution mass spectrometer, Thermo Exactive plus, operating at full scan mode (120,000 FWMH) in the range of 100−1,800 m/z. The settings for electrospray ionization were as follows: oven temperature of 150 °C and a source voltage of 3.6 kV. The acquired data were internally calibrated with diisooctyl phthalate as a contaminant in MeOH (m/z 391.2843).

Melting points were determined on VEB Analytik Dresden PHMK 78/1586 apparatus. Microwave experiments were conducted in the CEM Discover SP closed vessel microwave synthesizer.

### 3.3 General Synthetic procedure 1: Pyrimidine-4,5-diamines formation

To a solution of **1, 2**, or **3** (1.00 mmol) in butanol (10 mL), corresponding amine (1.00 mmol) and *N,N*-diisopropylethylamine (259 mg, 348 μL, 2.00 mmol) were added. The reaction mixture was heated at 85°C for 48 hours. Then, butanol was evaporated under reduced pressure and the crude product was purified by column chromatography (usually 1:1 hexane/EtOAc).

### 3.4 General Synthetic procedure 2: Cyclization with TFAA

0.75 mmol of compound **4** was put into a flask, toluene (1 mL) and trifluoroacetic anhydride (3 mL) were added. The mixture was heated at 70 °C for 16 hours. The crude mixture was diluted with dichloromethane (20 mL), then washed with 10% K_2_CO_3_ (30 mL) and distilled water (30 mL), and dried over MgSO_4_. The solvent was evaporated under reduced pressure (rotovap) and the residue was purified by column chromatography (usually 5:1 hexane/EtOAc).

### 3.5 General Synthetic procedure 3: Cyclization with phosgene

0.70 mmol of compound **4, 5**, or **6** was dissolved in anhydrous THF (10 mL) under inert atmosphere of nitrogen and cooled to -10 °C. After slow addition of 15 wt % COCl_2_ in toluene (560 μL, 0.85 mmol), 1M LiHMDS in hexane (1.4 mL, 1.40 mmol) was added portionwise. After 1 hour, the reaction mixture was cooled to ambient temperature and stirred for 20 minutes. The mixture was then evaporated under reduced pressure and the residue was purified by column chromatography (usually 2:1 EtOAc/hexane).

### 3.6 Preparation of intermediate **9** (R^1^ = Me, R^2^ = H)

2-Chloro-9-(2,4-dimethoxybenzyl)-7-methyl-7,9-dihydro-8*H*-purin-8-one (200 mg, 0.59 mmol) was suspended in anisole (3.15 mL) and TFA (2.6 mL) was added. The reaction mixture was stirred at 90°C for 18 hours. TFA was evaporated by the flow of nitrogen and anisole was evaporated using RVO. A crude mixture was dissolved in MeOH, solid NaHCO_3_ was added to neutralize the mixture and evaporated with silica gel. The product was purified by column chromatography (4:1 to 2:1 hexane/EtOAc). After evaporation and trituration with diethylether, the product was filtered-off to yield 2-chloro-7-methyl-7,9-dihydro-8*H*-purin-8-one 56 mg (52 %) as a white solid.

### 3.7 General Synthetic procedure 4: Chan-Lam cross-coupling reaction

A 10 mL vial was charged with Cu_2_S (8 mg, 0.05 mmol), MeCN (0.6 mL), and *N,N,N*′,*N*′-tetramethylethane-1,2-diamine (30 μL, 0.20 mmol). After 1h of stirring, 2-chloro-7-methyl-7,9-dihydro-8*H*-purin-8-one (18 mg, 0.10 mmol) and boronic acid (0.20 mmol) were added and the mixture was stirred at ambient temperature for 24 hours. A crude mixture was loaded on Celite, evaporated using RVO, and purified by column chromatography (2:1 hexane/EtOAc).

### 3.8 General Synthetic procedure 5: Buchwald-Hartwi amination and Boc deprotection

A microwave vial was charged with **7, 8, 9** or **10** (0.25 mmol), 1 mL of mixture 1,4-dioxane/H_2_O (4:1), *tert*-butyl 4-(4-aminophenyl)piperazine-1-carboxylate (55.4 mg, 0.20 mmol), K_2_CO_3_ (138 mg, 1.00 mmol), and XPhos Pd G2 (4 mg, 2 mol%). The reaction mixture was heated at 100 °C under microwave irradiation for 1 – 9 hours, the end of the reaction was monitored using HPLC/MS. The mixture was evaporated under reduced pressure and purified by column chromatography (1:1 hexane/EtOAc). All prepared Boc-protected intermediates (0.15 mmol) were dissolved in 2.8 mL of DCM/MeOH (1:1) and 36% HCl (240 μL) was added. After full conversion (checked by HPLC-MS; approximately 48 hours), 1 mL of distilled water was added together with solid K_2_CO_3_ to neutralize the mixture. Solvents were evaporated and product was purified on column chromatography (10:1 DCM/MeOH).

### 3.9 Cell cultures and viability assay

Human cell lines were obtained from the European Collection of Authenticated Cell Cultures (K562) and Cell lines service (MV4-11) and were cultivated according to the provider’s instructions. Briefly, cells were maintained in RPMI-1640 (MV4-11) or DMEM (K562) supplemented with 10% fetal bovine serum, penicillin (100 U/mL), and streptomycin (0.1 mg/mL) at 37 °C in 5% CO_2_. For the viability assays, cells were treated in triplicate with six different doses of each compound for 72 hours. After treatments, resazurin (Sigma Aldrich) solution was added for 4 hours, and fluorescence of resorufin corresponding to live cells was measured at 544 nm/590 nm (excitation/emission) using a Fluoroskan Ascent microplate reader (Labsystems). The GI_50_ value, the drug concentration lethal to 50% of the cells, was calculated from the dose response curves that resulted from the assays.

### 3.10 Kinase inhibition assays

CDK/cyclin complexes and FLT3 kinases were assayed as previously described [32,33]. Protein kinase selectivity was evaluated at a single concentration (1 μM) by screening against 30 enzymes at Eurofins Discovery (France).

### 3.11 Immunoblotting

Cell lysates were prepared, and then proteins were separated on SDS-polyacrylamide gels and electroblotted onto nitrocellulose membranes. After blocking, overnight incubation with specific primary antibodies, and incubation with peroxidase-conjugated secondary antibodies, peroxidase activity was detected with Super-Signal West Pico reagents (Thermo Scientific) using a CCD camera LAS-4000 (Fujifilm). The specific antibodies were purchased from Cell signaling (anti-FLT3, clone 8F2; anti-phospho-FLT3 Y589/591, clone 30D4; anti-STAT5; anti-phospho-STAT5 Y694; anti-ERK1/2; anti-phospho-ERK1/2 T202/Y204; peroxidase-conjugated secondary antibodies) or Merck (anti-α-tubulin, clone DM1A)

### 3.12 Flow cytometry

Asynchronously growing cells were seeded into a 96-well plate and, after a preincubation period, treated with tested compounds for 24 hours. The treated cells were stained directly (or after trypsinization for the adherent cells) with the 5× staining solution (17 mM trisodium citrate dihydrate, 0.5% IGEPAL CA-630, 7.5 mM spermine tetrahydrochloride, and 2.5 mM Tris; pH 7.6 containing 50 mg/mL propidium iodide). After the staining, DNA content was analyzed by flow cytometry using a 488 nm laser (BD FACS Verse with software BD FACSuite™, version 1.0.6.). Cell cycle distribution was analyzed using ModFit LT (Verity Software House, version 5.0). Palbociclib, trilaciclib and quizartinib were purchased from MedChemExpress.

### 3.13 Molecular docking

Molecular docking was performed into the previously published model of FLT3 in the active conformation [29]. The 3D structure of **15a** and the conformations with lowest energy were obtained by molecular mechanics with Avogadro 1.90.0. Polar hydrogens were added to ligand and protein with the AutoDock Tools program and rigid docking was performed using AutoDock Vina 1.059 [34]. Interactions between the **15a** and FLT3 were determined using PLIP (protein-ligand interaction profiler) [35] and figure was generated using Pymol ver. 2.0.4 (Schrödinger, LLC).

### 3.14 Experimental therapy

The experimental design was approved by the Institutional Animal Care and Use Committee (MSMT-37334/2020-4). Immunodeficient adult female NOD.Cg-*Prkdc*^*scid*^*Il2rg*^*tm1Wjl*^/SzJ mice (referred to as NSG mice) were purchased from The Jackson Laboratory and preserved in a pathogen-free environment in individually ventilated cages, provided with sterilized food and water. NSG mice were subcutaneously inoculated with 10 × 10^6^ leukemia cells. After all mice developed palpable tumors, they were stratified into cohorts with comparable tumor volumes and therapy with **15a** (20 mg/kg, daily i.p. dosing, 7 consecutive days) was initiated (=day 1, D1). Each cohort contained 6 animals. Three perpendicular dimensions (in millimeters) were measured daily with a digital caliper and tumor volumes were calculated using the following formula: π/6 × length × width × height. An experiment was terminated (and mice euthanized) when tumors exceeded 20 mm in the largest diameter.

## 4. Conclusion

In summary, we prepared a series of 2,7,9-trisubstituted-purin-8-nones to compare structure-activity relationships on CDK2, CDK4, and FLT3 kinases. It was revealed that FLT3 kinase was more tolerant in structural variety of substituents at positions 7 and 9 than CDK4 kinase. Introduction of the isopropyl group at position 7 was significantly beneficial for selectivity toward FLT3 kinase. The more detailed selectivity study on 30 kinases, known as off-targets of approved CDK4/6 inhibitors, with three representative compounds showed effective inhibition of four additional kinases STK16, TTK, NUAK2, Lyn and Src. Cellular analyses with MV4-11 cells treated the selected candidates **14e** and **15a** confirmed inhibition of autophosphorylation of FLT3 kinase, subsequent dephosphorylation of the downstream proteins STAT5 and ERK1/2 in nanomolar doses, and G1-cell cycle arrest. Pharmacological properties of the model compound studied on Caco-2 cells indicated substantially higher stability of the inhibitor **14e** in comparison with trilaciclib. To our delight, significant reduction of tumor growth was observed during the therapy of the MV4-11 xenograft model.

## Supplementary Materials

Figure S1: Kinase selectivity profiles of 14e and 14d; Procedures for the synthesis of intermediates 2-10; Characterization of final compounds 12, 13, 14 and 15; Copies of ^1^H and ^13^C NMR spectra of final compounds; HPLC of final compounds; References.

## Author Contributions

Synthesis, analytical data processing, writing, M.T.; synthesis, methodology, T.H. and L.J.; writing—original draft preparation, R.J., M.T.; biochemistry and biology, investigation, K.K., E.Ř., P.K., A.D.; docking, M.P.; conceptualization, supervision, project administration, funding acquisition, V.K., P.C.

## Funding

The work was supported by the European Regional Development Fund (CZ.02.1.01/0.0/0.0/16_019/0000868), Czech Science Foundation (20-25308S and 21-06553S), the Ministry of Health of the Czech Republic (NV19-08-00144) and Palacký University Olomouc (IGA_PrF_2022_007 and IGA_PrF_2022_022).

## Institutional Review Board Statement

The animal study protocol was approved by the Institutional Animal Care and Use Committee (Charles University in Prague).

## Data Availability Statement

All data are reported in the manuscript and in the Supplementary Material and are available from corresponding author upon request.

## Acknowledgements

We would like to thank Jana Hudcová and Veronika Vojáčková for technical assistance. Plasmids for the expression of CDK2 and cyclin A2 were kindly gifted by Dr. Daniel Fisher (IGMM, CNRS, Montpellier, France).

## Conflicts of Interest

The authors declare no conflict of interest.

## Notes

### Competing Interest Statement

The authors have declared no competing interest.

