## Supporting Information for "Synthesis and structural optimization of 2,7,9-trisubstituted purin-8-ones as FLT3-ITD inhibitors"

**Content:**

Kinase selectivity profiles of **14e** and **14d**

Procedures for the synthesis of intermediates **2-10**

Characterization of final compounds **12**, **13**, **14** and **15**

Copies of $^{1}H$ and $^{13}C$ NMR spectra of final compounds

HPLC of final compounds

References

**Figure S1.** Kinase selectivity profiles of **14e** (white bars) and **14d** (grey bars) assayed at 1 μM concentration.

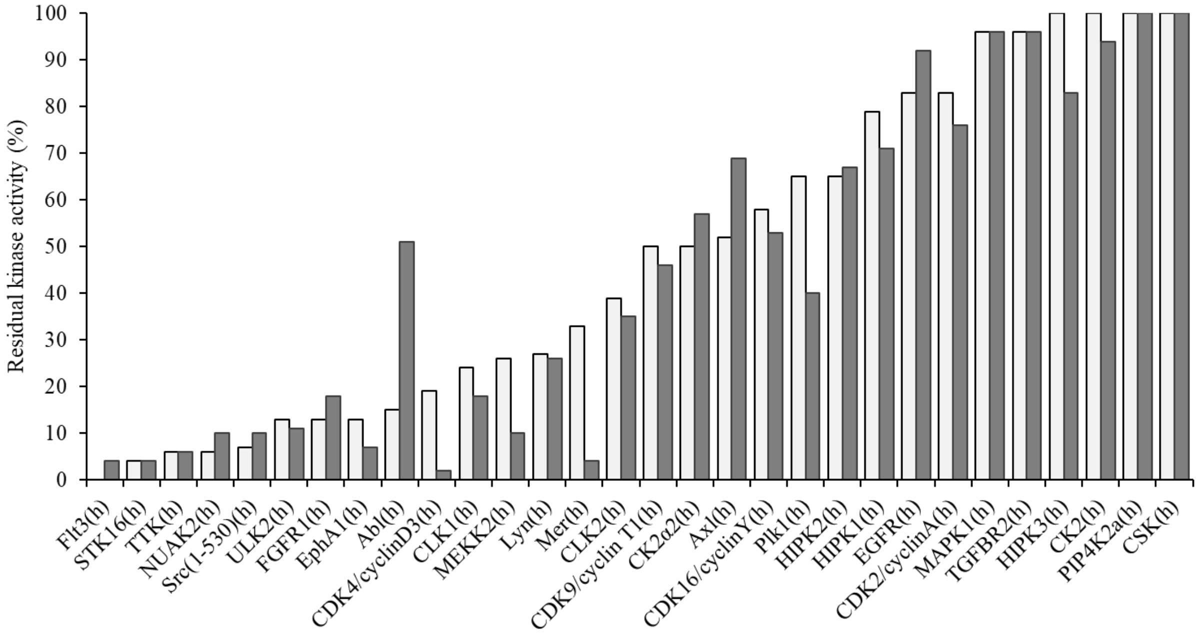

**Procedures for the synthesis of intermediates 2-10**

**Method i**

*2,4-dichloro-N-methylpyrimidin-5-amine* **2** [1]

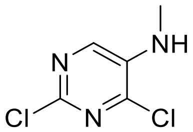

A solution of 2,4-dichloropyrimidine-5-amine **1** (5 g, 30.5 mmol) in MeOH (90 mL) was cooled to 0°C. Glacial acetic acid (10 mL, 183 mmol) was added, and then a solution of 37% aq. formaldehyde (2.7 mL, 36.6 mmol). A mixture was stirred for 1 hour, and then pulled out from an ice bath and an additional amount of acetic acid (20 mL) was added. After another 3 hours, $NaBH_3CN$ (4.6 g, 73.2 mmol) was added in two portions within two hours. After stirring a mixture for 20 hours at 5°C, a solvent was concentrated using RVO, diluted with water (100 mL), and cooled in an ice bath. Solid $NaHCO_3$ was added to neutralize acetic acid. Then, a water phase was extracted with DCM (2x 80 mL) and combined organic layers were washed with brine, dried over $MgSO_4$, and evaporated using RVO. The solid residue was diluted with MeOH (10 mL) and cooled in an ice bath and water (80 mL). After proper mixing, a solid was filtered off and washed with water to yield 4.4 g (81 %) of **2** as a white solid. $^1$H NMR (400 MHz, $CDCl_3$) $\delta$ 7.88 (s, 1H), 4.37 (s, 1H), 2.98 (s, 3H). $^{13}$C NMR (101 MHz, $CDCl_3$) $\delta$ 146.5, 146.0, 139.1, 138.7, 30.0. HRMS (ESI-TOF): calcd for $C_5H_6Cl_2N_3$ [M+H]$^+$ 177.9933, found 177.9934.

**Method ii**

*2,4-dichloro-N-isopropylpyrimidin-5-amine* **3** [2]

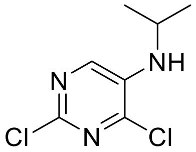

A round bottom flask was charged with **1** (5 g, 30.5 mmol), 2,2-dimethoxypropane (61 mL, 50 mmol), and glacial acetic acid (7.3 mL, 134 mmol). A mixture was cooled in an ice bath. Then, a suspension of $NaBH(OAc)_3$ (26 g, 122 mmol) in DCM (61 mL) was added. After 2 hours, a reaction mixture was pulled out from the ice bath and mixed for 20 hours. The mixture was concentrated under reduced pressure; diluted with DCM (150 mL), washed with 10% $K_2CO_3$ (100 mL), water (100 mL), and brine (100 mL), dried over $MgSO_4$, and evaporated using RVO. The crude product was purified by chromatography (1:1 hexane/EtOAc) to yield 6.18 g, 98 % of **3** as a yellow oil.

$^1$H NMR (400 MHz, $CDCl_3$) $\delta$ 7.89 (s, 1H), 4.09 (d, $J$ = 6.3 Hz, 1H), 3.75 – 3.60 (m, 1H), 1.29 (d, $J$ = 6.3 Hz, 6H). $^{13}$C NMR (101 MHz $CDCl_3$) $\delta$ 146.4, 145.6, 139.9, 137.2, 44.4, 22.6. HRMS (ESI-TOF): calcd for $C_7H_{10}Cl_2N_3$ $[M+H]^+$ 206.0246, found 206.0246.

**Method iii**

To a solution of **2 (R$^1$ = Me)** or **3 (R$^1$ = *i*-Pr)** (1.00 mmol) in butanol (10 mL), corresponding amine (1.00 mmol) and *N,N*-diisopropylethylamine (259 mg, 348 µL, 2.00 mmol) were added. The reaction mixture was heated at 85°C. After stirring for 48 hours, butanol was evaporated under reduced pressure and the residue was purified by column chromatography (usually in 1:1 hexane/EtOAc).

*2-chloro-N$^4$-cyclopentylpyrimidine-4,5-diamine* [3]

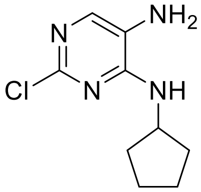


Yield 70 %, brown/pink solid. $^1$H NMR (400 MHz, $CDCl_3$) $\delta$ 7.57 (s, 1H), 5.06 (d, $J$ = 7.6 Hz, 1H), 4.40 (h, $J$ = 7.0 Hz, 1H), 2.99 (s, 2H) 2.19 – 2.06 (m, 2H), 1.81 – 1.56 (m, 4H), 1.51 – 1.35 (m, 2H). $^{13}$C NMR (101 MHz, $CDCl_3$) $\delta$ 157.2, 153.1, 142.1, 123.6, 52.7, 33.4 23.9. HRMS (ESI-TOF): calcd for $C_9H_{14}ClN_4$ $[M+H]^+$ 213.0902, found 213.0903.

*2-chloro-N$^4$-cyclohexylpyrimidine-4,5-diamine* [4]

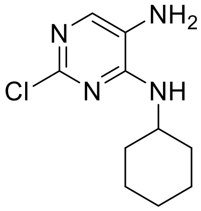


Yiled 73 %, brown/pink solid. $^1$H NMR (400 MHz, $CDCl_3$) $\delta$ 7.57 (s, 1H), 4.97 (d, $J$ = 7.2 Hz, 1H), 4.08 – 3.90 (m, 1H), 2.97 (s, 2H), 2.12 – 1.96 (m, 2H), 1.74 (dt, $J$ = 13.2, 3.6 Hz, 2H), 1.65 (dt, $J$ = 12.9,

3.8 Hz, 1H), 1.43 (qt, *J* = 12.2, 3.4 Hz, 2H), 1.20 (qd, *J* = 12.1, 3.5 Hz, 3H). $^{13}C$ NMR (101 MHz, $CDCl_3$) $\delta$ 156.8, 153.2, 142.3, 123.4, 49.4, 33.2, 25.7, 24.9. HRMS (ESI-TOF): calcd for $C_{10}H_{16}ClN_4$ $[M+H]^+$ 227.1058, found 227.1058.

*2-chloro-N⁴-(4-methylcyclohexyl)pyrimidine-4,5-diamine*

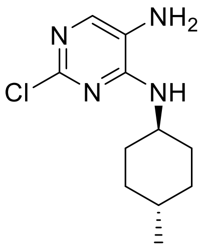

Yield 52 %, pale pink solid. $^1H$ NMR (400 MHz, $CDCl_3$) $\delta$ 7.55 (s, 1H), 4.91 (d, *J* = 7.7 Hz, 1H), 3.98 – 3.86 (m, 1H), 2.96 (s, 2H), 2.13 – 1.98 (m, 2H), 1.71 (d, *J* = 11.3 Hz, 2H), 1.34 (tdd, *J* = 12.9, 6.1, 3.4 Hz, 1H), 1.25 – 1.05 (m, 5H), 0.89 (d, *J* = 6.5 Hz, 3H), 0.86 – 0.78 (m, 1H). $^{13}C$ NMR (101 MHz, $CDCl_3$) $\delta$ 156.8, 153.2, 142.2, 123.5, 49.7, 33.9, 33.3, 32.2, 22.3. HRMS (ESI-TOF): calcd for $C_{11}H_{18}ClN_4$ $[M+H]^+$ 241.1215, found 241.1216.

*2-chloro-N⁴-cycloheptylpyrimidine-4,5-diamine*

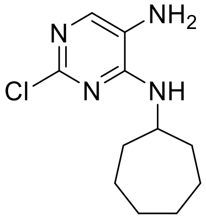

Yield 65 %, pale pink solid. $^1H$ NMR (400 MHz, DMSO-$d_6$) $\delta$ 7.35 (s, 1H), 6.54 (d, *J* = 7.5 Hz, 1H), 4.92 (s, 2H), 4.07 – 3.97 (m, 1H), 2.00 – 1.83 (m, 2H), 1.74 – 1.31 (m, 10H). $^{13}C$ NMR (101 MHz, DMSO-$d_6$) $\delta$ 152.4, 147.0, 135.5, 126.9, 50.9, 34.0, 28.0, 23.6. HRMS (ESI-TOF): calcd for $C_{11}H_{18}ClN_4$ $[M+H]^+$ 241.1215, found 241.1216.

*2-chloro-N⁴-cyclopropyl-N⁵-methylpyrimidine-4,5-diamine*

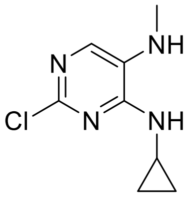

Yield 70 %, ochre solid. $^1$H NMR (400 MHz, $CDCl_3$) $\delta$ 7.45 (s, 1H), 5.36 (s, 1H), 3.01 (s, 1H), 2.94 – 2.86 (m, 1H), 2.80 (s, 3H), 0.94 – 0.76 (m, 2H), 0.64 – 0.47 (m, 2H). $^{13}$C NMR (101 MHz, $CDCl_3$) $\delta$ 157.3, 151.1, 135.6, 128.2, 31.1, 24.1, 7.3. HRMS (ESI-TOF): calcd for $C_8H_{12}ClN_4$ $[M+H]^+$ 199.0745, found 199.0745.

*2-chloro-$N^4$-cyclobutyl-$N^5$-methylpyrimidine-4,5-diamine*

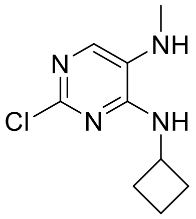


Yield 84 %, white solid.$^1$H NMR (400 MHz, $CDCl_3$) $\delta$ 7.44 (s, 1H), 5.25 (d, $J$ = 6.9 Hz, 1H), 4.65 – 4.51 (m, 1H), 3.02 (s, 1H), 2.81 (s, 3H), 2.55 – 2.34 (m, 2H), 1.95 – 1.80 (m, 2H), 1.80 – 1.68 (m, 2H). $^{13}$C NMR (101 MHz, $CDCl_3$) 155.4, 151.2, 136.0, 127.8, 46.2, 31.5, 31.12 15.3. HRMS (ESI-TOF): calcd for $C_9H_{14}ClN_4$ $[M+H]^+$ 213.0902, found 213.0903.

*2-chloro-$N^4$-cyclopentyl-$N^5$-methylpyrimidine-4,5-diamine* [5]

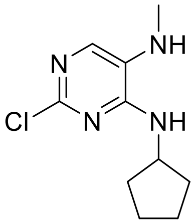


Yield 91 %, white solid. $^1$H NMR (400 MHz, $CDCl_3$) $\delta$ 7.41 (s, 1H), 5.13 (d, $J$ = 6.6 Hz, 1H), 4.41 (h, $J$ = 7.0 Hz, 1H), 3.15 (s, 1H), 2.81 (s, 3H), 2.18 – 2.05 (m, 2H), 1.93 – 1.55 (m, 4H), 1.55 – 1.33 (m, 2H). $^{13}$C NMR (101 MHz, $^{13}$C NMR (101 MHz, $CDCl_3$) $\delta$ 156.0, 150.9, 135.0, 128.0, 52.8, 33.3, 31.0, 23.9. HRMS (ESI-TOF): calcd for $C_{10}H_{16}ClN_4$ $[M+H]^+$ 227.1058, found 227.1058.

*2-chloro-$N^4$-cyclohexyl-$N^5$-methylpyrimidine-4,5-diamine*

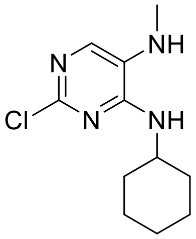

Yield 81 %, pale yellow solid. $^{1}$H NMR (400 MHz, $CDCl_3$) $\delta$ 7.98 (s, 1H), 4.84 (p, $J$ = 8.4 Hz, 1H), 3.42 (s, 3H), 2.33 – 2.14 (m, 2H), 2.10 – 1.89 (m, 4H), 1.83 – 1.48 (m, 4H). $^{13}$C NMR (101 MHz, $CDCl_3$) $\delta$ 152.7, 152.3, 151.2, 132.4, 122.3, 53.9, 29.5, 27.5, 24.9. HRMS (ESI-TOF): calcd for $C_{11}H_{18}ClN_4$ $[M+H]^+$ 241.1215, found 241.1214.

*4-((2-chloro-5-(methylamino)pyrimidin-4-yl)amino)cyclohexan-1-ol*

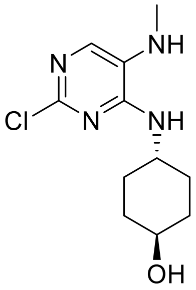


Yield 79 %, white solid. $^{1}$H NMR (400 MHz, DMSO-$d_6$) $\delta$ 7.17 (s, 1H), 6.60 (d, $J$ = 6.8 Hz, 1H), 5.18 – 5.10 (br. s, 1H), 4.58 (t, $J$ = 3.8 Hz, 1H), 3.84 – 3.73 (br. s, 1H), 3.49 – 3.36 (br. s, 1H), 2.72 – 2.65 (m, 3H), 1.95 – 1.79 (m, 4H), 1.25 (t, $J$ = 8.0 Hz, 4H). $^{13}$C NMR (101 MHz, DMSO-$d_6$) $\delta$ 152.8, 146.8, 130.8, 128.3, 68.3, 48.7, 34.0, 30.1, 29.4. HRMS (ESI-TOF): calcd for $C_{11}H_{18}ClN_4O$ $[M+H]^+$ 257.1164, found 257.1162.

*2-chloro-N$^{5}$-methyl-N$^{4}$-(4-methylcyclohexyl)pyrimidine-4,5-diamine*

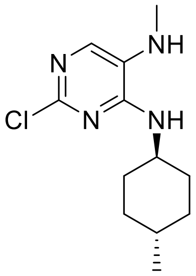


Yield 78 %, light brown solid. $^{1}$H NMR (400 MHz, $CDCl_3$) $\delta$ 7.44 (s, 1H), 4.81 (d, $J$ = 7.6 Hz, 1H), 4.00 – 3.88 (m, 1H), 2.81 (s, 3H), 2.17 – 1.99 (m, 2H), 1.78 – 1.68 (m, 2H), 1.22 – 1.07 (m, 4H), 0.91 (d, $J$ = 6.4 Hz, 3H). $^{13}$C NMR (101 MHz, $CDCl_3$) $\delta$ 155.9, 151.6, 136.2, 127.7, 49.7, 33.9, 33.3, 32.2, 31.2, 22.3. HRMS (ESI-TOF): calcd for $C_{12}H_{20}ClN_4$ $[M+H]^+$ 255.1371, found 255.1371.

*2-chloro-N^4-cycloheptyl-N^5-methylpyrimidine-4,5-diamine*

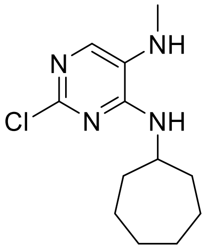

Yield 84 %, white solid. $^1$H NMR (400 MHz, $CDCl_3$) $\delta$ 7.43 (s, 1H), 4.97 (d, $J$ = 7.5 Hz, 1H), 4.27 – 4.13 (m, 1H), 2.94 (s, 1H), 2.81 (s, 3H), 2.06 – 1.95 (m, 2H), 1.71 – 1.58 (m, 4H), 1.57 – 1.43 (m, 6H). $^{13}$C NMR (101 MHz, $CDCl_3$) $\delta$ 155.5, 151.4, 135.9, 127.7, 51.5, 35.0, 31.1, 28.4, 24.0. HRMS (ESI-TOF): calcd for $C_{12}H_{20}ClN_4$ [M+H]$^+$ 255.1371, found 255.1375.

*2-chloro-N^4-cyclooctyl-N^5-methylpyrimidine-4,5-diamine*

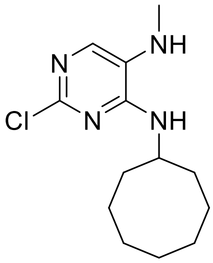

Yield 82 %, white solid. $^1$H NMR (400 MHz, $CDCl_3$) $\delta$ 7.44 (s, 1H), 4.93 (d, $J$ = 7.4 Hz, 1H), 4.28 – 4.17 (m, 1H), 2.81 (s, 3H), 1.98 – 1.80 (m, 2H), 1.79 – 1.35 (m, 12H). $^{13}$C NMR (101 MHz, $CDCl_3$) $\delta$ 155.6, 151.6, 136.1, 127.8, 50.6, 30.0, 31.2, 27.5, 25.6, 23.7. HRMS (ESI-TOF): calcd for $C_{12}H_{22}ClN_4$ [M+H]$^+$ 269.1528, found 269.1529.

*2-chloro-N^4-cyclododecyl-N^5-methylpyrimidine-4,5-diamine*

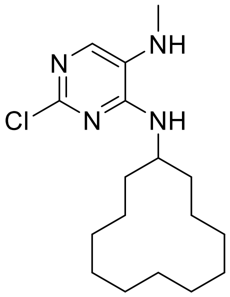

Yield 86 %, white solid. $^1$H NMR (500 MHz, DMSO-$d_6$) $\delta$ 7.16 (s, 1H), 6.54 (d, $J$ = 7.8 Hz, 1H), 5.12 (q, $J$ = 4.8 Hz, 1H), 4.22 (h, $J$ = 6.7 Hz, 1H), 2.70 (d, $J$ = 4.9 Hz, 3H), 1.65 (dq, $J$ = 13.3, 6.6, 6.1 Hz, 2H), 1.51 – 1.24 (m, 20H). $^{13}$C NMR (126 MHz, DMSO-$d_6$) $\delta$ 153.0, 146.8, 130.6, 128.1, 46.2, 29.6,

29.4, 23.4, 23.2, 23.1, 22.9, 21.3. HRMS (ESI-TOF): calcd for $C_{17}H_{30}ClN_4$ $[M+H]^+$ 325.2154, found 325.2152.

*2-chloro-N⁵-methyl-N⁴-(tetrahydro-2H-pyran-4-yl)pyrimidine-4,5-diamine*

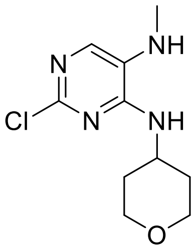


Yield 80 %, pale yellow solid. $^1$H NMR (400 MHz, $CDCl_3$) $\delta$ 7.26 (s, 1H), 4.93 (d, $J$ = 7.4 Hz, 1H), 4.11 – 4.00 (m, 1H), 3.79 (ddd, $J$ = 11.8, 3.9, 2.2 Hz, 2H), 3.35 (td, $J$ = 11.8, 2.5 Hz, 2H), 1.84 (ddd, $J$ = 12.6, 4.3, 2.1 Hz, 2H), 1.32 (qd, $J$ = 11.5, 4.5 Hz, 2H). $^{13}$C NMR (101 MHz, $CDCl_3$) $\delta$ 155.4, 150.9, 135.7, 128.0, 66.9, 47.1, 33.2, 31.0. HRMS (ESI-TOF): calcd for $C_{10}H_{16}ClN_4O$ $[M+H]^+$ 243.1007, found 243.1008.

*2-chloro-N⁵-methyl-N⁴-(1-methylpiperidin-4-yl)pyrimidine-4,5-diamine*

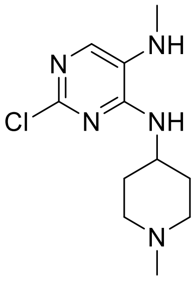


Yield 94 %, pale yellow solid. $^1$H NMR (400 MHz, DMSO-$d_6$) $\delta$ 7.19 (s, 1H), 6.93 (d, $J$ = 7.2 Hz, 1H), 5.41 (d, $J$ = 4.5 Hz, 1H), 4.01 – 3.86 (m, 1H), 3.09 – 3.00 (m, 2H), 2.69 (d, $J$ = 4.9 Hz, 3H), 2.41 (s, 3H), 2.01 – 1.88 (m, 2H), 1.65 (qd, $J$ = 13.4, 12.6, 3.7 Hz, 2H). $^{13}$C NMR (101 MHz, DMSO-$d_6$) $\delta$ 152.66, 146.43, 130.80, 128.47, 53.19, 46.04, 44.37, 29.93, 29.32. HRMS (ESI-TOF): calcd for $C_{11}H_{19}ClN_5$ $[M+H]^+$ 256.1323, found 256.1323.

*2-chloro-N⁵-methyl-N⁴-phenylpyrimidine-4,5-diamine*

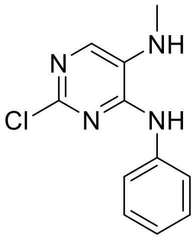

Yield 59 %, light brown solid. $^1H$ NMR (400 MHz, DMSO-$d_6$) $\delta$ 8.70 (s, 1H), 7.69 – 7.64 (m, 2H), 7.49 (s, 1H), 7.39 – 7.32 (m, 2H), 7.07 (tt, $J$ = 7.2, 1.1 Hz, 1H), 2.79 (s, 3H). $^{13}C$ NMR (126 MHz, DMSO-$d_6$) $\delta$ 150.4, 145.6, 139.1, 133.6, 129.2, 128.7, 123.1, 120.7, 29.6. HRMS (ESI-TOF): calcd for $C_{11}H_{12}ClN_4$ $[M+H]^+$ 235.0745, found 235.0745.

*2-chloro-$N^5$-methyl-$N^4$-(p-tolyl)pyrimidine-4,5-diamine*

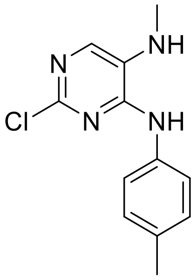


Yield 53 %, light brown solid. $^1H$ NMR (400 MHz, DMSO-$d_6$) $\delta$ 8.61 (s, 1H), 7.55 – 7.49 (m, 2H), 7.46 (s, 1H), 7.19 – 7.13 (m, 2H), 5.49 (d, $J$ = 4.5 Hz, 1H), 3.34 (s, 1H), 2.78 (d, $J$ = 4.2 Hz, 3H), 2.28 (s, 3H). $^{13}C$ NMR (101 MHz, DMSO-$d_6$) $\delta$ 150.6, 145.7, 136.4, 133.3, 132.3, 129.1, 129.1, 121.0, 29.6, 20.5. HRMS (ESI-TOF): calcd for $C_{12}H1_4ClN_4$ $[M+H]^+$ 249.0902, found 249.0902.

*2-chloro-$N^4$-(2,4-dimethoxybenzyl)-$N^5$-methylpyrimidine-4,5-diamine*

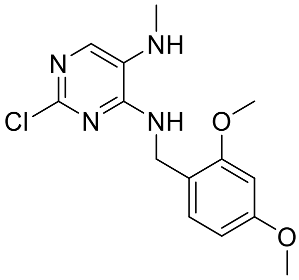


Yield 75 %, white solid. $^1H$ NMR (400 MHz, $CDCl_3$) $\delta$ 7.39 (s, 1H), 7.24 (s, 1H), 6.45 – 6.41 (m, 2H), 5.39 (d, $J$ = 6.3 Hz, 1H), 4.56 (d, $J$ = 5.5 Hz, 2H), 3.81 (s, 3H), 3.78 (s, 3H), 2.77 (s, 3H). $^{13}C$ NMR (101 MHz, $CDCl_3$) $\delta$ 160.8, 158.9, 156.0, 150.8, 135.1, 131.2, 128.2, 118.5, 104.1, 98.8, 55.5, 40.8, 30.9. HRMS (ESI-TOF): calcd for $C_{14}H_{18}ClN_4O_2$ $[M+H]^+$ 309.1113, found 309.1111.

*2-chloro-N^4^-cyclopentyl-N^5^-isopropylpyrimidine-4,5-diamine*

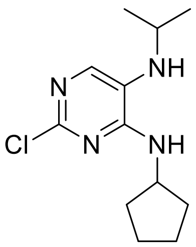


Yield 74 %, white solid.$^{1}$H NMR (400 MHz, $CDCl_3$) $\delta$ 7.49 (s, 1H), 5.21 (d, $J$ = 6.8 Hz, 1H), 4.39 (h, $J$ = 7.1 Hz, 1H), 3.39 (p, $J$ = 6.2 Hz, 1H), 2.54 (s, 1H), 2.17 – 2.06 (m, 2H), 1.78 – 1.57 (m, 4H), 1.49 – 1.37 (m, 2H), 1.18 (d, $J$ = 6.2 Hz, 6H). $^{13}$C NMR (101 MHz, $CDCl_3$) $\delta$ 157.7, 152.4, 140.2, 125.4, 52.7, 45.6, 33.3, 23.9, 22.9. HRMS (ESI-TOF): calcd for $C_{12}H_{20}ClN_4$ $[M+H]^+$ 255.1371, found 255.1373.

*2-chloro-$N^4$-cyclohexyl-$N^5$-isopropylpyrimidine-4,5-diamine*

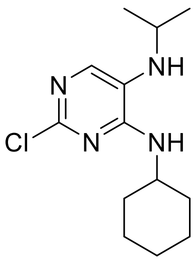


Yield 73 %, pale yellow solid. $^{1}$H NMR (400 MHz, $CDCl_3$) $\delta$ 7.49 (s, 1H), 5.15 (d, $J$ = 7.7 Hz, 1H), 4.04 – 3.93 (m, 1H), 3.39 (hept, $J$ = 6.2 Hz, 1H), 2.50 (s, 1H), 2.11 – 1.93 (m, 2H), 1.73 (dt, $J$ = 13.5, 3.6 Hz, 2H), 1.65 (dt, $J$ = 11.4, 3.5 Hz, 1H), 1.50 – 1.36 (m, 2H), 1.26 – 1.19 (m, 2H), 1.18 (d, $J$ = 6.2 Hz, 6H). $^{13}$C NMR (101 MHz, $CDCl_3$) $\delta$ 157.4, 152.6, 140.7, 125.2, 77.5, 49.3, 45.8, 33.2, 25.8, 24.9, 22.9. HRMS (ESI-TOF): calcd for $C_{13}H_{22}ClN_4$ $[M+H]^+$ 269.1528, found 269.1529.

*2-chloro-$N^5$-isopropyl-$N^4$-(4-methylcyclohexyl)pyrimidine-4,5-diamine*

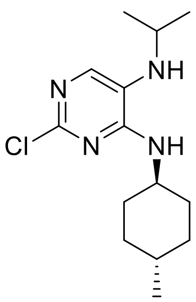


Yield 60 %, pale yellow solid. $^{1}$H NMR (400 MHz, $CDCl_3$) $\delta$ 7.48 (s, 1H), 5.08 (d, $J$ = 7.8 Hz, 1H), 3.93 (ddd, $J$ = 11.5, 7.8, 3.8 Hz, 1H), 3.39 (hept, $J$ = 6.2 Hz, 1H), 2.53 (s, 1H), 2.12 – 2.02 (m, 2H), 1.76 –

1.67 (m, 2H), 1.20 – 1.08 (m, 4H), 1.17 (d, *J* = 6.2 Hz, 6H), 0.91 (d, *J* = 6.5 Hz, 3H). $^{13}C$ NMR (101 MHz, $CDCl_3$) $\delta$ 157.3, 152.4, 140.4, 125.3, 49.7, 45.6, 33.9, 33.2, 32.2, 22.9, 22.3. HRMS (ESI-TOF): calcd for $C_{14}H_{24}ClN_4$ $[M+H]^+$ 283.1684, found 283.1682.

*2-chloro-$N^4$-cycloheptyl-$N^5$-isopropylpyrimidine-4,5-diamine*

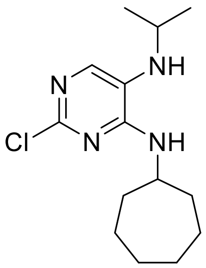


Yield 72 %, pale yellow solid. $^1H$ NMR (400 MHz, $CDCl_3$) $\delta$ 7.48 (s, 1H), 5.24 (d, *J* = 7.7 Hz, 1H), 4.23 – 4.10 (m, 1H), 3.38 (hept, *J* = 6.2 Hz, 1H), 2.55 (s, 1H), 2.09 – 1.89 (m, 2H), 1.71 – 1.39 (m, 11H), 1.17 (d, *J* = 6.2 Hz, 6H).$^{13}C$ NMR (101 MHz, $CDCl_3$) $\delta$ 157.09, 152.55, 140.50, 125.28, 51.49, 45.74, 34.98, 28.36, 24.09, 22.93. HRMS (ESI-TOF): calcd for $C_{14}H_{24}ClN_4$ $[M+H]^+$ 283.1684, found 283.1685.

**Method iv**

0.75 mmol of compound **4** (**$R^1$ = H, $R^2$ = cycloalkyl**) was weight into a flask, toluene (1 mL) and trifluoroacetic anhydride (3 mL) were added and the mixture was heated under a condenser at 70°C for 16 hours. The crude mixture was diluted with dichloromethane (20 mL), washed with 10% $K_2CO_3$ (30 mL), distilled water (30 mL), dried over $MgSO_4$, and evaporated.

*2-chloro-9-cyclopentyl-8-(trifluoromethyl)-9H-purine*

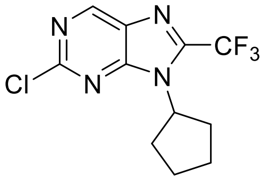


Yield 83 %, light brown solid. $^1H$ NMR (400 MHz, $CDCl_3$) $\delta$ 9.16 (s, 1H), 4.93 (p, *J* = 8.4 Hz, 1H), 2.54 – 2.40 (m, 2H), 2.26 – 2.07 (m, 4H), 1.87 – 1.59 (m, 2H). $^{13}C$ NMR (101 MHz, $CDCl_3$) $\delta$ 155.47, 153.66, 152.48, 144.39 (q, *J* = 39.6 Hz), 131.56, 118.37 (q, *J* = 272.6 Hz), 59.30, 31.33, 25.02. HRMS (ESI-TOF): calcd for $C_{11}H_{11}ClF_3N_4$ $[M+H]^+$ 291.0619, found 291.0619.

*2-chloro-9-cyclohexyl-8-(trifluoromethyl)-9H-purine*

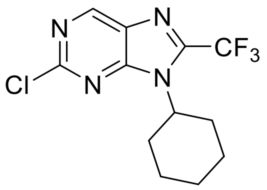

Yield 86 %, white solid. $^1$H NMR (400 MHz, $CDCl_3$) $\delta$ 9.14 (s, 1H), 4.42 (tt, $J$ = 12.1, 3.7 Hz, 1H), 2.61 (qd, $J$ = 12.2, 6.0 Hz, 2H), 2.04 – 1.88 (m, 4H), 1.82 – 1.75 (m, 1H), 1.54 – 1.35 (m, 3H). $^{13}$C NMR (101 MHz, $CDCl_3$) $\delta$ 155.42, 154.00, 152.50, 143.81 (q, $J$ = 39.7 Hz), 118.39 (q, $J$ = 272.5 Hz), 59.6, 30.9, 25.9, 24.8. HRMS (ESI-TOF): calcd for $C_{12}H_{16}ClN_4O$ [M+H]$^+$ 267.1007, found 267.1008. HRMS (ESI-TOF): calcd for $C_{12}H_{16}ClN_4O$ [M+H]$^+$ 305.0775, found 305.0778.

*2-chloro-9-cycloheptyl-8-(trifluoromethyl)-9H-purine*

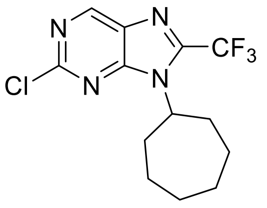

Yield 91 %, colourless oil. $^1$H NMR (400 MHz, $CDCl_3$) $\delta$ 9.14 (s, 1H), 4.59 (t, $J$ = 10.8 Hz, 1H), 2.83 – 2.60 (m, 2H), 2.15 – 1.89 (m, 4H), 1.82 – 1.66 (m, 4H), 1.66 – 1.48 (m, 2H). $^{13}$C NMR (101 MHz, $CDCl_3$) $\delta$ 155.6, 153.8, 152.5, 143.4 (q, $J$ = 39.6 Hz), 131.4, 118.4 (q, $J$ = 272.5 Hz), 61.5, 33.7, 27.3, 25.3. HRMS (ESI-TOF): calcd for $C_{13}H_{15}ClF3N_4$ [M+H]$^+$ 319.0932, found 319.0930.

*2-chloro-9-(trans-4-methylcyclohexyl)-8-(trifluoromethyl)-9H-purine*

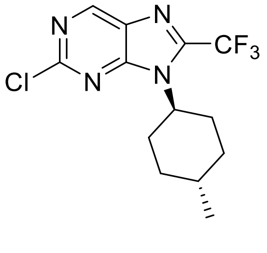

Yield 90%, colourless oil. $^1$H NMR (400 MHz, $CDCl_3$) $\delta$ 9.18 (s, 1H), 4.42 (tt, $J$ = 12.3, 3.6 Hz, 1H), 2.68 (qd, $J$ = 12.4, 3.8 Hz, 2H), 1.99 – 1.89 (m, 4H), 1.23 – 1.09 (m, 2H), 0.99 (d, $J$ = 6.5 Hz, 3H). $^{13}$C NMR (101 MHz, $CDCl_3$) $\delta$ 155.3, 154.0, 152.4, 144.08 (q, $J$ = 39.9 Hz), 131.3, 118.15 (qd, $J$ = 272.6 Hz), 59.6, 35.0, 31.4, 30.6, 22.1. HRMS (ESI-TOF): calcd for $C_{13}H_{15}ClF3N_4$ [M+H]$^+$ 319.0932, found 319.0927.

*2-chloro-9-cyclopentyl-7,9-dihydro-8H-purin-8-one* [3]

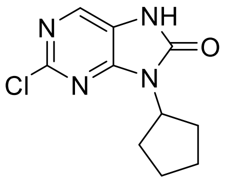

Yield 46 %, white solid. $^1$H NMR (400 MHz, DMSO-$d_6$) $\delta$ 11.57 (s, 1H), 8.10 (s, 1H), 4.69 (p, $J$ = 8.3 Hz, 1H), 3.31 (s, 1H), 2.19 – 2.02 (m, 2H), 2.01 – 1.80 (m, 4H), 1.65 – 1.55 (m, 2H). $^{13}$C NMR (101 MHz, DMSO-$d_6$) $\delta$ 152.7, 151.6, 150.0, 133.8, 121.2, 52.3, 28.9, 24.5. HRMS (ESI-TOF): calcd for $C_{10}H_{12}ClN_4O$ [M+H]$^+$ 239.0694, found 239.0694.

*2-chloro-9-cyclohexyl-7,9-dihydro-8H-purin-8-one* [3]

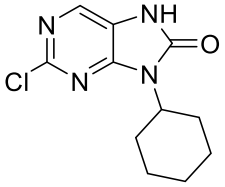

Yield 43 %, white solid. $^1$H NMR (400 MHz, DMSO-$d_6$) $\delta$ 11.52 (s, 1H), 8.07 (s, 1H), 4.10 (tt, $J$ = 12.3, 3.8 Hz, 1H), 2.13 (qd, $J$ = 12.6, 3.3 Hz, 2H), 1.83 – 1.60 (m, 5H), 1.40 – 1.23 (m, 3H), 1.22 – 1.12 (m, 1H). $^{13}$C NMR (101 MHz, DMSO-$d_6$) $\delta$ 152.5, 151.6, 150.0, 133.9, 121.1, 52.0, 29.1, 25.2, 24.8. HRMS (ESI-TOF): calcd for $C_{11}H_{14}ClN_4O$ [M+H]$^+$ 253.0851, found 253.0851.

*2-chloro-9-cycloheptyl-7,9-dihydro-8H-purin-8-one*

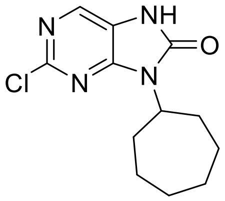

Yield 45 %, white solid. $^1$H NMR (400 MHz, DMSO-$d_6$) $\delta$ 11.54 (s, 1H), 8.10 (s, 1H), 4.30 (tt, $J$ = 10.8, 4.0 Hz, 1H), 2.29 – 2.15 (m, 2H), 1.90 – 1.72 (m, 4H), 1.70 – 1.39 (m, 6H). $^{13}$C NMR (101 MHz, DMSO-$d_6$) $\delta$ 152.4, 151.4, 150.1, 133.8, 121.1, 53.9, 31.9, 27.2, 24.6. HRMS (ESI-TOF): calcd for $C_{12}H_{16}ClN_4O$ [M+H]$^+$ 267.1007, found 267.1009.

*2-chloro-9-(trans-4-methylcyclohexyl)-7,9-dihydro-8H-purin-8-one*

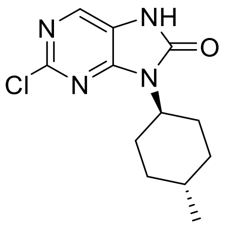

Yield 34%, white solid. $^{1}$H NMR (400 MHz, DMSO-$d_6$) $\delta$ 11.57 (s, 1H), 8.11 (s, 1H), 4.12 (tt, $J$ = 12.2, 3.7 Hz, 1H), 2.23 (qd, $J$ = 12.5, 2.8 Hz, 2H), 1.84 – 1.66 (m, 4H), 1.16 – 1.01 (m, 2H), 0.91 (d, $J$ = 6.5 Hz, 3H). $^{13}$C NMR (101 MHz, DMSO-$d_6$) $\delta$ 152.5, 151.7, 150.0, 133.9, 121.1, 51.9, 33.7, 31.2, 28.8, 22.1. HRMS (ESI-TOF): calcd for $C_{12}H_{16}ClN_4O$ $[M+H]^+$ 267.1007, found 267.1008.

**Method v**

0.70 mmol of compound **4**, **5** or **6** was dissolved in anhydrous THF (10 mL) under an inert atmosphere of nitrogen and cooled to -10°C. After slow addition of 15 wt % $COCl_2$ in toluene (560 µL, 0.85 mmol), 1M LiHMDS in hexane (1.4 mL, 1.40 mmol) was added portionwise. After 1 hour, the reaction mixture was cooled to ambient temperature and stirred for 20 minutes. The mixture was then evaporated under reduced pressure and purified by column chromatography (2:1 EtOAc/hexane) to yield **8-10**.

*2-chloro-9-cyclopropyl-7-methyl-7,9-dihydro-8H-purin-8-one*

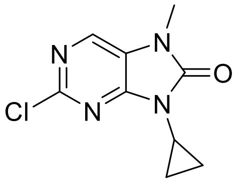

Yield 55 %, white solid. $^{1}$H NMR (400 MHz, $CDCl_3$) $\delta$ 7.98 (s, 1H), 3.42 (s, 3H), 3.03 – 2.97 (m, 1H), 1.18 – 1.12 (m, 4H). $^{13}$C NMR (101 MHz, $CDCl_3$) $\delta$ 153.3, 152.7, 1521., 132.6, 122.2, 27.6, 23.2, 5.9. HRMS (ESI-TOF): calcd for $C_9H_{10}ClN_4O$ $[M+H]^+$ 225.0538, found 225.0537.

*2-chloro-9-cyclobutyl-7-methyl-7,9-dihydro-8H-purin-8-one*

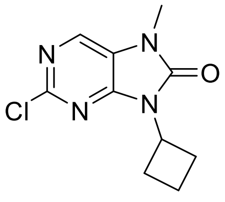

Yield 77 %, white solid. $^{1}$H NMR (500 MHz, DMSO-$d_6$) $\delta$ 8.33 (s, 1H), 4.87 – 4.79 (m, 1H), 2.97 – 2.84 (m, 2H), 2.24 (dtt, $J$ = 13.4, 8.1, 2.5 Hz, 2H), 1.91 – 1.82 (m, 1H), 1.79 (tdd, $J$ = 10.5, 8.1, 2.4 Hz, 1H). $^{13}$C NMR (126 MHz, DMSO-$d_6$) $\delta$ 152.8, 151.3, 150.8, 134.2, 123.2, 46.8, 27.9, 27.6, 15.2. HRMS (ESI-TOF): calcd for $C_{10}H_{12}ClN_4O$ $[M+H]^+$ 239.0695, found 239.0695.

*2-chloro-9-cyclopentyl-7-methyl-7,9-dihydro-8H-purin-8-one* [3]

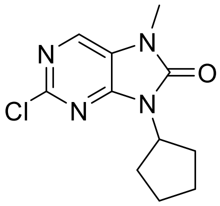


Yield 46 %, white solid. $^{1}$H NMR (400 MHz, $CDCl_3$) $\delta$ 7.97 (s, 1H), 4.84 (p, $J$ = 8.4 Hz, 1H), 3.42 (s, 3H), 2.30 – 2.11 (m, 2H), 2.08 – 1.89 (m, 4H), 1.74 – 1.53 (m, 2H). $^{13}$C NMR (101 MHz, $CDCl_3$) $\delta$ 152.7, 152.3, 151.2, 132.4, 122.3, 53.9, 29.5, 27.5, 24.9. HRMS (ESI-TOF): calcd for $C_{11}H_{14}ClN_4O$ $[M+H]^+$ 253.0851, found 253.0852.

*2-chloro-9-cyclohexyl-7-methyl-7,9-dihydro-8H-purin-8-one* [3]

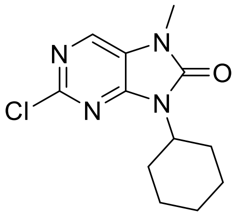


Yield 75 %, light yellow solid. $^{1}$H NMR (400 MHz, $CDCl_3$) $\delta$ 7.96 (s, 1H), 4.33 (tt, $J$ = 12.4, 3.9 Hz, 1H), 3.41 (s, 3H), 2.31 (qd, $J$ = 12.6, 3.7 Hz, 2H), 1.91 – 1.82 (m, 2H), 1.81 – 1.73 (m, 2H), 1.50 – 1.29 (m, 4H). $^{13}$C NMR (101 MHz, $CDCl_3$) $\delta$ 152.7, 152.3, 151.2, 132.4, 122.3, 53.6, 29.7, 27.6, 25.8, 25.0. HRMS (ESI-TOF): calcd for $C_{12}H_{16}ClN_4O$ $[M+H]^+$ 267.1007, found 267.1010.

*2-chloro-9-(trans-4-hydroxycyclohexyl)-7-methyl-7,9-dihydro-8H-purin-8-one* [3]

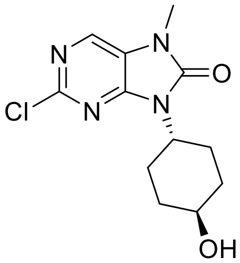

Yield 77 %, light yellow solid. $^{1}H$ NMR (400 MHz, DMSO-$d_6$) $\delta$ 8.33 (s, 1H), 4.67 (s, 1H), 4.15 (tt, $J$ = 12.3, 3.9 Hz, 1H), 3.47 (tq, $J$ = 10.7, 3.8 Hz, 1H), 3.34 (s, 3H), 2.24 (qd, $J$ = 13.4, 3.2 Hz, 2H), 1.97 – 1.88 (m, 2H), 1.77 – 1.66 (m, 2H), 1.36 – 1.23 (m, 2H). $^{13}C$ NMR (101 MHz, DMSO-$d_6$) $\delta$ 152.0, 150.4, 150.2, 133.7, 122.7, 67.9, 51.9, 34.3, 27.4, 27.1. $C_{12}H_{16}ClN_4O$ [M+H]$^+$ 267.1007, found 267.1010. HRMS (ESI-TOF): calcd for $C_{12}H_{16}ClN_4O_2$ [M+H]$^+$ 283.0956, found 283.0954.

*2-chloro-N$^5$-methyl-N$^4$-(trans-4-methylcyclohexyl)pyrimidine-4,5-diamine*

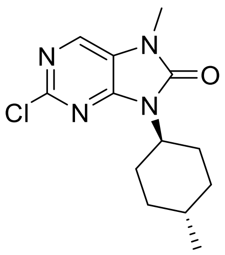


Yield 81 %, white solid. $^{1}H$ NMR (400 MHz, $CDCl_3$) $\delta$ 7.96 (s, 1H), 4.32 (tt, $J$ = 12.4, 4.0 Hz, 1H), 3.41 (s, 3H), 2.38 (qd, $J$ = 12.7, 3.5 Hz, 2H), 1.89 – 1.70 (m, 4H), 1.12 (qd, $J$ = 13.3, 3.3 Hz, 2H), 0.93 (d, $J$ = 6.5 Hz, 3H). $^{13}C$ NMR (101 MHz, $CDCl_3$) $\delta$ 152.7, 152.3, 151.2, 132.3, 122.3, 53.5, 34.3, 31.5, 29.4, 27.5, 22.3. HRMS (ESI-TOF): calcd for $C_{13}H_{18}ClN_4O$ [M+H]$^+$ 281.1164, found 281.1163.

*2-chloro-9-cycloheptyl-7-methyl-7,9-dihydro-8H-purin-8-one*

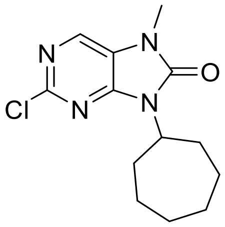


Yield 84 %, pale yellow solid. $^{1}H$ NMR (400 MHz, $CDCl_3$) $\delta$ 7.96 (s, 1H), 4.49 (tt, $J$ = 11.0, 3.8 Hz, 1H), 3.42 (s, 3H), 2.45 – 2.27 (m, 2H), 1.93 – 1.78 (m, 4H), 1.73 – 1.63 (m, 4H), 1.61 – 1.46 (m, 2H). $^{13}C$ NMR (101 MHz, $CDCl_3$) $\delta$ 152.6, 152.4, 150.9, 132.3, 122.3, 55.7, 32.6, 27.6, 27.5, 25.2. HRMS (ESI-TOF): calcd for $C_{13}H_{18}ClN_4O$ [M+H]$^+$ 281.1164, found 282.1161.

*2-chloro-9-cyclooctyl-7-methyl-7,9-dihydro-8H-purin-8-one*

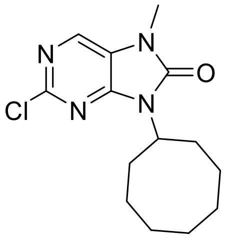

Yield 84 %, white solid. $^{1}H$ NMR (400 MHz, $CDCl_3$) $\delta$ 7.96 (s, 1H), 4.59 (tt, $J$ = 10.5, 3.3 Hz, 1H), 3.44 (d, $J$ = 14.6 Hz, 3H), 2.48 – 2.37 (m, 2H), 1.90 – 1.74 (m, 4H), 1.68 – 1.53 (m, 8H). $^{13}C$ NMR (101 MHz, $CDCl_3$) $\delta$ 152.6, 152.4, 150.9, 132.2, 122.2, 54.6, 31.4, 27.5, 26.5, 25.9, 24.9. HRMS (ESI-TOF): calcd for $C_{14}H_{20}ClN_4O$ $[M+H]^+$ 295.1320, found 295.1323.

*2-chloro-9-cyclododecyl-7-methyl-7,9-dihydro-8H-purin-8-one*

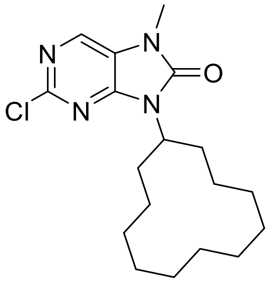

Yield 69 %, white solid. $^{1}H$ NMR (400 MHz, $CDCl_3$) $\delta$ 7.97 (s, 1H), 4.65 (p, $J$ = 6.9 Hz, 1H), 3.43 (s, 3H), 2.31 – 2.13 (m, 2H), 1.93 – 1.76 (m, 2H), 1.65 – 1.48 (m, 3H), 1.39 (t, $J$ = 11.0 Hz, 15H). $^{13}C$ NMR (101 MHz, $CDCl_3$) $\delta$ 153.2, 152.4, 151.4, 132.3, 122.3, 50.3, 28.2, 27.6, 24.3, 24.1, 22.7, 22.7, 22.7. HRMS (ESI-TOF): calcd for $C_{18}H_{28}ClN_4O$ $[M+H]^+$ 351.1946, found 351.1947.

*2-chloro-7-methyl-9-(tetrahydro-2H-pyran-4-yl)-7,9-dihydro-8H-purin-8-one* [3]

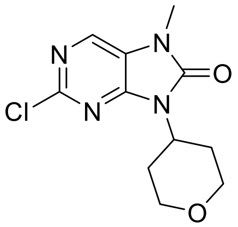

Yield 83 %, pale yellow solid. $^{1}H$ NMR (400 MHz, $CDCl_3$) $\delta$ 8.00 (s, 1H), 4.58 (tt, $J$ = 12.3, 4.2 Hz, 1H), 4.15 – 4.08 (m, 2H), 3.52 (td, $J$ = 12.6, 1.5 Hz, 2H), 3.44 (s, 3H), 2.72 (qd, $J$ = 12.5, 4.5 Hz, 2H), 1.70 (dd, $J$ = 13.1, 3.1 Hz, 2H). $^{13}C$ NMR (126 MHz, $CDCl_3$) $\delta$ 152.5, 151.0, 132.7, 122.3, 67.4, 50.7, 29.8, 27.6. HRMS (ESI-TOF): calcd for $C_{11}H_{14}ClN_4O_2$ $[M+H]^+$ 269.0800, found 269.0801.

*2-chloro-7-methyl-9-(1-methylpiperidin-4-yl)-7,9-dihydro-8H-purin-8-one*

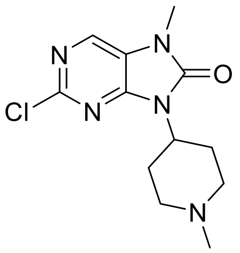

Yield 91 %, White solid. $^1H$ NMR (400 MHz, $CDCl_3$) $\delta$ 7.98 (s, 1H), 4.34 (tt, $J$ = 12.3, 4.2 Hz, 1H), 3.42 (s, 3H), 3.11 – 2.95 (m, 2H), 2.73 (qd, $J$ = 12.6, 4.0 Hz, 2H), 2.35 (s, 3H), 2.14 (td, $J$ = 12.2, 2.2 Hz, 2H), 1.84 – 1.60 (m, 2H). $^{13}C$ NMR (101 MHz, $CDCl_3$) $\delta$ 155.71, 152.97, 150.70, 147.50, 133.15, 132.26, 120.38, 117.39, 116.95, 55.46, 51.48, 50.85, 46.37, 46.32, 28.95, 27.35.HRMS (ESI-TOF): calcd for $C_{12}H_{17}ClN_5O$ $[M+H]^+$ 282.1116, found 282.1116.

*2-chloro-7-methyl-9-phenyl-7,9-dihydro-8H-purin-8-one* [3]

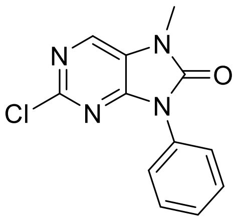

Yield 93 %, white solid. $^1H$ NMR (400 MHz, DMSO-$d_6$) $\delta$ 8.48 (s, 1H), 7.61 – 7.57 (m, 4H), 7.53 – 7.46 (m, 1H), 3.44 (s, 3H). $^{13}C$ NMR (101 MHz, DMSO-$d_6$) $\delta$ 151.9, 150.6, 150.6, 134.4, 132.0, 129.2, 128.5, 126.4, 123.1, 27.7. HRMS (ESI-TOF): calcd for $C_{12}H_{10}ClN_4O$ $[M+H]^+$ 261.0538, found 261.0541.

*2-chloro-7-methyl-9-(p-tolyl)-7,9-dihydro-8H-purin-8-one*

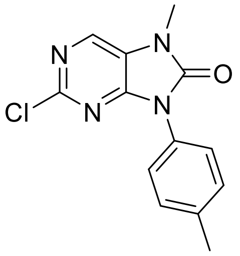

Yield 88 %, light yellow solid. $^1H$ NMR (400 MHz, DMSO-$d_6$) $\delta$ 8.46 (s, 1H), 7.48 – 7.43 (m, 2H), 7.39 – 7.36 (m, 2H), 3.43 (s, 3H), 2.39 (s, 3H). $^{13}C$ NMR (101 MHz, DMSO-$d_6$) $\delta$ 152.0, 150.7, 150.6, 138.2,

134.2, 129.6, 129.4, 126.3, 123.0, 39.7, 27.7, 20.8. HRMS (ESI-TOF): calcd for $C_{13}H_{12}ClN_4O$ $[M+H]^+$ 275.0694, found 275.0692.

*2-chloro-9-(2,4-dimethoxybenzyl)-7-methyl-7,9-dihydro-8H-purin-8-one*

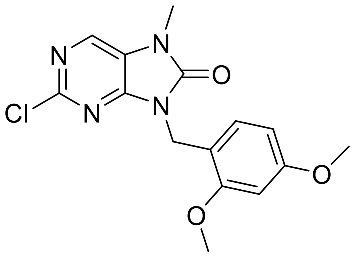

Yield 94 %, white solid. $^1H$ NMR (400 MHz, $CDCl_3$) $\delta$ 7.97 (s, 1H), 7.21 (d, $J$ = 8.9 Hz, 1H), 6.43 – 6.40 (m, 2H), 5.07 (s, 2H), 3.79 (s, 3H), 3.77 (s, 3H), 3.42 (s, 3H). $^{13}C$ NMR (101 MHz, $CDCl_3$) $\delta$ 161.1, 158.6, 153.1, 152.7, 151.4, 132.4, 130.8, 122.5, 115.6, 104.2, 98.7, 55.5, 39.5, 27.6. HRMS (ESI-TOF): calcd for $C_{15}H_{16}ClN_4O_3$ $[M+H]^+$ 335.0905, found 335.0907.

*2-chloro-9-cyclopentyl-7-isopropyl-7,9-dihydro-8H-purin-8-one*

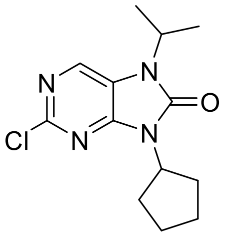

Yiled 92 %, yellow solid. $^1H$ NMR (400 MHz, $CDCl_3$) $\delta$ 8.10 (s, 1H), 4.85 (q, $J$ = 8.1 Hz, 1H), 4.70 (q, $J$ = 7.3 Hz, 1H), 2.29 – 2.15 (m, 2H), 2.06 – 1.93 (m, 4H), 1.73 – 1.62 (m, 2H), 1.50 (s, 6H). $^{13}C$ NMR (101 MHz, $CDCl_3$) $\delta$ 151.9, 151.9, 151.3, 133.4, 120.5, 53.8, 45.7, 29.5, 25.0, 20.7. HRMS (ESI-TOF): calcd for $C_{13}H_{18}ClN_4O$ $[M+H]^+$ 281.1164, found 281.1164.

*2-chloro-9-cyclohexyl-7-isopropyl-7,9-dihydro-8H-purin-8-one*

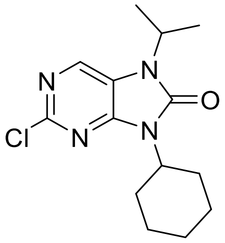

Yield 41 %, pale yellow solid. $^1H$ NMR (400 MHz, $CDCl_3$) $\delta$ 8.09 (s, 1H), 4.71 (hept, $J$ = 7.0 Hz, 1H), 4.33 (tt, $J$ = 12.4, 3.9 Hz, 1H), 2.33 (qd, $J$ = 12.4, 3.2 Hz, 2H), 1.90 (dd, $J$ = 9.6, 6.8 Hz, 2H), 1.83 –

1.75 (m, 2H), 1.74 – 1.67 (m, 1H), 1.50 (s, 6H), 1.47 – 1.28 (m, 3H). $^{13}$C NMR (101 MHz, $CDCl_3$) $\delta$ 151.9, 151.9, 151.3, 133.3, 120.5, 53.6, 45.7, 29.7, 25.9, 25.1, 20.7. HRMS (ESI-TOF): calcd for $C_{14}H_{20}ClN_4O$ $[M+H]^+$ 295.1320, found 295.1321.

*2-chloro-7-isopropyl-9-(trans-4-methylcyclohexyl)-7,9-dihydro-8H-purin-8-one*

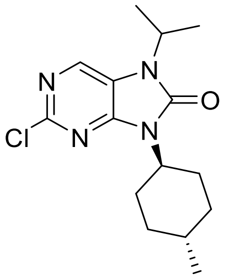


Yield 64 %, yellow oil. $^1$H NMR (400 MHz, $CDCl_3$) $\delta$ 8.08 (s, 1H), 4.69 (hept, $J$ = 7.0 Hz, 1H), 4.30 (tt, $J$ = 12.4, 4.0 Hz, 1H), 2.38 (qd, $J$ = 12.7, 3.5 Hz, 2H), 1.89 – 1.79 (m, 2H), 1.79 – 1.69 (m, 2H), 1.48 (d, $J$ = 7.0 Hz, 6H), 1.10 (qd, $J$ = 13.2, 3.3 Hz, 2H), 0.92 (d, $J$ = 6.5 Hz, 3H). $^{13}$C NMR (101 MHz, $CDCl_3$) $\delta$ 151.9, 151.4, 133.3, 120.5, 53.5, 45.7, 34.3, 31.5, 29.3, 22.3, 20.6. HRMS (ESI-TOF): calcd for $C_{15}H_{22}ClN_4O$ $[M+H]^+$ 309.1477, found 309.1476.

*2-chloro-$N^4$-cycloheptyl-$N_5$-isopropylpyrimidine-4,5-diamine*

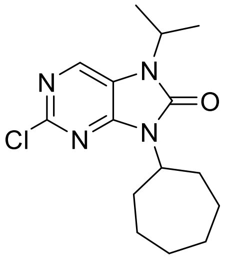


Yield 55 %, pale yellow solid. $^1$H NMR (400 MHz, $CDCl_3$) $\delta$ 8.07 (s, 1H), 4.69 (hept, $J$ = 7.0 Hz, 1H), 4.47 (tt, $J$ = 11.0, 3.8 Hz, 1H), 2.42 – 2.19 (m, 2H), 1.94 – 1.72 (m, 4H), 1.69 – 1.60 (m, 4H), 1.59 – 1.50 (m, 2H), 1.47 (d, $J$ = 7.0 Hz, 6H). $^{13}$C NMR (101 MHz, $CDCl_3$) $\delta$ 152.0, 151.8, 151.0, 133.3, 120.5, 55.6, 45.7, 32.6, 27.6, 25.3, 20.7. HRMS (ESI-TOF): calcd for $C_{15}H_{22}ClN_4O$ $[M+H]^+$ 309.1477, found 309.1476.

***Scheme S2:*** *Synthesis of intermediates for Buchwald Hartwig amination prepared by Chan-Lam reaction.*

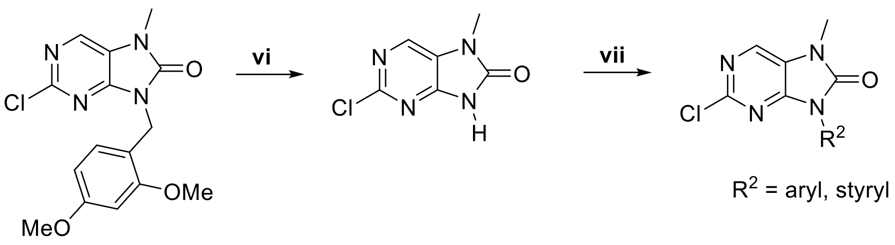


**Method vi**

2-Chloro-9-(2,4-dimethoxybenzyl)-7-methyl-7,9-dihydro-8*H*-purin-8-one (200 mg, 0.59 mmol) was suspended in anisole (3.15mL) and TFA (2.6 mL) was added. The reaction mixture was stirred at 90°C for 18 hours. TFA was evaporated by the flow of nitrogen and anisole was evaporated using RVO. A crude mixture was dissolved in MeOH, solid $NaHCO_3$ was added to neutralize the mixture and evaporated with silica gel. The product was purified by column chromatography (4:1 to 2:1 hexane/EtOAc). After evaporation and trituration with diethylether, the product was filtered-off to yield **7** 2-chloro-7-methyl-7,9-dihydro-8*H*-purin-8-one 56 mg (52%) as a white solid. $^1$H NMR (400 MHz, DMSO-$d_6$) $\delta$ 12.35 (s, 1H), 8.27 (s, 1H), 3.30 (s, 3H) ppm. $^{13}$C NMR (100 MHz, DMSO-$d_6$) $\delta$ 153.1, 151.5, 150.4, 133.3, 123.7, 26.9. HRMS (ESI-TOF): calcd for $C_6H_6ClN_4O$ $[M+H]^+$ 185.0225, found 185.0226. [6]

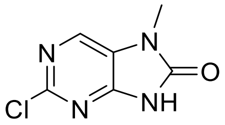

**Method vii**

A 10 mL vial was charged with $Cu_2S$ (8 mg, 0.05 mmol), MeCN (0.6 mL), and *N*,*N*,*N′*,*N′*-tetramethylethane-1,2-diamine (30 μL, 0.20 mmol). After 1h of stirring, 2-chloro-7-methyl-7,9-dihydro-8*H*-purin-8-one (18 mg, 0.10 mmol) and boronic acid (0.20 mmol) were added and the mixture was stirred at ambient temperature for 24 hours. A crude mixture was loaded on Celite, evaporated using RVO, and purified by column chromatography (2:1 hexane/EtOAc).

*2-chloro-7-methyl-9-(o-tolyl)-7,9-dihydro-8H-purin-8-one* [3]

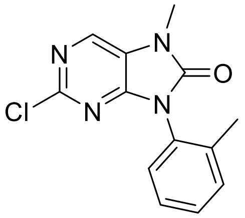

Yield 88 %, pale yellow solid. $^{1}H$ NMR (400 MHz, DMSO-$d_6$) $\delta$ 7.48 – 7.45 (m, 2H), 7.41 – 7.36 (m, 2H), 3.46 (s, 3H), 2.11 (s, 3H). $^{13}C$ NMR (100 MHz, DMSO-$d_6$) $\delta$ 189.4, 188.4, 188.3, 173.9, 172.0, 168.6, 168.4, 167.3, 166.5, 164.5, 160.9, 65.4, 54.9. HRMS (ESI-TOF): calcd for $C_{13}H_{11}ClN_4O$ $[M+H]^+$ 275.0694, found 275.0694.

*2-chloro-9-(4-methoxyphenyl)-7-methyl-7,9-dihydro-8H-purin-8-one* [3]

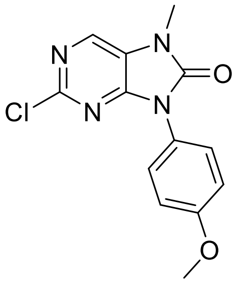

White solid 57%. $^{1}H$ NMR (400 MHz, DMSO-$d_6$) $\delta$ 8.45 (s, 1H), 7.49 – 7.45 (m, 2H), 7.14 – 7.09 (m, 2H), 3.83 (s, 3H), 3.43 (s, 3H). $^{13}C$ NMR (101 MHz, DMSO-$d_6$) $\delta$ 159.1, 152.1, 150.8, 150.6, 134.1, 128.0, 124.5, 123.0, 114.4, 55.5, 27.6. HRMS (ESI-TOF): calcd for $C_{13}H_{11}ClN_4O_2$ $[M+H]^+$ 291.0643, found 291.0640.

2-chloro-9-(2-methoxyphenyl)-7-methyl-7,9-dihydro-8*H*-purin-8-one

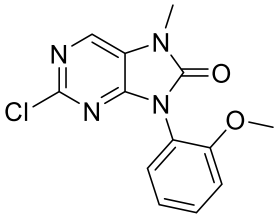

White foam 72%. $^{1}H$ NMR (400 MHz, DMSO-$d_6$) $\delta$ 8.45 (s, 1H), 7.55 (td, $J$ = 8.5, 1.6 Hz, 1H), 7.43 (dd, $J$ = 7.8, 1.6 Hz, 1H), 7.28 (dd, $J$ = 7.8, 0.8 Hz, 1H), 7.13 (td, $J$ = 7.8, 0.8 Hz, 1H), 3.75 (s, 3H), 3.45 (s, 3H). $^{13}C$ NMR (101 MHz, DMSO-$d_6$) $\delta$ = 155.3, 151.8, 151.0, 150.8, 134.3, 131.3, 130.0, 123.0,

120.7, 119.9, 112.8, 55.9, 27.7. HRMS (ESI-TOF): calcd for $C_{13}H_{11}ClN_4O_2$ $[M+H]^+$ 291.0643, found 291.0640.

2-chloro-7-methyl-9-(naphthalen-1-yl)-7,9-dihydro-8*H*-purin-8-one

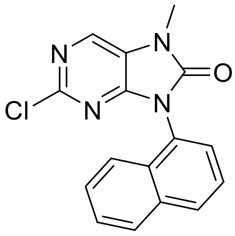

Pale yellow solid 24%. $^1$H NMR (400 MHz, DMSO-$d_6$) $\delta$ 8.54 (s, 1H), 8.19 (d, $J$ = 4.7 Hz, 1H), 8.11 (d, $J$ = 8.1 Hz, 1H), 7.73 - 7.62 (m, 4H), 7.55 (ddd, $J$ = 8.1, 6.9, 1.3 Hz, 1H), 3.51 (s, 3H). $^{13}$C NMR (101 MHz, DMSO-$d_6$) $\delta$ 152.3, 151.6, 150.7, 134.4, 133.9, 130.1, 129.7, 128.4, 128.2, 127.3, 127.3, 126.8, 125.7, 123.6, 122.7, 27.9. HRMS (ESI-TOF): calcd for $C_{16}H_{11}ClN_4O$ $[M+H]^+$ 311.0694, found 311.0692.

(*E*)-2-chloro-9-(4-fluorostyryl)-7-methyl-7,9-dihydro-8*H*-purin-8-one

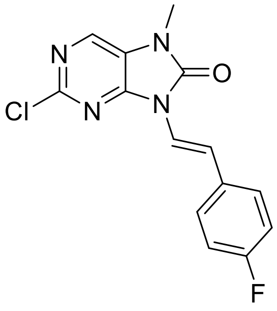

White solid 53%. $^1$H NMR (400 MHz, DMSO-$d_6$) $\delta$ 8.47 (s, 1H), 7.62 – 7.59 (m, 2H), 7.55 (d, $J$ = 15.3 Hz, 1H), 7.37 (d, $J$ = 15.3 Hz, 1H), 7.22 – 7.17 (m, 2H), 3.40 (s, 3H). $^{13}$C NMR (101 MHz, DMSO-$d_6$) $\delta$ 161.7 (d, $J$ = 245 Hz), 150.9, 150.7, 148.9, 134.7, 131.4 (d, $J$ = 3 Hz), 128.2 (d, $J$ = 8 Hz), 122.9, 119.4, 118.3 (d, $J$ = 2 Hz), 115.7 (d, $J$ = 22 Hz), 27.6. HRMS (ESI-TOF): calcd for $C_{14}H_{10}ClFN_4O$ $[M + H]^+$ 305.0600, found 305.060

**Characterization of final compounds 12, 13, 14 and 15.**

*9-Cyclopentyl-N-(4-(piperazin-1-yl)phenyl)-8-(trifluoromethyl)-9H-purin-2-amine (**12a**)*

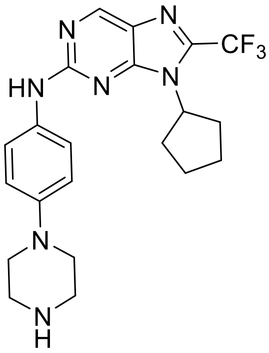


Yield 21 %, yellow solid, m. p. 232-234 °C. HPLC purity: >99.9 %. $^{1}$H NMR (400 MHz, DMSO-$d_6$) $\delta$ 9.59 (s, 2H), 8.95 (s, 1H), 7.73 – 7.57 (m, 2H), 7.16 – 7.00 (m, 2H), 4.80 (p, $J$ = 8.6 Hz, 1H), 3.54 – 3.33 (m, 4H), 3.32 – 3.19 (m, 4H), 2.45 – 2.32 (m, 2H), 2.10 – 1.92 (m, 4H), 1.73 – 1.62 (m, 2H). $^{13}$C NMR (101 MHz, DMSO-$d_6$) $\delta$ 156.5, 152.5, 151.8, 143.5, 138.6 (q, $J$ = 37.7 Hz), 134.3, 126.0, 120.5, 120.3, 118.7 (q, $J$ = 272.5 Hz), 117.3, 57.5, 46.9, 42.0, 30.0, 24.4. HRMS (ESI): m/z calcd $C_{21}H_{25}F_3N_7$ for $[M+H]^+$ 432.218, found $[M+H]^+$ 432.2117.

*9-Cyclohexyl-N-(4-(piperazin-1-yl)phenyl)-8-(trifluoromethyl)-9H-purin-2-amine (**12b**)*

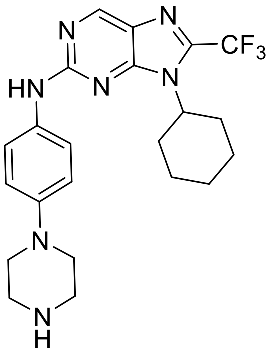


Yield 43 %, yellow solid, m. p. 218-220 °C. HPLC purity: 98.2 %. $^{1}$H NMR (400 MHz, DMSO-$d_6$) $\delta$ 9.71 (s, 1H), 8.93 (s, 1H), 7.71 – 7.60 (m, 2H), 6.94 – 6.82 (m, 2H), 4.22 (t, $J$ = 12.3 Hz, 1H), 3.06 – 2.94 (m, 4H), 2.93 – 2.79 (m, 4H), 2.57 (qd, $J$ = 11.8, 2.9, 2H), 1.93 – 1.80 (m, 4H), 1.77 – 1.70 (m, 1H), 1.44 – 1.32 (m, 2H), 1.32 – 1.23 (m, 1H). $^{13}$C NMR (101 MHz, DMSO-$d_6$) $\delta$ 156.6, 152.6, 152.5,

144.8, 137.6 (q, $J$ = 38.3 Hz), 133.8, 125.8, 120.0, 118.8 (q, $J$ = 272.8 Hz), 116.8, 57.2, 46.4, 42.5, 29.9, 25.2, 25.0. HRMS (ESI): m/z calcd $C_{22}H_{27}F_3N_7$ for $[M+H]^+$ 446.2275, found $[M+H]^+$ 446.2273.

*9-(trans-4-Methylcyclohexyl)-N-(4-(piperazin-1-yl)phenyl)-8-(trifluoromethyl)-9H-purin-2-amine* ***(12c)***

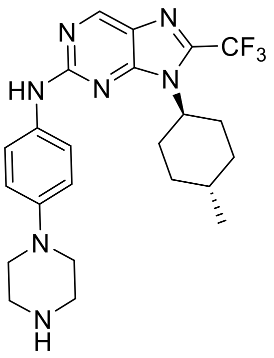

Yield 57 %, yellow solid, m. p. 228-230 °C. HPLC purity: 99.6 %. $^1$H NMR (400 MHz, DMSO-$d_6$) $\delta$ 9.82 (s, 1H), 9.37 (s, 2H), 8.99 (s, 1H), 7.82 – 7.66 (m, 2H), 7.04 – 6.92 (m, 2H), 4.24 (tt, $J$ = 12.3, 3.2 Hz, 1H), 3.53 (s, 3H), 3.38 – 3.30 (m, 4H), 3.28 – 3.15 (m, 4H), 2.64 (qd, $J$ = 13.1, 3.6 Hz, 2H), 1.94 – 1.79 (m, 4H), 1.62 – 1.45 (m,1.2 1H), 1.15 (qd, $J$ = 12.5, 12.3, 2.5 Hz, 2H), 0.98 (d, $J$ = 6.5 Hz, 3H). $^{13}$C NMR (101 MHz, DMSO-$d_6$) $\delta$ 156.7, 152.6, 152.5, 145.0, 137.5 (q, $J$ = 38.3 Hz), 133.6, 125.8, 120.1, 118.8 (q, $J$ = 271.6 Hz), 116.8, 57.1, 46.4, 42.6, 33.6, 31.4, 29.6, 22.1. HRMS (ESI): m/z calcd $C_{23}H_{29}F_3N_7$ for $[M+H]^+$ 460.2431, found $[M+H]^+$ 460.2429.

*9-Cycloheptyl-N-(4-(piperazin-1-yl)phenyl)-8-(trifluoromethyl)-9H-purin-2-amine* ***(12d)***

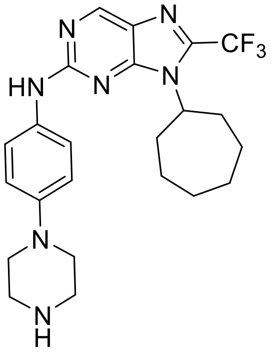

Yield 34 %, yellow solid, m. p. 216-218 °C. HPLC purity: 98.30 %. $^1$H NMR (400 MHz, DMSO-$d_6$) $\delta$ 9.74 (s, 1H), 8.97 (s, 1H), 7.76 – 7.64 (m, 2H), 6.99 – 6.86 (m, 2H), 5.99 (br. s, 1H), 4.48 – 4.32 (m,

1H), 3.10 (s, 4H), 2.99 (s, 4H), 2.68 – 2.55 (m, 2H), 2.02 – 1.92 (m, 2H), 1.91 – 1.79 (m, 2H), 1.78 – 1.61 (m, 4H), 1.59 – 1.46 (m, 2H). $^{13}$C NMR (126 MHz, DMSO-$d_6$) $\delta$ 156.4, 152.6, 152.1, 142.4, 137.7 (q, $J$ = 37.5 Hz), 135.6, 126.0, 120.2, 118.7 (q, $J$ = 271.2 Hz), 118.1, 59.7, 47.6, 42.0, 32.7, 26.8, 24.2. HRMS (ESI): m/z calcd $C_{23}H_{29}F_3N_7$ for $[M+H]^+$ 460.2431, found $[M+H]^+$ 460.2430.

*9-Cyclopentyl-2-((4-(piperazin-1-yl)phenyl)amino)-7,9-dihydro-8H-purin-8-one (**13a**)*

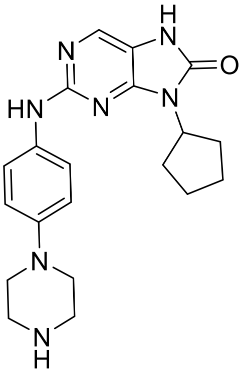

Yield 30 %, pale yellow solid, m. p. 200-202 °C. HPLC purity: 98.2 %. $^1$H NMR (400 MHz, DMSO-$d_6$) $\delta$ 8.94 (s, 1H), 7.89 (s, 1H), 7.63 – 7.51 (m, 2H), 6.92 – 6.82 (m, 2H), 4.66 (p, $J$ = 8.6 Hz, 1H), 3.12 – 2.99 (m, 4H), 2.97 – 2.84 (m, 4H), 2.29 – 2.11 (m, 2H), 2.00 – 1.80 (m, 4H), 1.70 – 1.54 (m, 2H). $^{13}$C NMR (101 MHz, DMSO-$d_6$) $\delta$ 155.5, 153.4, 151.1, 146.3, 134.3, 134.3, 120.0, 116.7, 114.9, 52.4, 49.8, 45.3, 29.2, 25.0. HRMS (ESI): m/z calcd $C_{20}H_{26}N_7O$ for $[M+H]^+$ 380.2193, found $[M+H]^+$ 380.2192.

*9-Cyclohexyl-2-((4-(piperazin-1-yl)phenyl)amino)-7,9-dihydro-8H-purin-8-one (**13b**)*

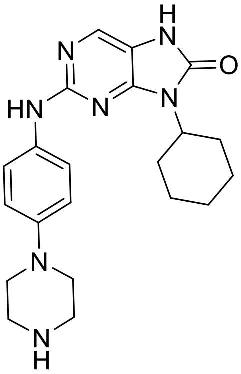

Yield 76 %, pale yellow solid, m. p. 210-212 °C. HPLC purity: 99.8 %. $^{1}$H NMR (400 MHz, DMSO-$d_6$) $\delta$ 8.93 (s, 1H), 7.84 (s, 1H), 7.59 – 7.51 (m, 2H), 6.90 – 6.72 (m, 2H), 4.08 (tt, $J$ = 12.1, 3.5 Hz, 1H), 3.06 – 2.94 (m, 4H), 2.92 – 2.91 (m, 4H), 2.27 (qd, $J$ = 13.0, 3.2 Hz, 2H), 1.90 – 1.74 (m, 2H), 1.72 – 1.59 (m, 3H), 1.36 – 1.25 (m, 2H), 1.21 – 1.16 (m, 1H). $^{13}$C NMR (101 MHz, DMSO-$d_6$) $\delta$ 154.9, 152.7, 150.6, 145.8, 133.8, 133.8, 119.3, 116.2, 114.3, 51.2, 49.4, 44.9, 29.2, 25.4, 25.0. HRMS (ESI): m/z calcd $C_{21}H_{28}N_7O$ for $[M+H]^+$ 394.2350, found $[M+H]^+$ 394.2349.

*9-(trans-4-Methylcyclohexyl)-2-((4-(piperazin-1-yl)phenyl)amino)-7,9-dihydro-8H-purin-8-one*

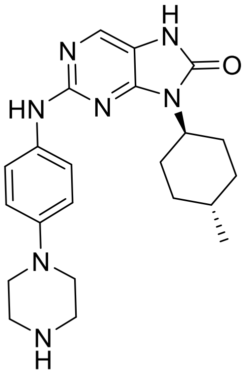


Yield 46 %, pale yellow solid, m. p. 274-276 °C. HPLC purity: >99.9 %. $^{1}$H NMR (400 MHz, DMSO-$d_6$) $\delta$ 9.00 (s, 1H), 7.88 (s, 1H), 7.61 (d, $J$ = 9.1 Hz, 2H), 6.88 (d, $J$ = 9.1 Hz, 2H), 4.11 (tt, $J$ = 12.5, 4.1 Hz, 1H), 3.25 – 3.15 (m, 4H), 3.09 (dd, $J$ = 6.4, 3.5 Hz, 4H), 2.37 (qd, $J$ = 12.9, 4.2 Hz, 1H), 1.80 (d, $J$ = 13.1 Hz, 2H), 1.69 (dd, $J$ = 12.3, 2.7 Hz, 2H), 1.53 – 1.39 (m, 1H), 1.06 (qd, $J$ = 12.8, 2.5 Hz, 2H), 0.94 (d, $J$ = 6.5 Hz, 3H). $^{13}$C NMR (126 MHz, DMSO-$d_6$) $\delta$ 154.8, 152.7, 150.7, 144.9, 134.4, 133.7, 119.3, 116.5, 114.4, 51.1, 47.6, 43.5, 34.0, 31.4, 28.9, 22.2. HRMS (ESI): m/z calcd $C_{22}H_{30}N_7O$ for $[M+H]^+$ 408.2506, found $[M+H]^+$ 408.2504.

*9-Cycloheptyl-2-((4-(piperazin-1-yl)phenyl)amino)-7,9-dihydro-8H-purin-8-one (**13d**)*

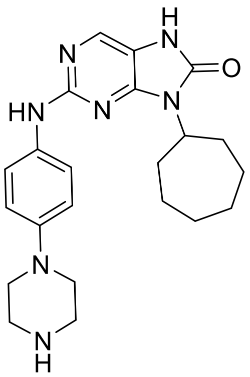

Yield 38 %, pale yellow solid, m. p. 230-232 °C. HPLC purity: 99.8 %. $^{1}$H NMR (400 MHz, DMSO-*d*$_6$) *δ* 11.06 (s, 1H), 9.59 (s, 2H), 9.31 (s, 1H), 7.90 (s, 1H), 7.71 – 7.54 (M, 2H), 7.07 – 6.86 (m, 2H), 4.27 (ddt, *J* = 11.5, 7.9, 3.4 Hz, 1H), 3.41 – 3.25 (m, 4H), 3.23 – 3.05 (m, 4H), 2.31 (q, *J* = 10.9 Hz, 2H), 1.85 – 1.72 (m, 4H), 1.70 – 1.54 (m, 4H), 1.53 – 1.39 (m, 2H). $^{13}$C NMR (101 MHz, DMSO-*d*$_6$) *δ* 153.83, 152.56, 151.20, 144.95, 133.56, 119.99, 116.68, 114.56, 42.53, 31.96, 27.10, 24.50. HRMS (ESI): m/z calcd $C_{22}H_{30}N_7O$ for $[M+H]^+$ 408.2506, found $[M+H]^+$ 408.2505.

*9-Cyclopropyl-7-methyl-2-((4-(piperazin-1-yl)phenyl)amino)-7,9-dihydro-8H-purin-8-one (**14b**)*

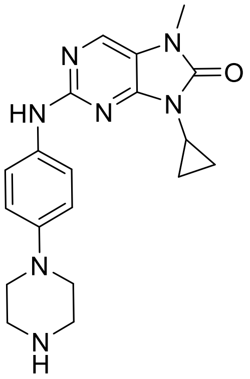

Yeild 26 %, pale yellow solid, m. p. 156-158 °C. HPLC purity: 99.8 %. $^{1}$H NMR (400 MHz, DMSO-*d*$_6$) *δ* 9.04 (s, 1H), 8.03 (s, 1H), 7.66 – 7.57 (m, 2H), 6.90 – 6.80 (m, 2H), 3.26 (s, 3H), 2.98 – 2.93 (m, 4H), 2.92 – 2.89 (m, 1H), 2.87 – 2.78 (m, 4H), 1.09 – 1.04 (m, 2H), 1.03 – 0.94 (m, 2H). $^{13}$C NMR (101 MHz, DMSO-*d*$_6$) *δ* 155.5, 152.9, 150.6, 146.3, 133.5, 133.1, 119.3, 116.0, 115.9, 50.5, 46.7, 27.0, 22.24, 5.4. HRMS (ESI): m/z calcd $C_{19}H_{24}N_7O$ for $[M+H]^+$ 366.2037, found $[M+H]^+$ 366.2036.

*9-Cyclobutyl-7-methyl-2-((4-(piperazin-1-yl)phenyl)amino)-7,9-dihydro-8H-purin-8-one (**14c**)*

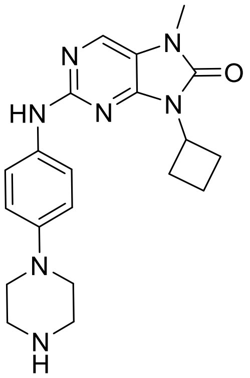

Yield 35 %, white solid, m. p. 112-114 °C. HPLC purity: >99.9 %. $^{1}$H NMR (400 MHz, DMSO-$d_6$) $\delta$ 9.08 – 9.02 (m, 1H), 8.07 (s, 1H), 7.65 – 7.54 (m, 2H), 6.91 – 6.80 (m, 2H), 4.82 (p, $J$ = 8.8 Hz, 1H), 3.34 – 3.31 (m, 2H), 3.27 (s, 3H), 3.03 (pd, $J$ = 9.9, 1.7 Hz, 2H) 3.11 – 2.97 (m, 2H), 2.95 – 2.92 (m, 2H), 2.91 – 2.84 (m, 2H), 2.84 – 2. 80 (m, 2H), 2.23 (qt, $J$ = 8.3, 2.3 Hz, 2H), 1.95 – 1.75 (m, 2H). $^{13}$C NMR (101 MHz, DMSO-$d_6$) $\delta$ 155.3, 152.1, 149.7, 146.4, 146.4, 133.7, 133.3, 119.4, 116.2, 115.9, 115.8, 50.4, 50.1, 45.7, 45.7, 44.4, 39.9, 27.1, 27.0, 14.6. HRMS (ESI): m/z calcd $C_{20}H_{26}N_7O$ for $[M+H]^+$ 380.2193, found $[M+H]^+$ 380.1353.

*9-Cyclopentyl-7-methyl-2-((4-(piperazin-1-yl)phenyl)amino)-7,9-dihydro-8H-purin-8-one (**14d**)*

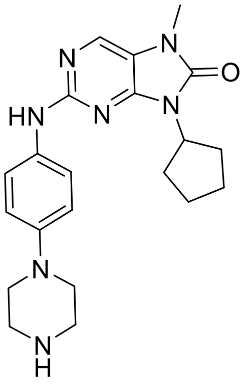

Yield 47 %, pale yellow solid, m. p. 110-112 °C. HPLC purity: 99.8 %. $^{1}$H NMR (400 MHz, DMSO-$d_6$) $\delta$ 9.01 (s, 1H), 8.06 (s, 1H), 7.61 – 7.51 (m, 2H), 6.88 – 6.80 (m, 2H), 4.70 (p, $J$ = 8.4 Hz, 1H), 3.29 (s, 3H), 3.06 – 2.94 (m, 4H), 2.95 – 2.85 (m, 4H), 2.25 – 2.12 (m, 2H), 2.00 – 1.83 (m, 4), 1.69 – 1.53 (m, 2H). $^{13}$C NMR (126 MHz, DMSO-$d_6$) $\delta$ 155.2, 152.2, 149.4, 146.0, 133.4, 119.5, 116.1, 52.4, 49.7, 45.1, 28.8, 27.0, 24.4. HRMS (ESI): m/z calcd $C_{21}H_{28}N_7O$ for $[M+H]^+$ 394.2350, found $[M+H]^+$ 394.2350.

*9-Cyclohexyl-7-methyl-2-((4-(piperazin-1-yl)phenyl)amino)-7,9-dihydro-8H-purin-8-one (**14e**)*

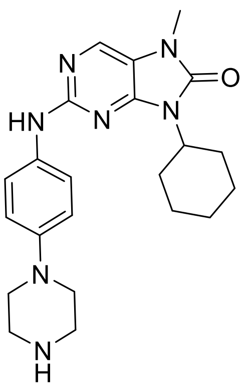


Yield 56 %, pale yellow solid, m. p. 198-200 °C. HPLC purity: 98.1 %. $^{1}$H NMR (400 MHz, DMSO-*d*$_6$) *δ* 9.00 (s, 1H), 8.02 (s, 1H), 7.58 – 7.51 (m, 2H), 6.83 – 6.77 (m, 2H), 4.12 (tt, *J* = 13.2, 3.7 Hz, 1H), 3.24 (s, 3H), 2.94 – 2.87 (m, 4H), 2.87 – 2.68 (m, 4H), 2.26 (qd, *J* = 12.9, 3.1 Hz, 2H), 1.87 – 1.76 (m, 2H), 1.72 – 1.61 (m, 3H), 1.40 – 1.10 (m, 3H). $^{13}$C NMR (101 MHz, DMSO-*d*$_6$) *δ* 155.2, 152.0, 149.4, 146.4, 133.4, 133.4, 119.4, 115.9, 51.7, 50.5, 45.7, 29.3, 27.0, 25.4, 25.0. HRMS (ESI): m/z calcd $C_{22}H_{30}N_7O$ for $[M+H]^+$ 408.2506, found $[M+H]^+$ 408.2503.

*9-(trans-4-Hydroxycyclohexyl)-7-methyl-2-((4-(piperazin-1-yl)phenyl)amino)-7,9-dihydro-8H-purin-8-one (**14f**)*

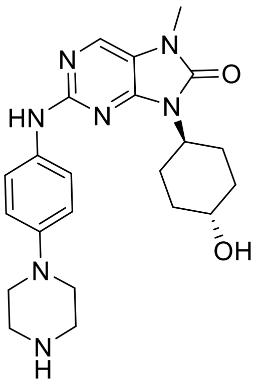


Yield 46 %, yellow solid, m. p. 272-274 °C. HPLC purity: 98.5 %. $^{1}$H NMR (400 MHz, DMSO-*d*$_6$) *δ* 9.03 (s, 1H), 8.06 (s, 1H), 7.60 – 7.55 (m, 2H), 6.87 – 6.81 (m, 2H), 4.71 (s, 1H), 4.14 (tt, *J* = 12.3, 4.1 Hz, 1H), 3.58 – 3.47 (m, 1H), 3.28 (s, 3H) 2.97 – 2.83 (m, 4H), 2.86 – 2.80 (m, 4H), 2.38 (qd, *J* = 13.0, 3.1 Hz, 2H), 2.01 – 1.91 (m, 2H), 1.74 – 1.65 (m, 2H), 1.37 – 1.23 (m, 2H). $^{13}$C NMR (101 MHz,

DMSO-$d_6$) $\delta$ 155.2, 152.0, 149.4, 146.3, 133.5, 133.4, 119.4, 116.0, 68.2, 51.1, 50.4, 45.6, 34.6, 27.0, 27.0. HRMS (ESI): m/z calcd $C_{22}H_{30}N_7O_2$ for $[M+H]^+$ 424.2455, found $[M+H]^+$ 424.2452.

*7-Methyl-9-(trans-4-methylcyclohexyl)-2-((4-(piperazin-1-yl)phenyl)amino)-7,9-dihydro-8H-purin-8-one (**14g**)*

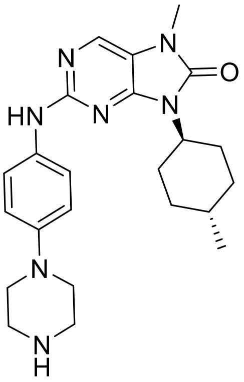

Yield 32 %, white solid, m. p. 114-116 °C. HPLC purity: 99.3 %. $^1H$ NMR (400 MHz, DMSO-$d_6$) $\delta$ 9.00 (s, 1H), 8.05 (s, 1H), 7.61 – 7.53 (m, 2H), 6.86 – 6.80 (m, 2H), 4.14 (tt, $J$ = 12.2, 3.9 Hz, 1H), 3.28 (s, 3H), 2.97 – 2.91 (m, 4H), 2.85 – 2.79 (m, 4H), 2.36 (qd, $J$ = 12.6, 3.2 Hz, 2H), 1.85 – 1.77 (m, 2H), 1.75 – 1.63 (m, 2H), 1.07 (qd, 12.3, 3.2 Hz, 2H), 0.94 (d, $J$ = 6.5 Hz, 3H). $^{13}C$ NMR (101 MHz, DMSO-$d_6$) $\delta$ 155.2, 152.1, 149.5, 146.4, 133.4, 119.4, 116.0, 115.9, 51.6, 50.5, 45.7, 33.9, 31.4, 28.9, 27.0, 22.2. HRMS (ESI): m/z calcd $C_{23}H_{32}N_7O$ for $[M+H]^+$ 422.2663, found $[M+H]^+$ 422.2661.

*9-Cycloheptyl-7-methyl-2-((4-(piperazin-1-yl)phenyl)amino)-7,9-dihydro-8H-purin-8-one (**14h**)*

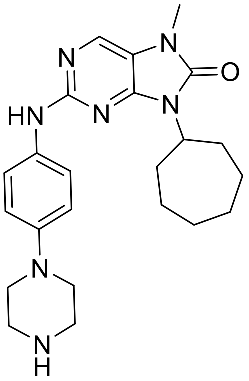

Yield 23 %, pale yellow solid, m. p. 256-258 °C. HPLC purity: 98.9 %. $^1H$ NMR (400 MHz, DMSO-$d_6$) $\delta$ 8.97 (s, 1H), 8.01 (s, 1H), 7.59 – 7.50 (m, 2H), 6.84 – 6.77 (m, 2H), 4.27 (tt, $J$ = 11.0, 3.7 Hz,

1H), 3.24 (s, 3H), 2.96 – 2.92 (m, 4H), 2.85 – 2. 81 (m, 4H), 2.29 (qd, *J* = 10.8, 3.7 Hz, 2H), 1.82 – 1.71 (m, 4H), 1.68 – 1.49 (m, 4H), 1.51 – 1.37 (m, 2H). $^{13}$C NMR (101 MHz, DMSO-$d_6$) $\delta$ 155.3, 151.9, 149.2, 146.1, 133.5, 133.4, 119.5, 116.0, 53.9, 50.0, 45.4, 32.1, 27.1, 27.0, 24.5. HRMS (ESI): m/z calcd $C_{23}H_{32}N_7O$ for $[M+H]^+$ 422.2663, found $[M+H]^+$ 422.2188.

*9-Cyclooctyl-7-methyl-2-((4-(piperazin-1-yl)phenyl)amino)-7,9-dihydro-8H-purin-8-one (**14i**)*

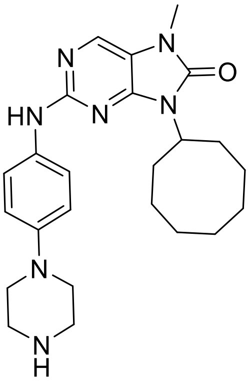


Yield 55 %, pale yellow solid, m. p. 122-124 °C. HPLC purity: 98.7 %. $^1$H NMR (400 MHz, DMSO-$d_6$) $\delta$ 9.00 (s, 1H), 8.05 (s, 1H), 7.62 – 7.55 (m, 2H), 6.87 – 6.79 (m, 2H), 4.40 (tt, *J* = 10.5, 3.0 Hz, 1H), 3.28 (s, 3H), 3.01 – 2.92 (m, 4H), 2.87 – 2.85 (m, 4H), 2.47 – 2.30 (m, 2H), 1.85 – 1.68 (m, 5H), 1.63 – 1.48 (m, 7H). $^{13}$C NMR (101 MHz, DMSO-$d_6$) δ 155.3, 151.9, 149.2, 146.2, 133.5, 133.4, 119.5, 115.9, 52.8, 50.2, 45.5, 30.7, 27.0, 26.2, 24.9, 24.3. HRMS (ESI): m/z calcd $C_{24}H_{34}N_7O$ for $[M+H]^+$ 436.2819, found $[M+H]^+$ 436.2818.

*9-Cyclododecyl-7-methyl-2-((4-(piperazin-1-yl)phenyl)amino)-7,9-dihydro-8H-purin-8-one (**14j**)*

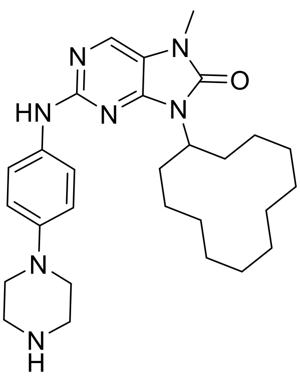

Yield 61 %, white solid, m. p. 114 °C. HPLC purity: >99.9 %. $^{1}$H NMR (400 MHz, DMSO-$d_6$) $\delta$ 9.03 (s, 1H), 8.07 (s, 1H), 7.62 – 7.56 (m, 2H), 6.85 – 6.79 (m, 2H), 4.55 (p, $J$ = 6.7 Hz, 1H), 3.29 (s, 3H), 3.00 – 2.89 (m, 4H), 2.89 – 2.77 (m, 4H), 2.16 – 2.09 (m, 2H), 1.85 – 1.78 (m, 2H), 1.63 – 1.49 (m, 3H), 1.43 – 1.31 (m, 15H). $^{13}$C NMR (101 MHz, DMSO-$d_6$) $\delta$ 155.4, 152.4, 149.7, 146.1, 133.7, 133.4, 119.2, 115.9, 115.7, 50.2, 47.7, 45.6, 27.5, 27.0, 23.8, 23.6, 22.1, 22.1, 22.0. HRMS (ESI): m/z calcd $C_{28}H_{42}N_7O$ for $[M+H]^+$ 492.3445, found $[M+H]^+$ 492.3445.

*7-Methyl-2-((4-(piperazin-1-yl)phenyl)amino)-9-(tetrahydro-2H-pyran-4-yl)-7,9-dihydro-8H-purin-8-one (**14k**)*

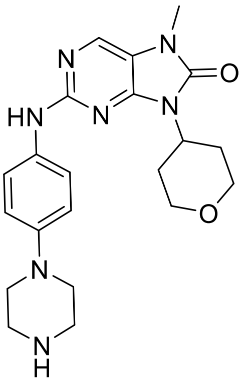


Yield 64 %, pale yellow solid, m. p. 124-126 °C. HPLC purity: > 99.9 %. $^{1}$H NMR (400 MHz, DMSO-$d_6$) $\delta$ 9.05 (s, 1H), 8.07 (s, 1H), 7.63 – 7.55 (m, 2H), 6.89 – 6.80 (m, 2H), 4.41 (tt, $J$ = 12.1, 4.2 Hz, 1H), 3.99 (dd, $J$ = 11.3, 4.2 Hz, 2H), 3.42 – 3.38 (m, 2H), 3.29 (s, 3H), 3.02 – 2.88 (m, 5H), 2.93 – 2.70 (m, 3H), 2.57 (qd, 11.6, 3.4 Hz, 2H), 1.66 (dd, $J$ = 12.7, 2.9 Hz, 2H). $^{13}$C NMR (101 MHz, DMSO-$d_6$) $\delta$ 155.2, 152.0, 149.6, 146.3, 133.4, 119.4, 116.1, 116.0, 66.6, 50.4, 49.1, 45.6, 29.4, 27.1. HRMS (ESI): m/z calcd $C_{21}H_{28}N_7O$ for $[M+H]^+$ 410.2299, found $[M+H]^+$ 410.2298.

*7-Methyl-9-(1-methylpiperidin-4-yl)-2-((4-(piperazin-1-yl)phenyl)amino)-7,9-dihydro-8H-purin-8-one (**14l**)*

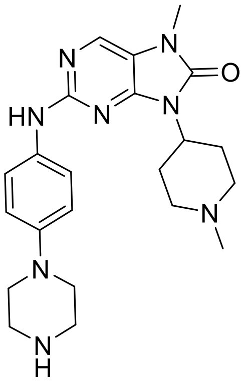

Yield 29 %, pale yellow solid, m. p. decomposition at 282 °C: HPLC purity: 98.7 %. $^{1}$H NMR (400 MHz, $CDCl_3$) $\delta$ 7.81 (s, 1H), 7.50 – 7.45 (m, 2H), 6.94 – 6.89 (m, 2H), 6.87 (s, 1H), 4.26 (tt, $J$ = 12.3, 4.2 Hz, 1H), 3.35 (s, 3H), 3.10 – 3.06 (m, 4H), 3.05 – 3.02 (m, 4H), 2.98 (dt, $J$ = 12.0, 1.5 Hz, 2H), 2.77 (qd, $J$ = 12.5, 3.9 Hz, 2H), 2.33 (s, 3H), 2.09 (td, $J$ = 12.2, 2.1 Hz, 2H), 1.79 – 1.66 (m, 2H). $^{13}$C NMR (101 MHz, $CDCl_3$) $\delta$ 155.7, 153.0, 150.7, 147.5, 133.2, 132.3, 120.4, 117.4, 117.0, 55.5, 51.5, 50.9, 46.4, 46.3, 29.0, 27.4. HRMS (ESI): m/z calcd $C_{22}H_{31}N_8O$ for $[M+H]^+$ 423.2615, found $[M+H]^+$ 423.2605.

*7-Methyl-9-phenyl-2-((4-(piperazin-1-yl)phenyl)amino)-7,9-dihydro-8H-purin-8-one (**14m**)*

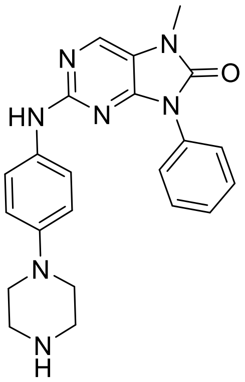

Yield 35 %, yellow solid, m. p. 120-122 °C. HPLC purity: 99.2 %. $^{1}$H NMR (400 MHz, DMSO-$d_6$) $\delta$ 9.09 (s, 1H), 8.19 (s, 1H), 7.68 – 7.62 (m, 2H), 7.61 – 7.51 (m, 4H), 7.47 – 7.43 (m, 1H), 6.84 – 6.76 (m, 2H), 3.38 (s, 3H), 2.96 – 2.87 (m, 4H), 2.86 – 2.75 (m, 4H). $^{13}$C NMR (101 MHz, DMSO-$d_6$) $\delta$ 155.6, 151.7, 149.7, 146.4, 134.0, 133.3, 133.0, 128.9, 127.8, 126. 4, 119.4, 116.3, 115.9, 50.5, 45.7, 27.3. HRMS (ESI): m/z calcd $C_{22}H_{24}N_7O$ for $[M+H]^+$ 402.2037, found $[M+H]^+$ 402.2035.

*7-Methyl-2-((4-(piperazin-1-yl)phenyl)amino)-9-(o-tolyl)-7,9-dihydro-8H-purin-8-one (**14n**)*

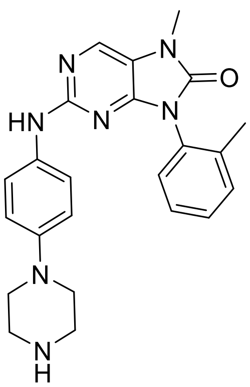

Yield 44 %, yellow solid, m. p. 152-154 °C. HPLC purity: 98.4 %. $^{1}$H NMR (400 MHz, DMSO-$d_6$) $\delta$ 9.11 (s, 1H), 8.18 (s, 1H), 7.55 – 7.51 (m, 2H), 7.46 – 7.34 (m, 4H), 6.82 – 6.77 (m, 2H), 3.3 (s, 3H), 3.04 – 3.00 (m, 4H), 2.97 – 2.92 (m, 4H), 2.13 (s, 3H). $^{13}$C NMR (100 MHz, DMSO-$d_6$) $\delta$ 155.6, 151.7, 150.3, 145.7, 136.5, 133.8, 133.5, 131.8, 130.8, 129.3, 129.2, 126.8, 119.3, 116.7, 116.2, 49.0, 44.6, 27.3, 17.4. HRMS (ESI): m/z calcd $C_{23}H_{27}N_7O$ for $[M+H]^+$ 416.2193, found $[M+H]^+$ 416.2190.

*7-Methyl-2-((4-(piperazin-1-yl)phenyl)amino)-9-(p-tolyl)-7,9-dihydro-8H-purin-8-one (**14o**)*

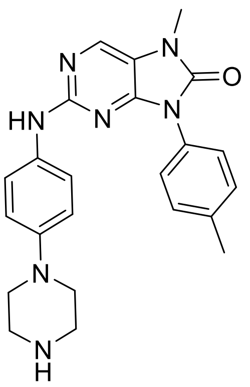

Yield 56 %, pale yellow solid, m. p. 126-128 °C. HPLC purity: 99.8 %. $^{1}$H NMR (400 MHz, DMSO-$d_6$) $\delta$ 9.07 (s, 1H), 8.17 (s, 1H), 7.57 – 7.47 (m, 4H), 7.39 – 7.32 (m, 2H), 6.83 – 6.76 (m, 2H), 3.37 (s, 3H), 2.92 – 2.91 (m, 4H), 287 – 2.73 (m, 4H), 2.39 (s, 3H). $^{13}$C NMR (101 MHz, DMSO-$d_6$) $\delta$ 151.9, 151.9, 149.9, 146.4, 137.4, 133.8, 133.33, 130.4, 129.4, 126.3, 119.4, 116.3, 116.0, 50.5, 45.7, 27.3, 20.8. HRMS (ESI): m/z calcd $C_{23}H_{26}N_7O$ for $[M+H]^+$ 416.2193, found $[M+H]^+$ 415.2190.

*9-(2-Methoxyphenyl)-7-methyl-2-((4-(piperazin-1-yl)phenyl)amino)-7,9-dihydro-8H-purin-8-one* ***(14p)***

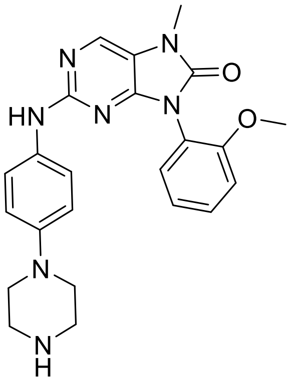

Yield 28 %, pale yellow solid, m. p. 185-187 °C. HPLC purity: 99.3 %. $^{1}$H NMR (400 MHz, DMSO-$d_6$) $\delta$ 9.14 (s, 1H), 8.15 (s, 1H), 7.58 - 7.54 (m, 2H), 7.51 (ddd, $J$ = 8.2, 7.5, 1.5 Hz, 1H), 7.41 (dd, $J$ = 7.5, 1.5 Hz, 1H), 7.25 (dd, $J$ = 8.2, 1.5 Hz, 1H), 7.10 (td, $J$ = 7.5 Hz, 1H), 3.75 (s, 3H), 3.38 (s, 3H), 3.20 - 3.15 (m, 4H), 3.13 - 3.09 (m, 4H). $^{13}$C NMR (101 MHz, DMSO-$d_6$) $\delta$ 155.6, 155.5, 151.9, 150.6, 144.8, 134.3, 133.3, 130.9, 130.4, 121.0, 120.6, 119.2, 116.7, 116.6, 112.6, 55.8, 47.2, 43.2, 27.3. HRMS (ESI): m/z calcd $C_{23}H_{25}N_7O_2$ for 432.2142, found $[M+H]^+$ 432.2140.

*9-(4-Methoxyphenyl)-7-methyl-2-((4-(piperazin-1-yl)phenyl)amino)-7,9-dihydro-8H-purin-8-one* ***(14q)***

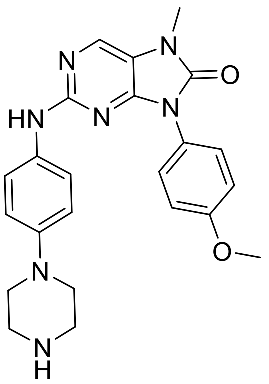

Yield 21 %, pale yellow solid, m. p. 270-272°C. HPLC purity: 99.4 %. $^{1}$H NMR (400 MHz, DMSO-$d_6$) $\delta$ 9.09 (s, 1H), 8.17 (s, 1H), 7.59 – 7.54 (m, 2H), 7.53 – 7.50 (m, 2H), 7.13 – 7.09 (m, 2H), 6.86 – 6.81 (m, 2H), 3.83 (s, 3H), 3.37 (s, 3H), 3.10 – 3.05 (m, 4H), 3.01 – 2.97 (m, 4H).$^{13}$C NMR (100

MHz, DMSO-$d_6$) $\delta$ 158.7, 155.5, 152.0, 150.0, 145.4, 133.9, 133.7, 128.0, 125.5, 119.3, 116.4, 116.3, 114.2, 55.5, 48.5, 44.2, 27.3. HRMS (ESI): m/z calcd $C_{23}H_{25}N_7O_2$ for $[M+H]^+$ 432.2142, found $[M+H]^+$ 432.2136.

*9-(2,4-Dimethoxybenzyl)-7-methyl-2-((4-(piperazin-1-yl)phenyl)amino)-7,9-dihydro-8H-purin-8-one* ***(14r)***

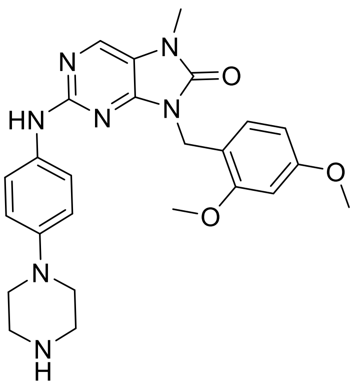

Yield 54 %, white solid, m. p. 102-104 °C. HPLC purity: 99.6 %. $^1$H NMR (400 MHz, DMSO-$d_6$) $\delta$ 9.02 (s, 1H), 8.09 (s, 1H), 7.56 – 7.49 (m, 2H), 6.84 (d, $J$ = 8.4 Hz, 1H), 6.82 – 6.77 (m, 2H), 6.58 (d, $J$ = 2.4 Hz, 1H), 6.43 (dd, $J$ = 8.4, 2.4 Hz, 1H), 4.88 (s, 2H), 3.81 (s, 3H), 3.72 (s, 3H), 3.33 (s, 3H), 2.94 – 2.91 (m, 4H), 2.83 – 2.80 (m, 4H). $^{13}$C NMR (101 MHz, DMSO-$d_6$) $\delta$ 159.9, 157.4, 155.5, 152.7, 150.0, 146.3, 133.3, 133.3, 127.8, 119.3, 116.4, 116.0, 115.9, 104.5, 98.3, 55.5, 55.2, 45.7, 37.8, 27.2. HRMS (ESI): m/z calcd $C_{25}H_{30}N_7O_3$ for $[M+H]^+$ 476.24059, found $[M+H]^+$ 476.2403.

*7-Methyl-9-(naphthalen-1-yl)-2-((4-(piperazin-1-yl)phenyl)amino)-7,9-dihydro-8H-purin-8-one* ***(14s)***

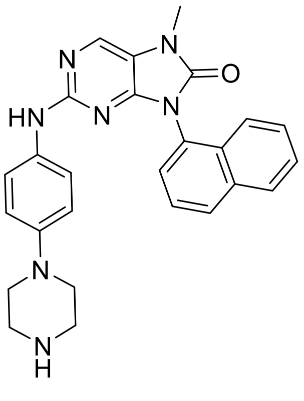

Yield 17 %, pale yellow solid, m. p. 178-180 °C. HPLC purity: 98.3 %. $^1$H NMR (400 MHz, DMSO-*d*$_6$) $\delta$ 9.00 (s, 1H), 8.24 (s, 1H), 8.15 - 8.11 (m, 1H), 8.11 - 8.07 (m, 1H), 7.72 - 7.69 (m, 1H), 7.69 - 7.67 (m, 1H), 7.64 - 7.62 (m, 1H), 7.60 (s, 1H), 7.57 - 7.53 (m, 1H), 7.47 - 7.43 (m, 2H), 6.75 - 6.71 (m, 2H), 3.45 (s, 3H), 3.41 (br. s, 1H), 2.97 - 2.93 (m, 4H), 2.91 - 2.87 (m, 4H). $^{13}$C NMR (101 MHz, DMSO-*d*$_6$) $\delta$ 155.6, 152.3, 151.1, 145.8, 133.9, 133.6, 133.5, 130.1, 129.6, 129.3, 128.3, 127.5, 127.2, 126.7, 125.8, 122.8, 119.2, 116.9, 116.0, 49.5, 45.0, 27.5. HRMS (ESI): m/z calcd $C_{26}H_{25}N_7O$ for $[M+H]^+$ 452.2193, found $[M+H]^+$ 452.2190.

*(E)-9-(4-Fluorostyryl)-7-methyl-2-((4-(piperazin-1-yl)phenyl)amino)-7,9-dihydro-8H-purin-8-one* ***(14t)***

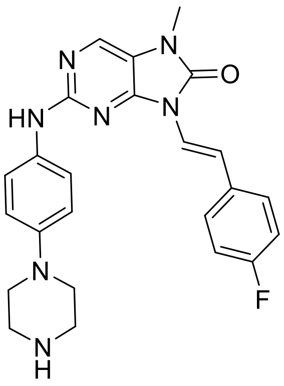


Yield 23 %, pale yellow solid, m. p. 248-250 °C. HPLC purity: 96.2 %. $^1$H (400 MHz, DMSO-*d*$_6$) $\delta$ 9.17 (s, 1H), 8.16 (s, 1H), 7.72 (d, $J$ = 14.8 Hz, 1H), 7.53 – 7.48 (m, 4H), 7.37 (d, $J$ = 14.8 Hz, 1H), 7.22 – 7.16 (m, 2H), 6.89 – 6.85 (m, 2H), 3.31 (s, 3H), 3.31 (br. s, 1H), 3.01 – 2.96 (m, 4H), 2.89 – 2.85 (m, 4H). $^{13}$C (101 MHz, DMSO-*d*$_6$) $\delta$ 161.4 (d, $J$ = 240 Hz), 155.7, 150.6, 148.1, 146.4, 134.4, 132.8, 131.9 (d, $J$ = 3 Hz), 127.4 (d, $J$ = 8 Hz), 120.3, 118.8, 117.7, 116.0, 115.8, 115.5 (d, $J$ = 22 Hz), 49.6, 45.1, 27.1. HRMS (ESI): m/z calcd $C_{24}H_{24}FN_7O$ for $[M + H]^+$ 446.2095, found $[M + H]^+$ 446.2099.

*9-Cyclopentyl-7-isopropyl-2-((4-(piperazin-1-yl)phenyl)amino)-7,9-dihydro-8H-purin-8-one* ***(15a)***

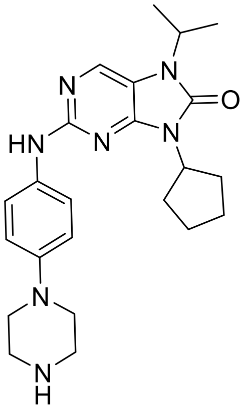


Yield 20 %, white solid, m. p. 130-132 °C. HPLC purity: 98.6 %. $^{1}$H NMR (400 MHz, DMSO-$d_6$) $\delta$ 8.99 (s, 1H), 8.24 (s, 1H), 7.58 – 7.51 (m, 2H), 6.87 – 6.81 (m, 2H), 4.70 (p, $J$ = 8.5 Hz, 1H), 4.54 (hept, $J$ = 7.1 Hz, 1H), 2.99 – 2.92 (m, 4H), 2.88 – 2.81 (m, 4H), 2.27 – 2.12 (m, 2H), 1.98 – 1.83 (m, 4H), 1.69 – 1.56 (m, 2H), 1.41 (d, $J$ = 6.9 Hz, 6H). $^{13}$C NMR (101 MHz, DMSO-$d_6$) $\delta$ 154.8, 151.2, 149.6, 146.3, 134.3, 133.3, 119.5, 115.9, 113.9, 52.2, 50.3, 45.6, 44.5, 28.7, 24.4, 20.1. HRMS (ESI): m/z calcd $C_{23}H_{31}N_7O$ for $[M+H]^+$ 422.2663, found $[M+H]^+$ 422.2661.

*9-Cyclohexyl-7-isopropyl-2-((4-(piperazin-1-yl)phenyl)amino)-7,9-dihydro-8H-purin-8-one (**15b**)*

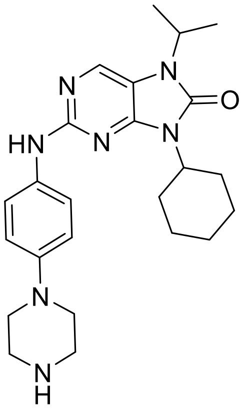


Yield 27 %, white solid, m. p. 132-134 °C. HPLC purity: 98.4 %. $^{1}$H NMR (400 MHz, DMSO-$d_6$) $\delta$ 9.02 (s, 1H), 8.23 (s, 1H), 7.62 – 7.55 (m, 2H), 6.88 – 6.81 (m, 2H), 4.54 (hept, $J$ = 7.0 Hz, 1H), 4.16 (tt, $J$ = 12.2, 3.8 Hz, 1H), 2.99 – 2.91 (m, 4H), 2.87 – 2.79 (m, 4H), 2.31 (qd, $J$ = 12.5, 3.3 Hz, 2H), 1.90 – 1.80 (m, 2H), 1.76 – 1.66 (m, 3H), 1.40 (d, $J$ = 6.9 Hz, 6H), 1.38 – 1.29 (m, 2H), 1.28 – 1.18 (m, 1H). $^{13}$C NMR (101 MHz, DMSO-$d_6$) $\delta$ 154.8, 151.0, 149.6, 146.4, 134.3, 133.4, 119.4, 115.9,

113.8, 51.7, 50.4, 45.7, 44.5, 29.2, 25.4, 25.0, 20.1. HRMS (ESI): m/z calcd $C_{24}H_{33}N_7O$ for $[M+H]^+$ 436.2819, found $[M+H]^+$ 436.2817.

*7-Isopropyl-9-(4-methylcyclohexyl)-2-((4-(piperazin-1-yl)phenyl)amino)-7,9-dihydro-8H-purin-8-one **(15c)***

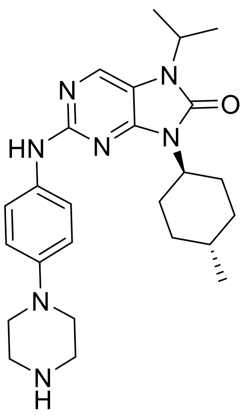

Yield 26 %, pale yellow solid, m. p. 264-266 °C. HPLC purity: 99.4 %. $^1$H NMR (500 MHz, DMSO-$d_6$) $\delta$ 9.08 (s, 1H), 8.24 (s, 1H), 7.66 – 7.60 (m, 2H), 6.93 – 6.85 (m, 2H), 4.53 (hept, $J$ = 6.9 Hz, 1H), 4.14 (tt, $J$ = 12.3, 3.9 Hz, 1H), 3.25 – 3.20 (m, 4H), 3.17 – 3.12 (m, 4H), 2.36 (qd, $J$ = 12.7, 3.3 Hz, 2H), 1.84 – 1.76 (m, 2H), 1.73 – 1.64 (m, 2H), 1.40 (d, $J$ = 6.9 Hz, 6H), 1.06 (qd, $J$ = 13.4, 3.0 Hz, 2H), 0.94 (d, $J$ = 6.5 Hz, 3H). $^{13}$C NMR (126 MHz, DMSO-$d_6$) $\delta$ 154.6, 151.1, 149.7, 144.7, 134.4, 134.2, 119.3, 116.6, 114.0, 51.6, 47.0, 44.5, 43.1, 34.0, 31.34 28.9, 22.2, 20.1. HRMS (ESI): m/z calcd $C_{25}H_{35}N_7O$ for $[M+H]^+$ 450.2976, found $[M+H]^+$ 450.2975.

*9-Cycloheptyl-7-isopropyl-2-((4-(piperazin-1-yl)phenyl)amino)-7,9-dihydro-8H-purin-8-one **(15d)***

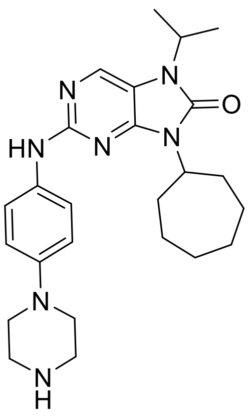

Yield 68 %, pale yellow solid, m. p. 266-268 °C. HPLC purity: 98.6 %. $^{1}$H NMR (500 MHz, DMSO-$d_6$) $\delta$ 9.05 (s, 1H), 8.23 (s, 1H), 7.66 – 7.59 (m, 2H), 6.93 – 6.84 (m, 2H), 4.53 (hept, $J$ = 6.9 Hz, 1H), 4.31 (tt, 10.9, 3.4, 1H), 3.22 – 3.09 (m, 4H), 3.08 – 2.94 (m, 4H), 2.33 (q, $J$ = 10.8 Hz, 2H), 1.85 – 1.72 (m, 4H), 1.71 – 1.54 (m, 4H), 1.52 – 1.43 (m, 2H), 1.40 (d, $J$ = 6.9 Hz, 6H). $^{13}$C NMR (126 MHz, DMSO-$d_6$) $\delta$ 154.7, 151.0, 134.2, 134.1, 149.4, 145.2, 119.4, 116.4, 113.9, 53.9, 48.0, 44.5, 43.9, 32.0, 27.1, 24.5, 20.1. HRMS (ESI): m/z calcd $C_{25}H_{35}N_7O$ for $[M+H]^+$ 450.2976, found $[M+H]^+$ 450.2976.

*7-Methyl-2-((4-(piperazin-1-yl)phenyl)amino)-7,9-dihydro-8H-purin-8-one (**14a**)*

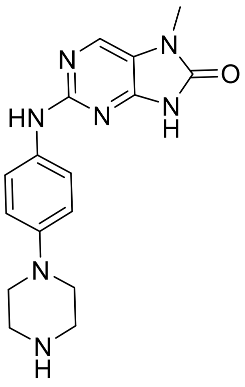


9-(2,4-Dimethoxybenzyl)-7-methyl-2-((4-(piperazin-1-yl)phenyl)amino)-7,9-dihydro-8*H*-purin-8-one (**14r**) (109 mg, 0.23 mmol) was dissolved in anisole (1.2 mL), then trifluoroacetic acid (1 mL) was added and a reaction mixture was heated in a round bottom flask with a condenser at 90°C for 5 days. TFA was evaporated by the flow of nitrogen and anisole was evaporated using RVO. Product was purified by column chromatography (5:1 DCM/MeOH). After evaporation, the product was diluted with $H_2O$/MeOH (2 mL) and filtered off to yield 7-methyl-2-((4-(piperazin-1-yl)phenyl)amino)-7,9-dihydro-8*H*-purin-8-one 26 mg (35 %).

M. p. 232-234 °C. HPLC purity: >99.9 %. $^{1}$H NMR (400 MHz, DMSO-$d_6$) $\delta$ 9.03 (s, 1H), 8.01 (s, 1H), 7.63 – 7.56 (m, 2H), 6.94 – 6.87 (m, 2H), 3.25 (s, 3H), 3.24 – 3.20 (m, 8H). $^{13}$C NMR (101 MHz, DMSO-$d_6$) $\delta$ 155.5, 153.2, 150.5, 144.4, 134.7, 132.8, 119.4, 117.5, 116.8, 46.7, 42.9, 26.6. HRMS (ESI): m/z calcd $C_{16}H_{20}N_7O$ for $[M+H]^+$ 326.1724, found $[M+H]^+$ 326.1724.

[1]H NMR spectrum of **12a**

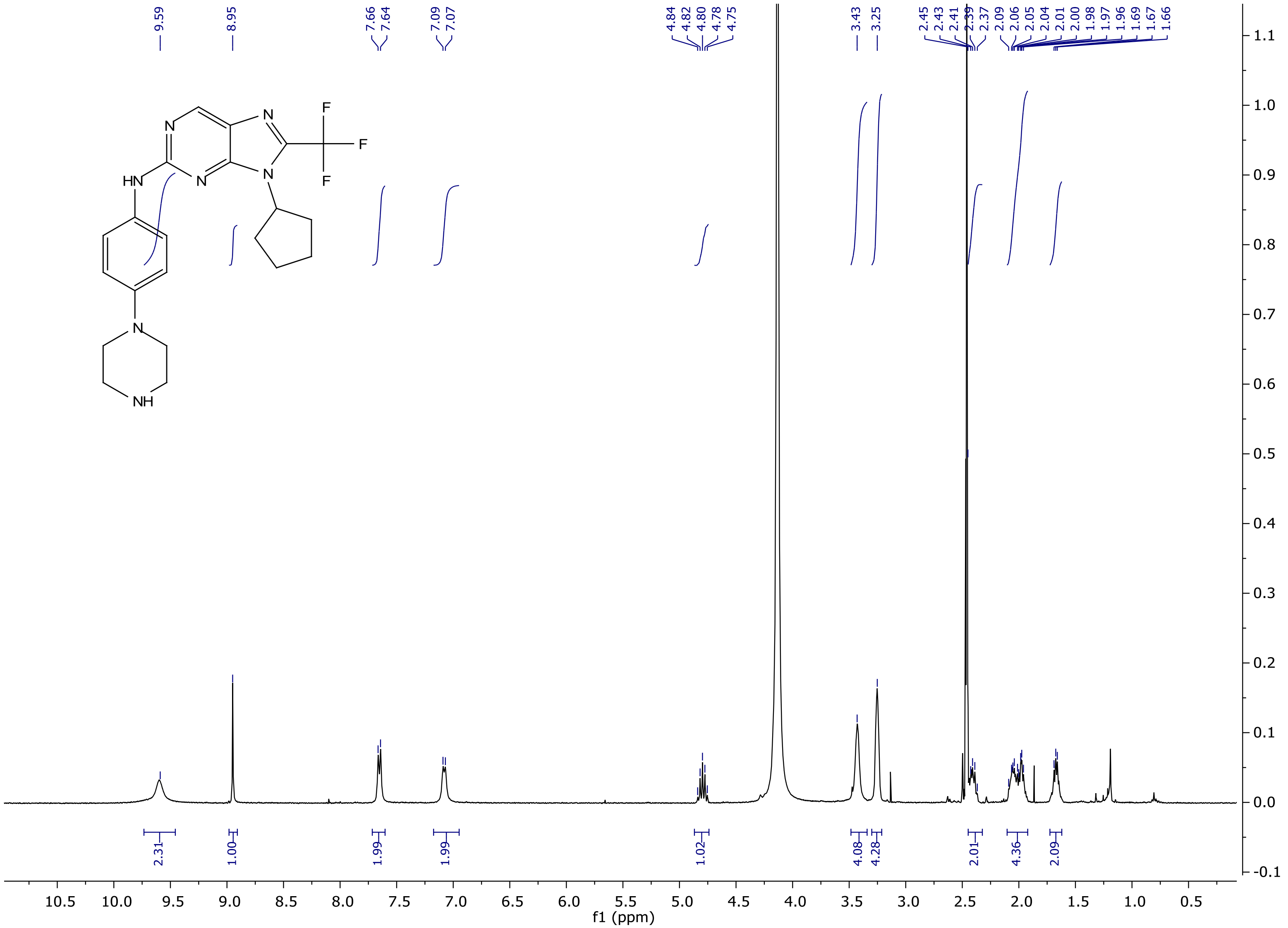

$^{13}C$ NMR spectrum of **12a**

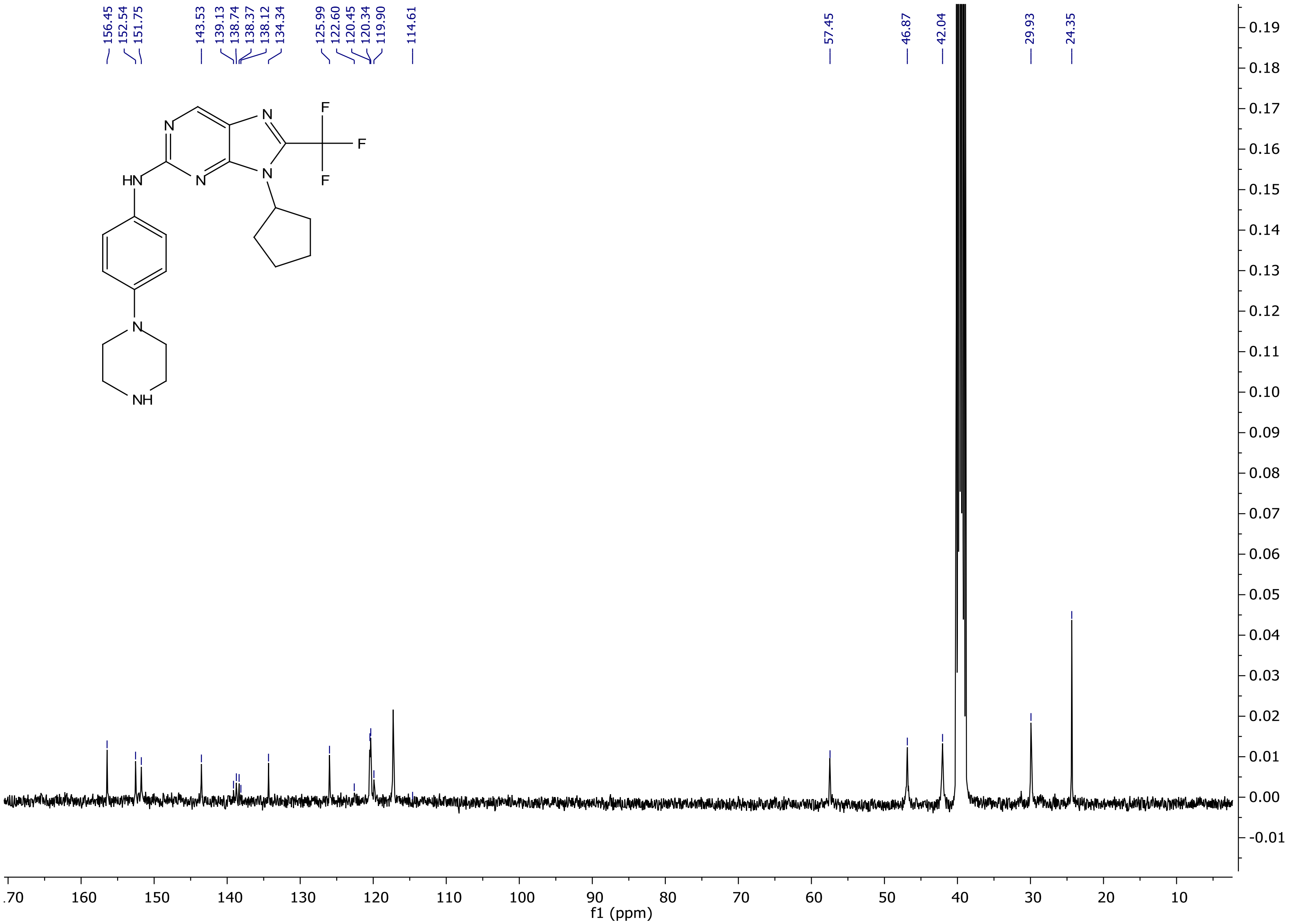

[1]H NMR spectrum of **12b**

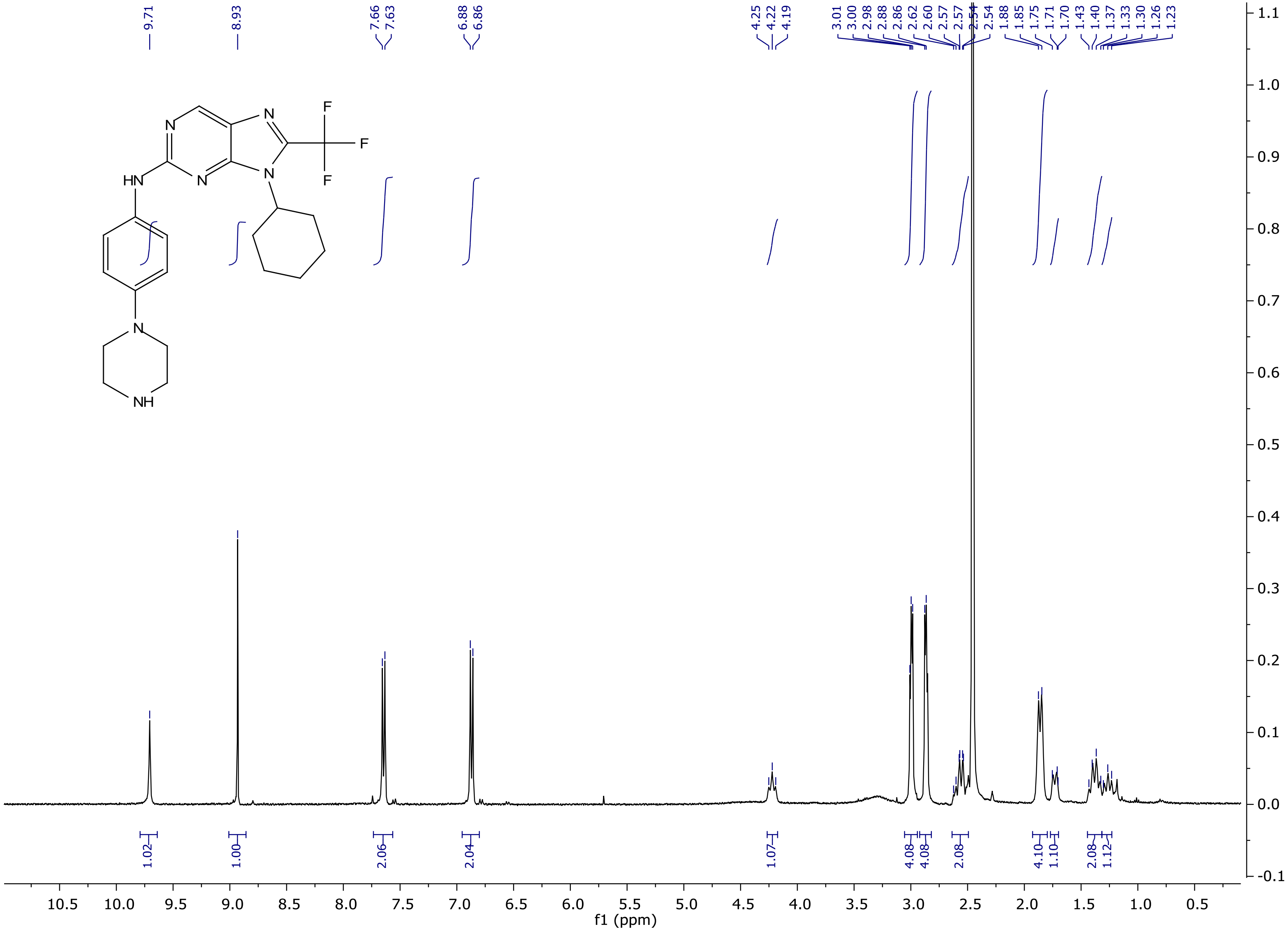

$^{13}$C NMR spectrum of **12b**

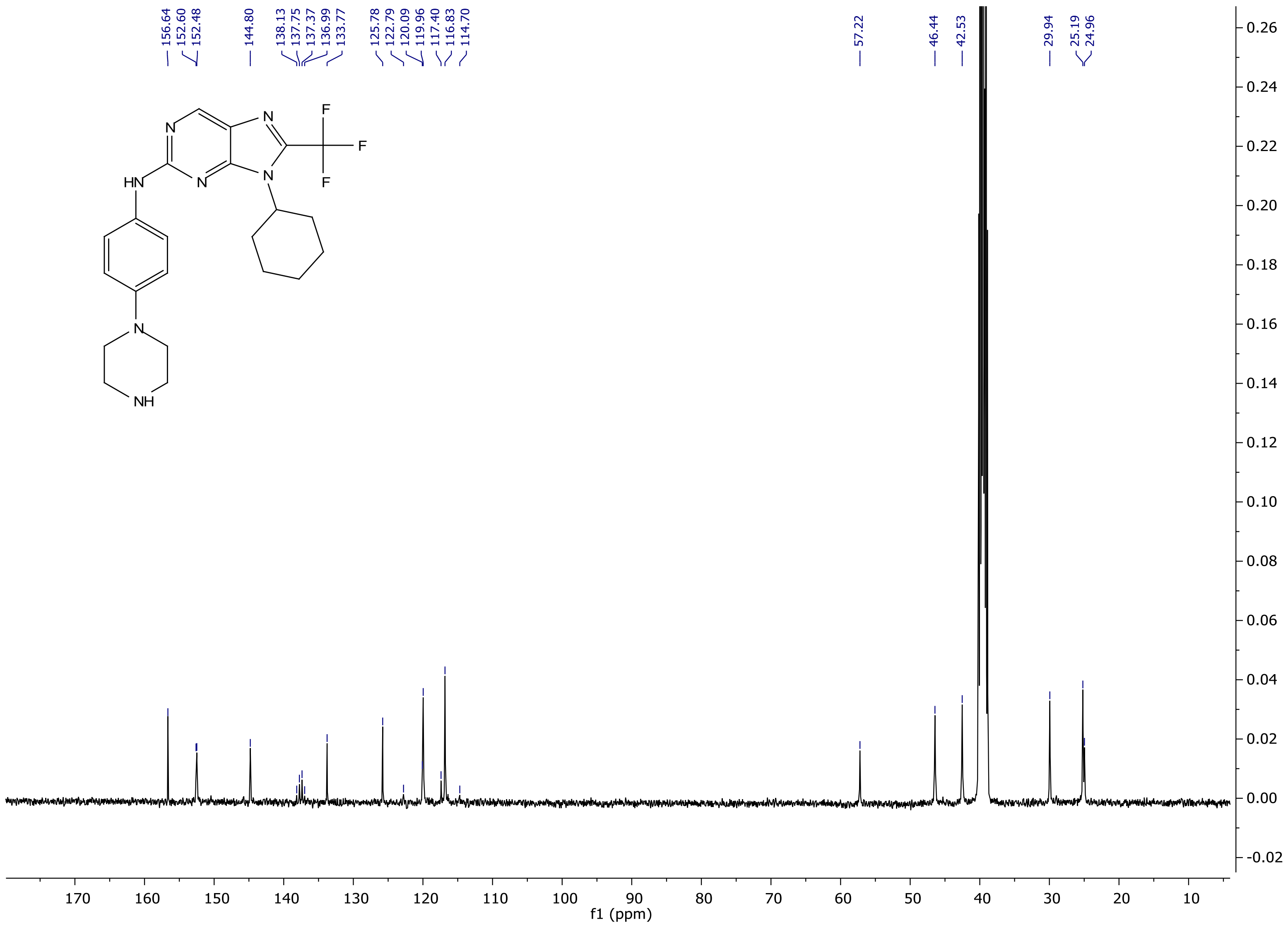

$^{1}$H NMR spectrum of **12c**

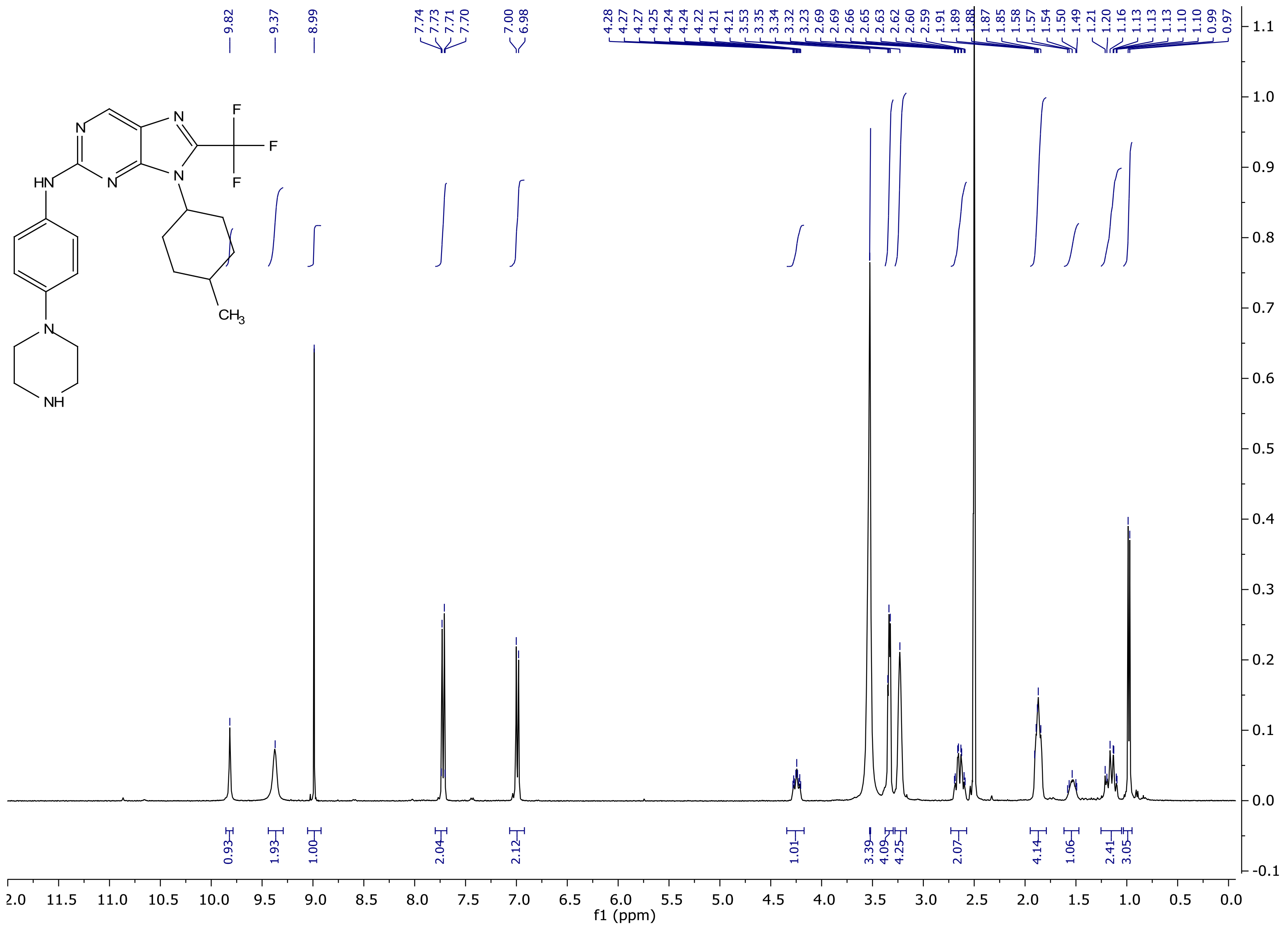

[13]C NMR spectrum of **12c**

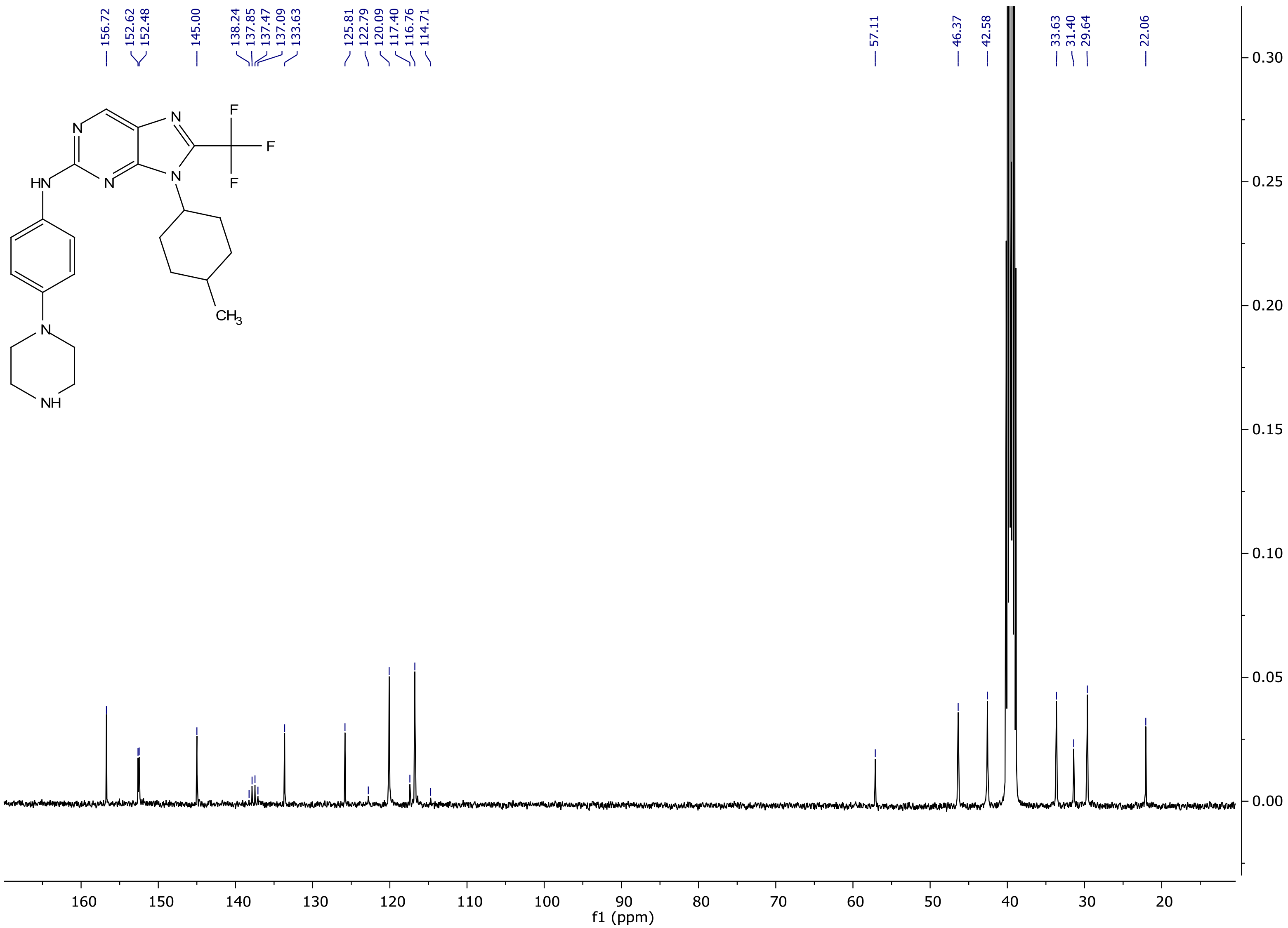

[1]H NMR spectrum of **12d**

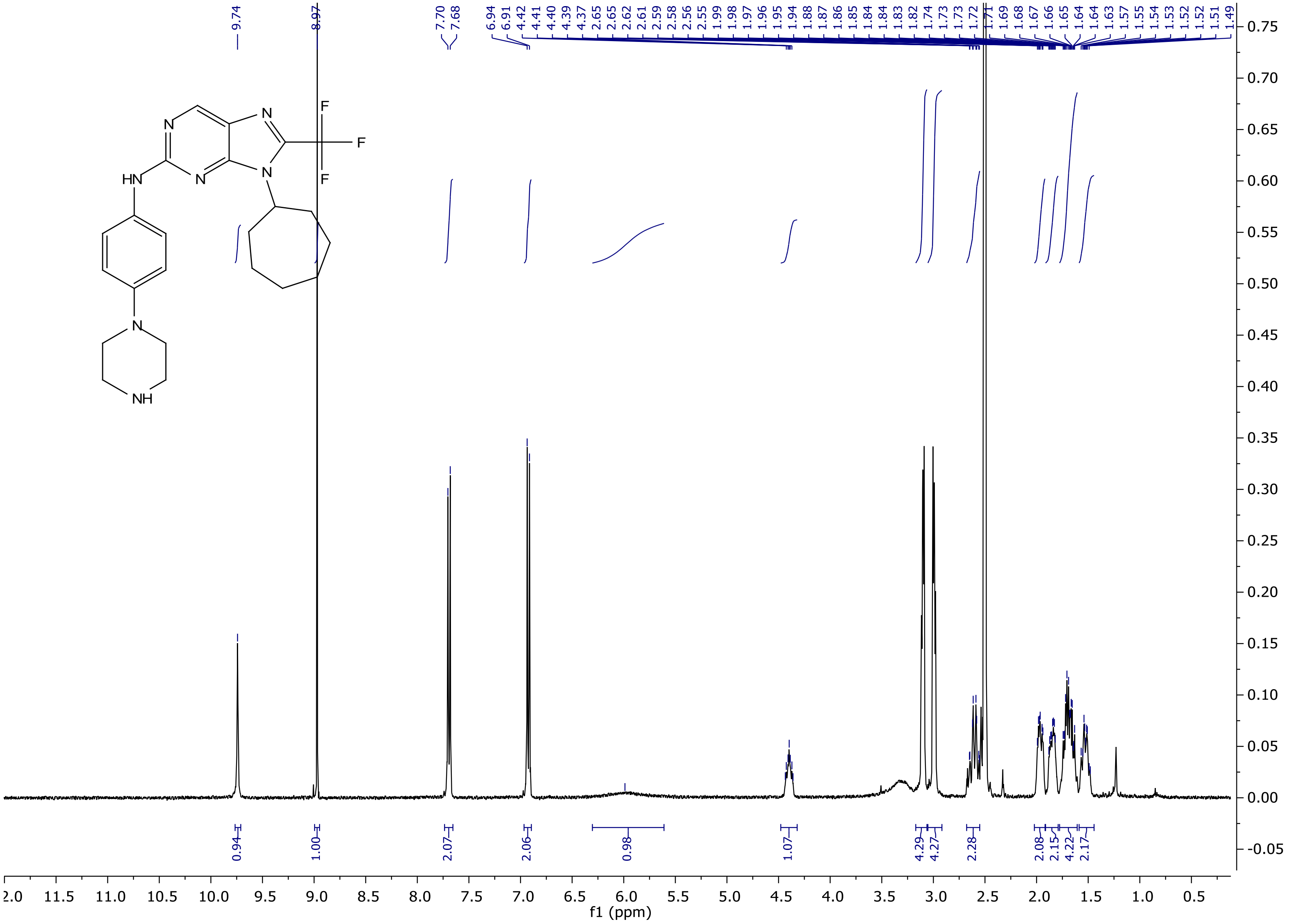

[13]C NMR spectrum of **12d**

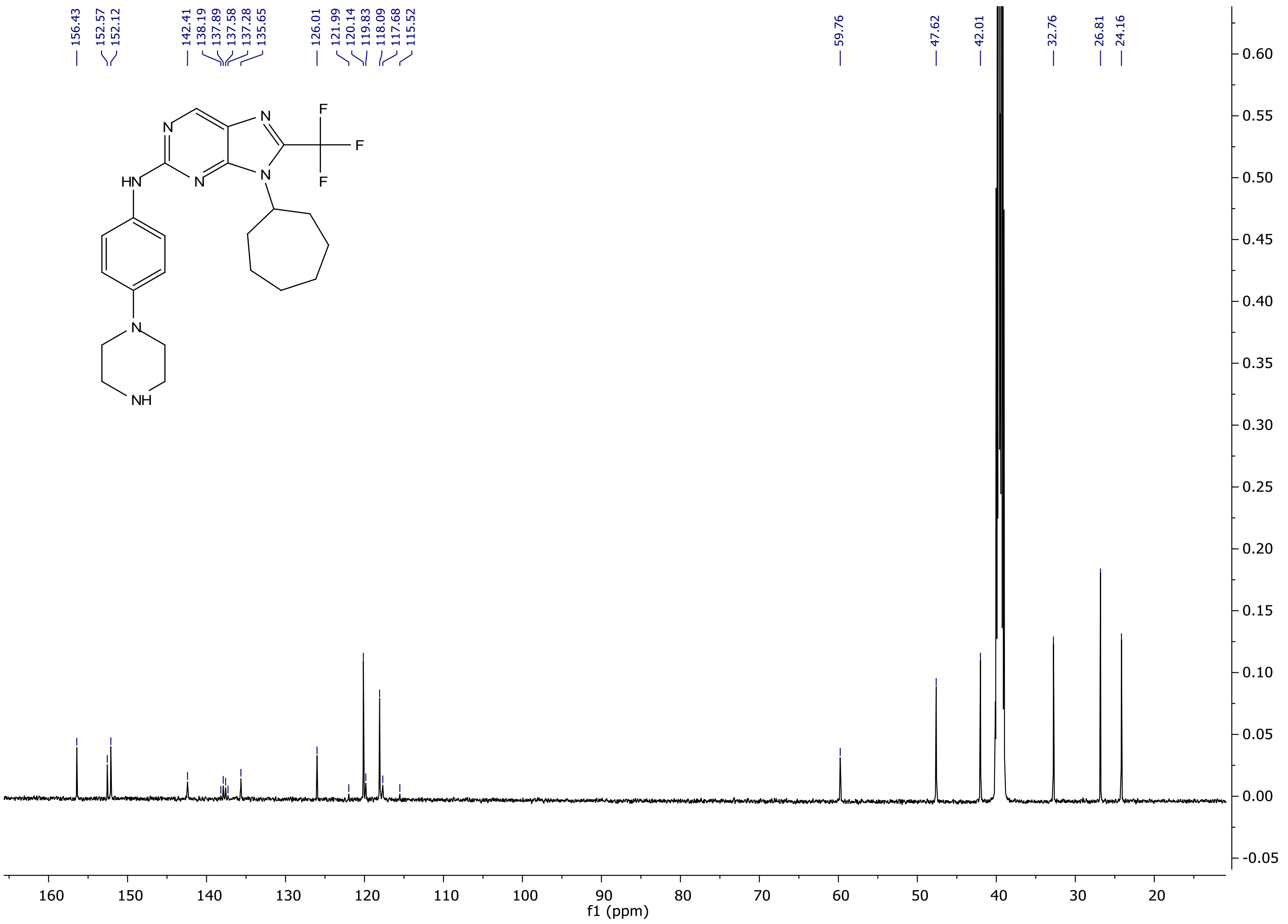

[1]H NMR spectrum of **13a**

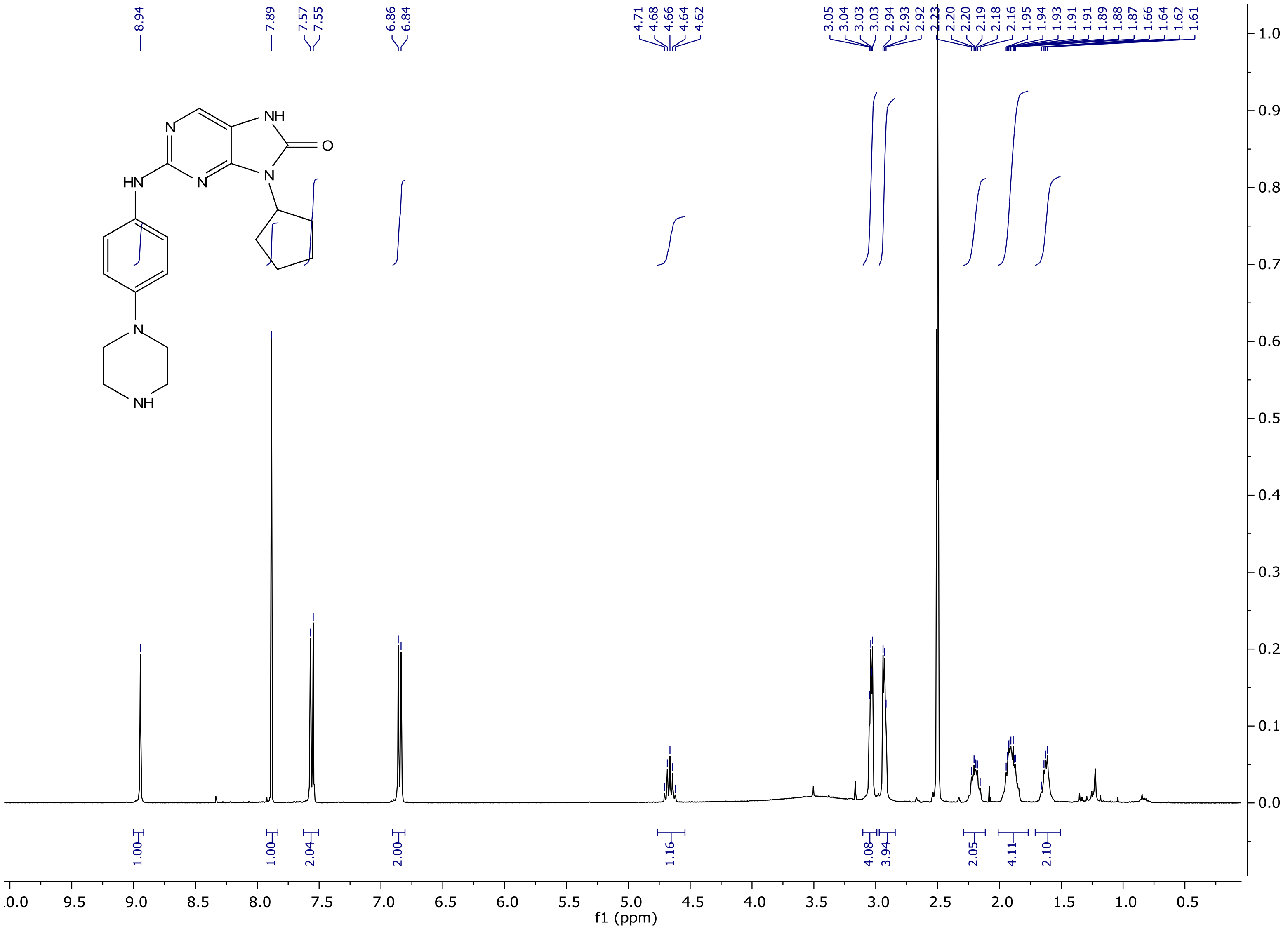

$^{13}$C NMR spectrum of **13a**

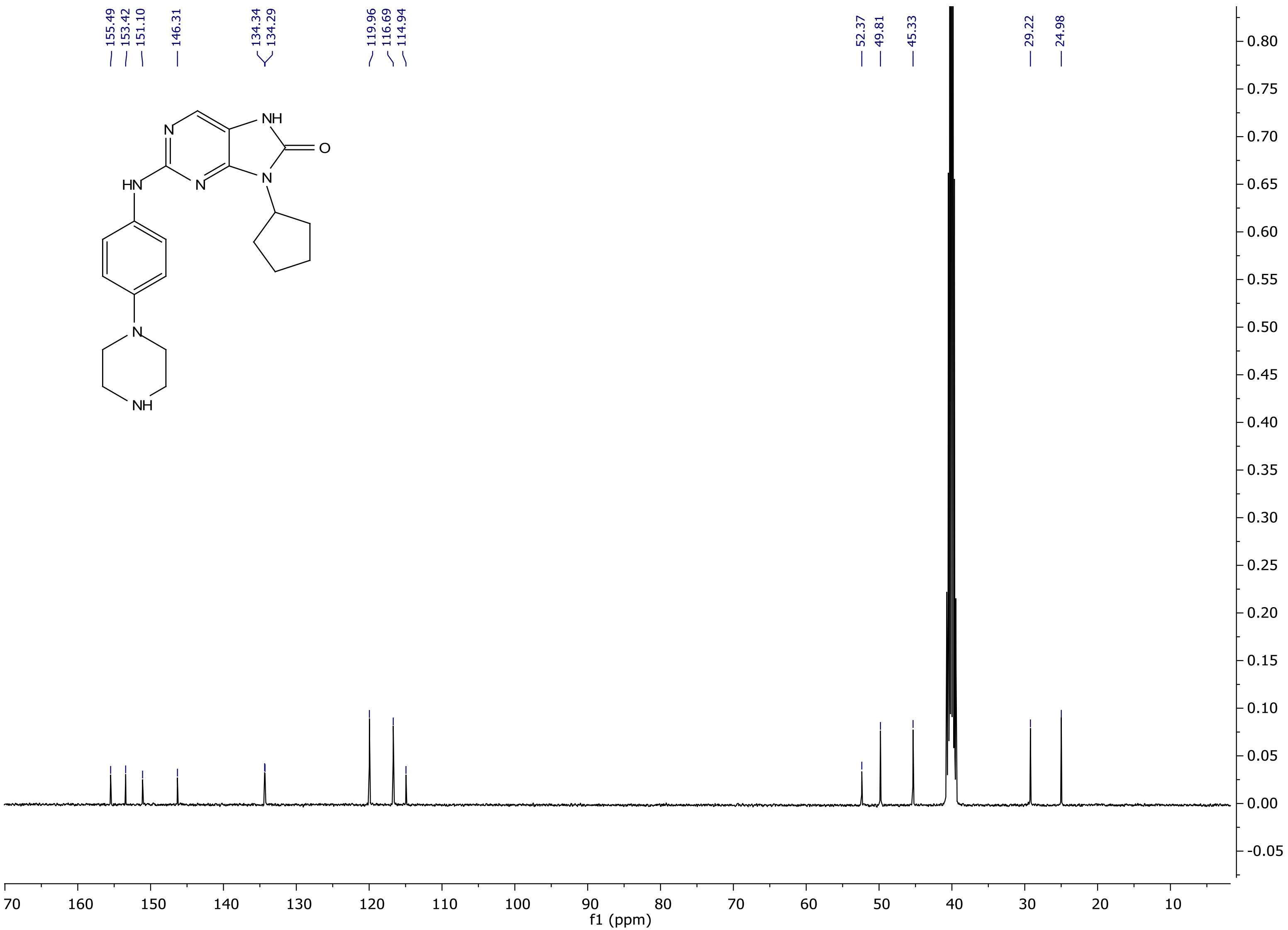

$^{1}$H NMR spectrum of **13b**

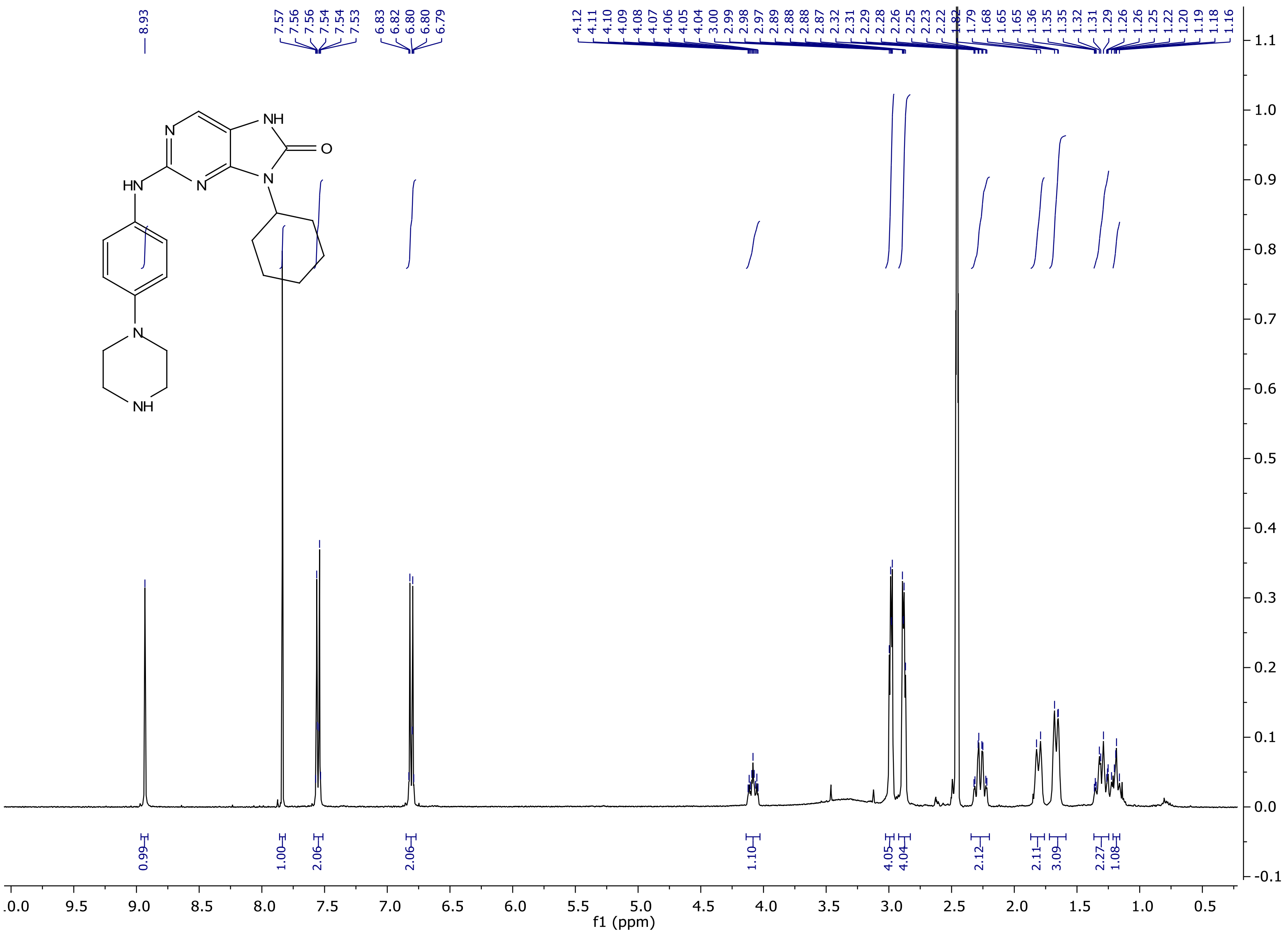

$^{13}$C NMR spectrum of **13b**

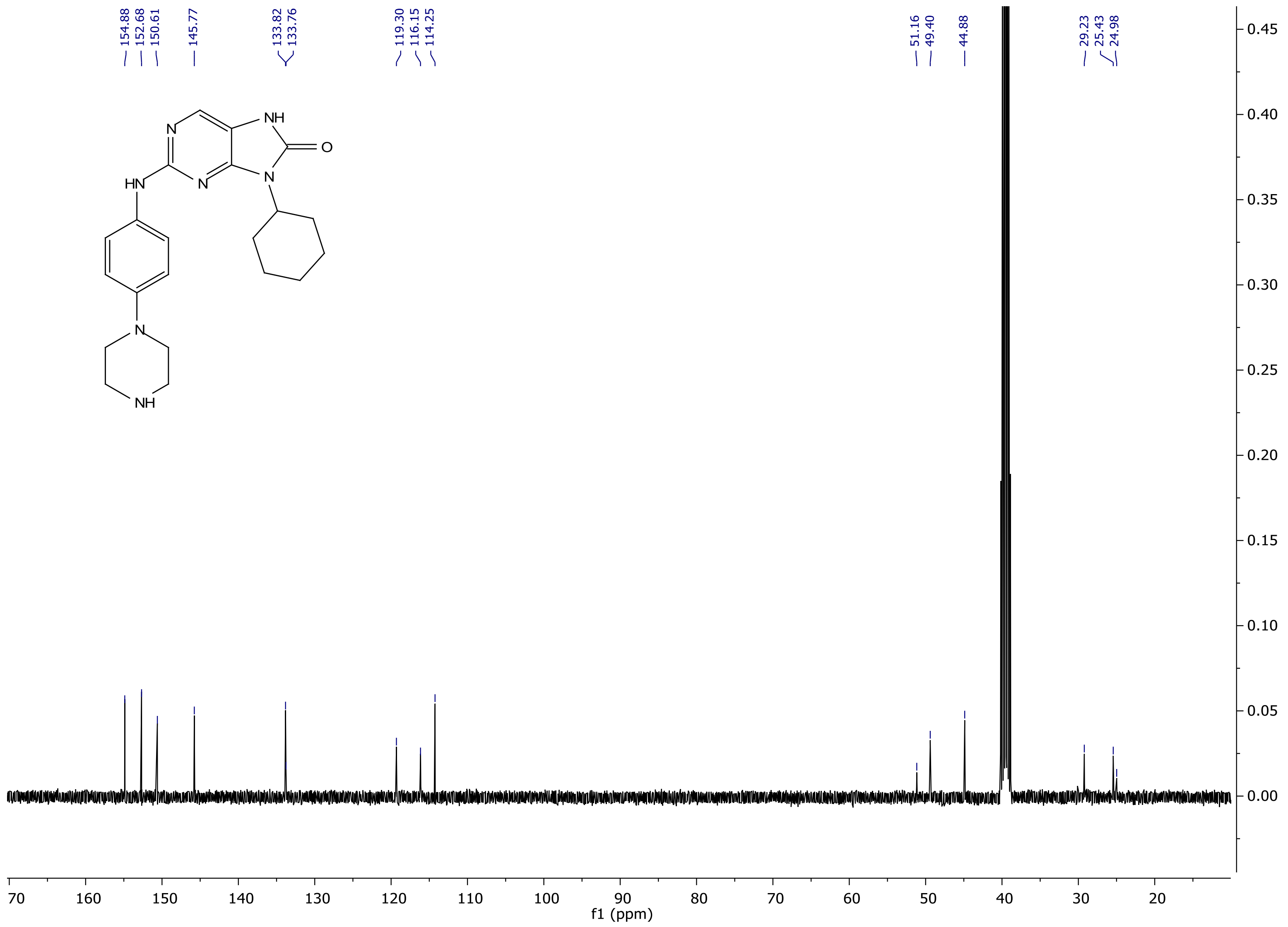

[1]H NMR spectrum of **13c**

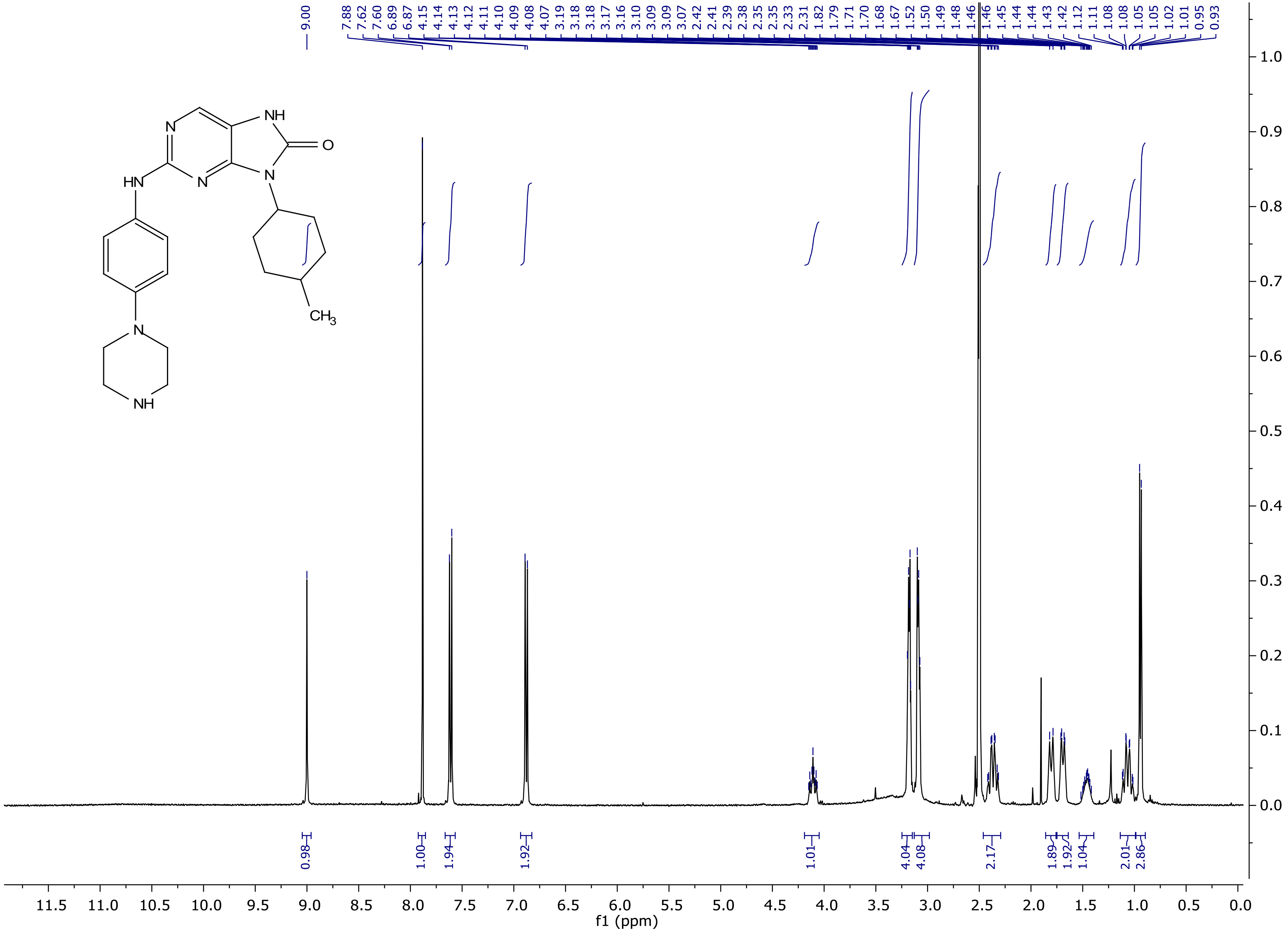

[13]C NMR spectrum of **13c**

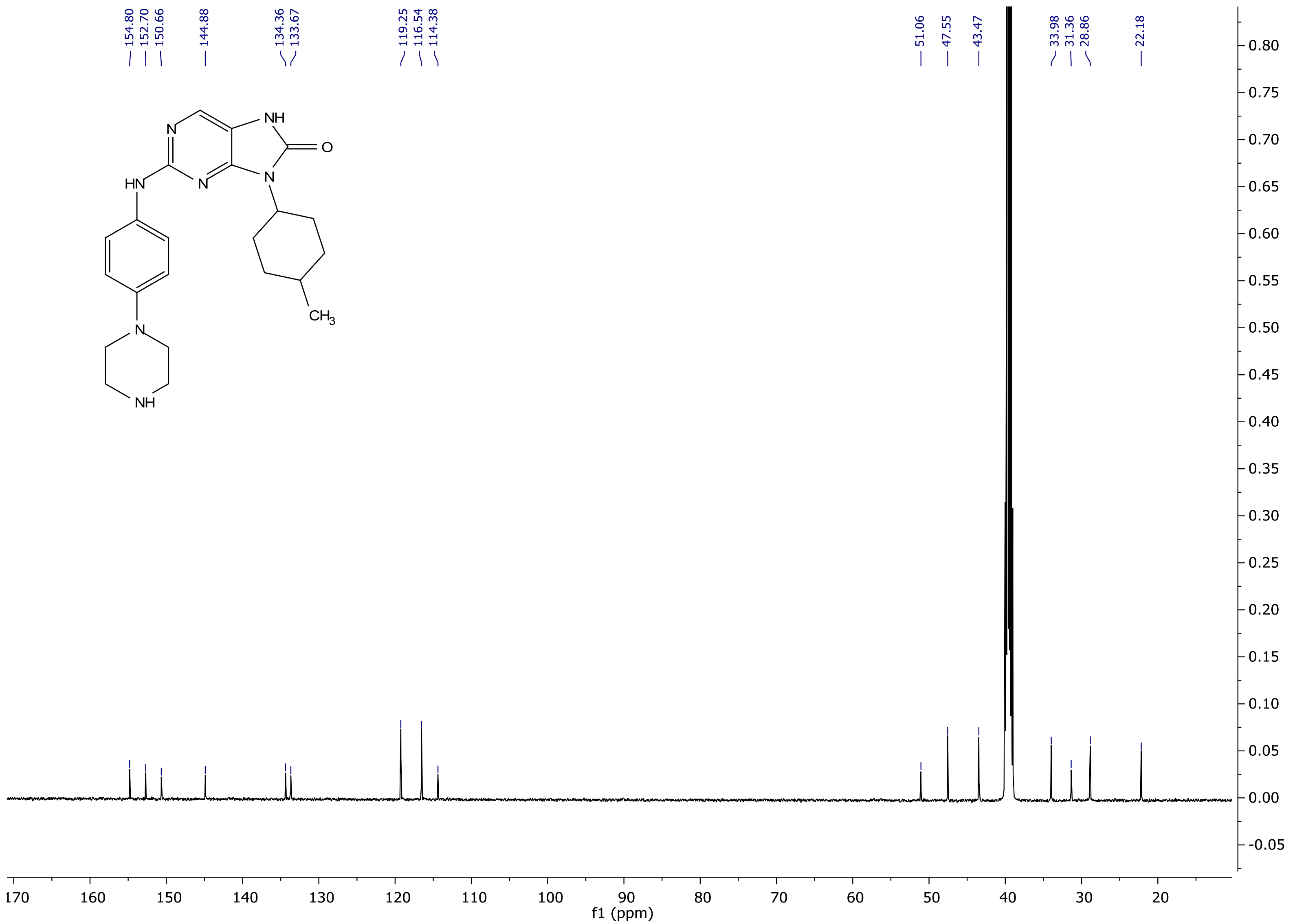

[1]H NMR spectrum of **13d**

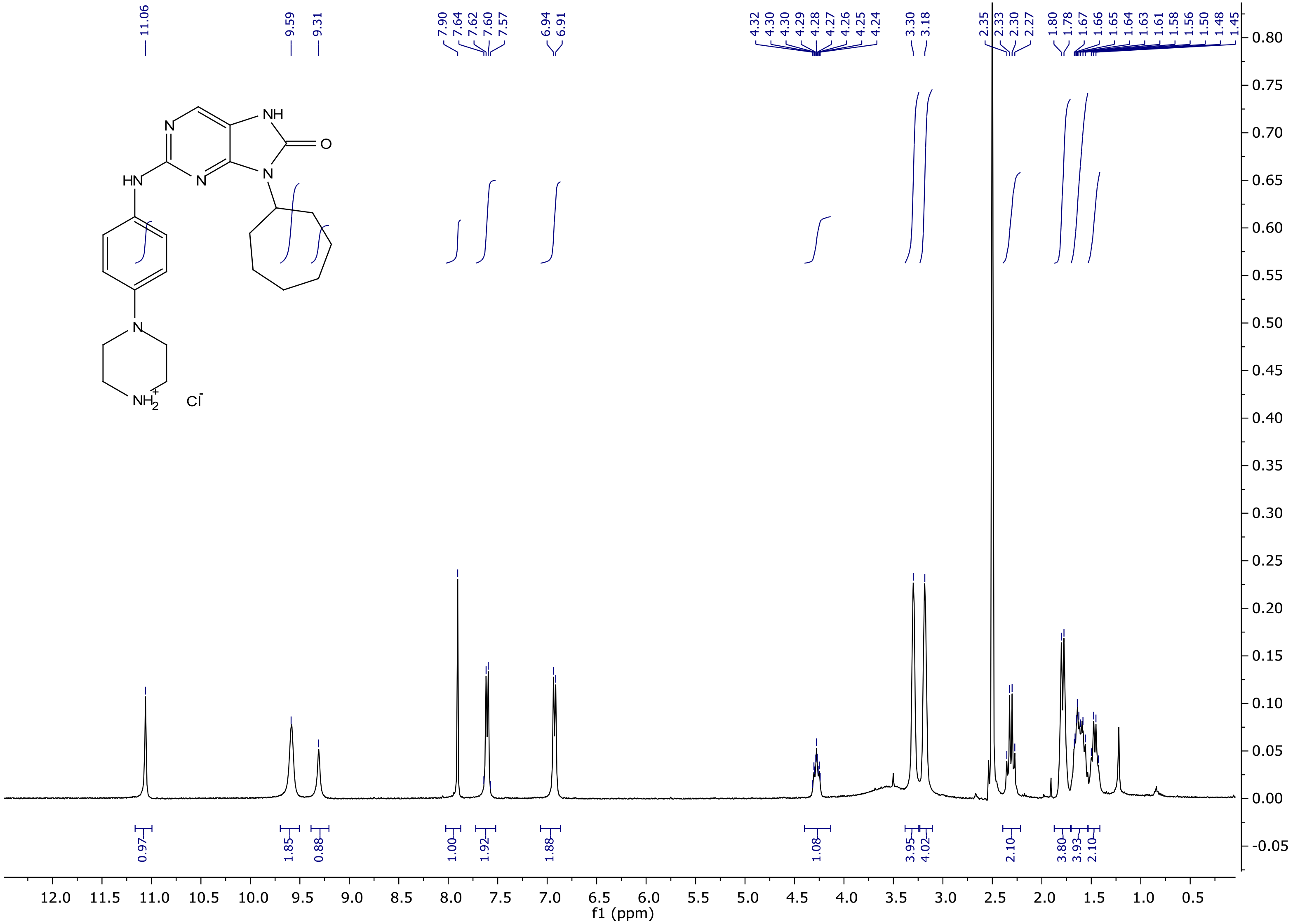

$^{13}$C NMR spectrum of **13d**

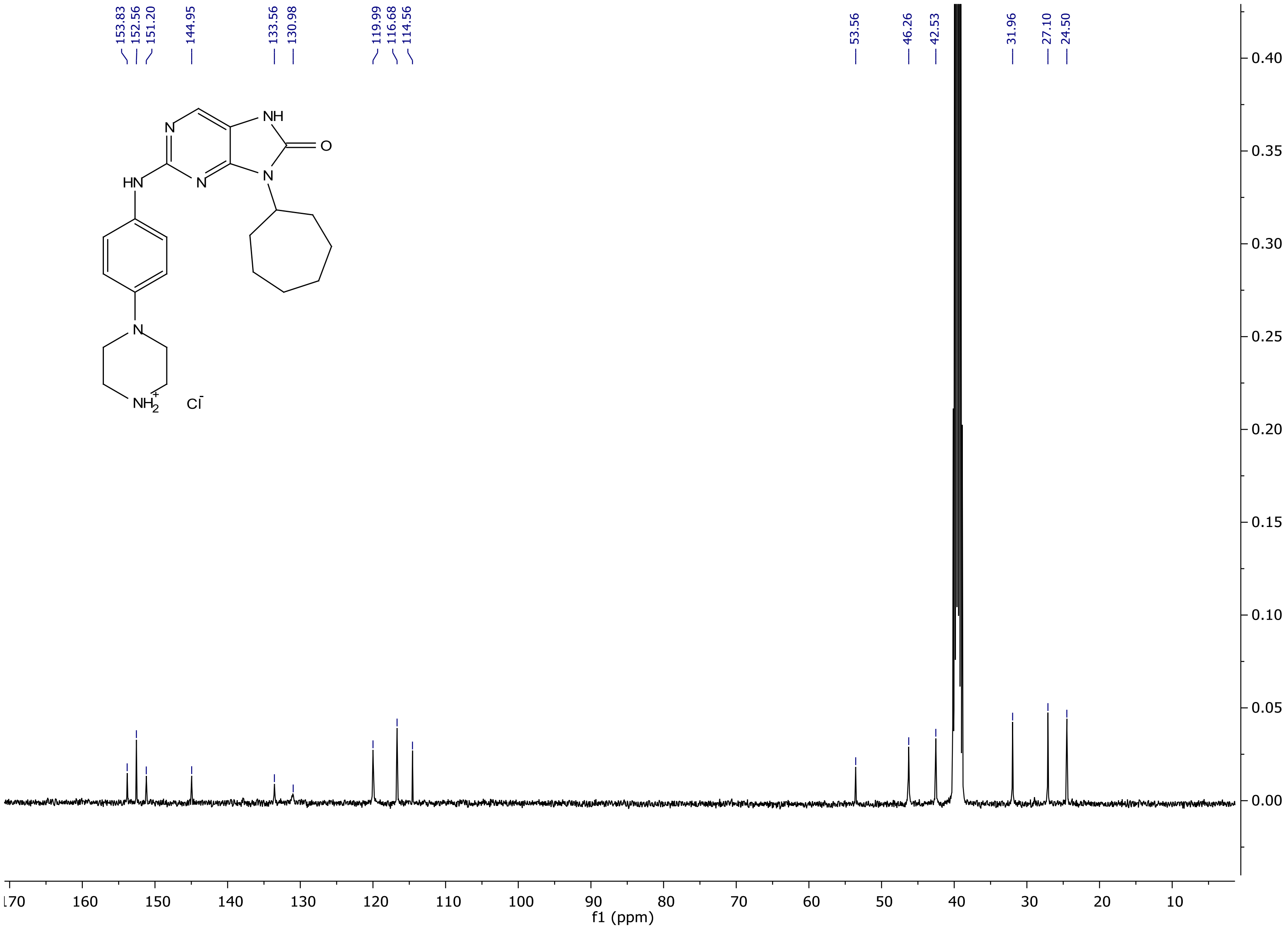

[1]H NMR spectrum of **14a**

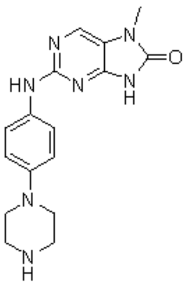

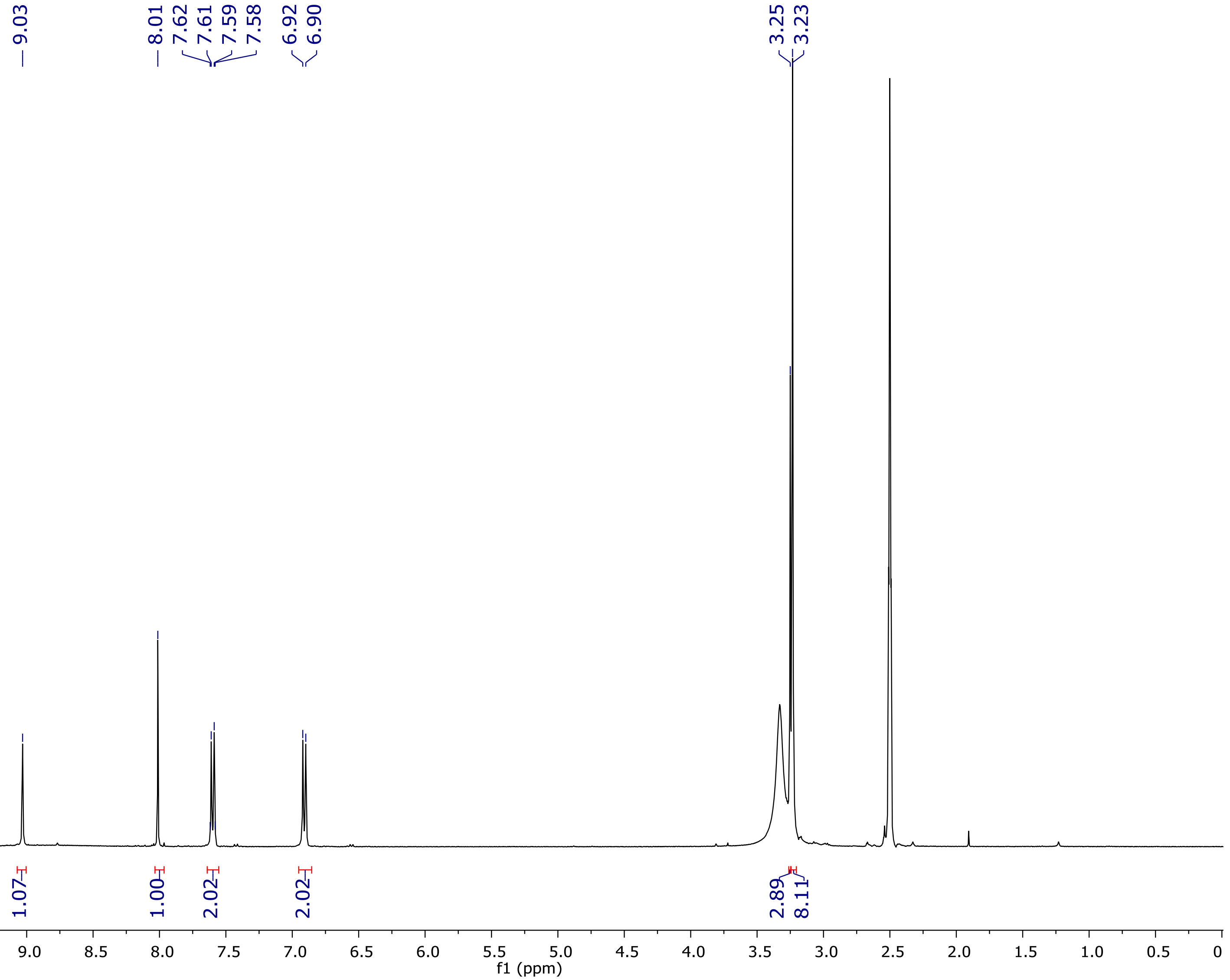

$^{13}$C NMR spectrum of **14a**

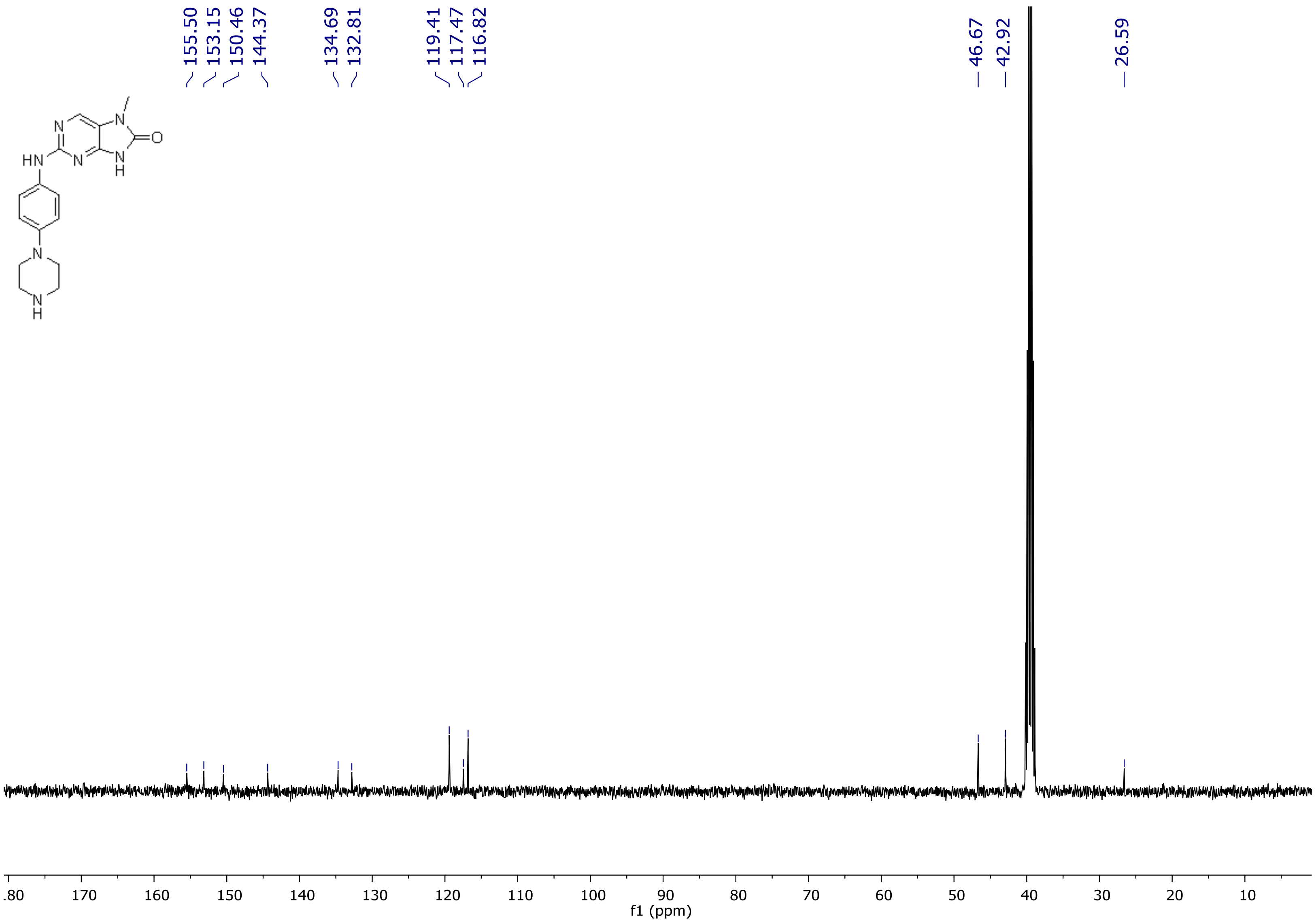

[1]H NMR spectrum of **14b**

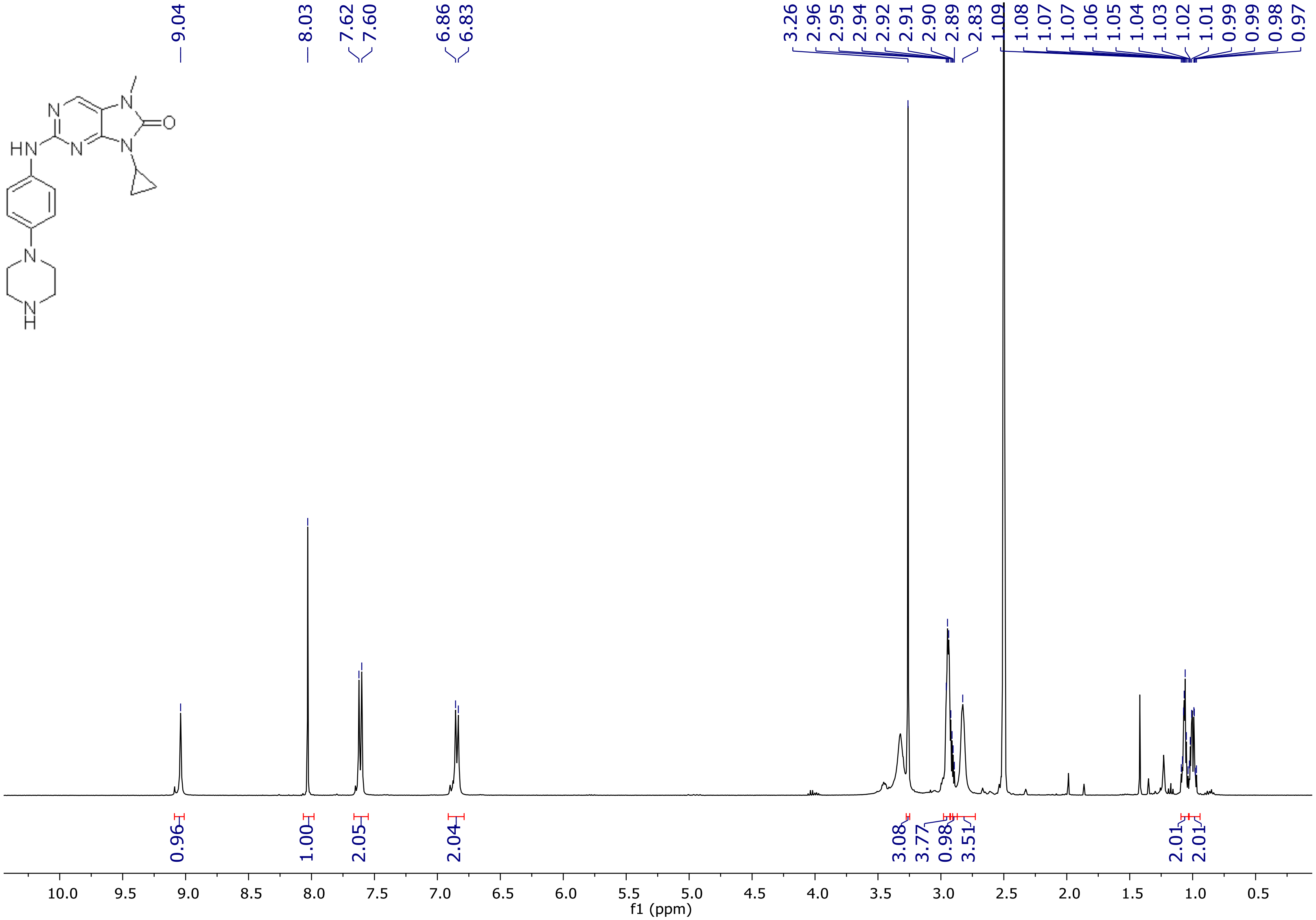

$^{13}$C NMR spectrum of **14b**

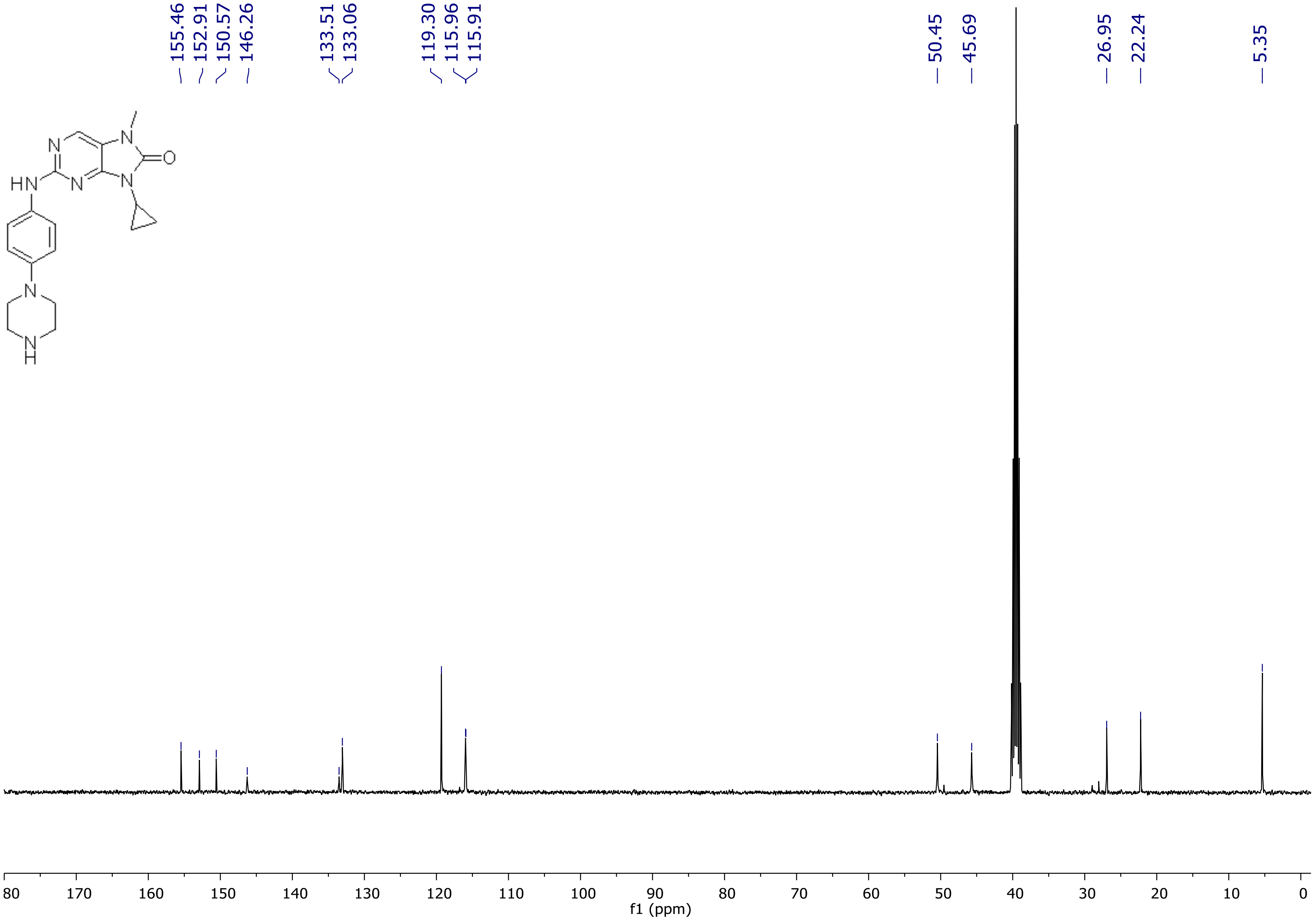

[1]H NMR spectrum of **14c**

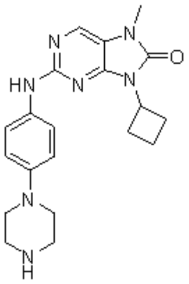


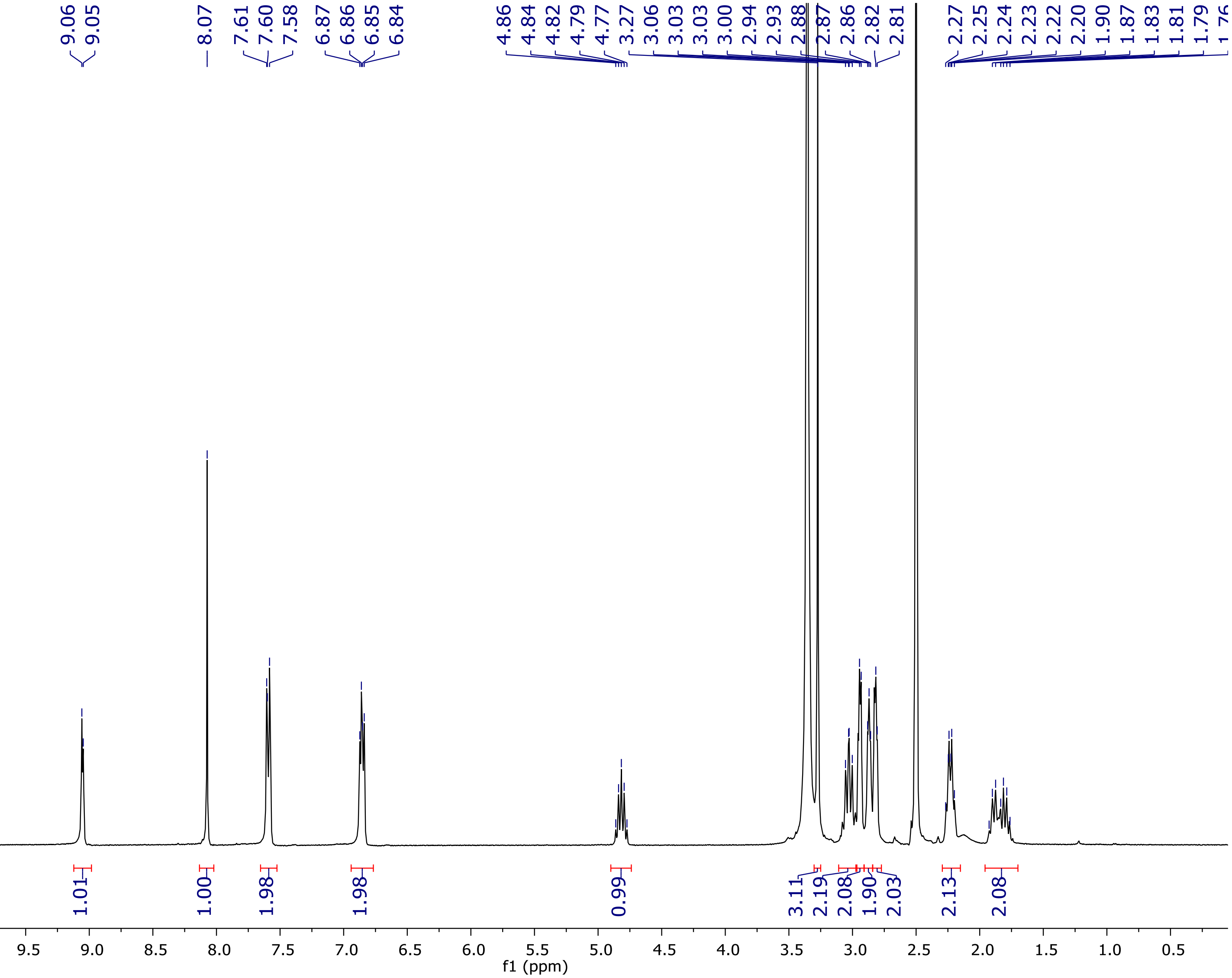

$^{13}$C NMR spectrum of **14c**

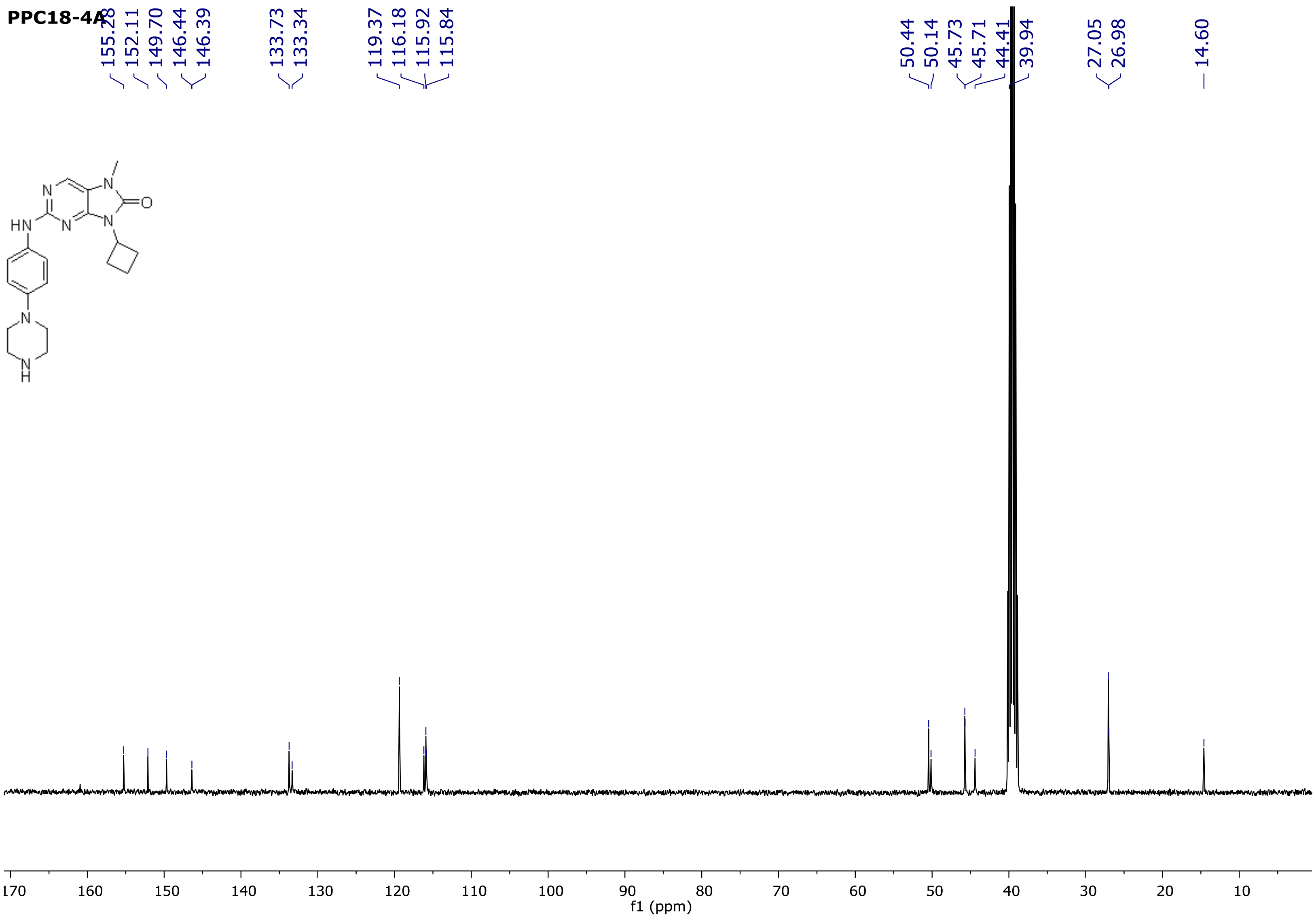

[1]H NMR spectrum of **14d**

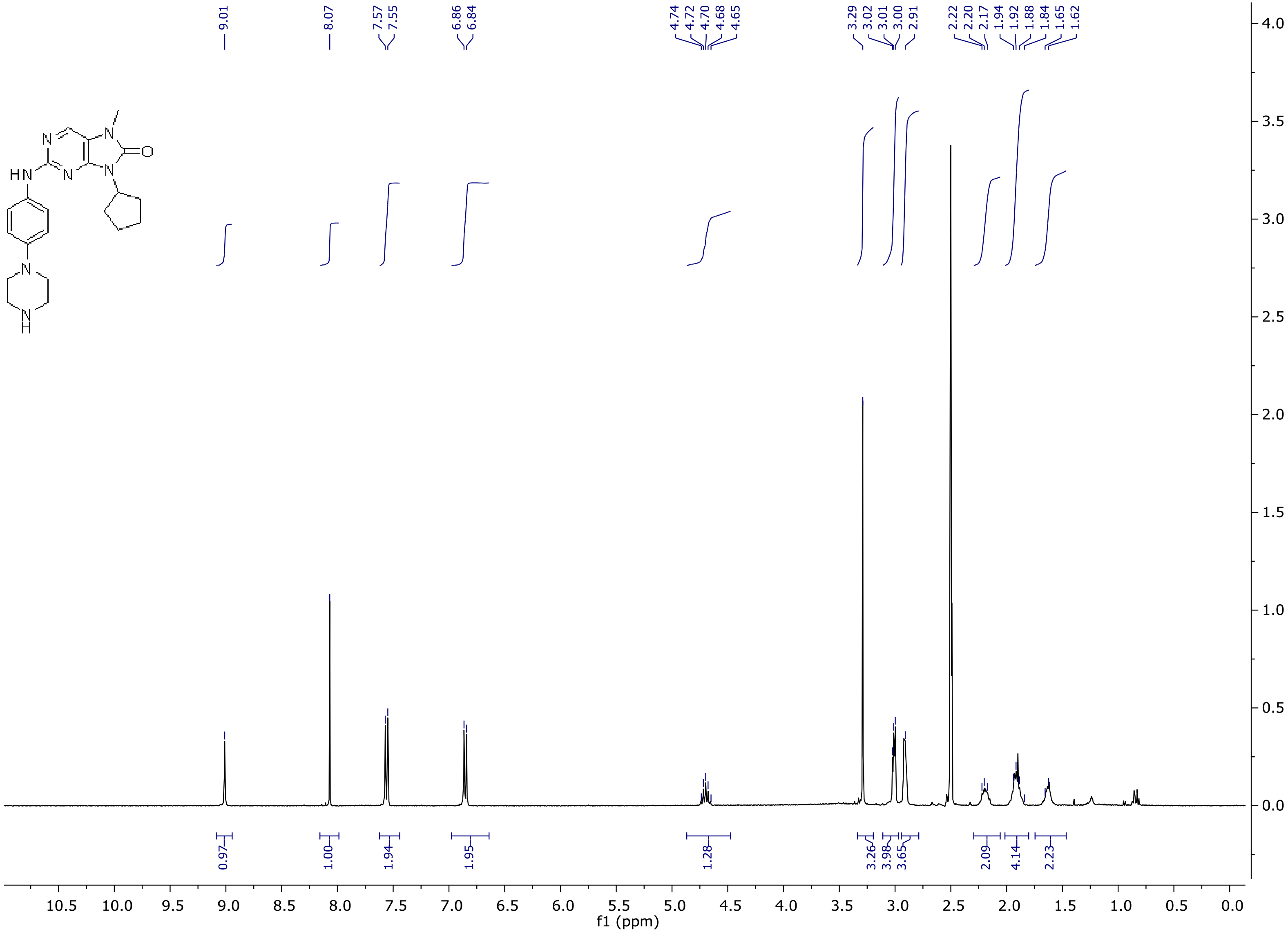

$^{1}$H NMR spectrum of **14d**

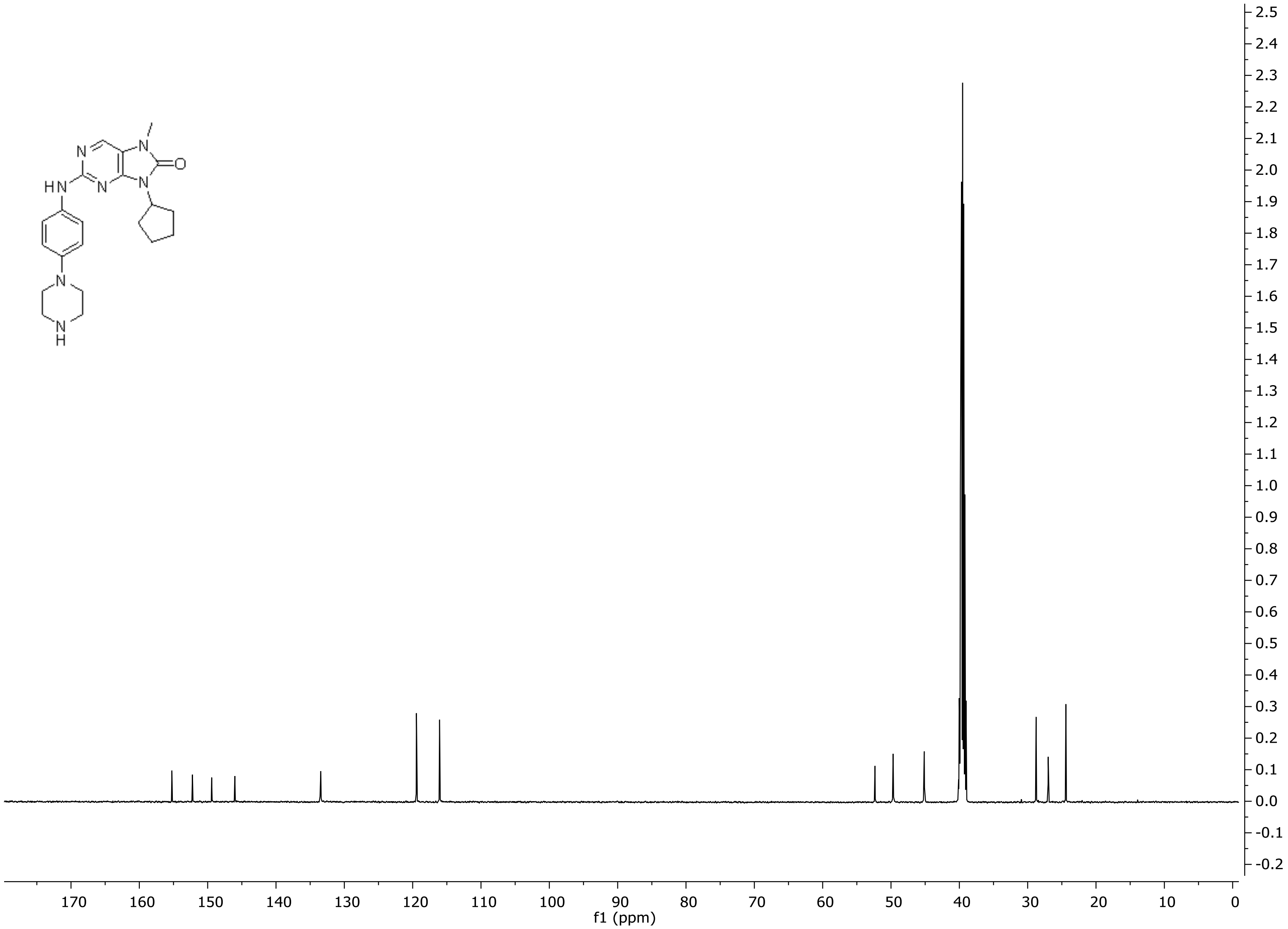

[1]H NMR spectrum of **14e**

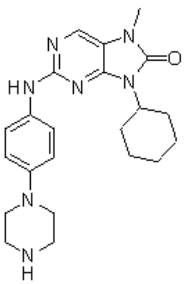


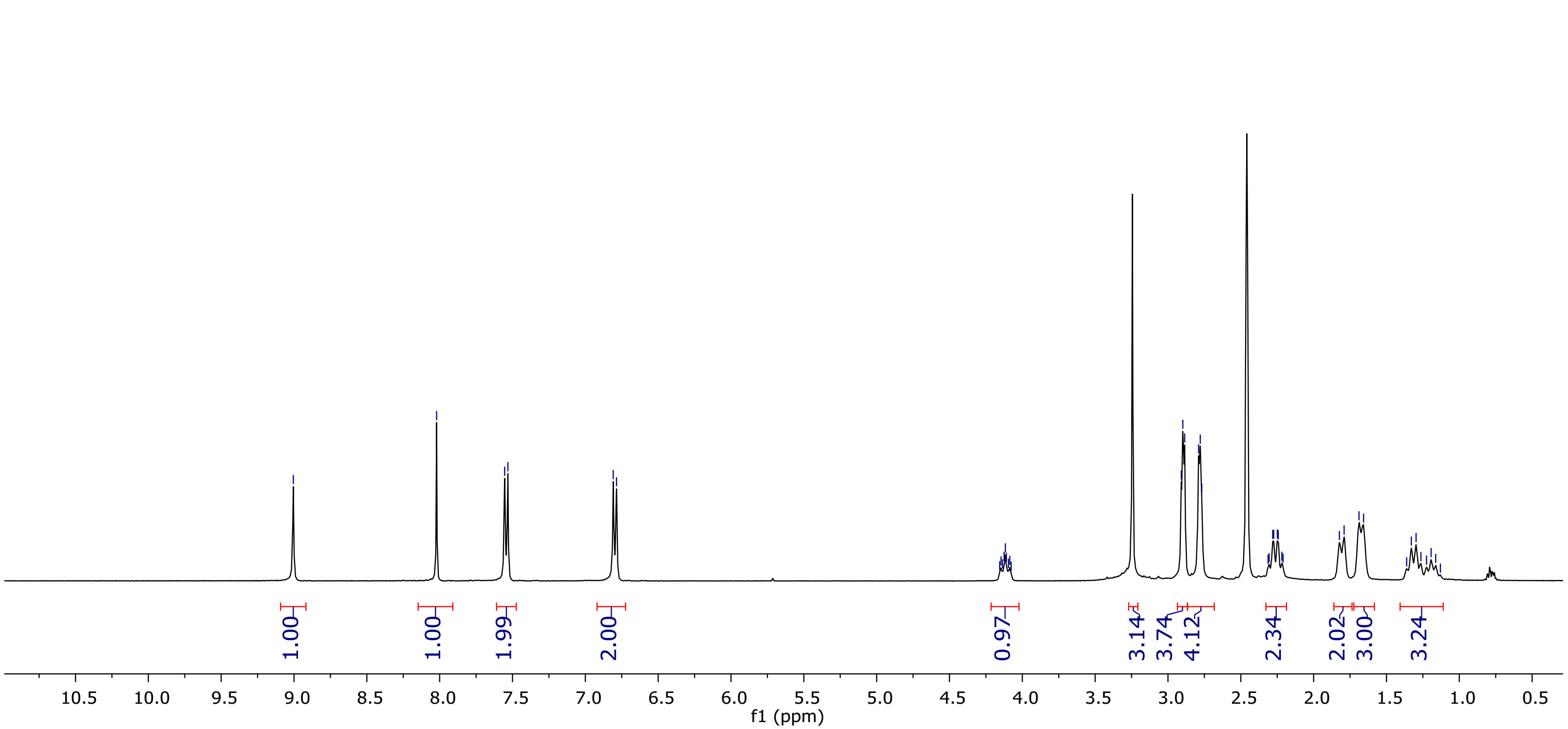

[1]H NMR spectrum of **14e**

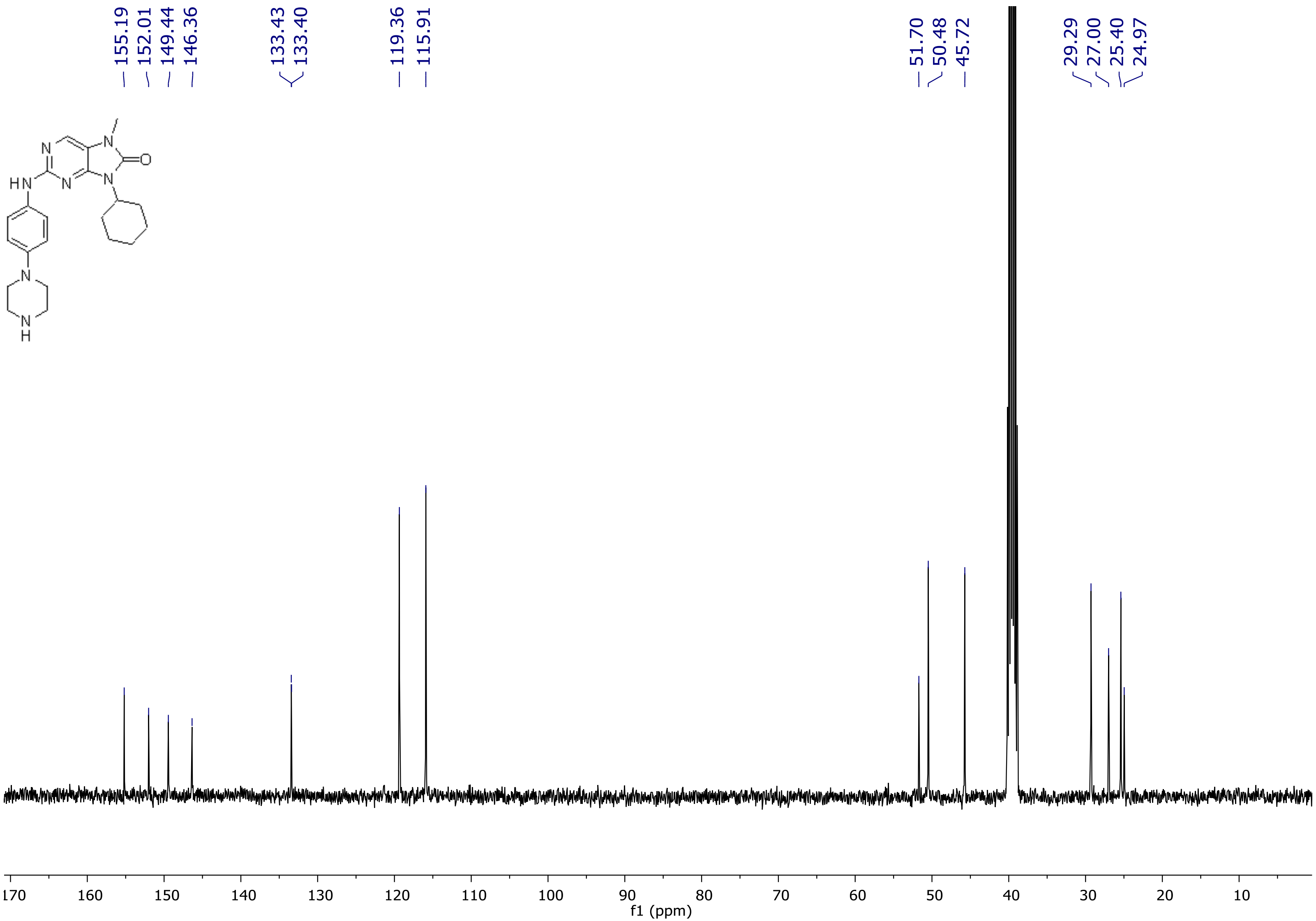

[1]H NMR spectrum of **14f**

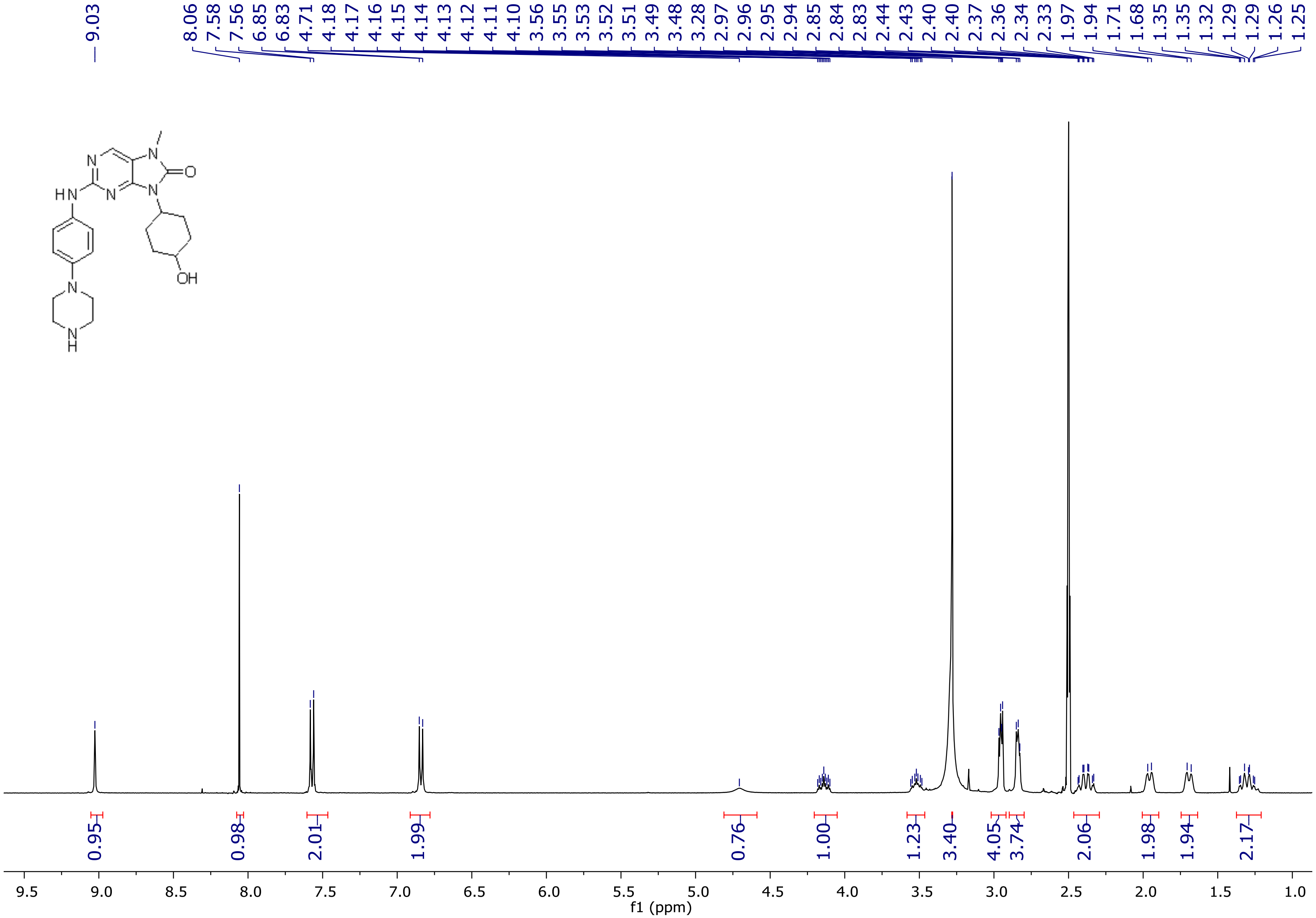

[1]H NMR spectrum of **14f**

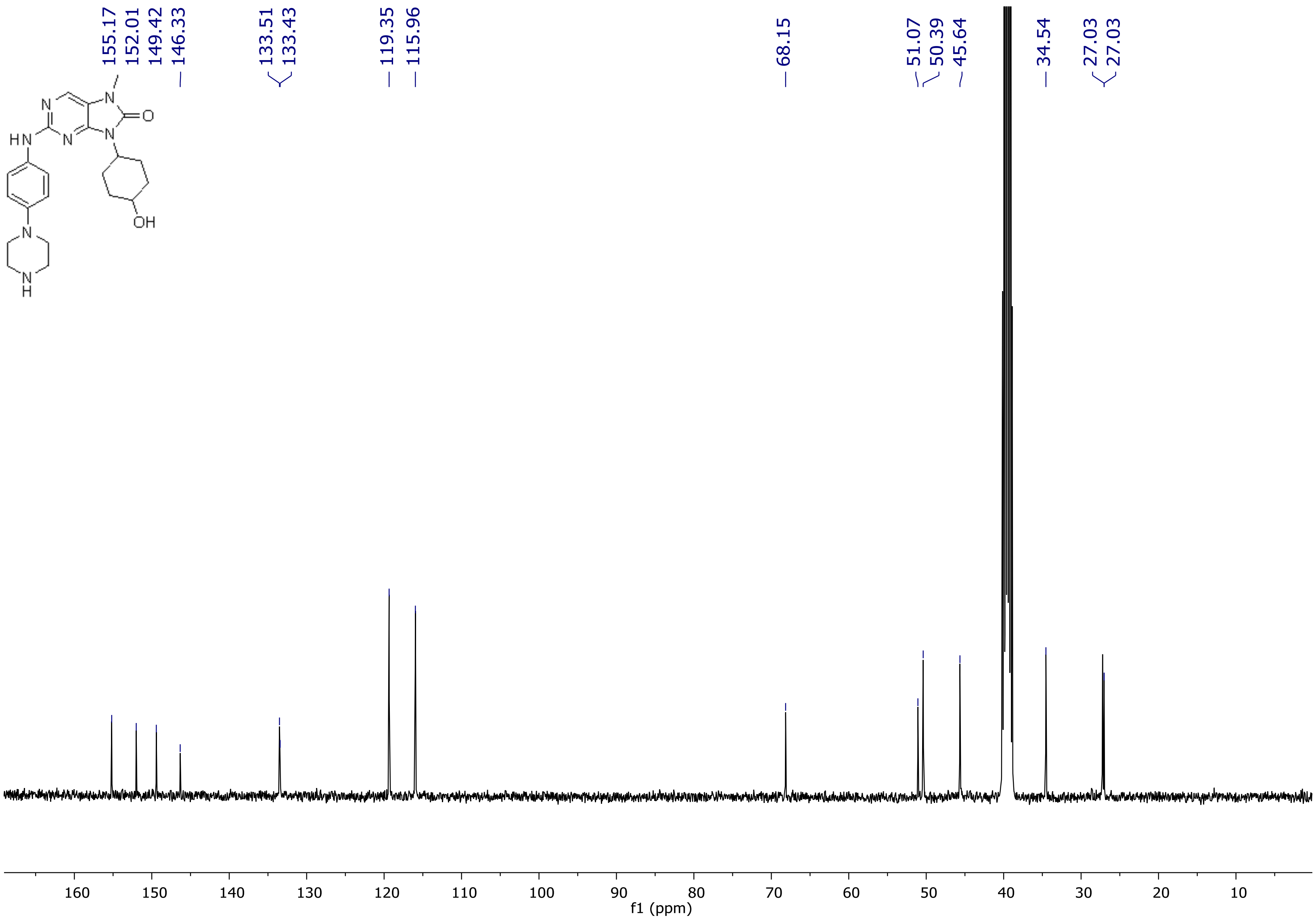

$^{1}$H NMR spectrum of **14g**

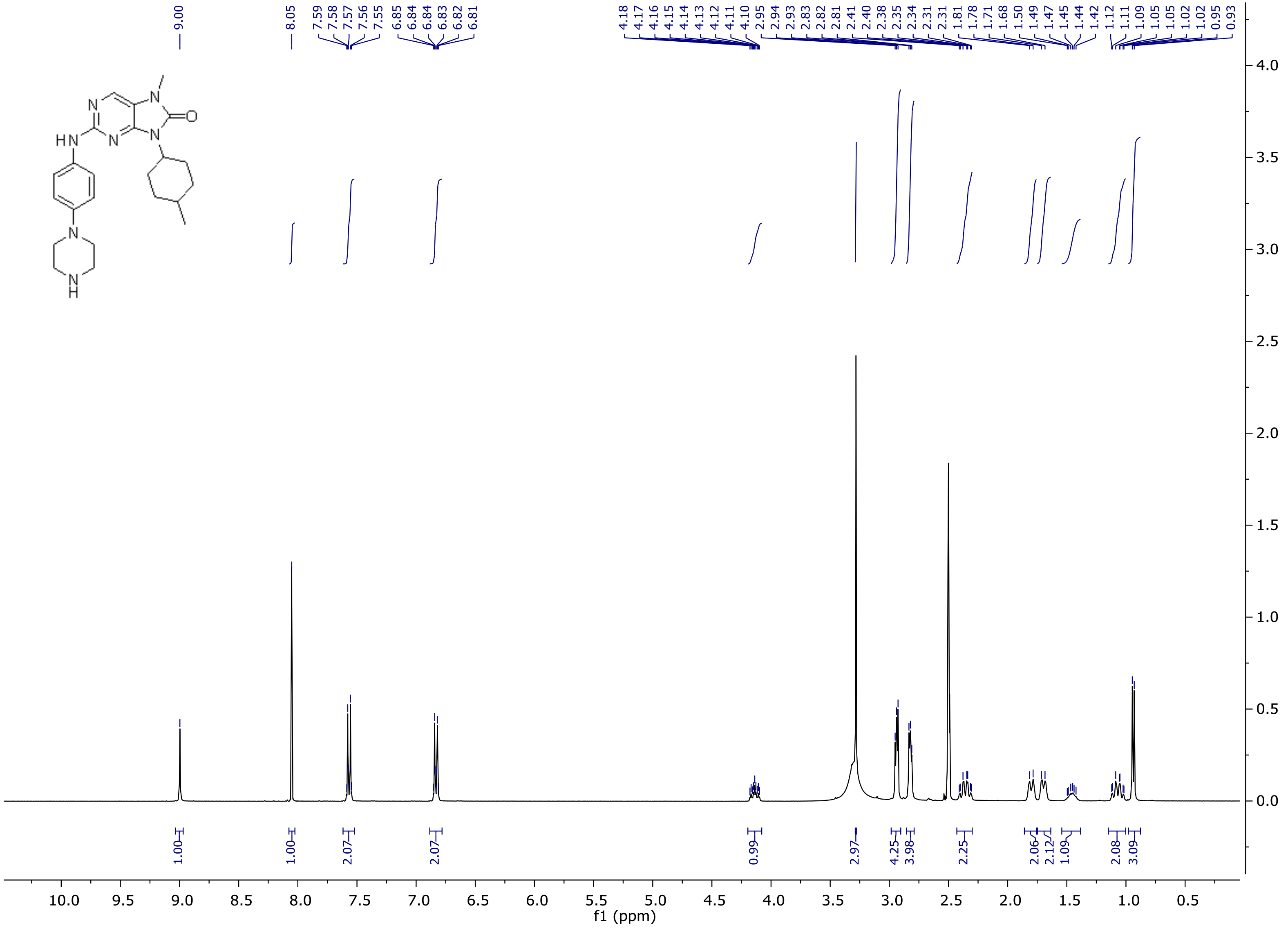

[1]H NMR spectrum of **14g**

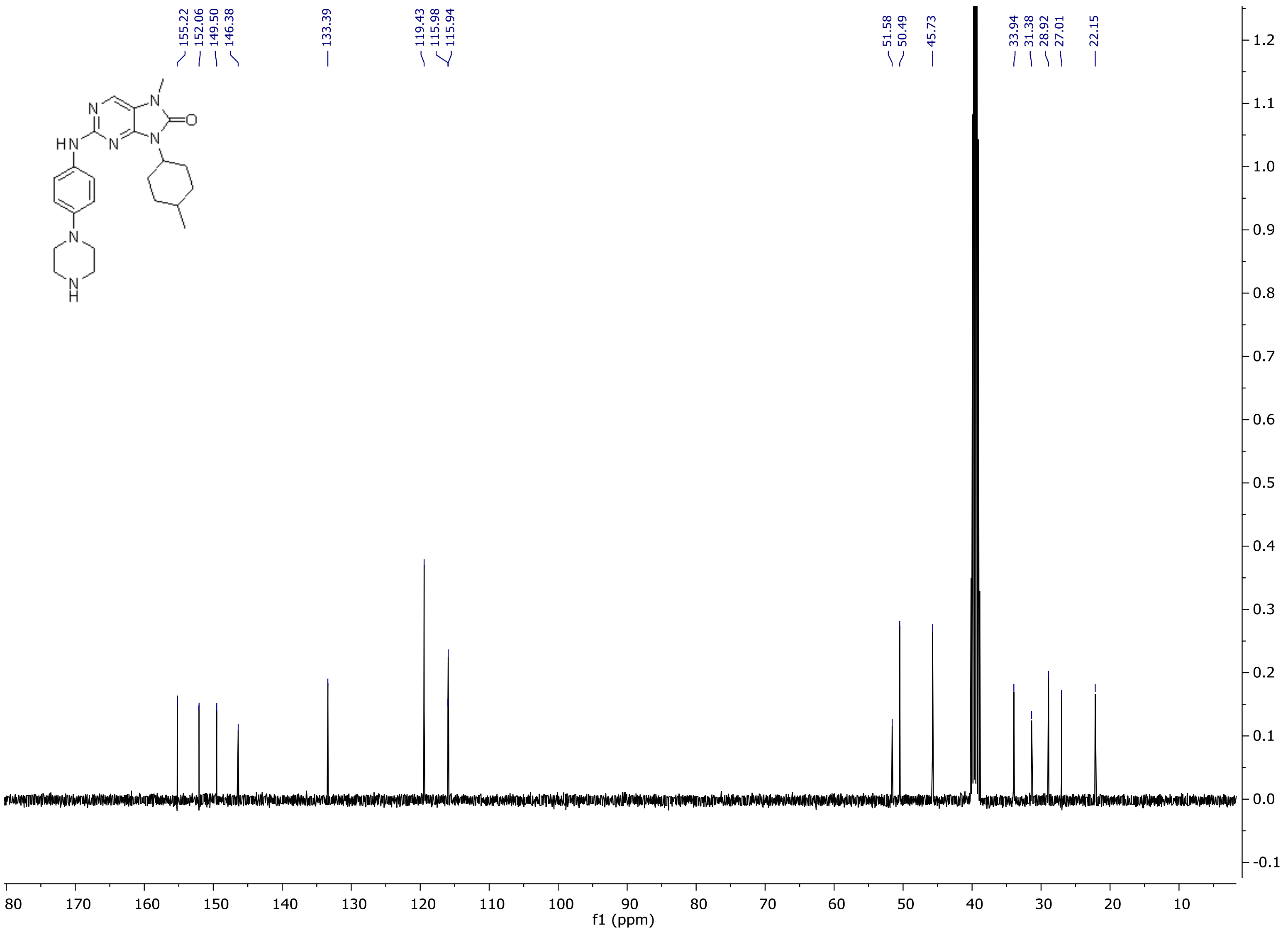

[1]H NMR spectrum of **14h**

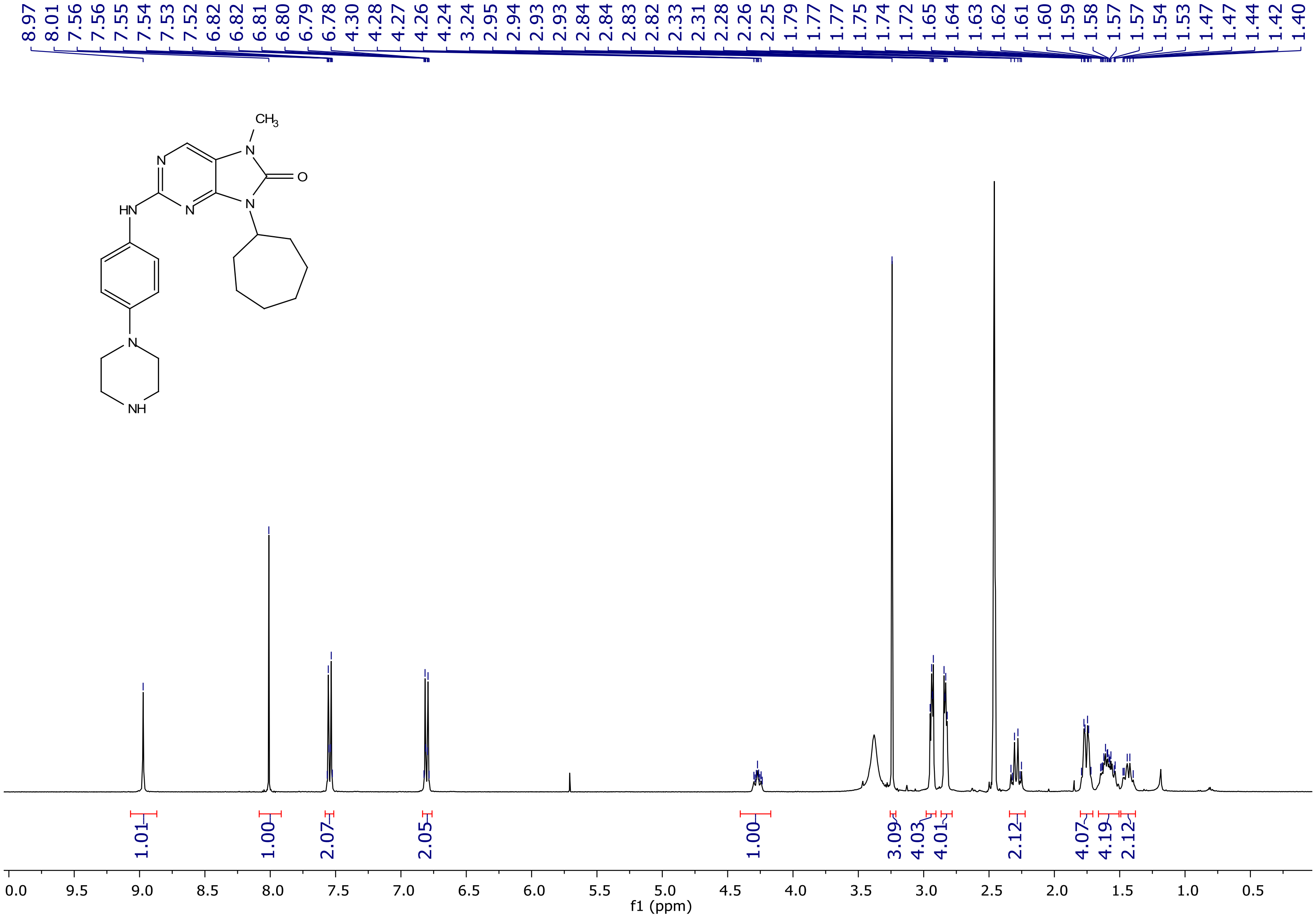

[1]H NMR spectrum of **14h**

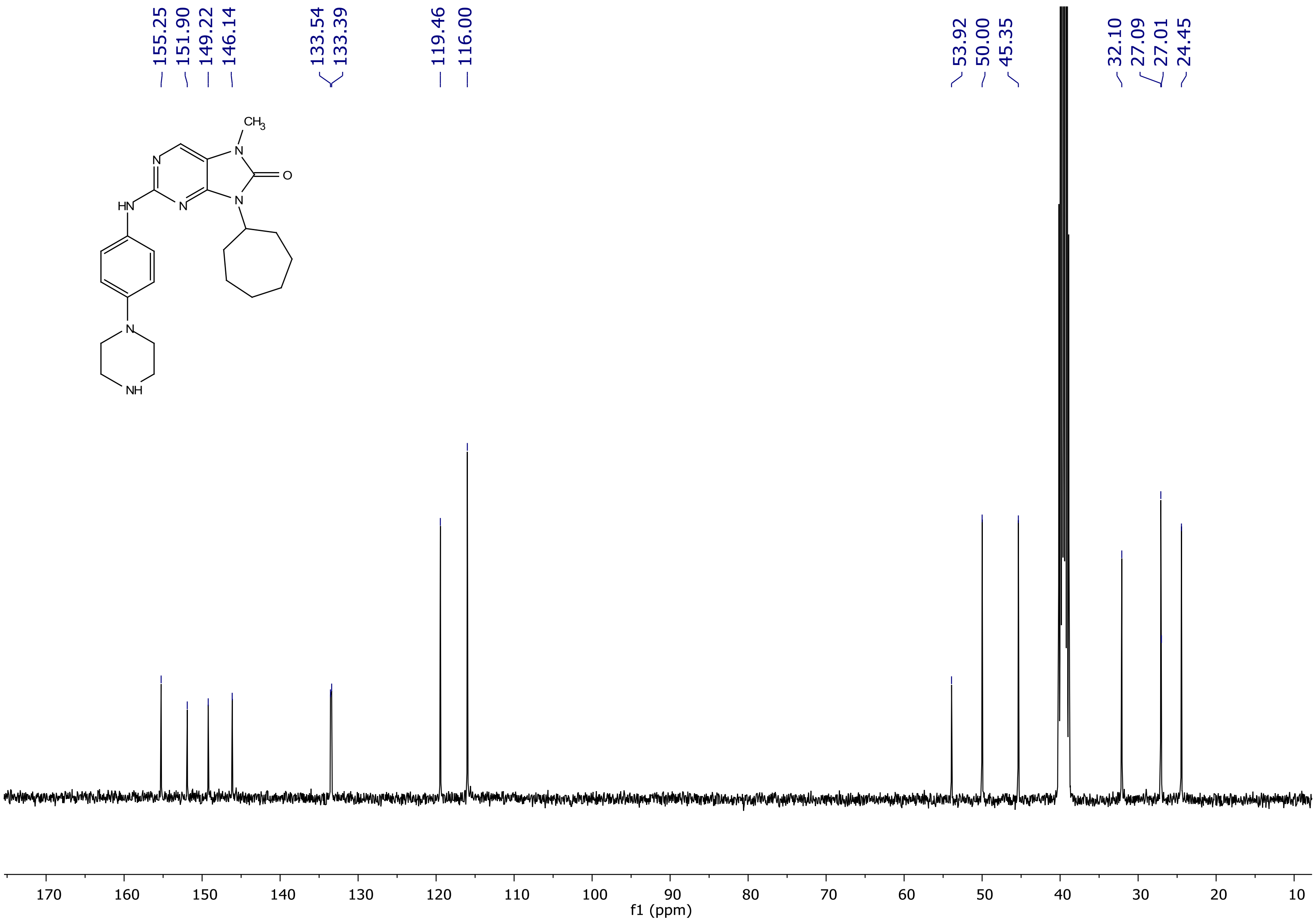

[1]H NMR spectrum of **14i**

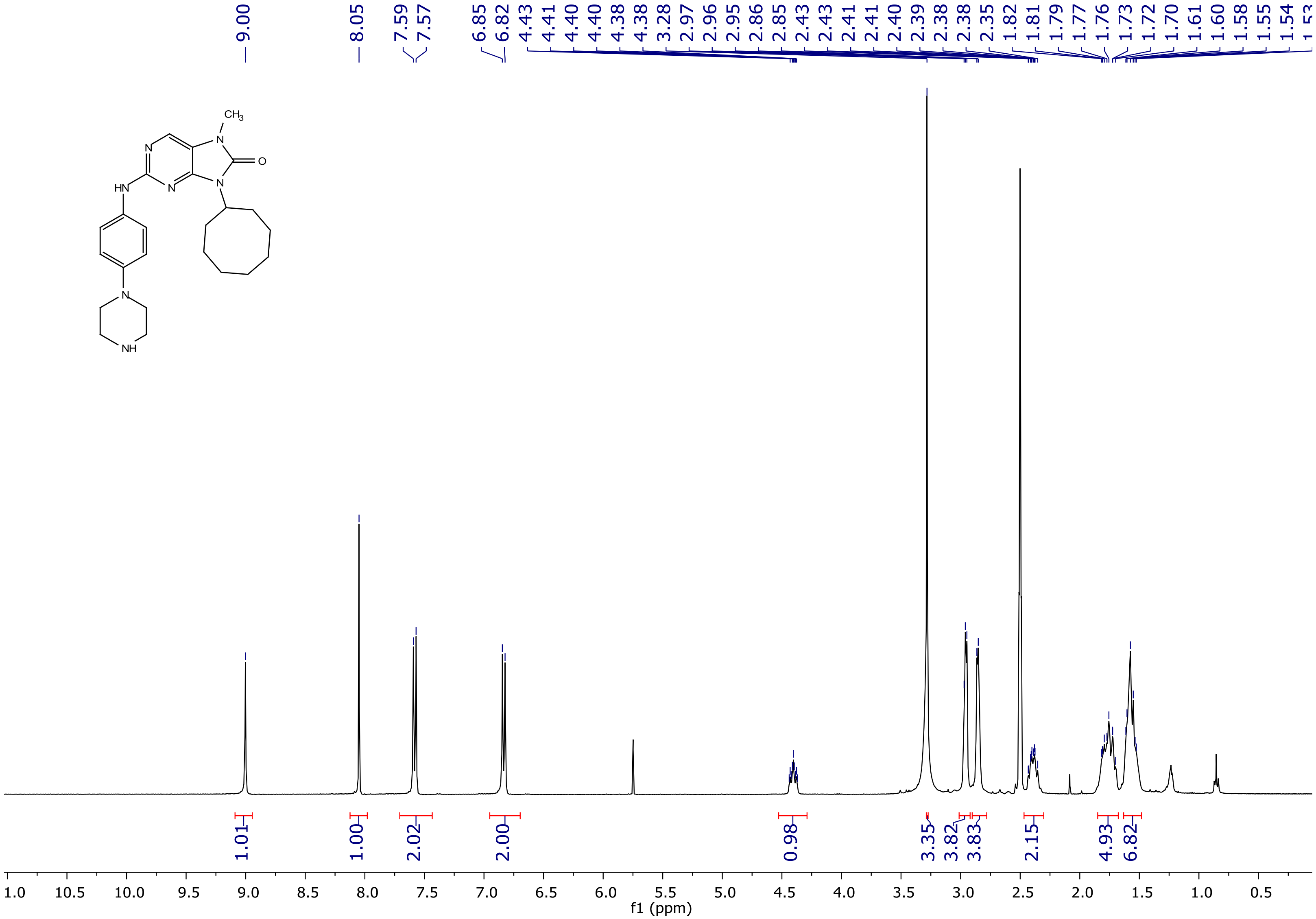

$^{1}$H NMR spectrum of **14i**

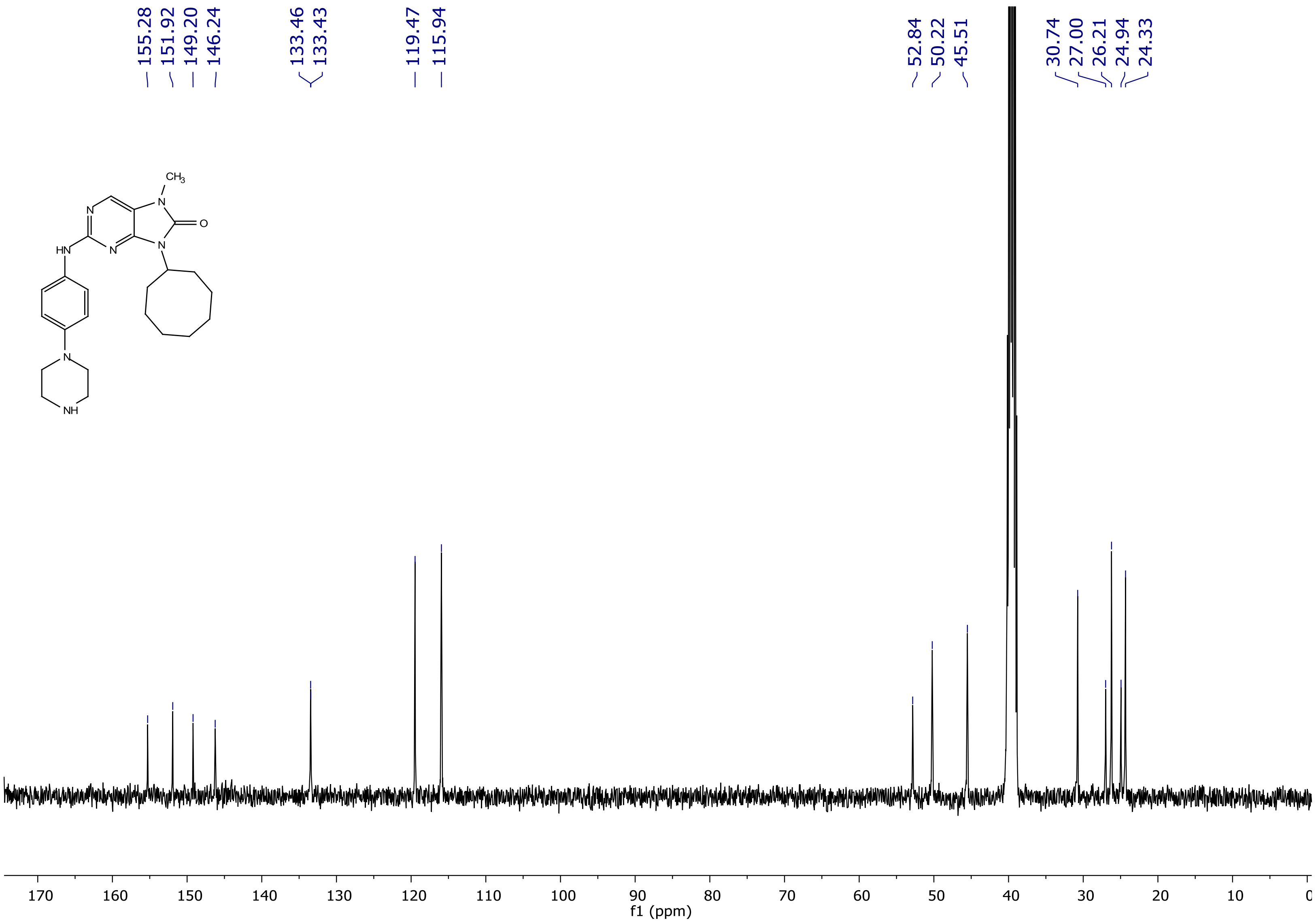

[1]H NMR spectrum of **14j**

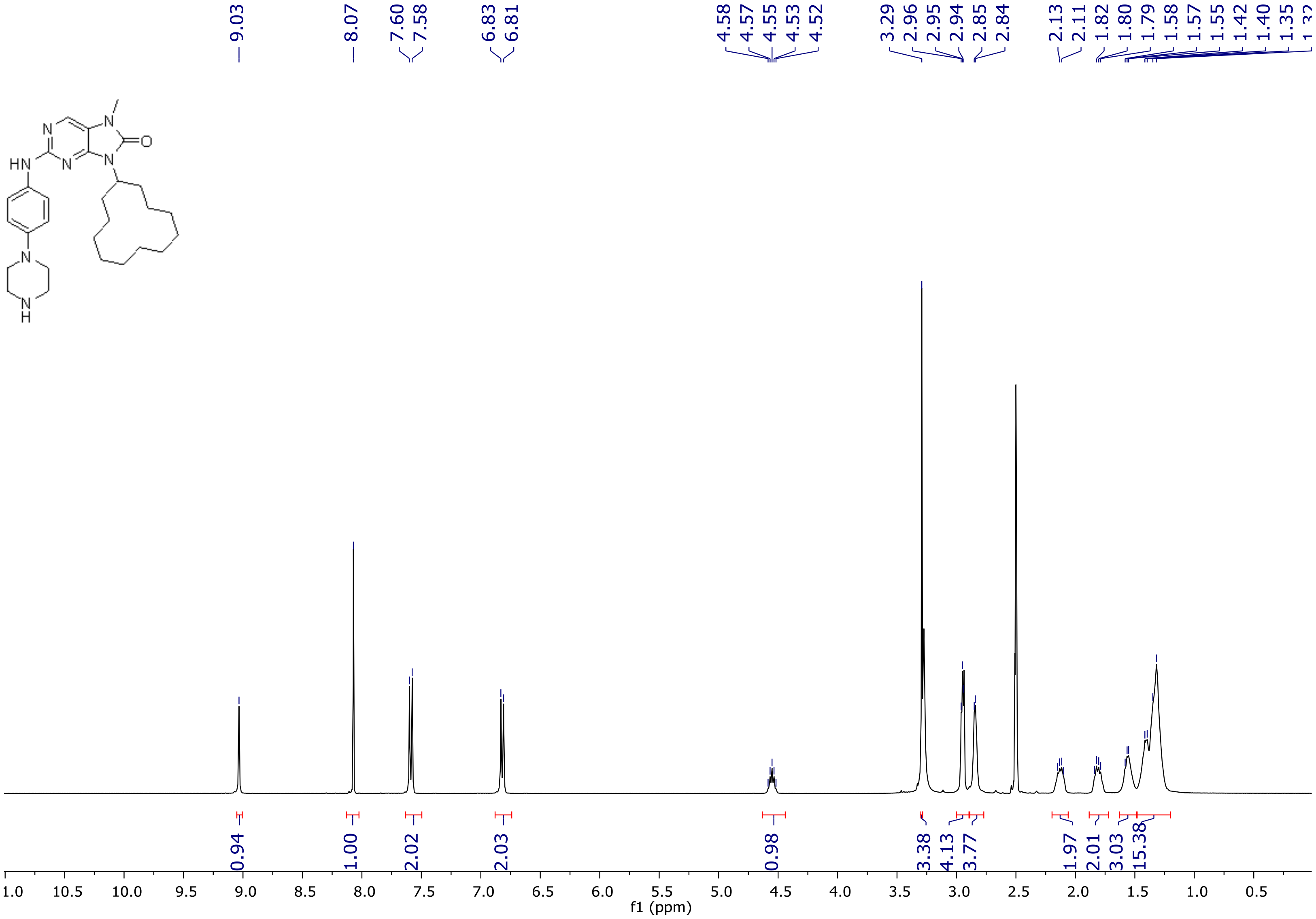

$^{1}$H NMR spectrum of **14j**

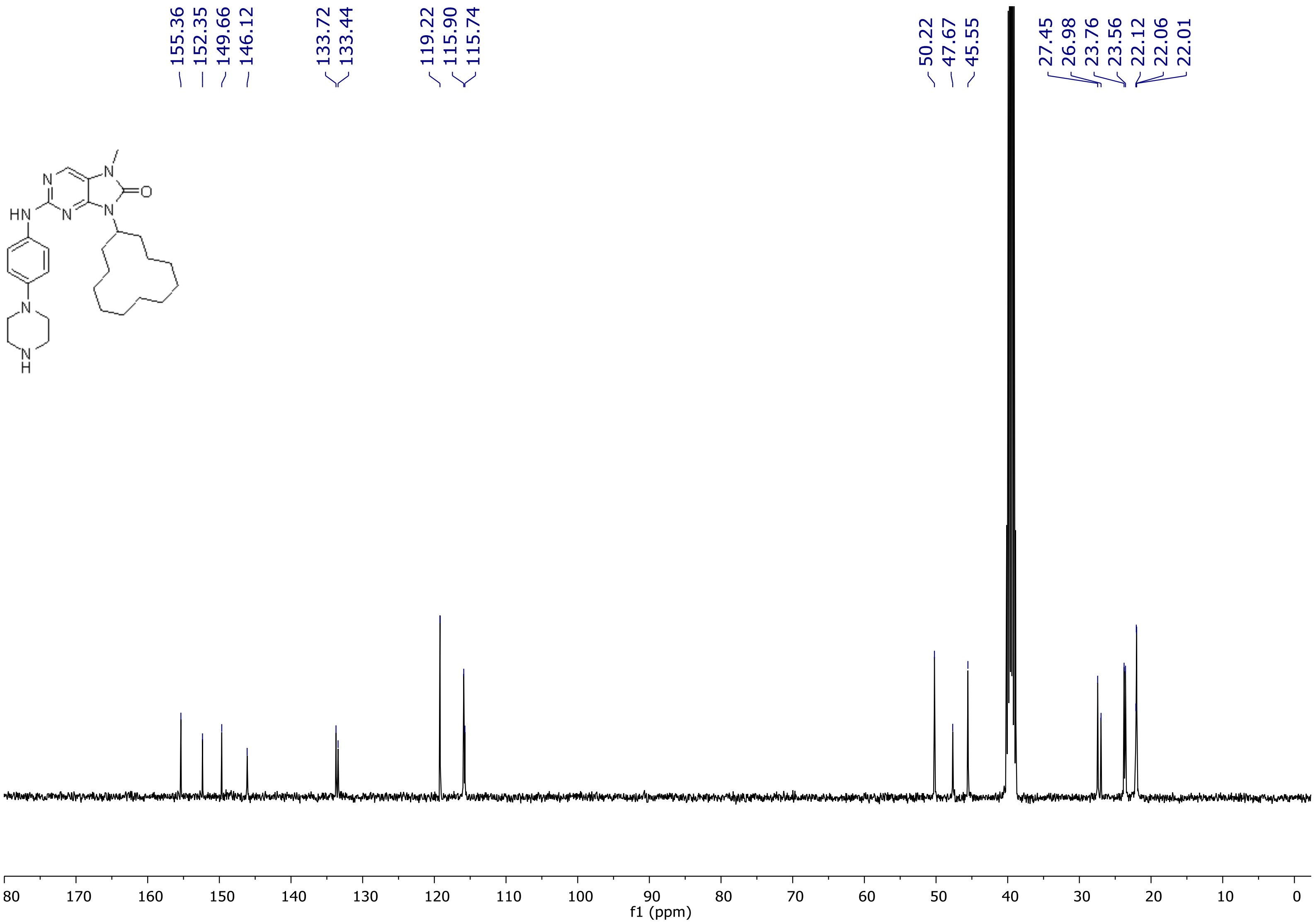

[1]H NMR spectrum of **14k**

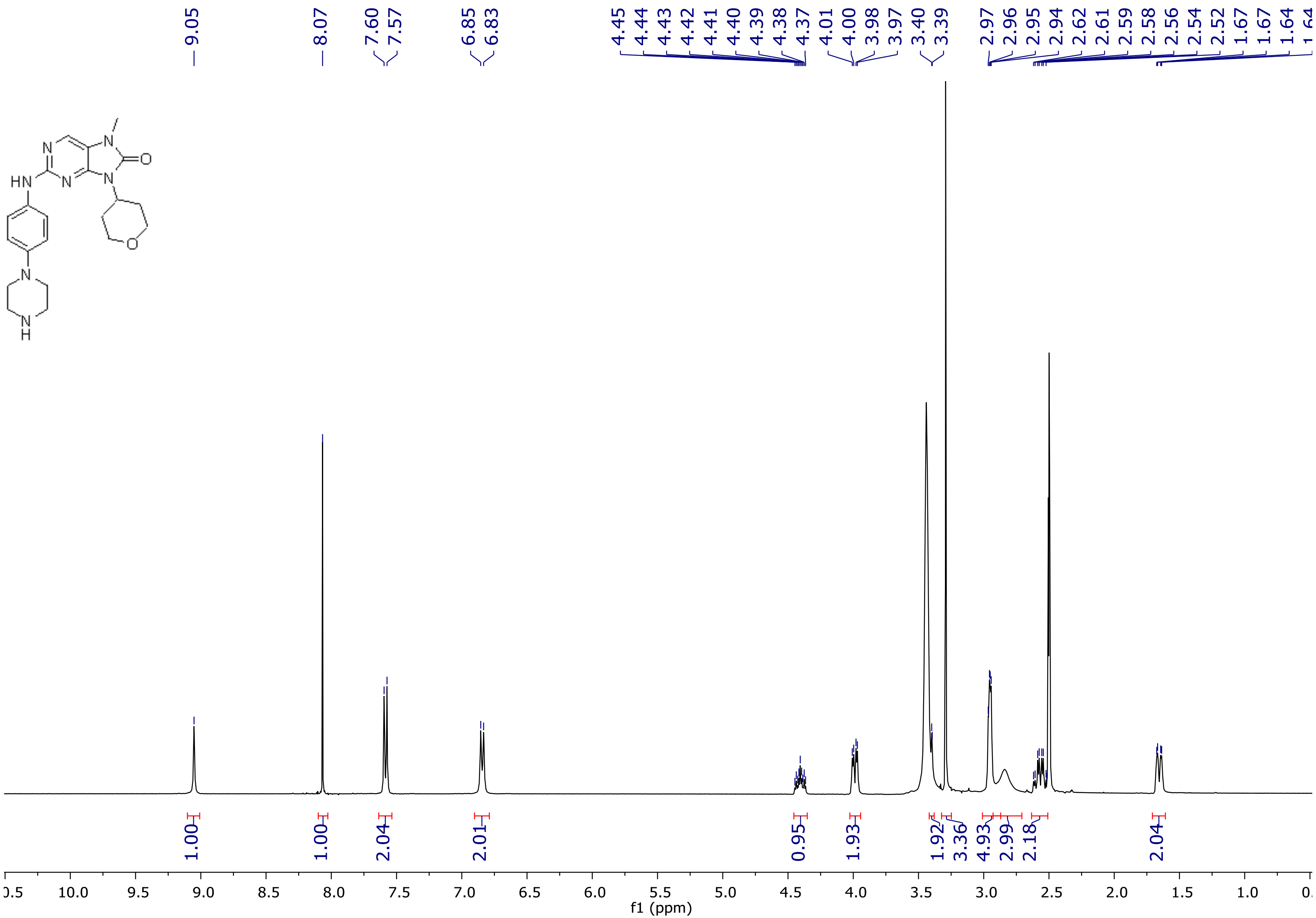

[1]H NMR spectrum of **14k**

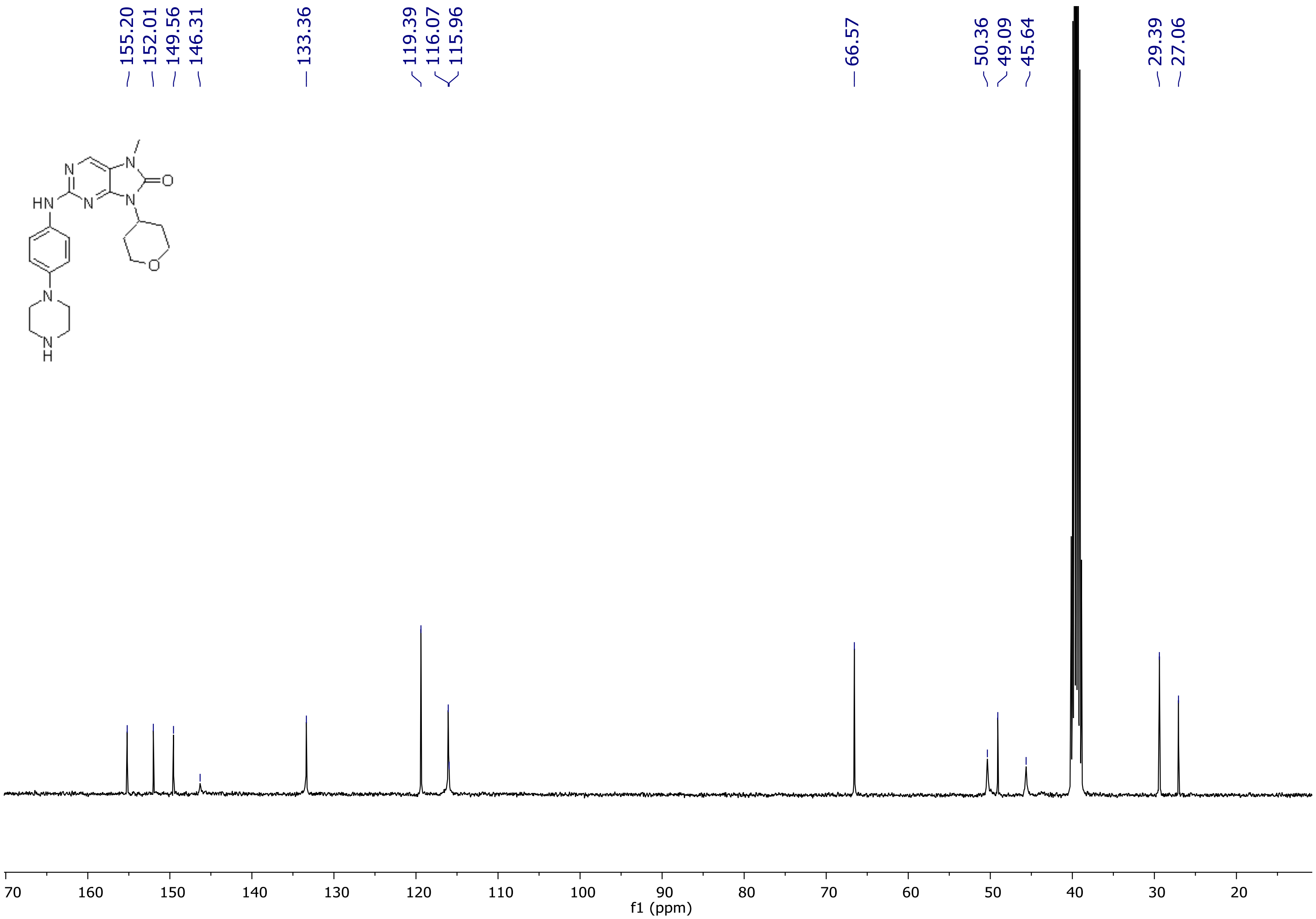

[1]H NMR spectrum of **14l**

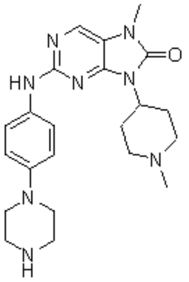

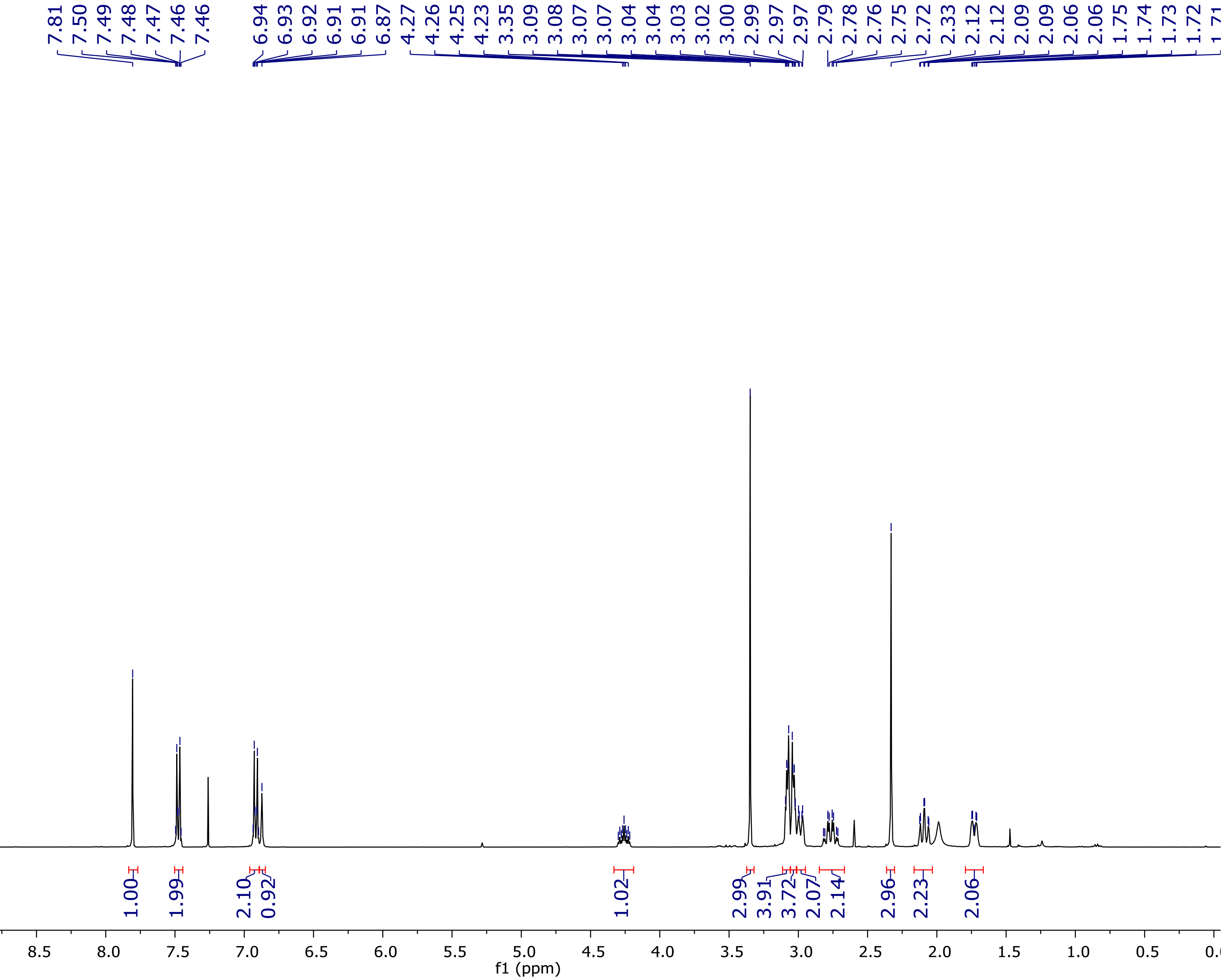

[1]H NMR spectrum of **14l**

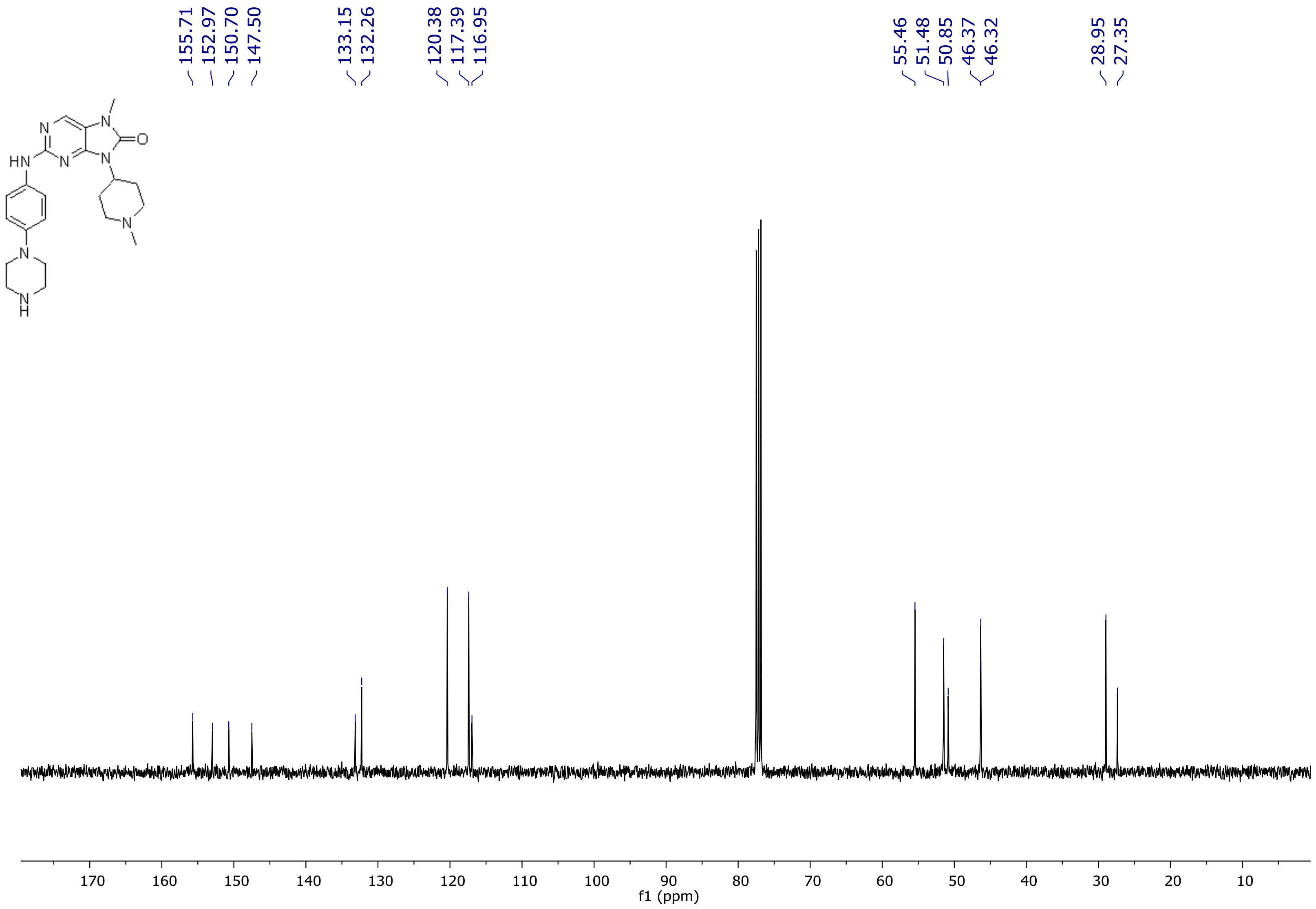

[1]H NMR spectrum of **14m**

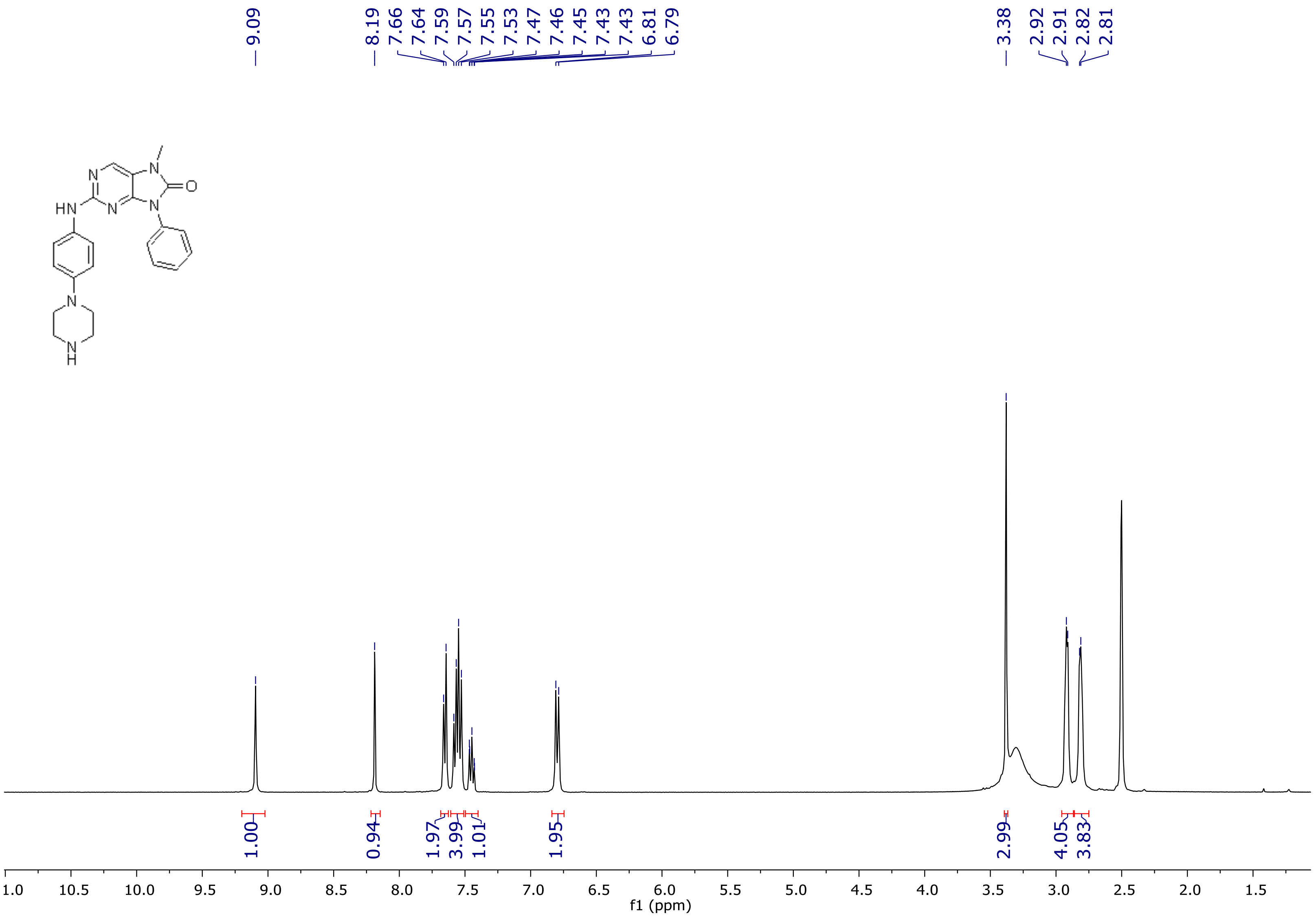

[1]H NMR spectrum of **14m**

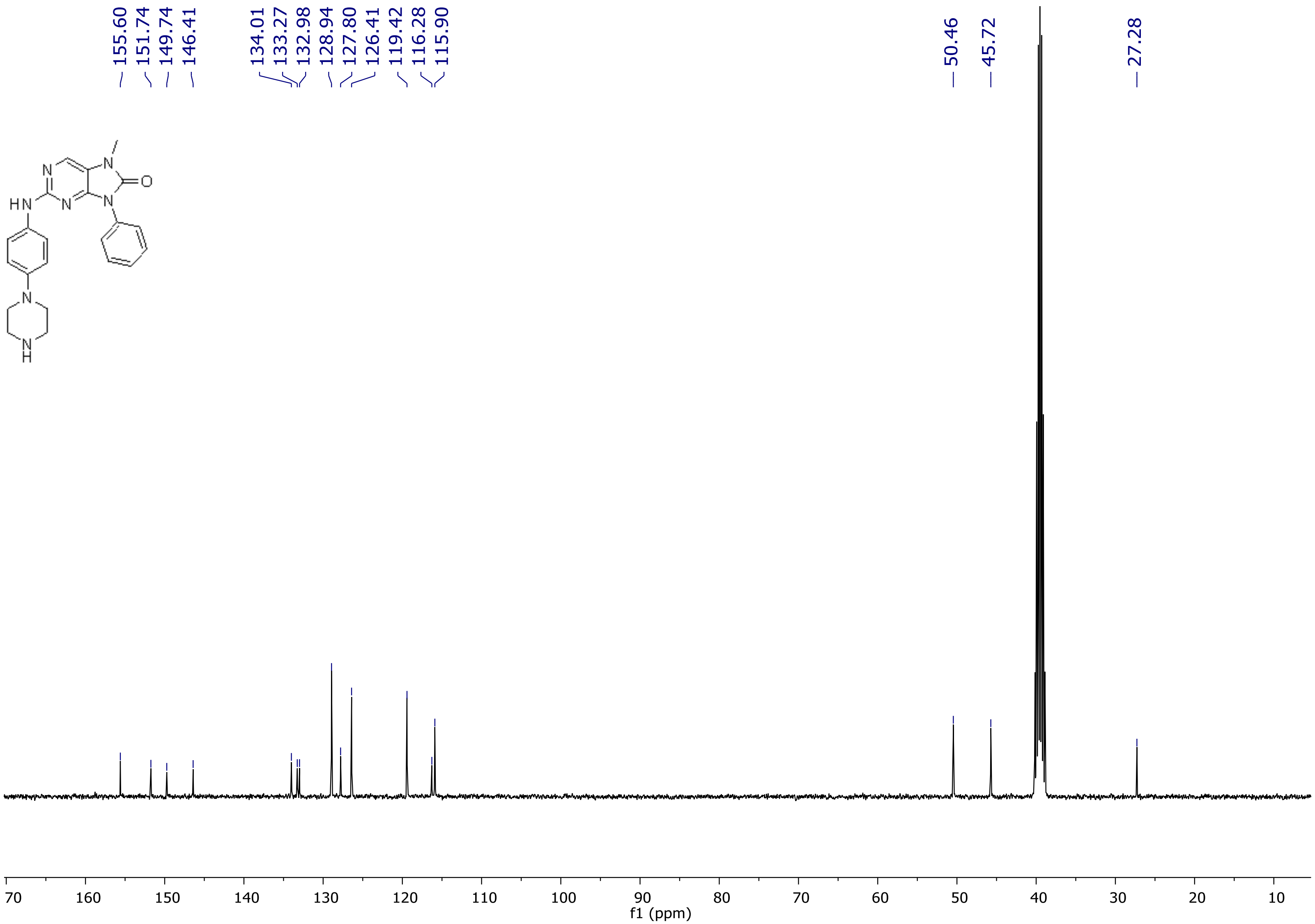

$^{1}$H NMR spectrum of **14n**

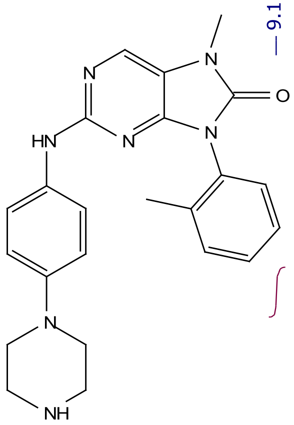

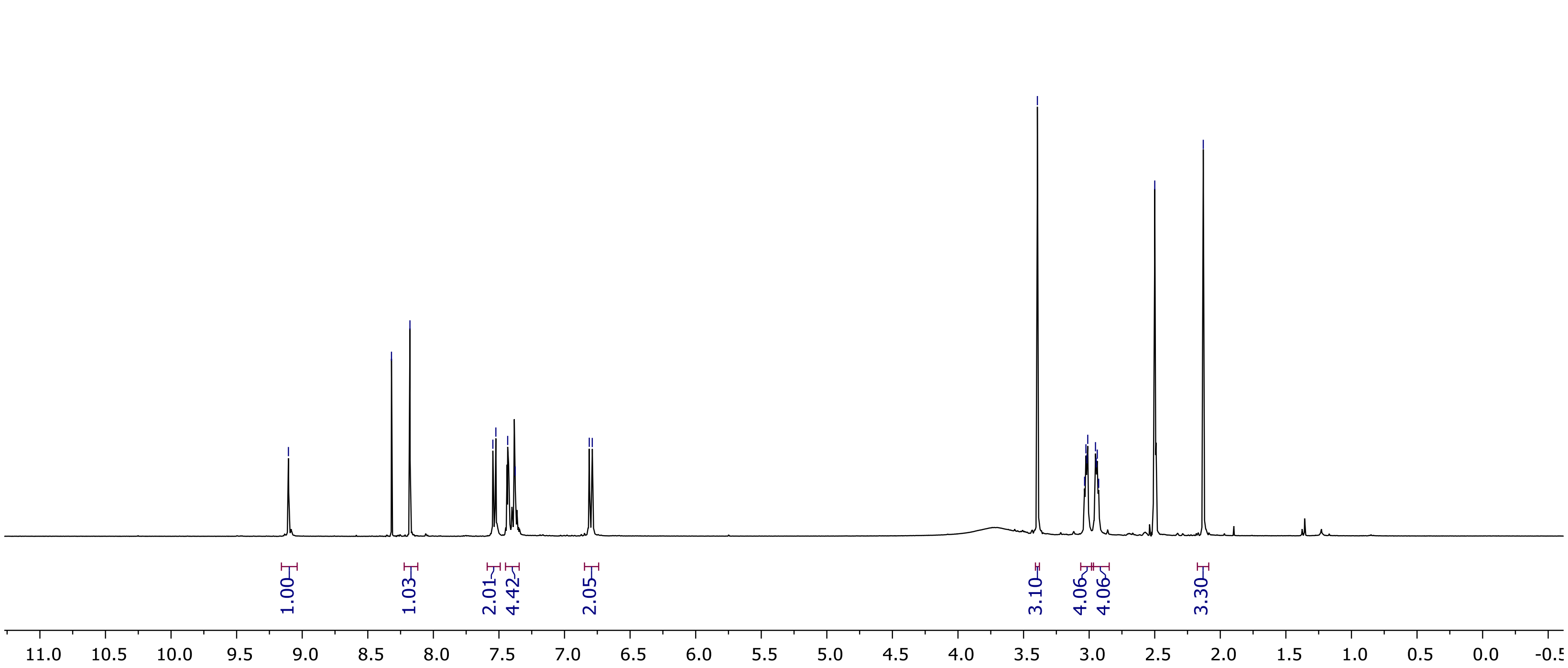

[13]C NMR spectrum of **14n**

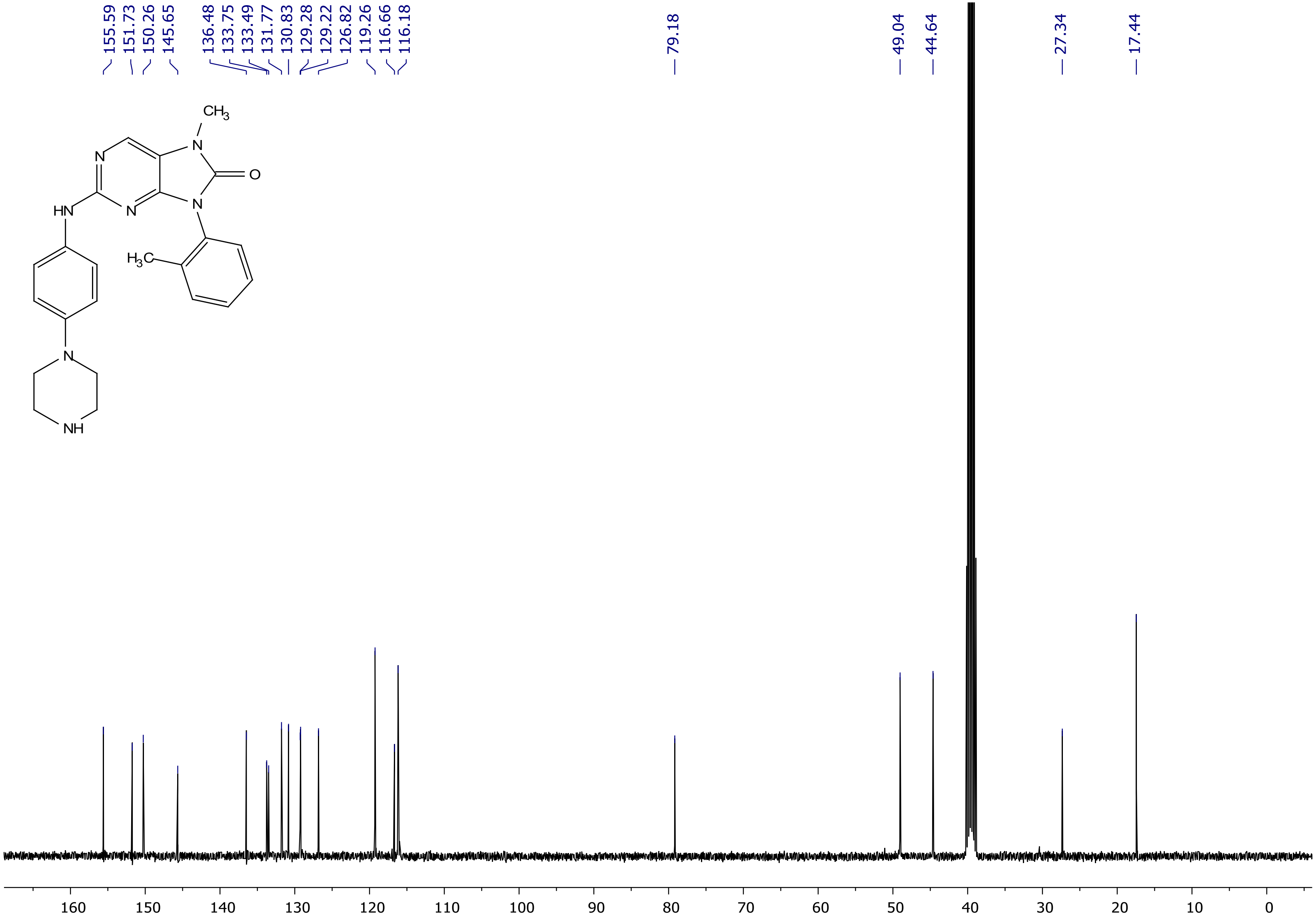

[1]H NMR spectrum of **14o**

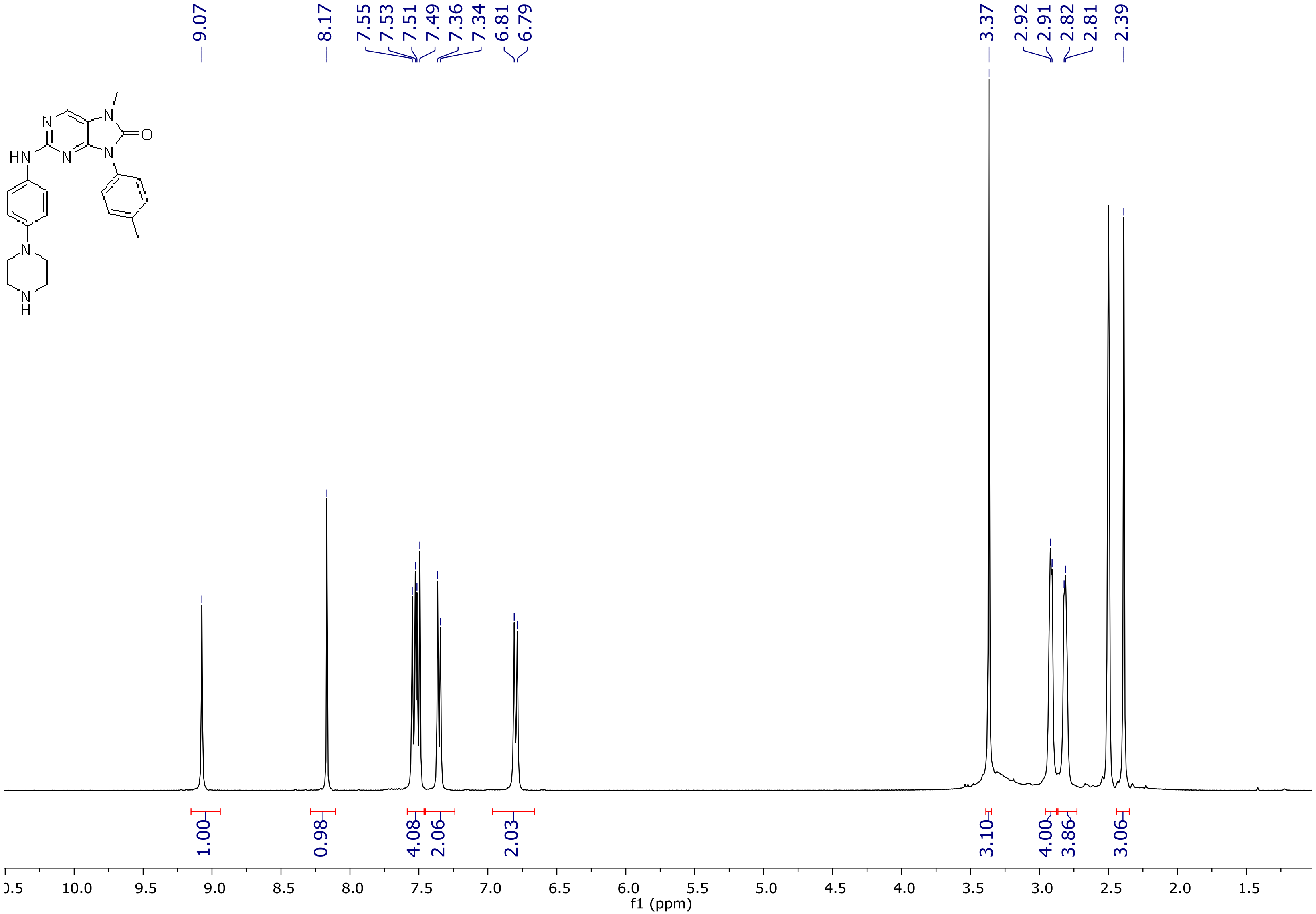

[13]C NMR spectrum of **14o**

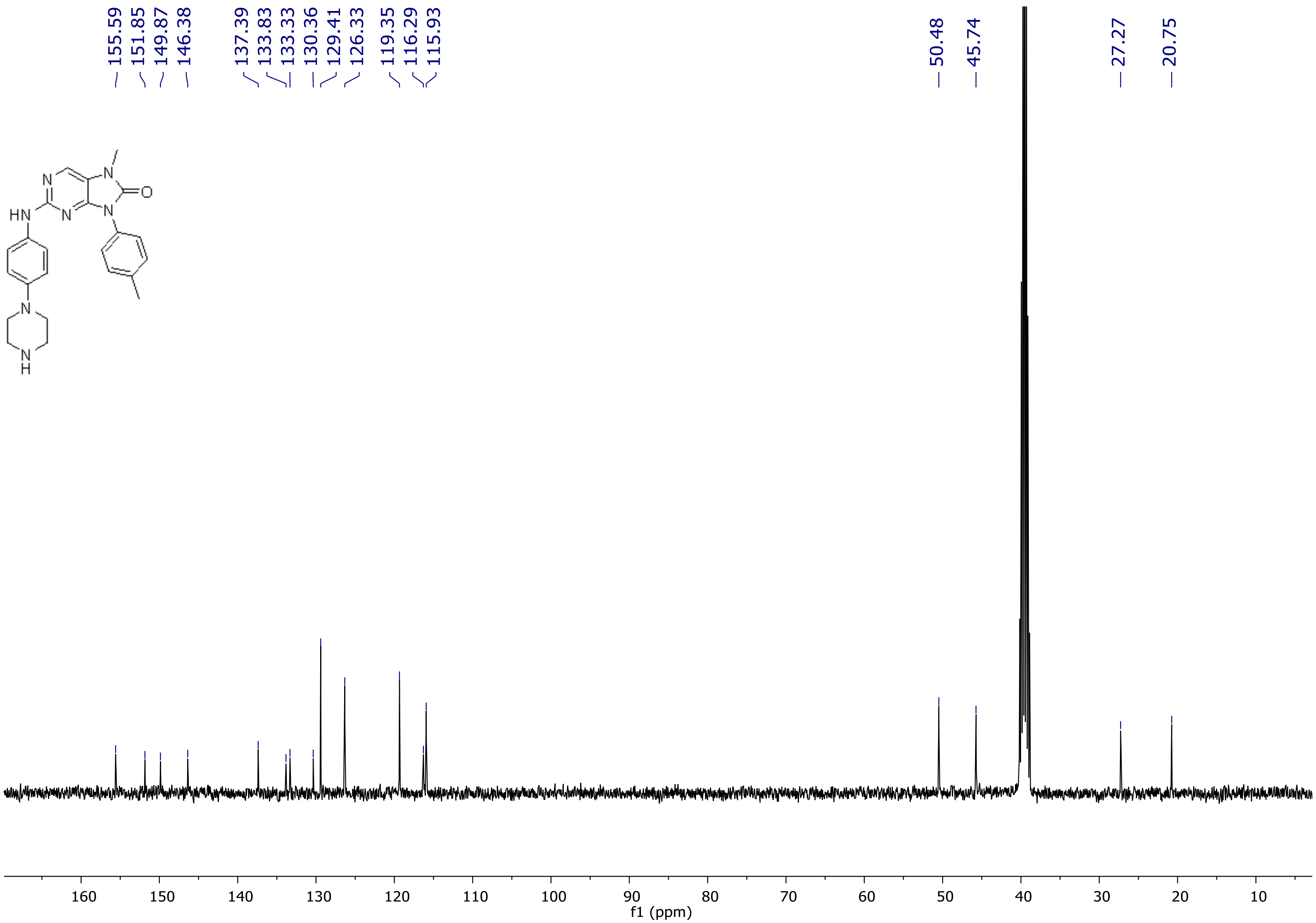

[1]H NMR spectrum of **14p**

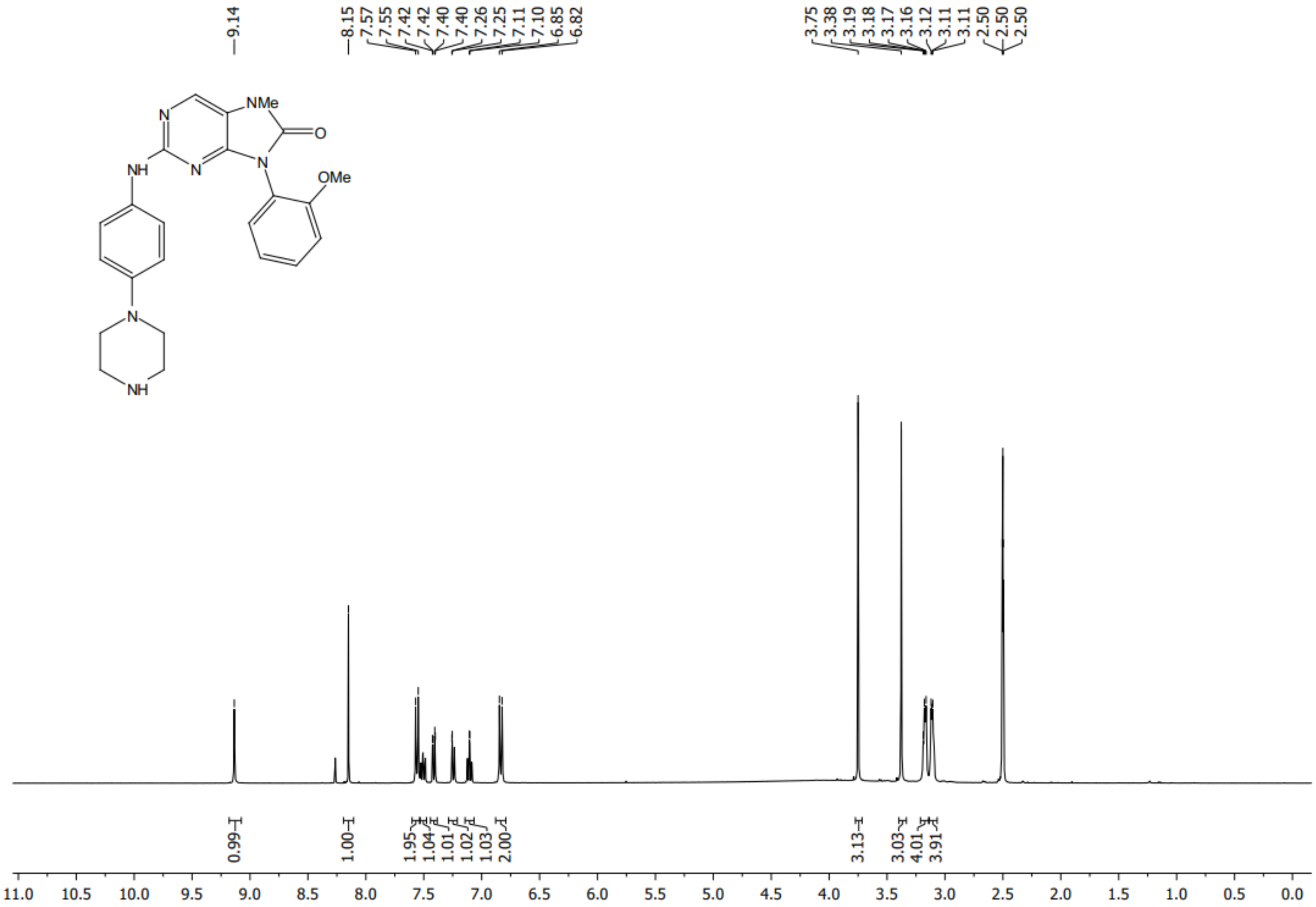

[13]C NMR spectrum of **14p**

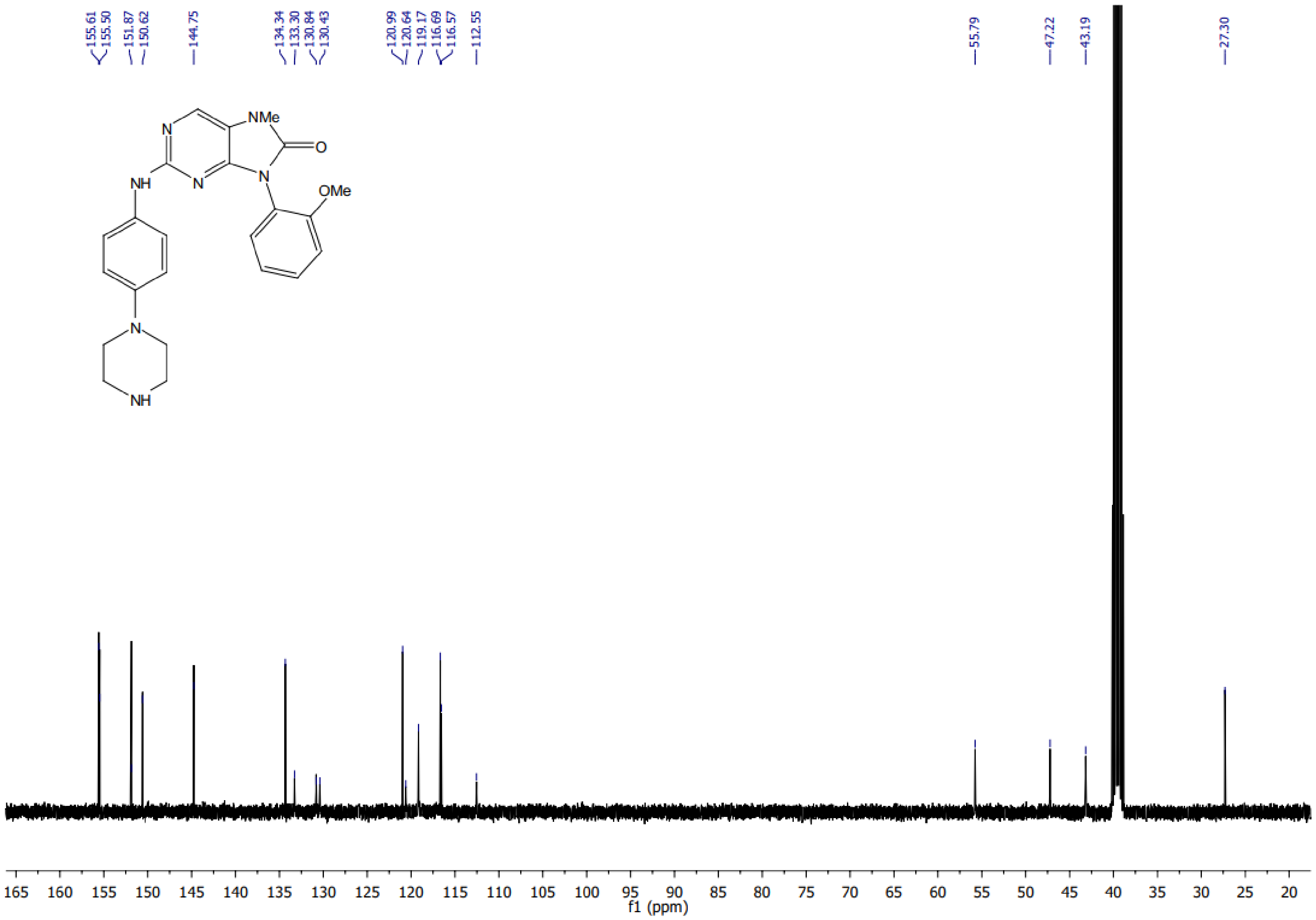

<sup></sup>

[1]H NMR spectrum of **14q**

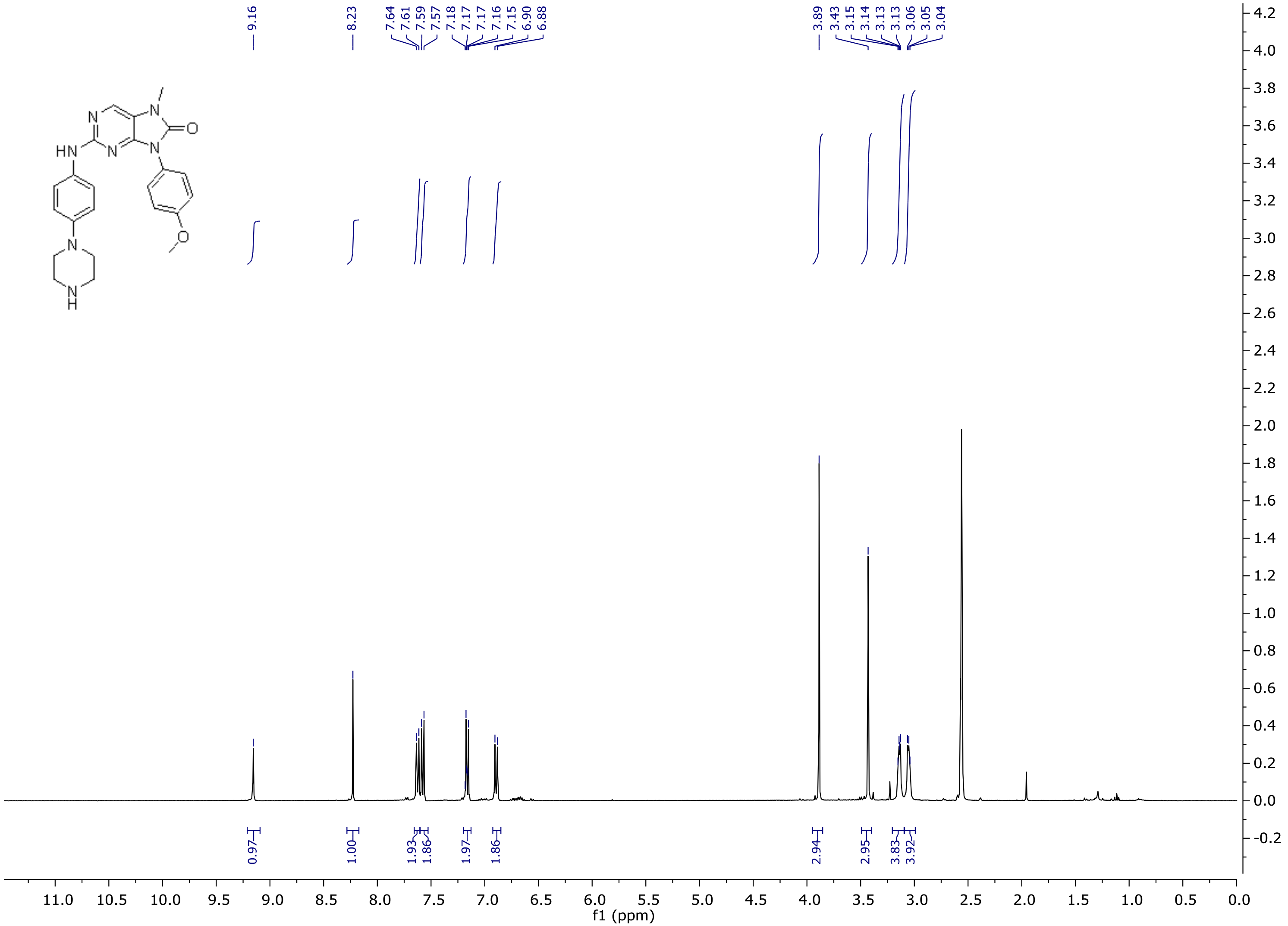

[13]C NMR spectrum of **14q**

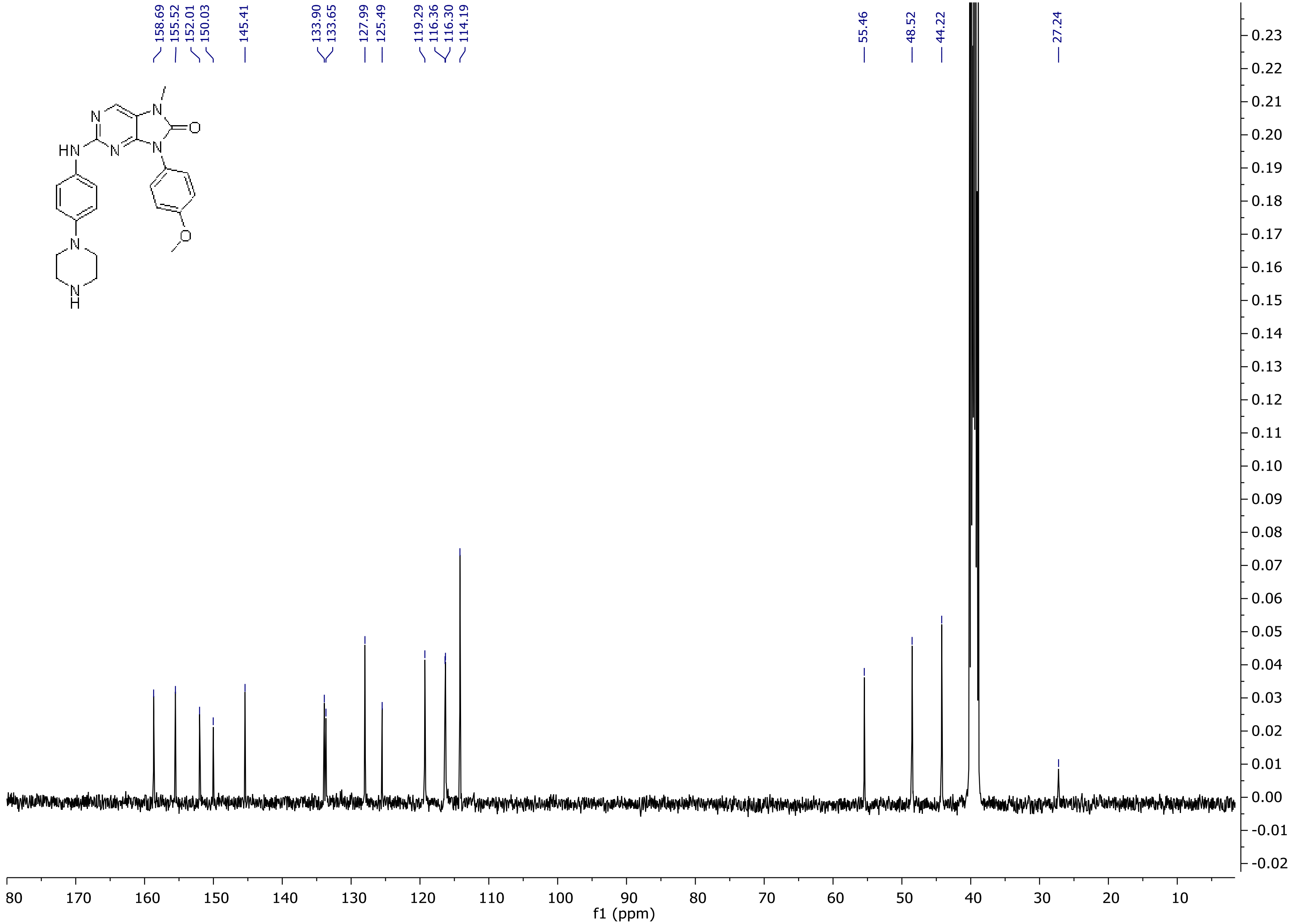

<sup></sup>[1]H NMR spectrum of **14r**

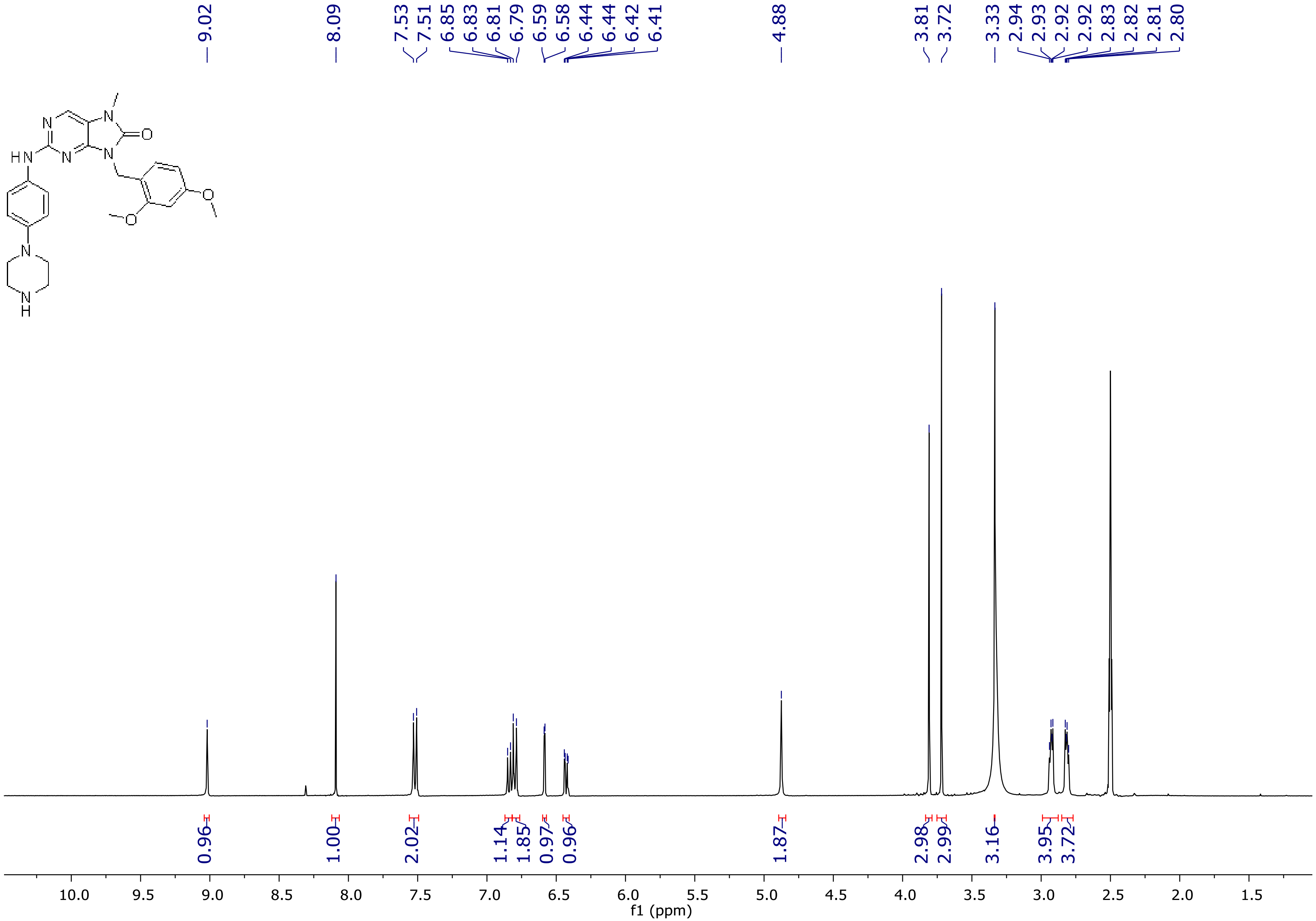

$^{13}$C NMR spectrum of **14r**

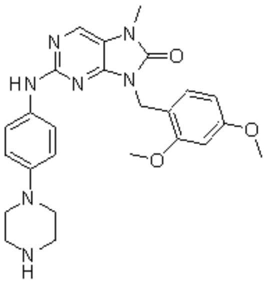

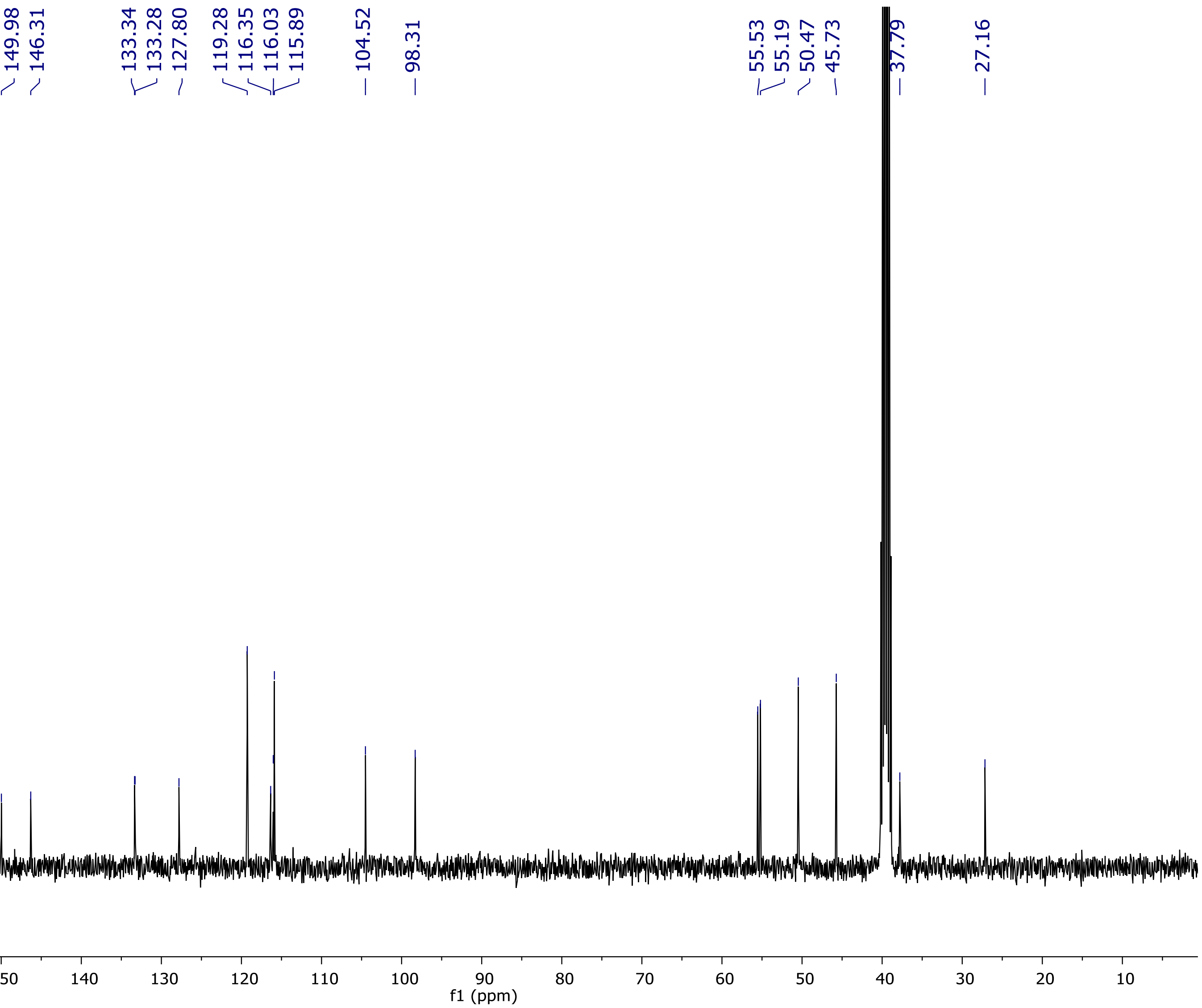

[1]H NMR spectrum of **14s**

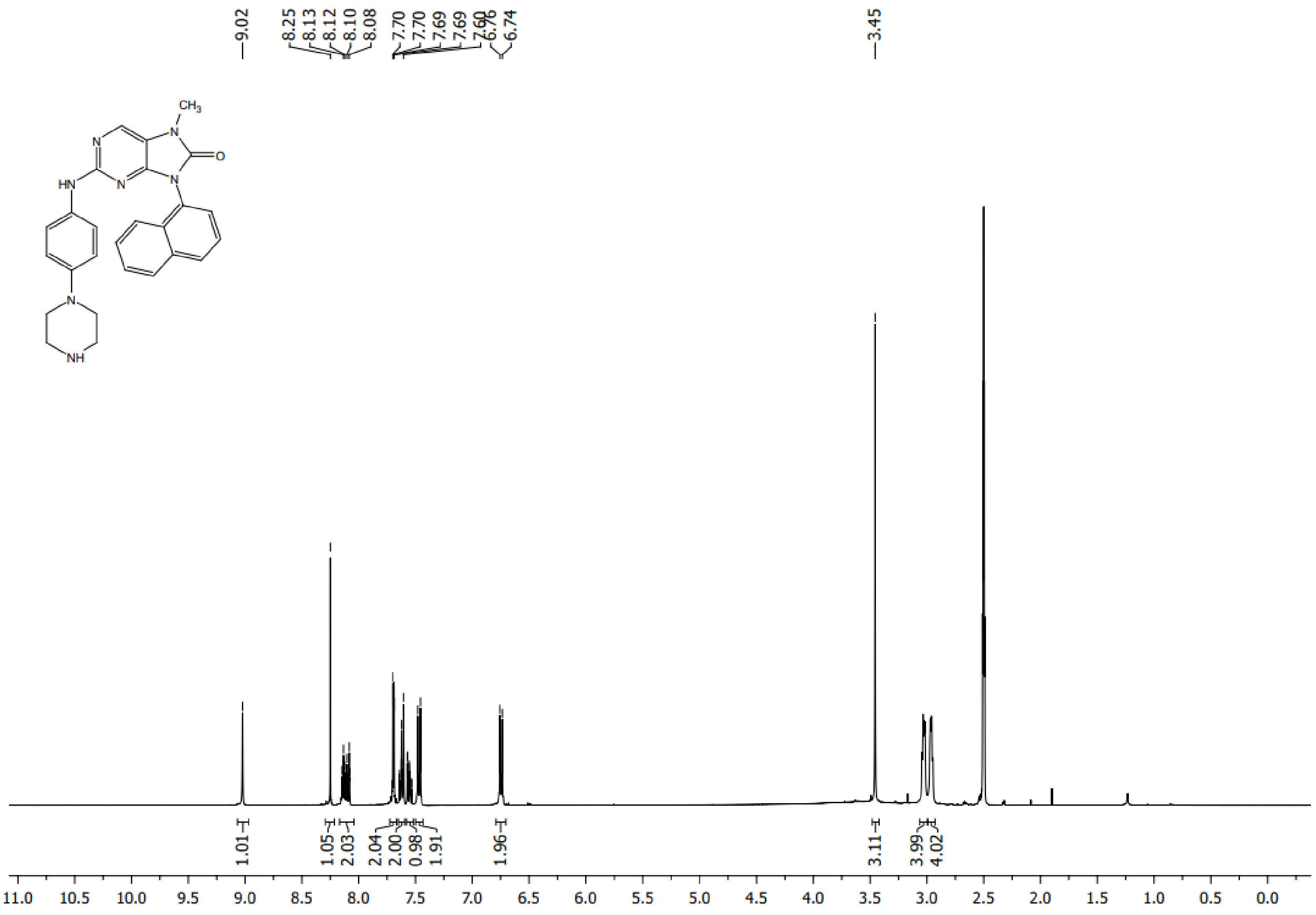

$^{13}$C NMR spectrum of **14s**

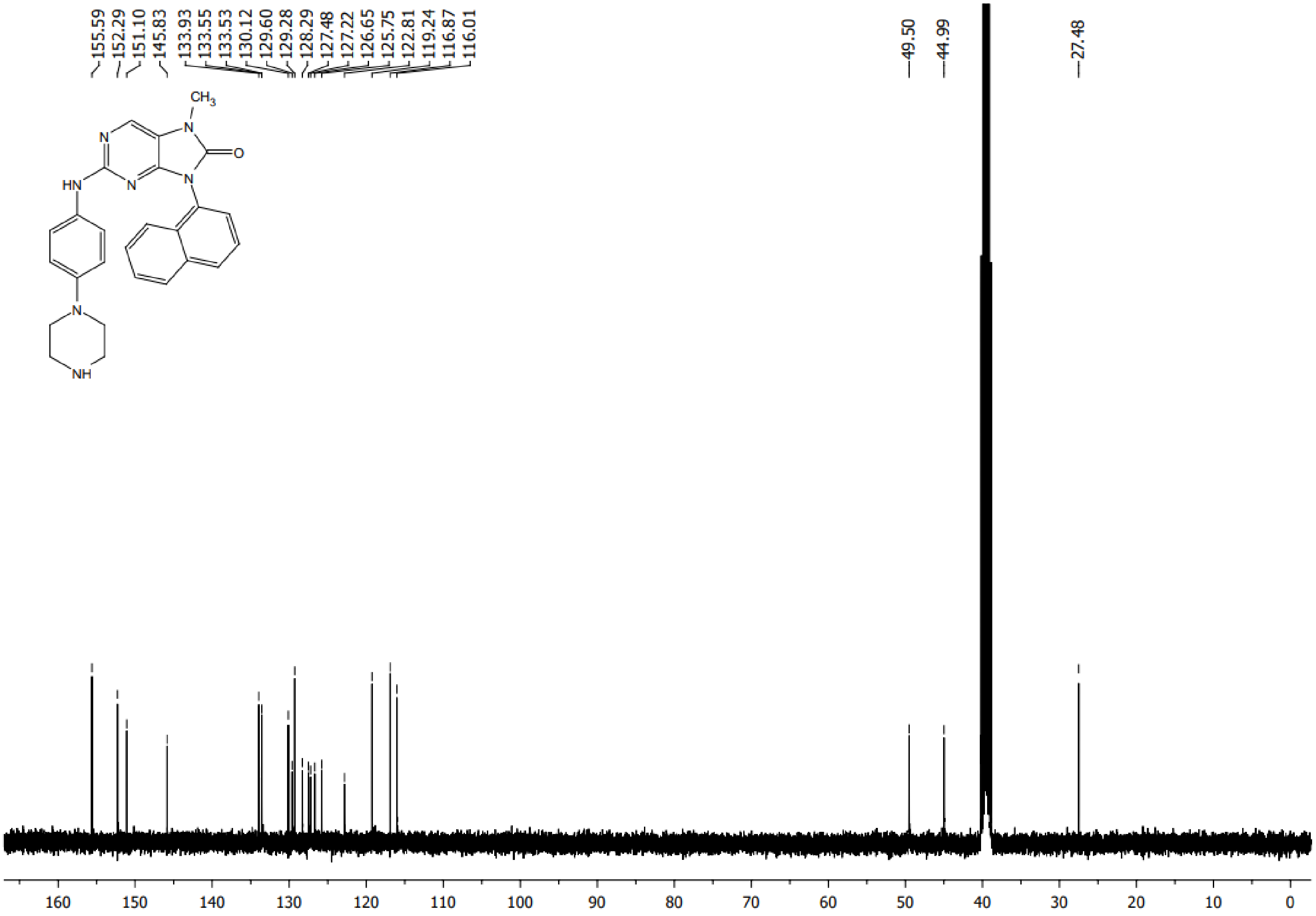

[1]H NMR spectrum of **14t**

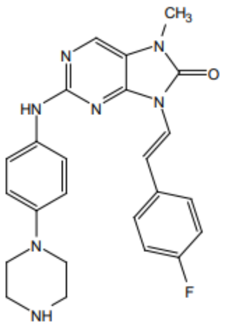

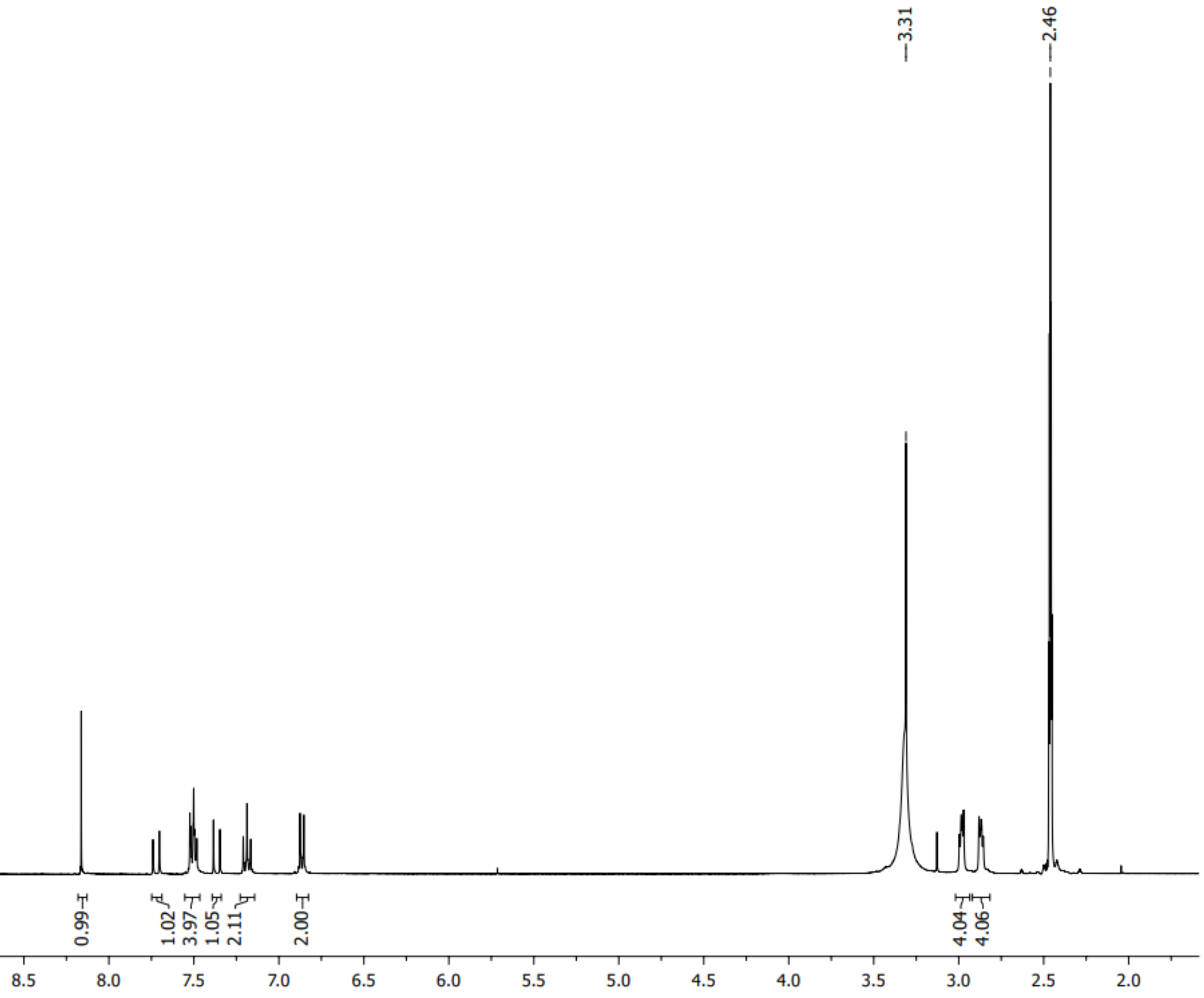

$^{13}$C NMR spectrum of **14t**

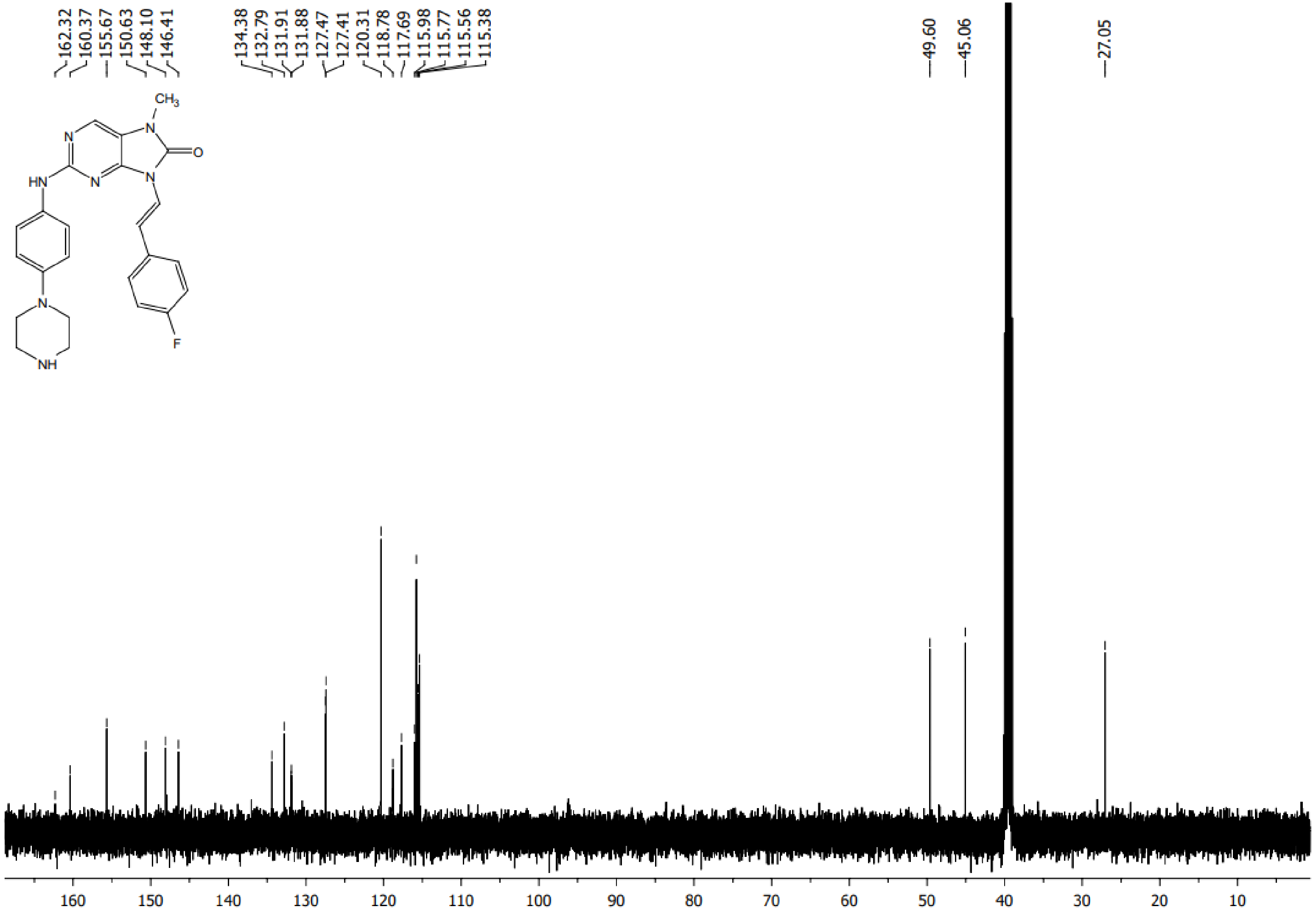

[1]H NMR spectrum of **15a**

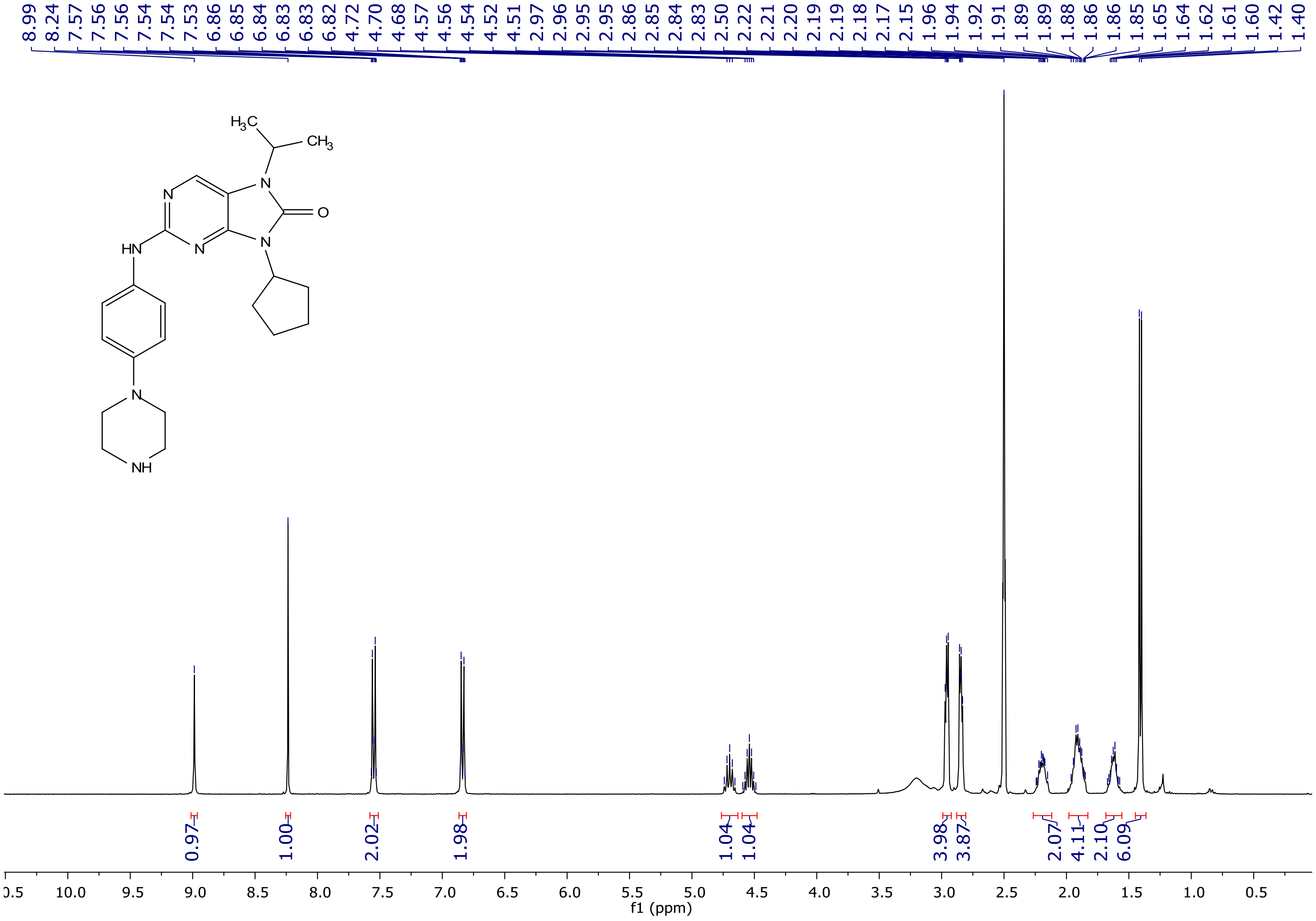

[13]C NMR spectrum of **15a**

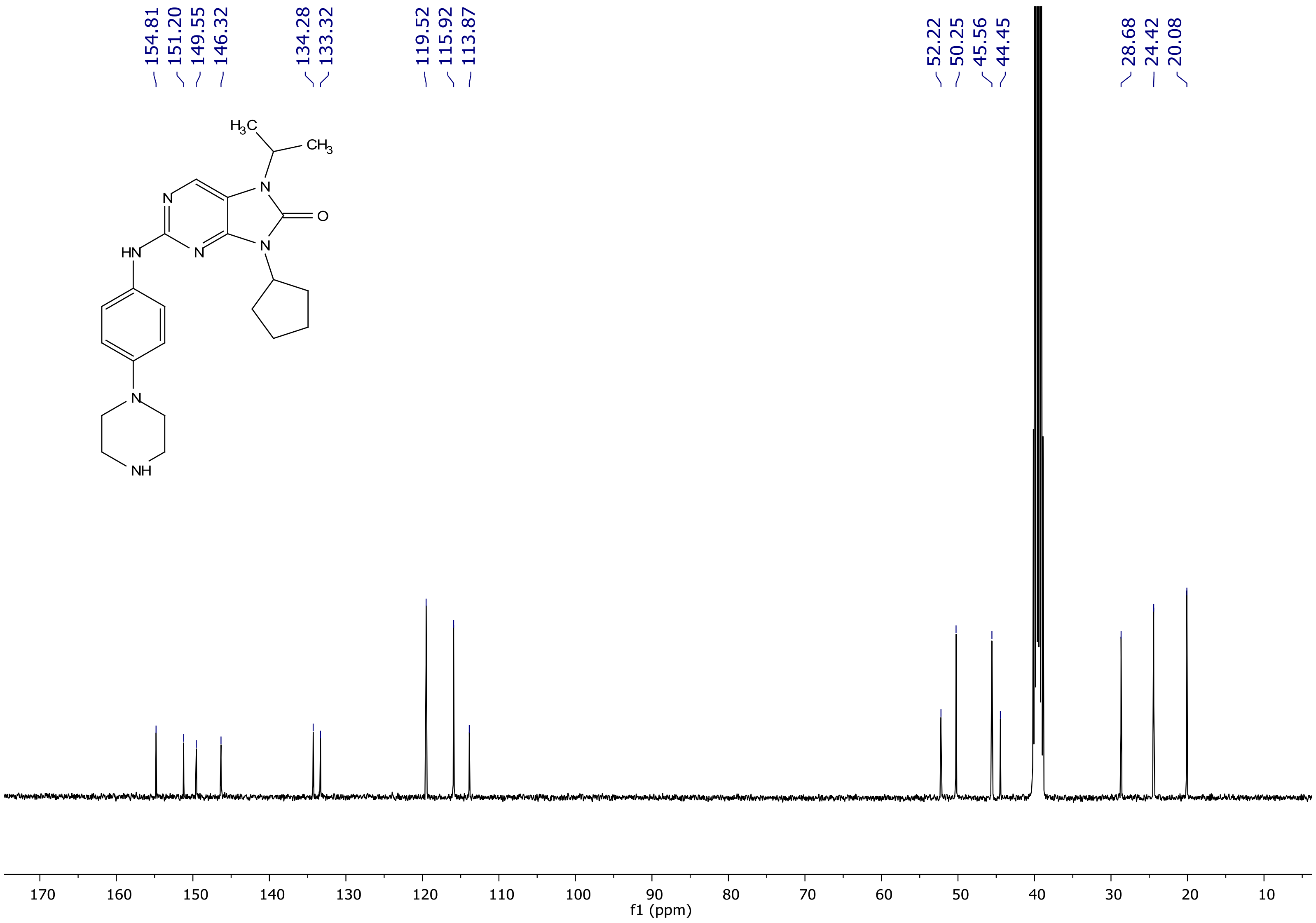

[1]H NMR spectrum of **15b**

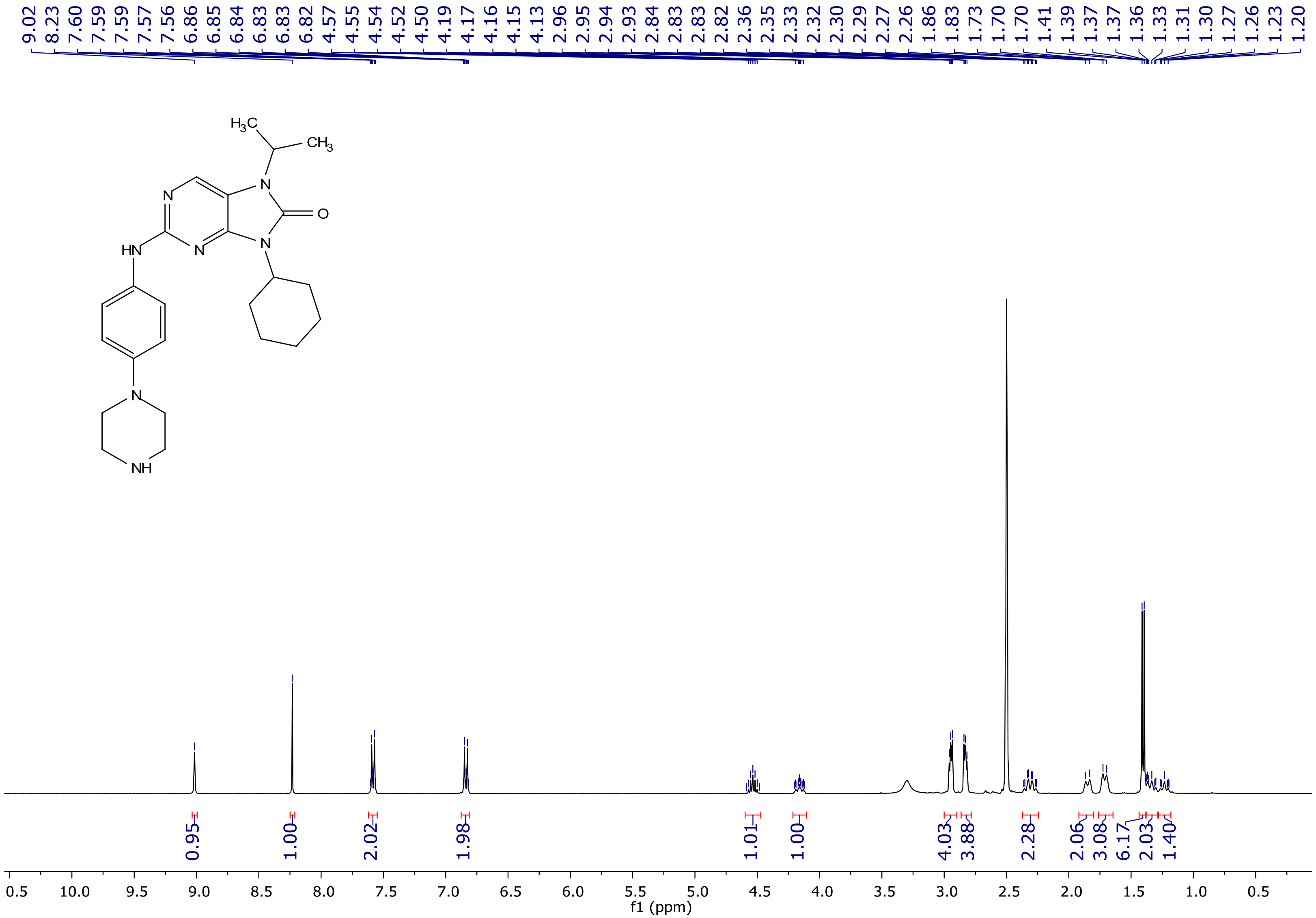

$^{13}C$ NMR spectrum of **15b**

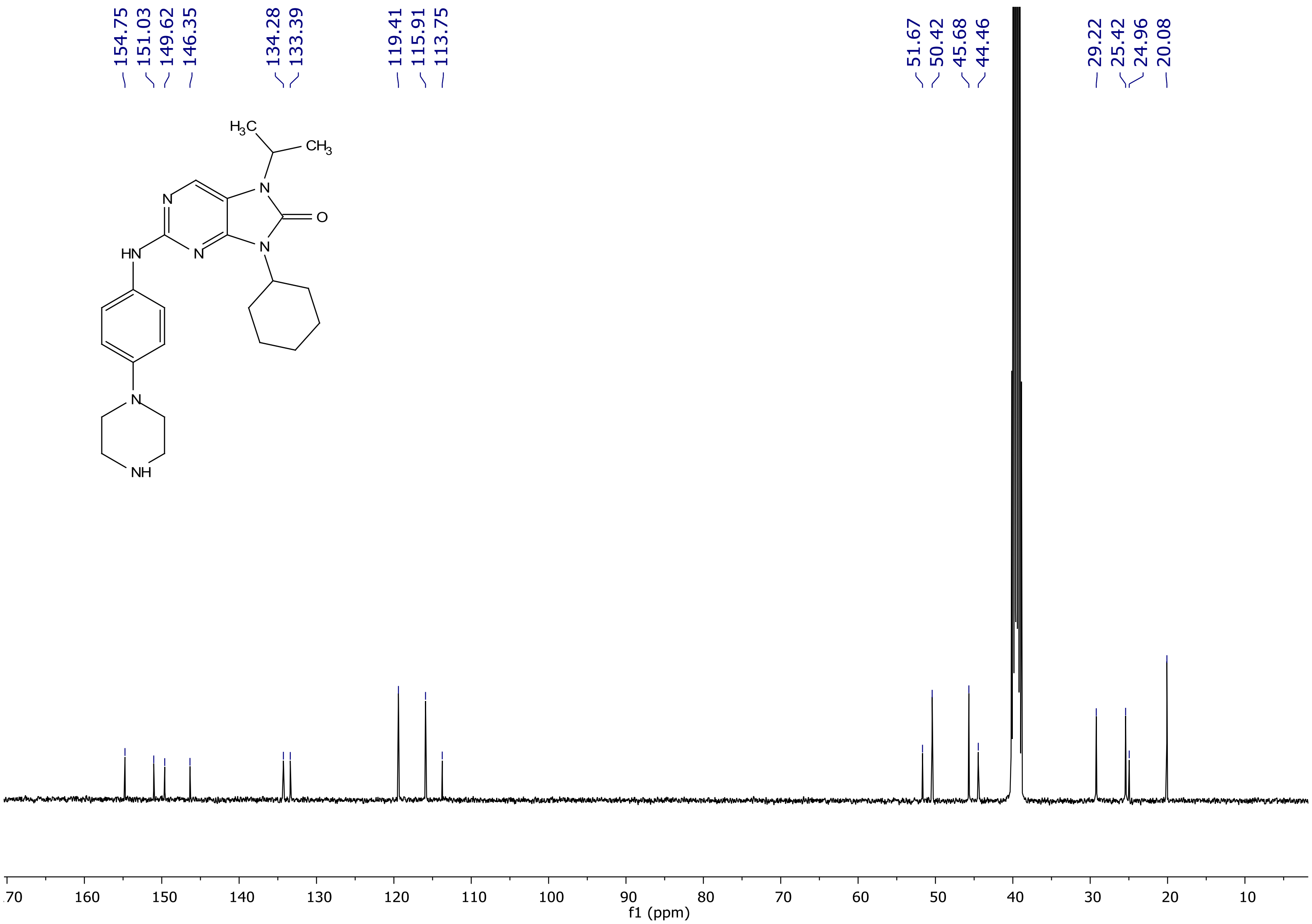
154.75
151.03
149.62
146.35
134.28
133.39
119.41
115.91
113.75
51.67
50.42
45.68
44.46
29.22
25.42
24.96
20.08
170
160
150
140
130
120
110
100
90
80
70
60
50
40
30
20
10
f1 (ppm)

[1]H NMR spectrum of **15c**

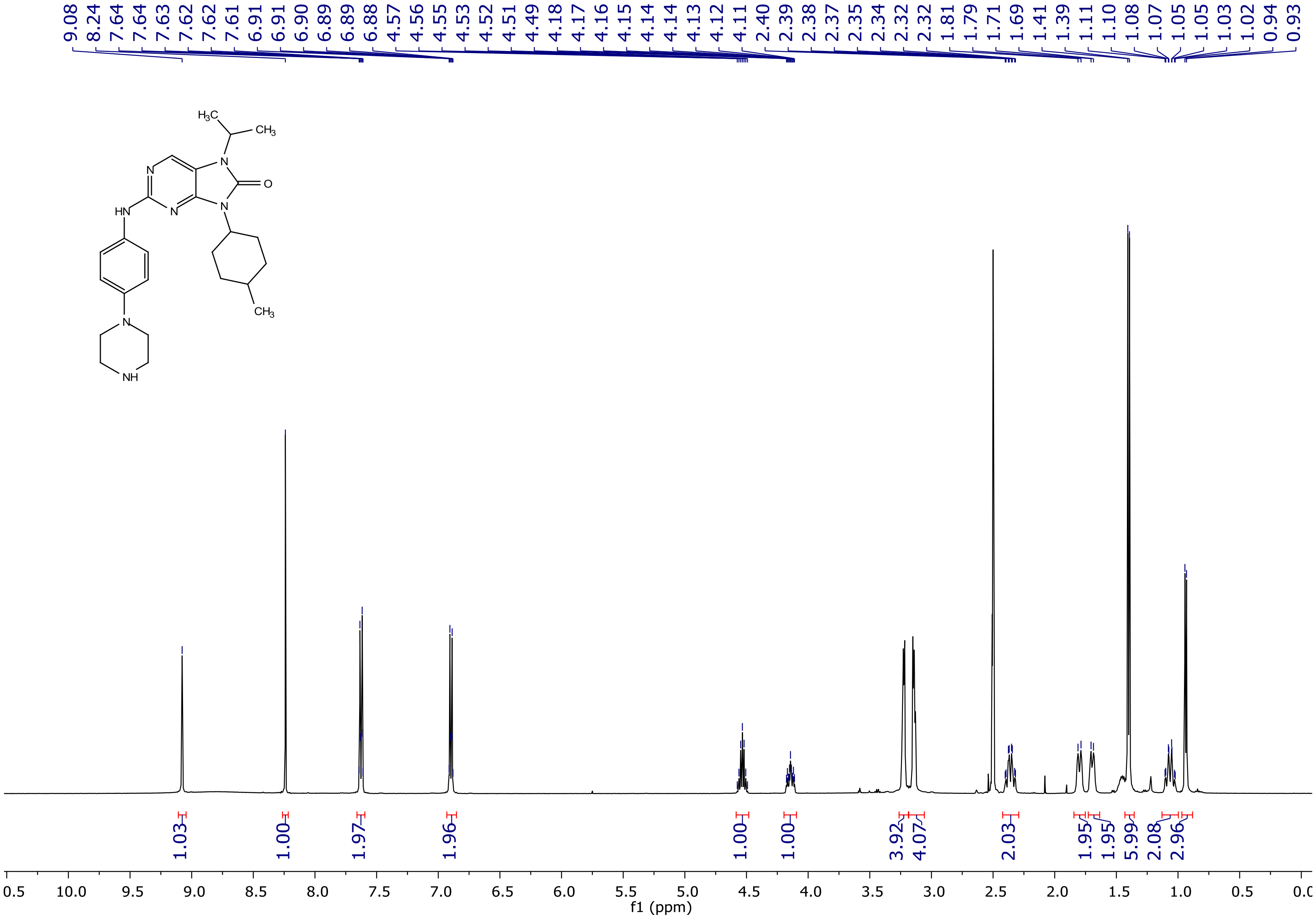

[13]C NMR spectrum of **15c**

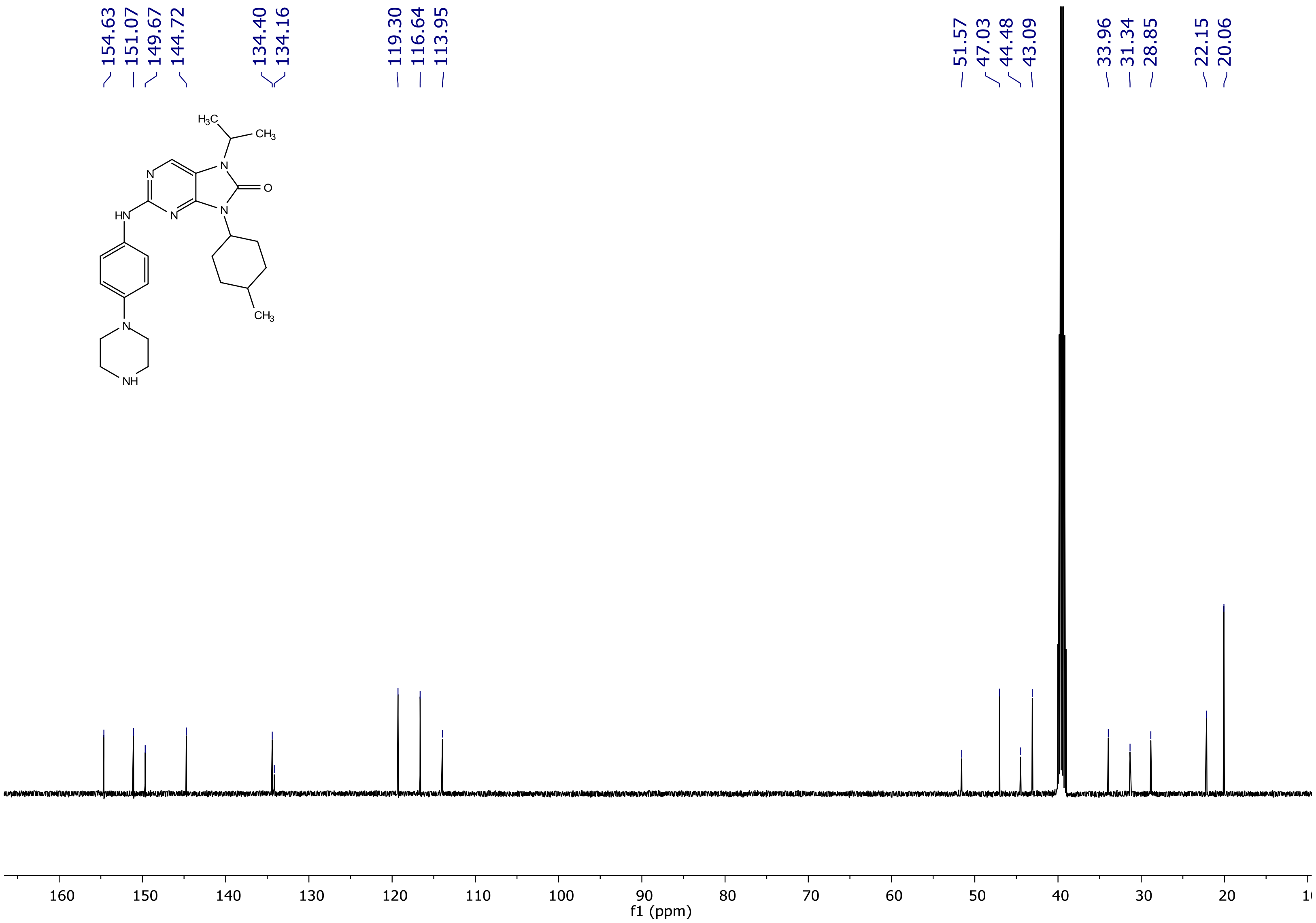

<sup></sup>

[1]H NMR spectrum of **15d**

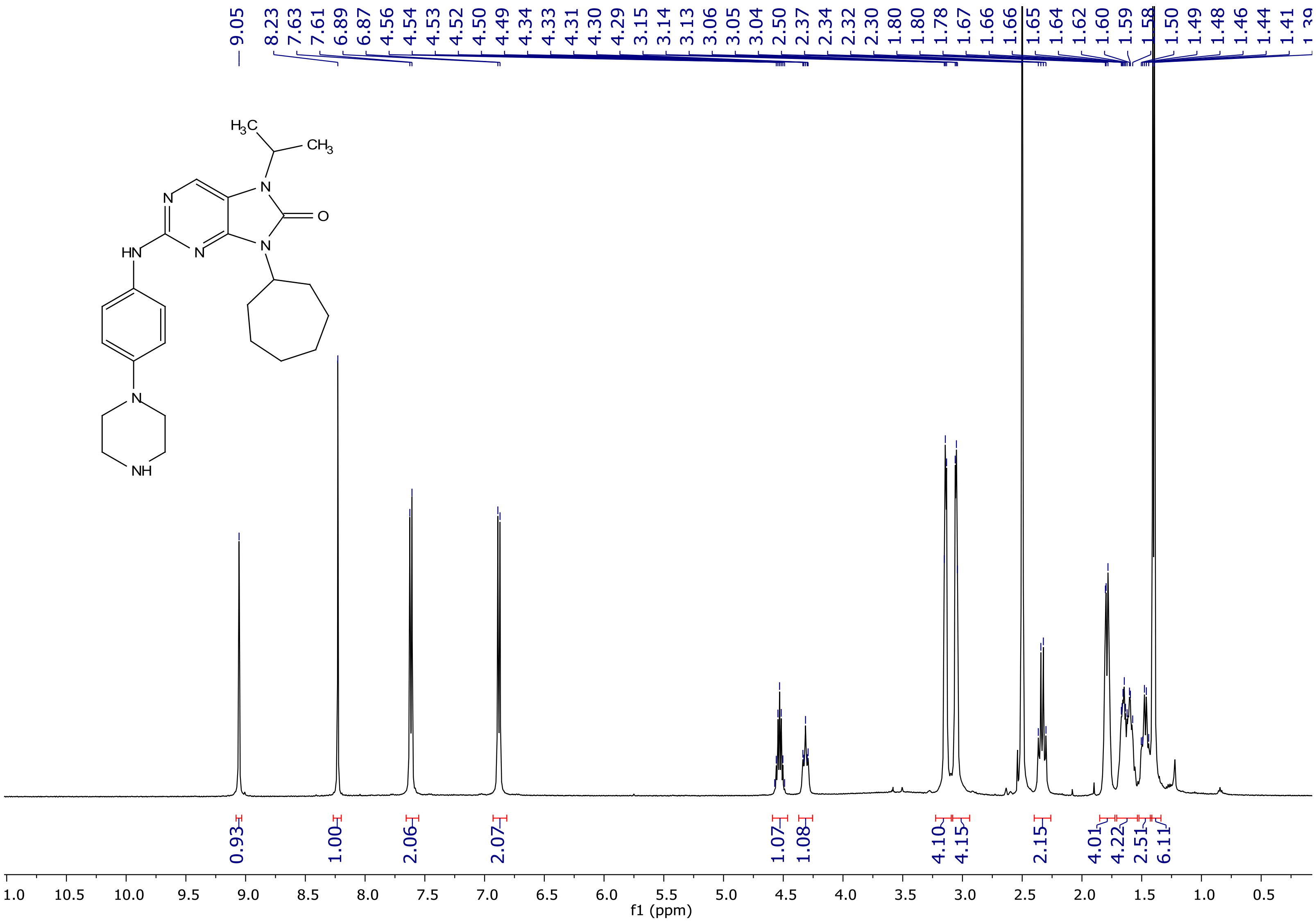

[13]C NMR spectrum of **15d**

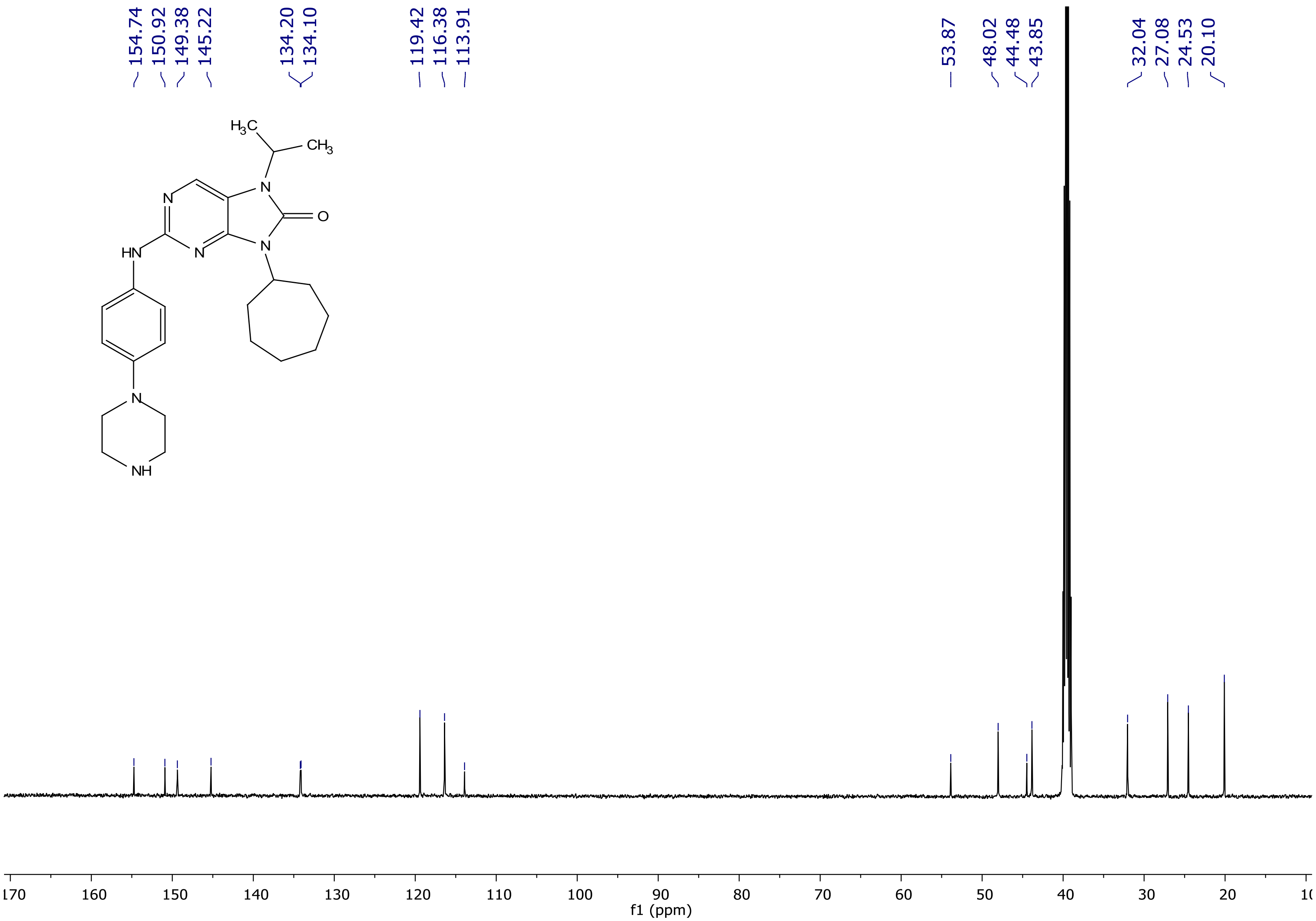

### HPLC of compound **12a**

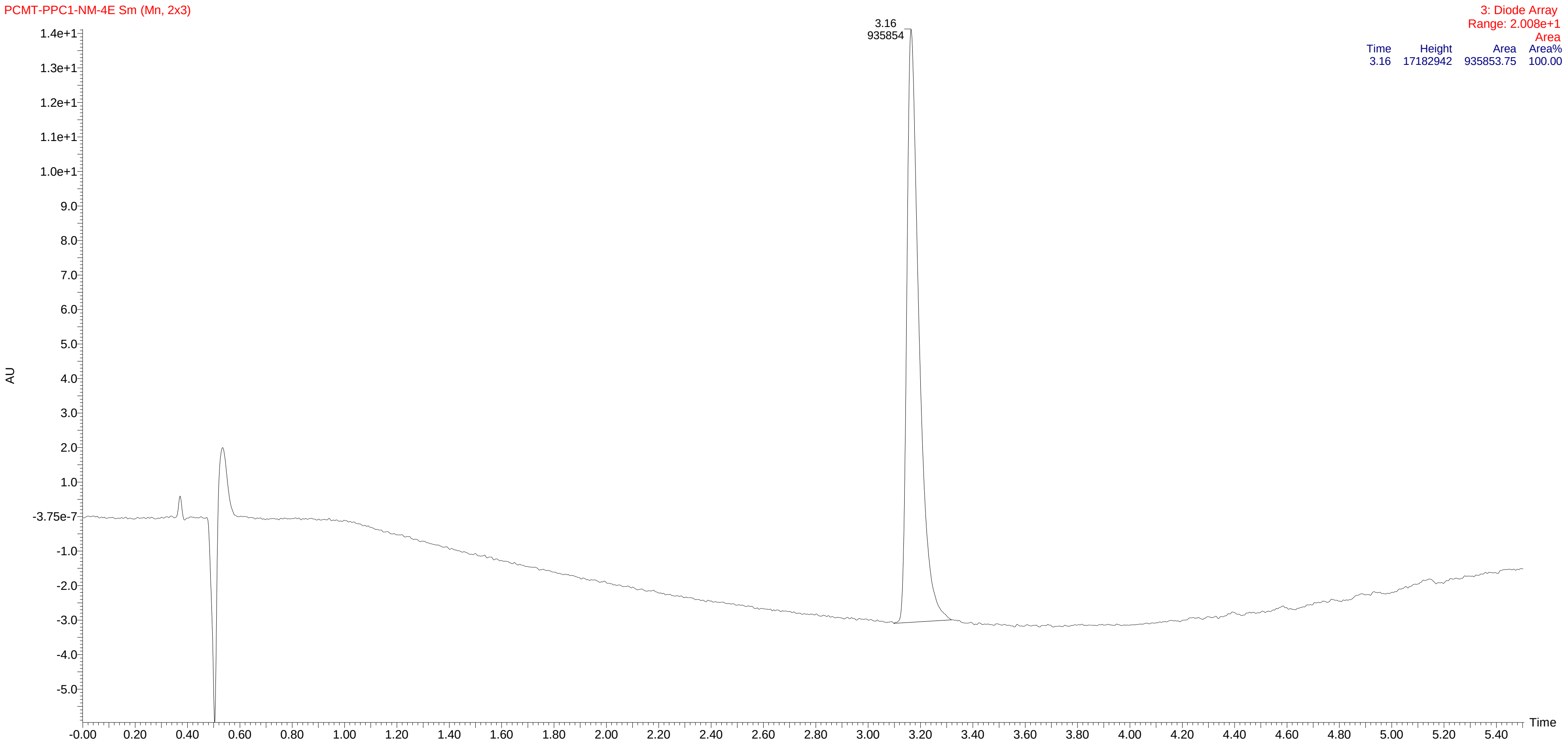

HPLC of compound **12b**

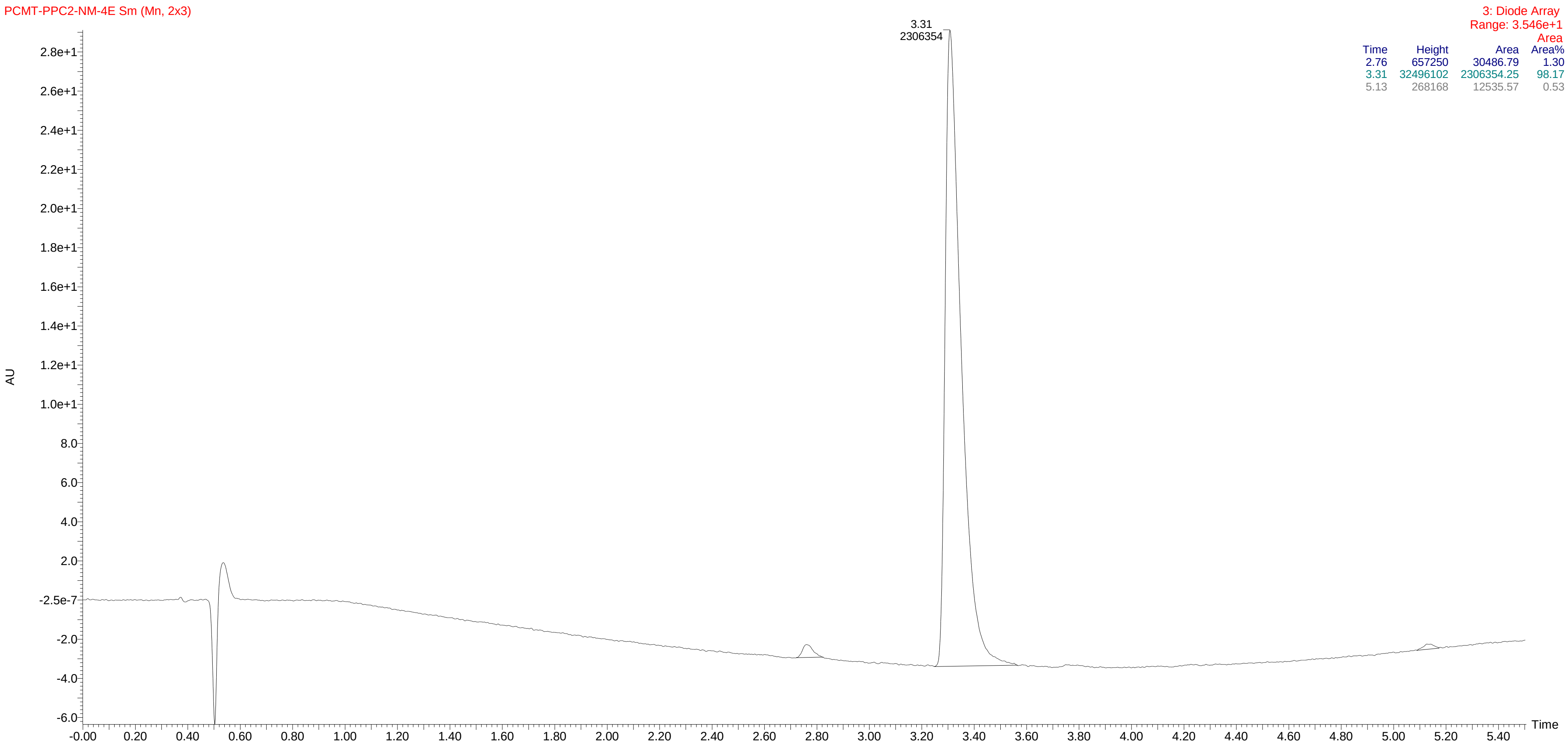

HPLC of compound **12c**

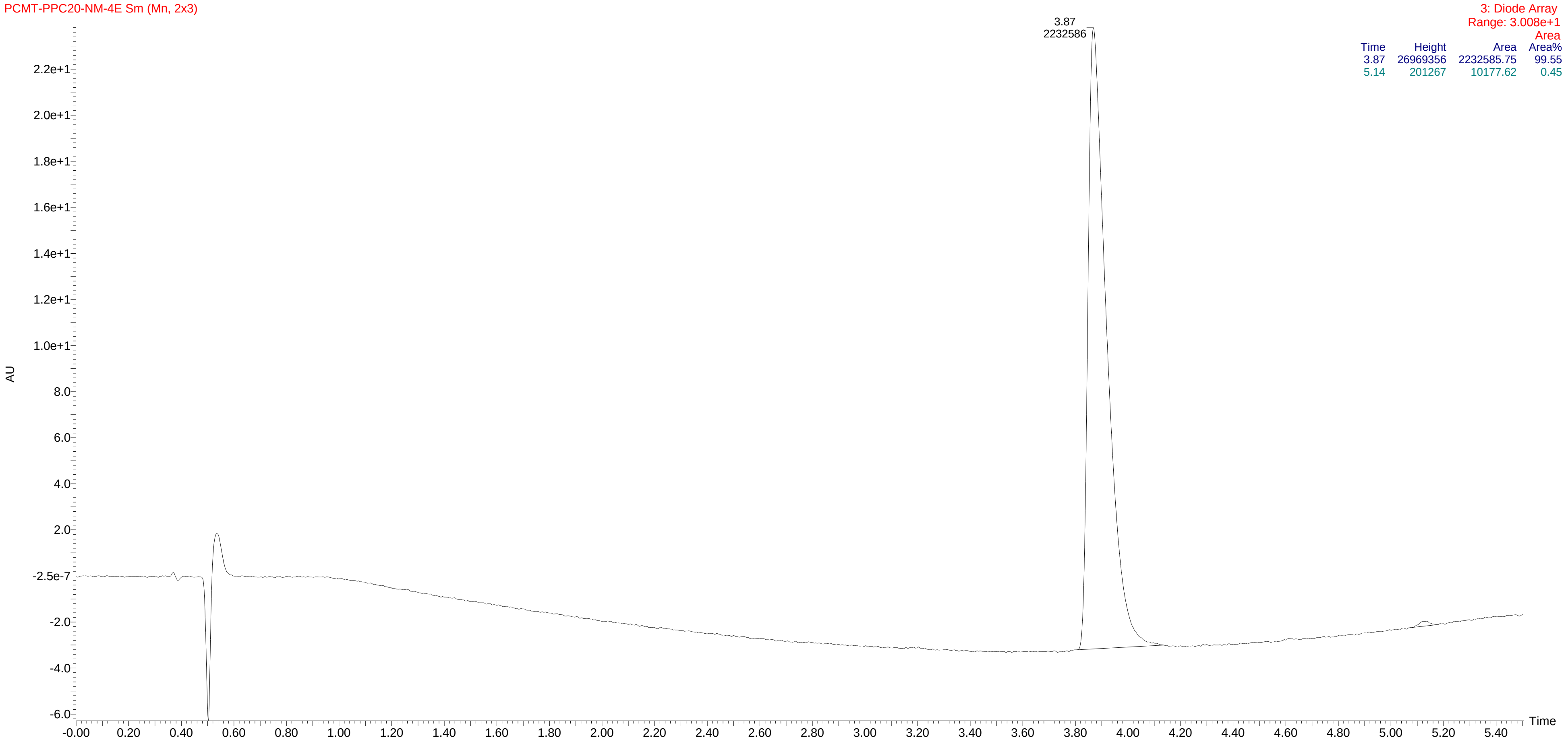


| Time | Height | Area | Area% |
|---|---|---|---|
| 3.87 | 26969356 | 2232585.75 | 99.55 |
| 5.14 | 201267 | 10177.62 | 0.45 |

HPLC of compound **12d**

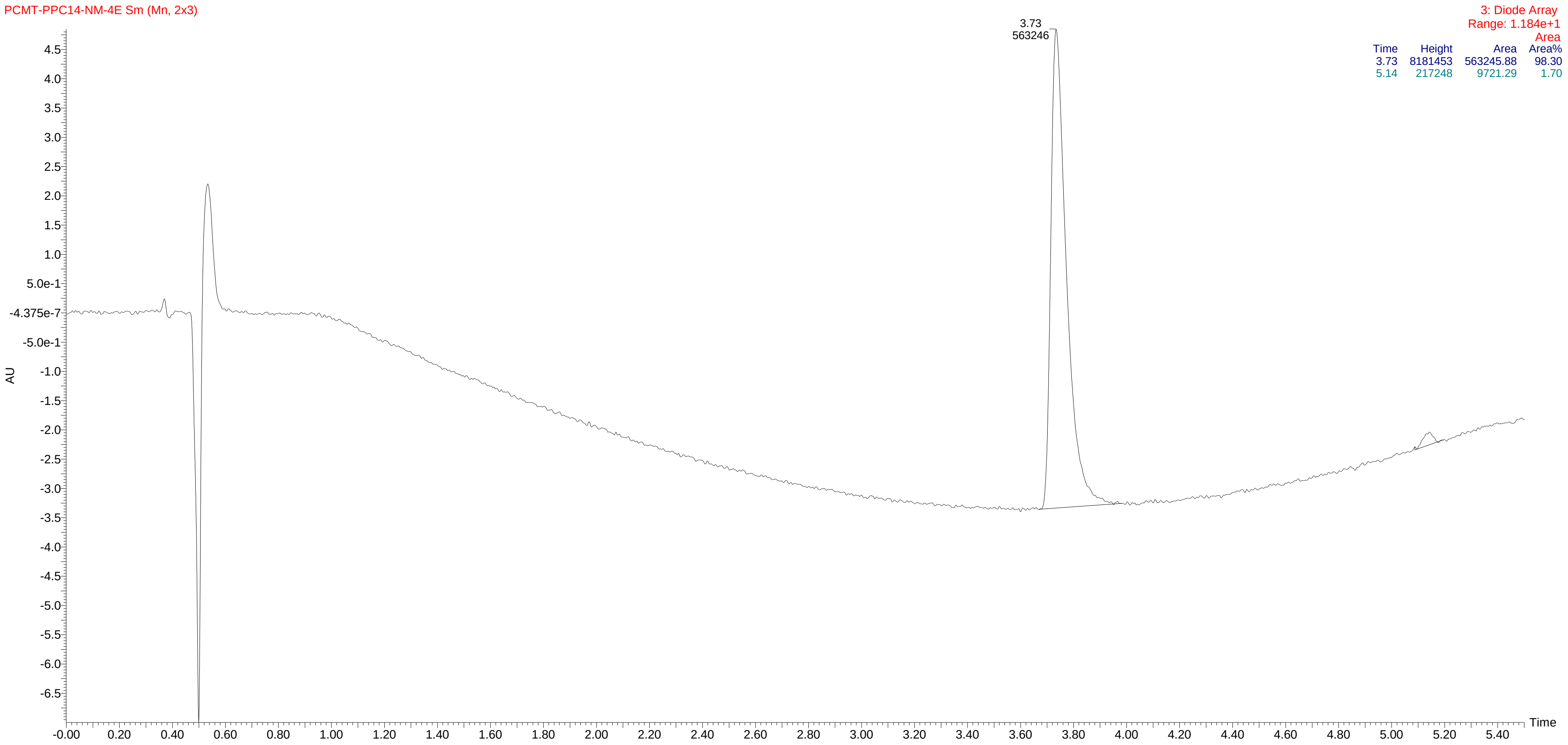

HPLC of compound **13a**

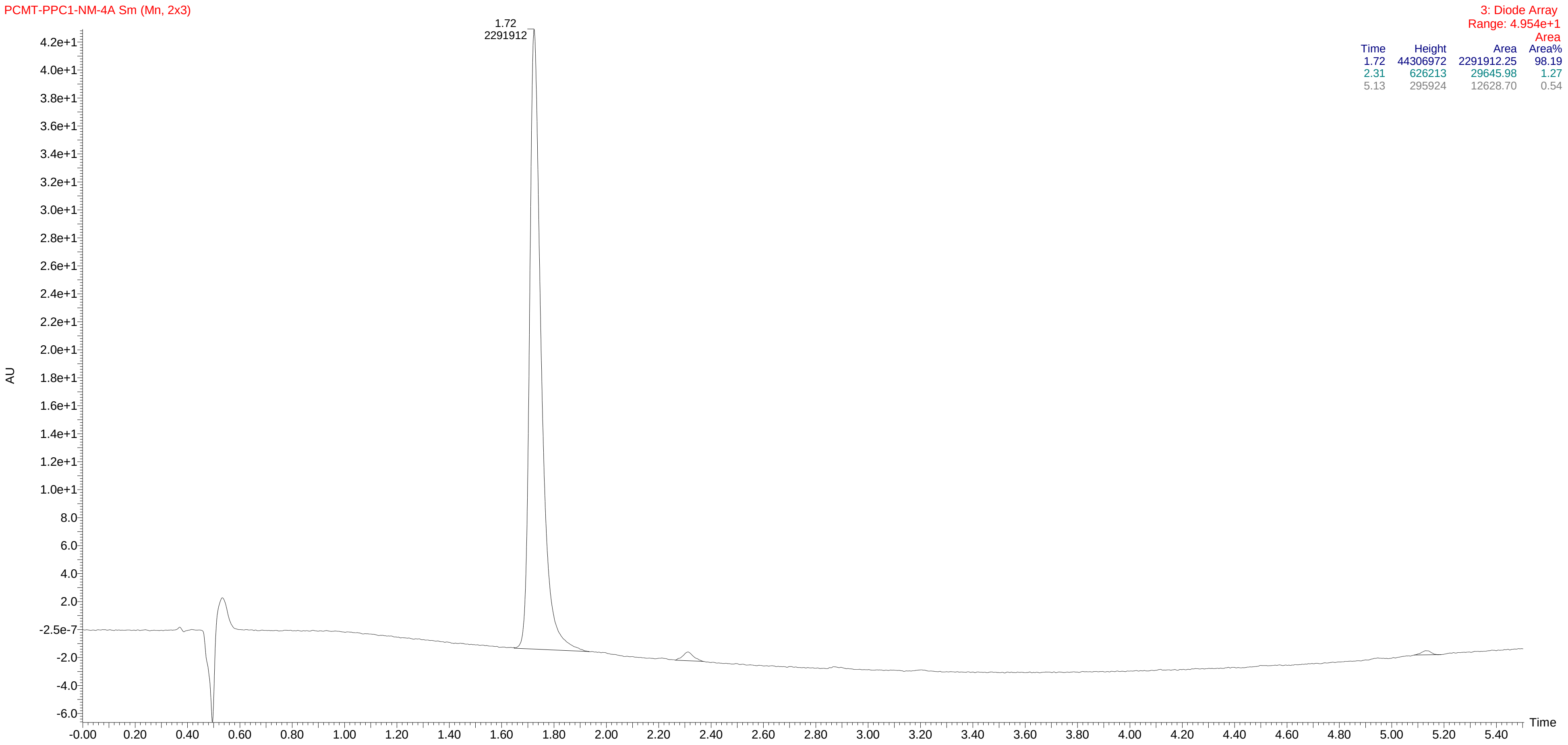

HPLC of compound **13b**

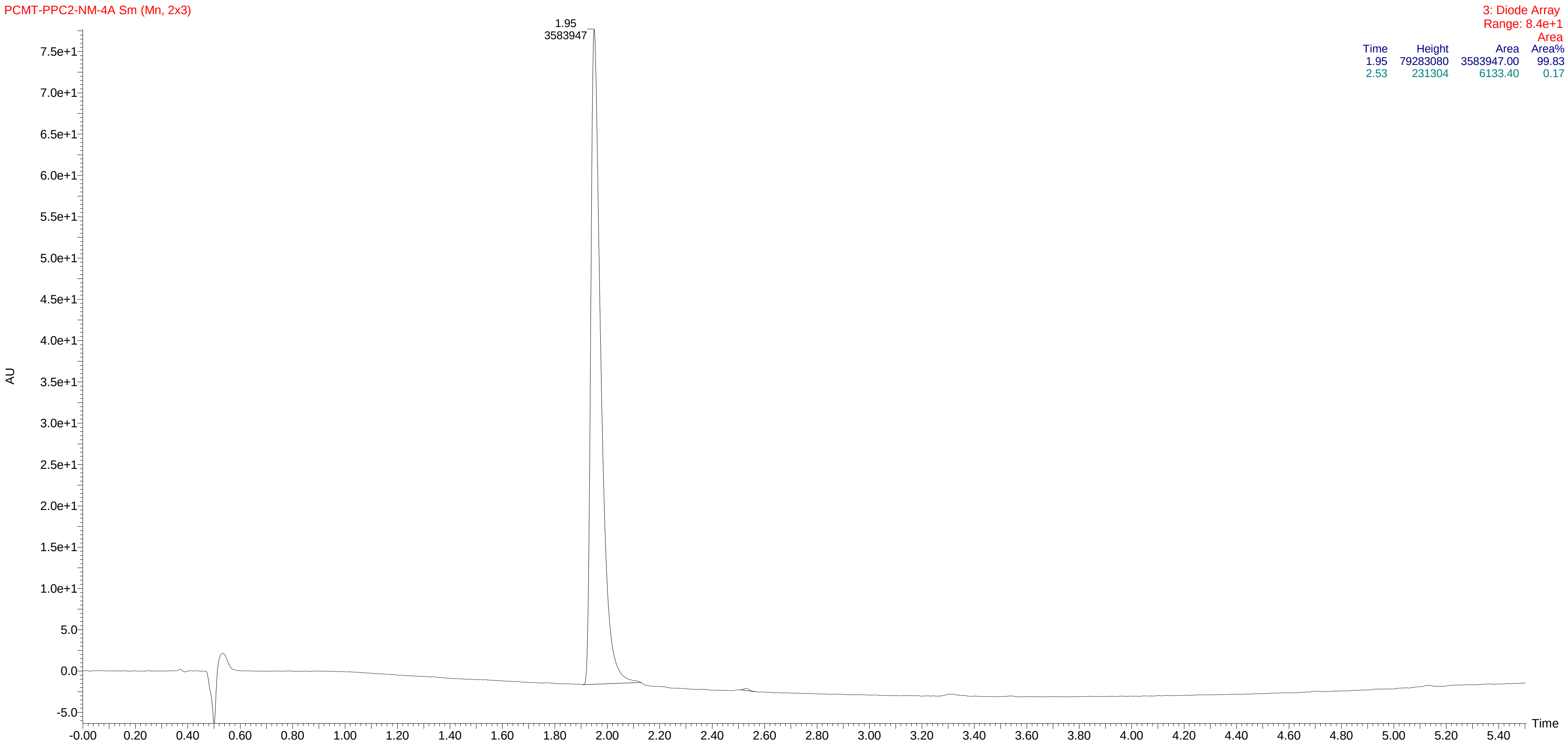


| Time | Height | Area | Area% |
|---|---|---|---|
| 1.95 | 79283080 | 3583947.00 | 99.83 |
| 2.53 | 231304 | 6133.40 | 0.17 |

HPLC of compound **13c**

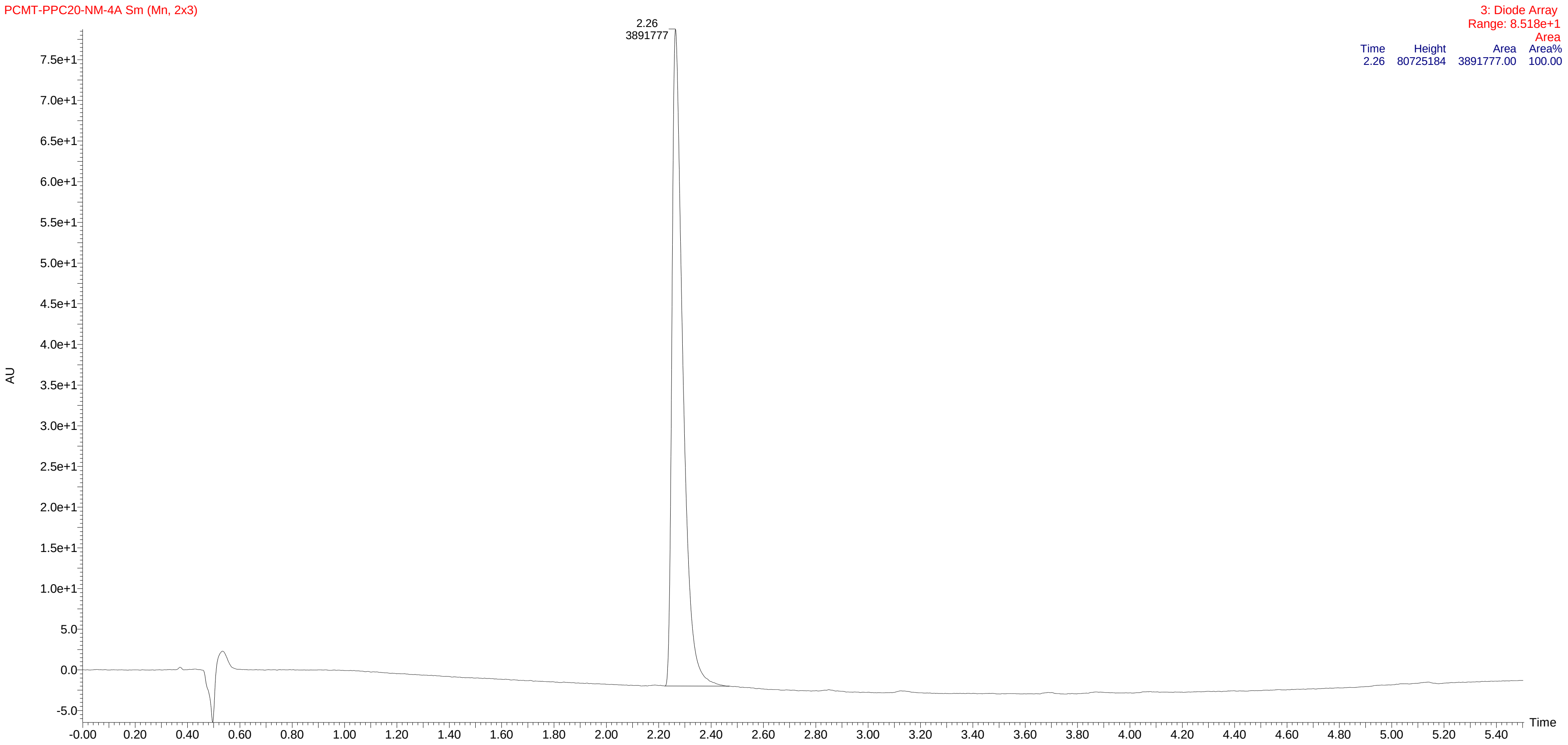

HPLC of compound **13d**

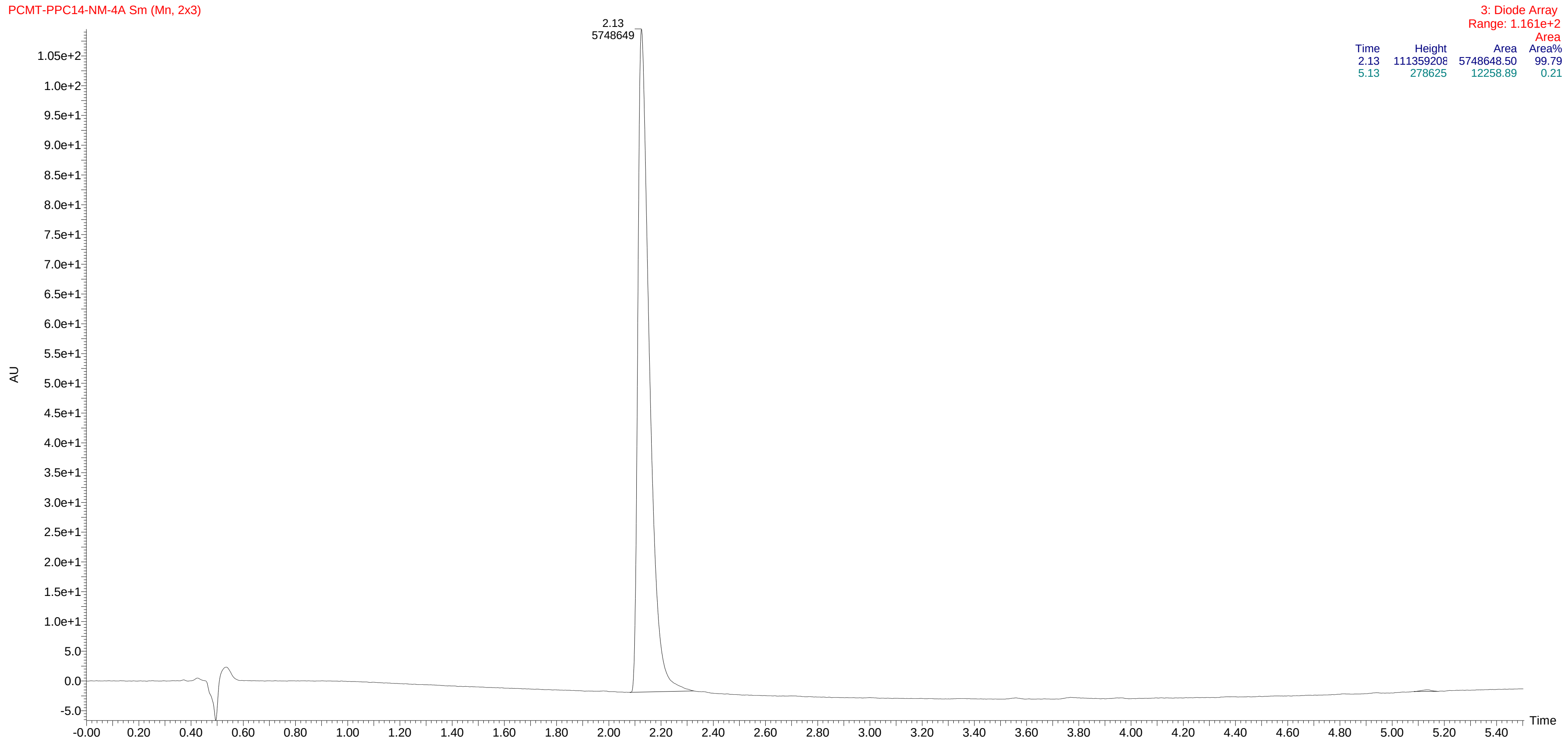

HPLC of compound **14a**

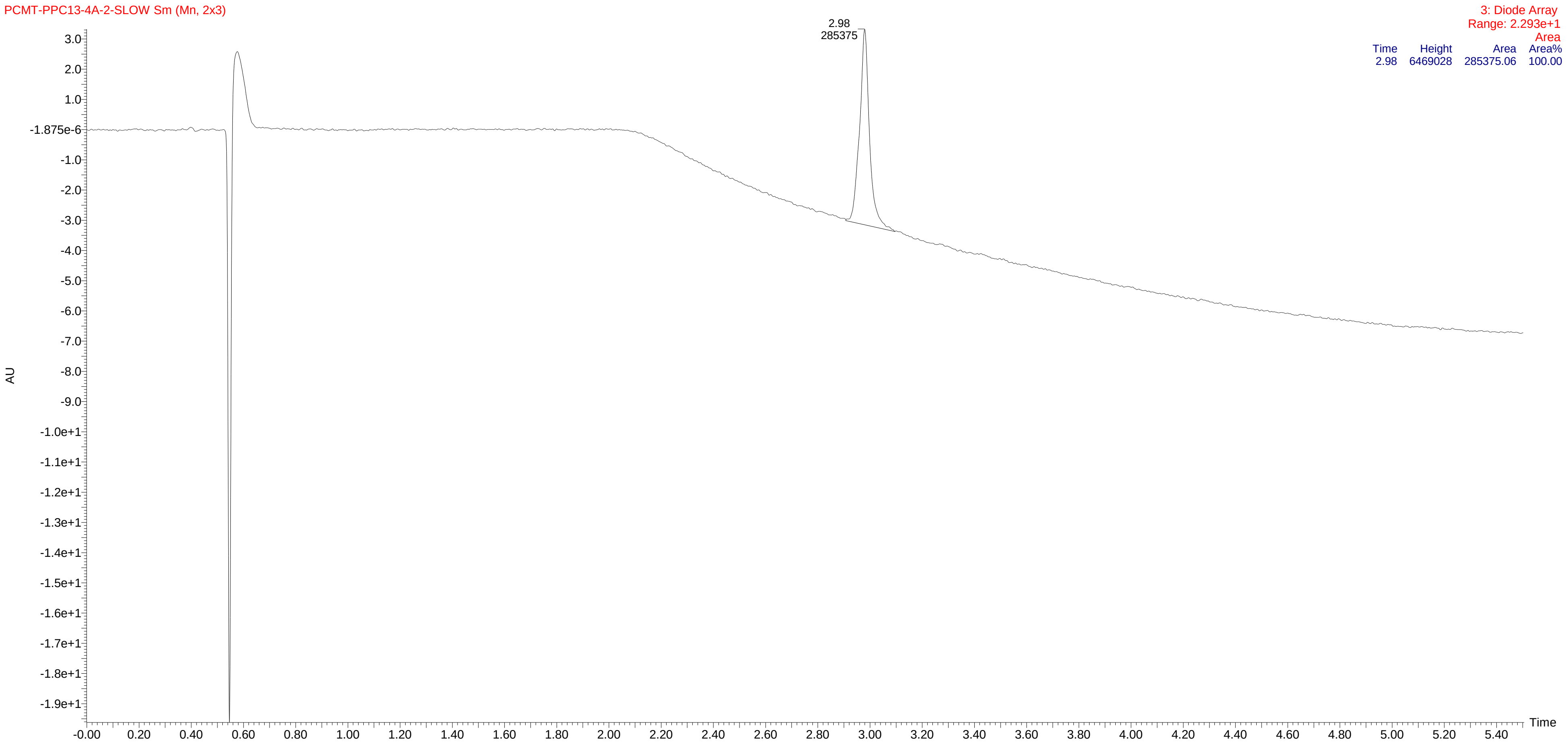

HPLC of compound **14b**

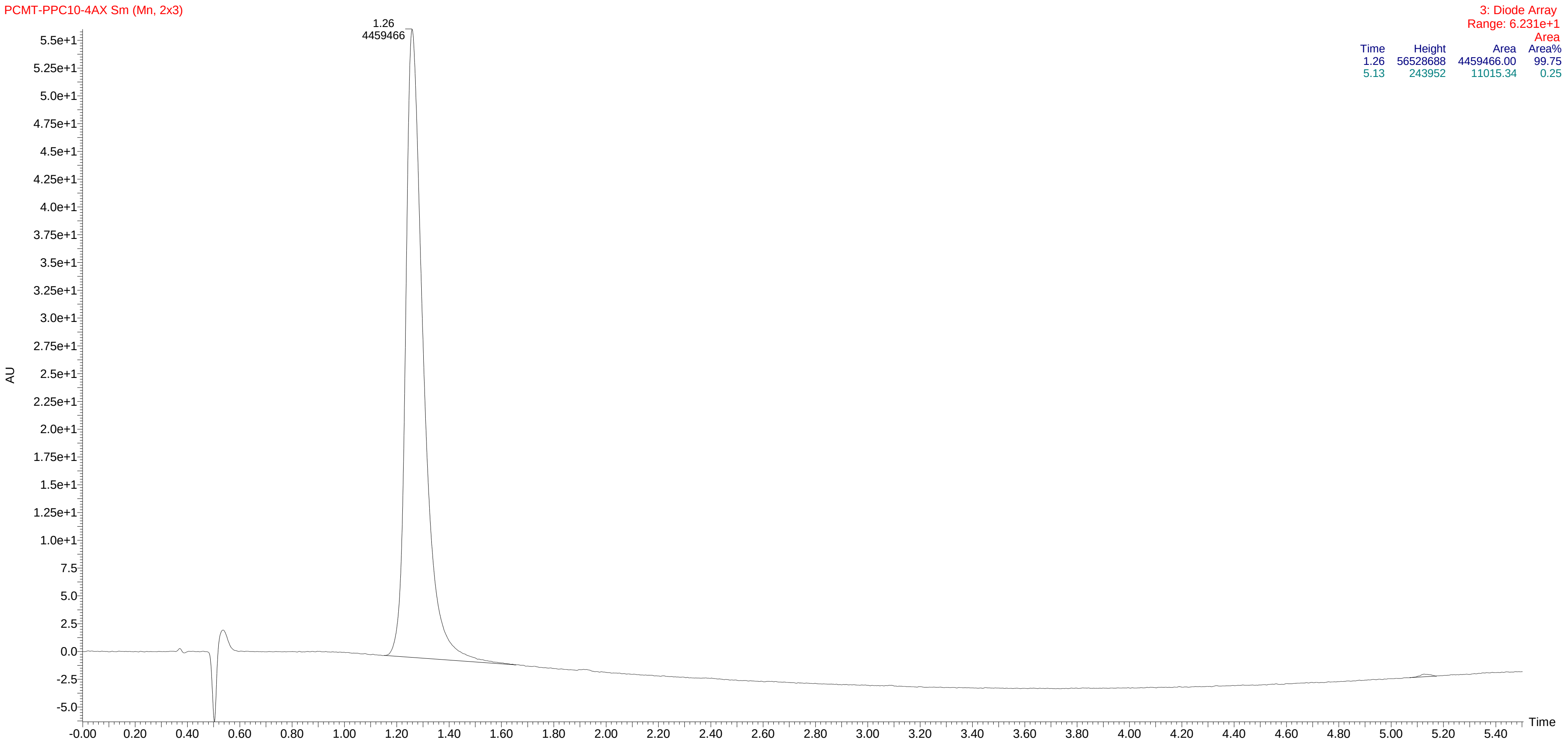

HPLC of compound **14c**

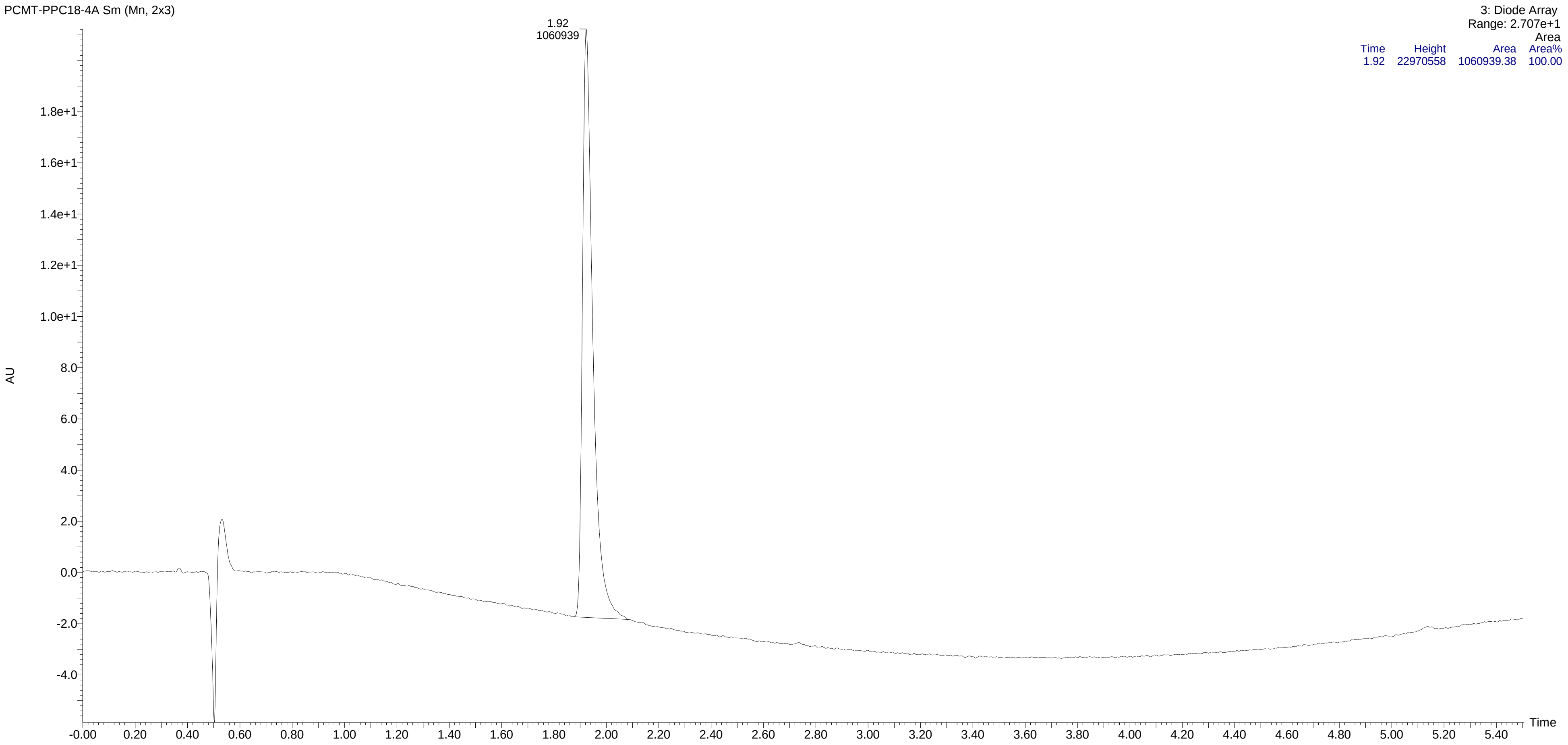


| Time | Height | Area | Area% |
|---|---|---|---|
| 1.92 | 22970558 | 1060939.38 | 100.00 |

HPLC of compound **14d**

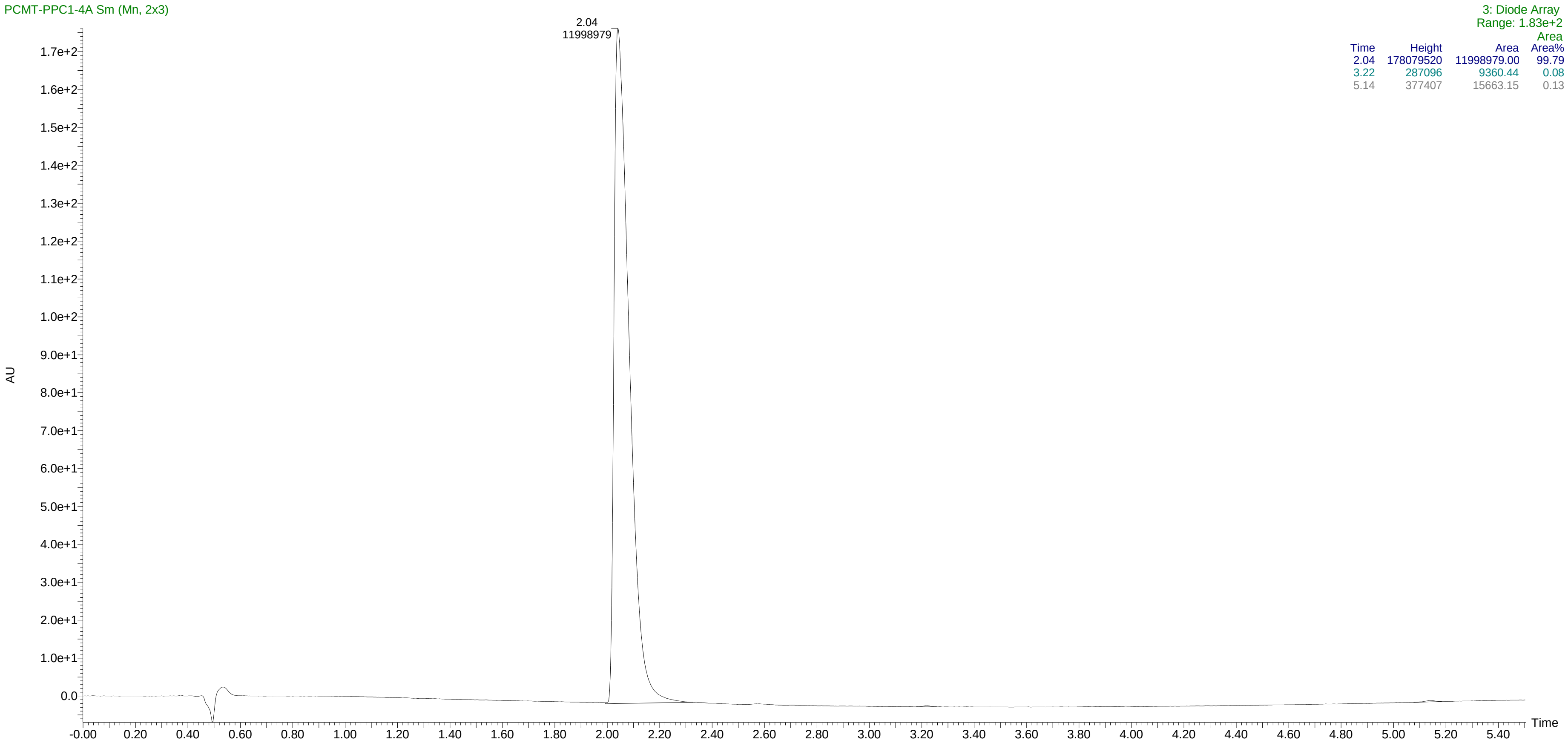


| Time | Height | Area | Area% |
|---|---|---|---|
| 2.04 | 178079520 | 11998979.00 | 99.79 |
| 3.22 | 287096 | 9360.44 | 0.08 |
| 5.14 | 377407 | 15663.15 | 0.13 |

HPLC of compound **14e**

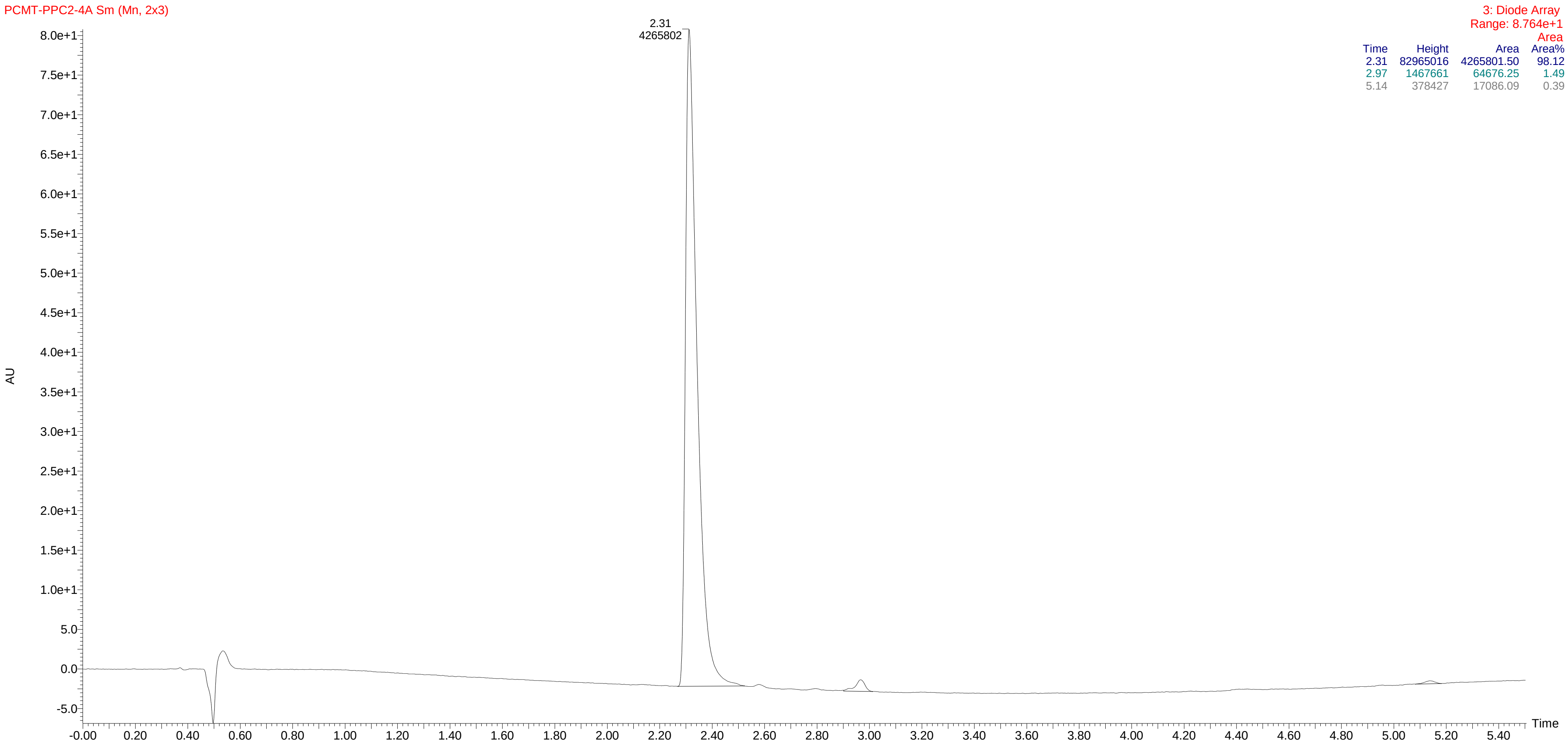


| Time | Height | Area | Area% |
|---|---|---|---|
| 2.31 | 82965016 | 4265801.50 | 98.12 |
| 2.97 | 1467661 | 64676.25 | 1.49 |
| 5.14 | 378427 | 17086.09 | 0.39 |

HPLC of compound **14f**

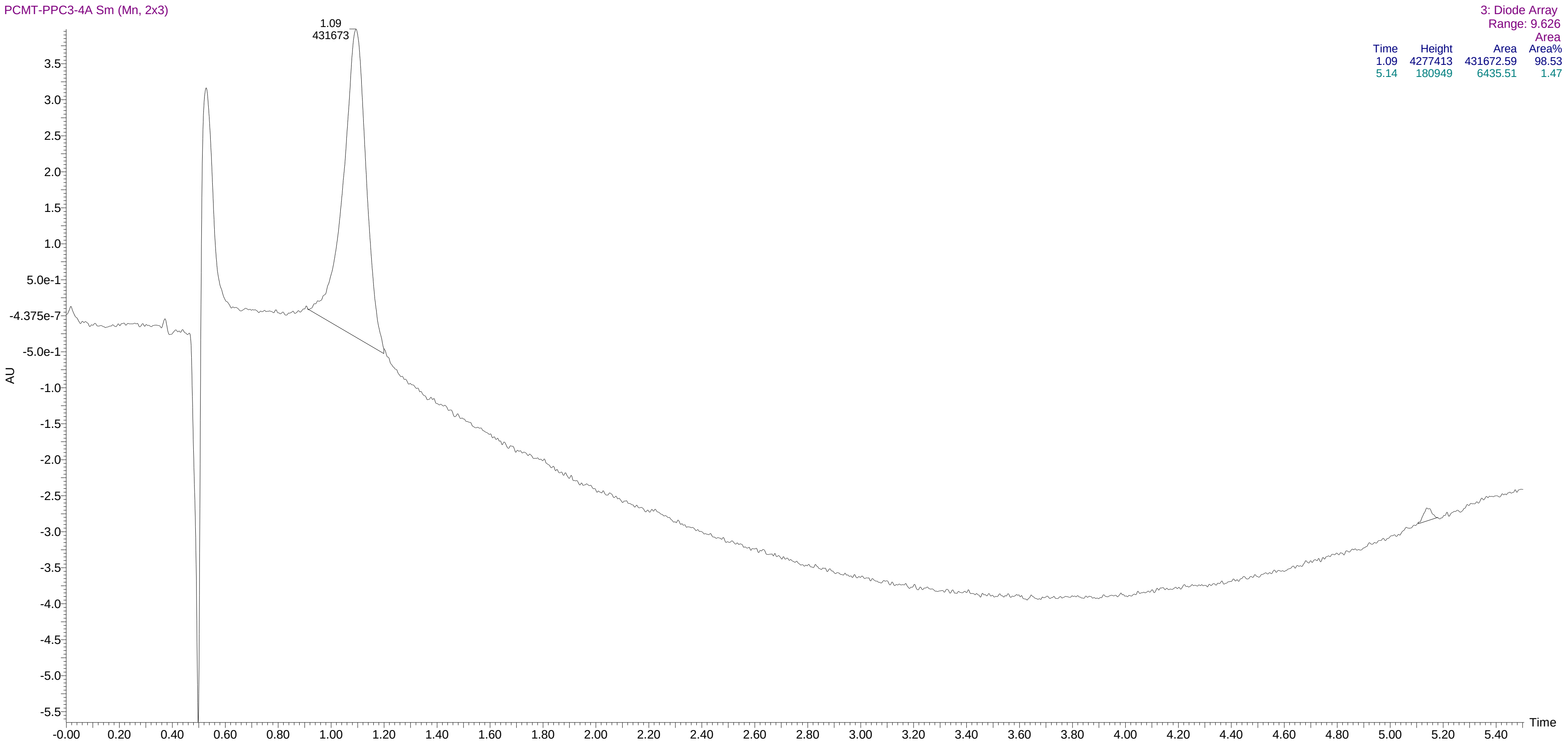


| Time | Height | Area | Area% |
|---|---|---|---|
| 1.09 | 4277413 | 431672.59 | 98.53 |
| 5.14 | 180949 | 6435.51 | 1.47 |

### HPLC of compound **14f**

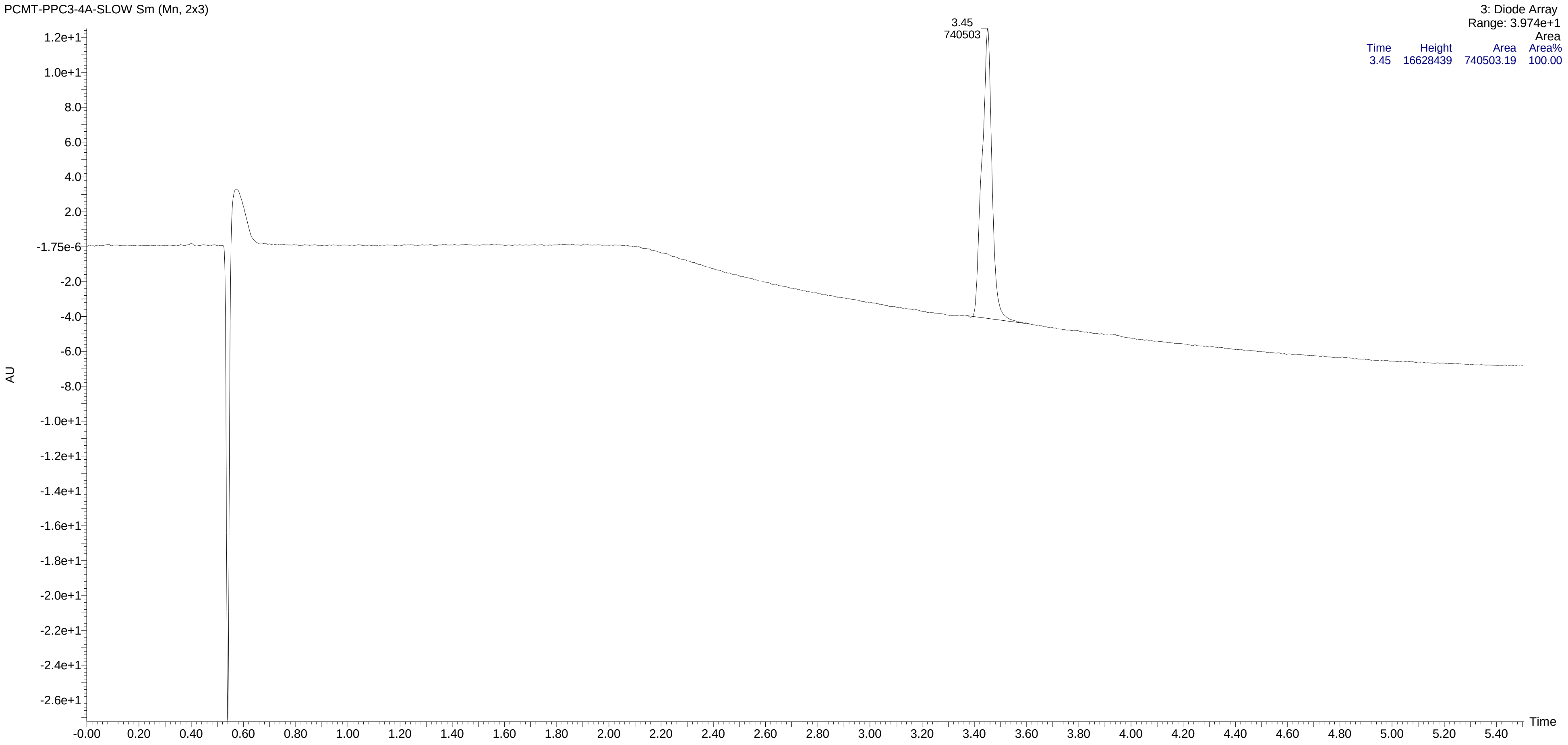

HPLC of compound **14g**

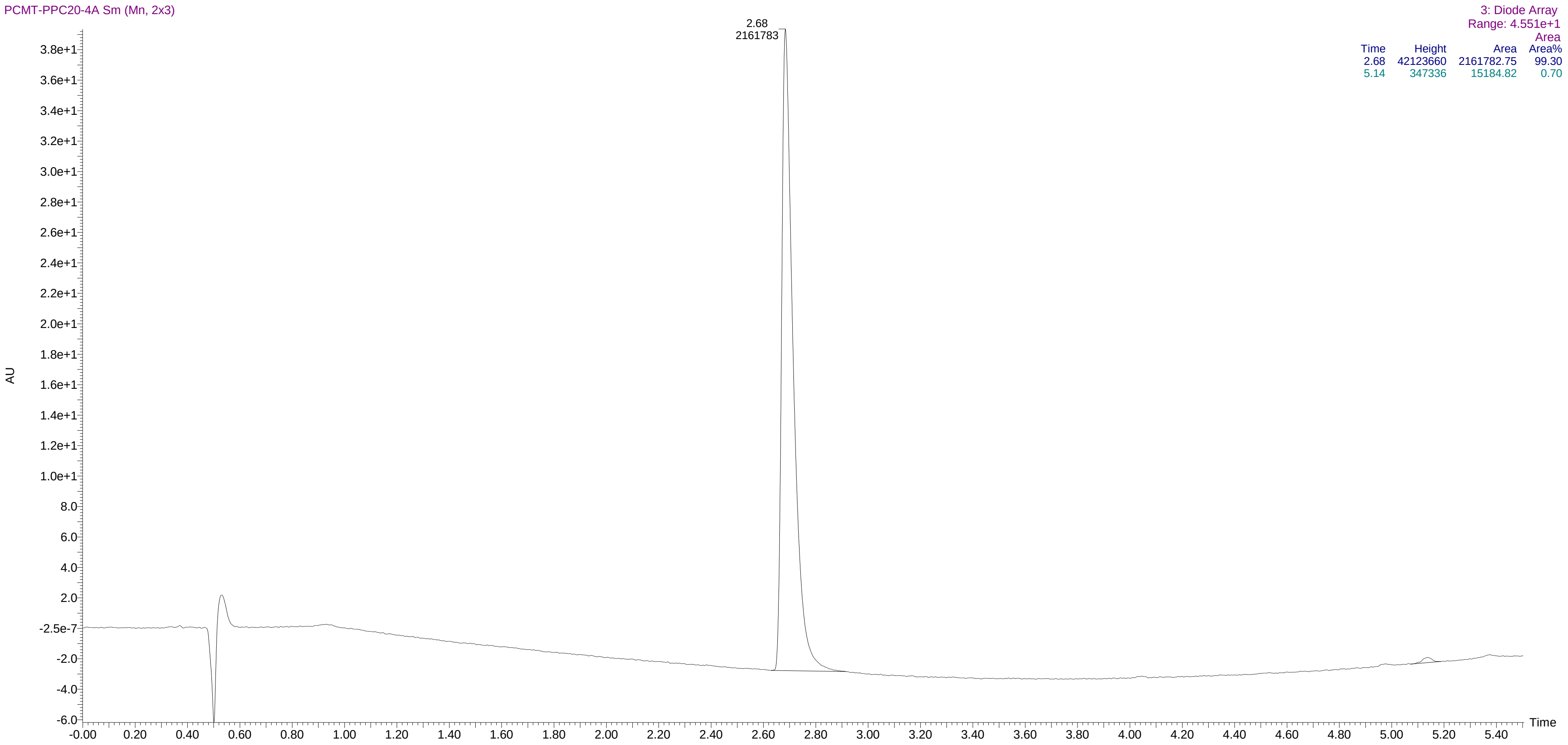


| Time | Height | Area | Area Area% |
|---|---|---|---|
| 2.68 | 42123660 | 2161782.75 | 99.30 |
| 5.14 | 347336 | 15184.82 | 0.70 |

HPLC of compound **14h**

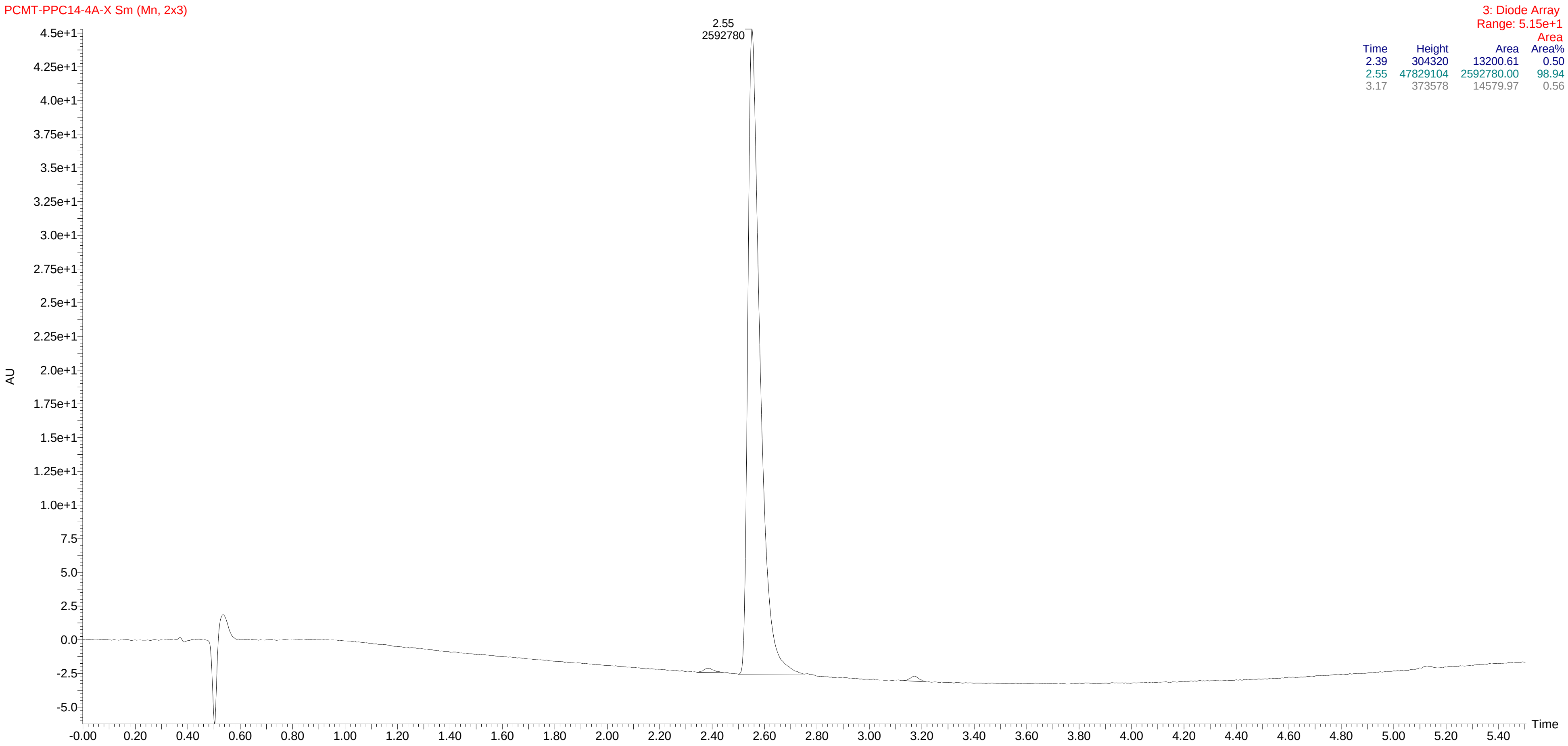


| Time | Height | Area | Area% |
|---|---|---|---|
| 2.39 | 304320 | 13200.61 | 0.50 |
| 2.55 | 47829104 | 2592780.00 | 98.94 |
| 3.17 | 373578 | 14579.97 | 0.56 |

HPLC of compound **14i**

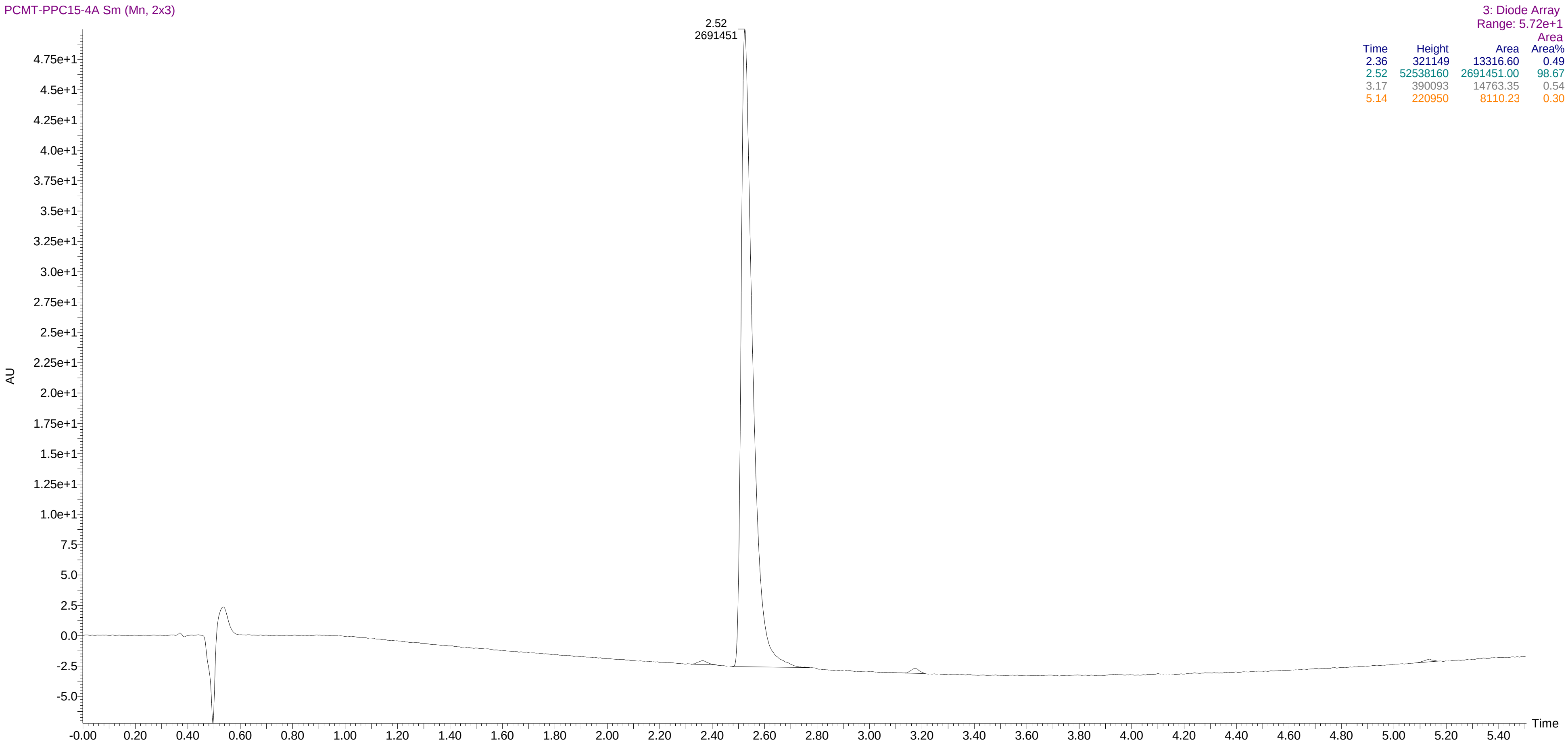


| Time | Height | Area | Area% |
|---|---|---|---|
| 2.36 | 321149 | 13316.60 | 0.49 |
| 2.52 | 52538160 | 2691451.00 | 98.67 |
| 3.17 | 390093 | 14763.35 | 0.54 |
| 5.14 | 220950 | 8110.23 | 0.30 |

HPLC of compound **14j**

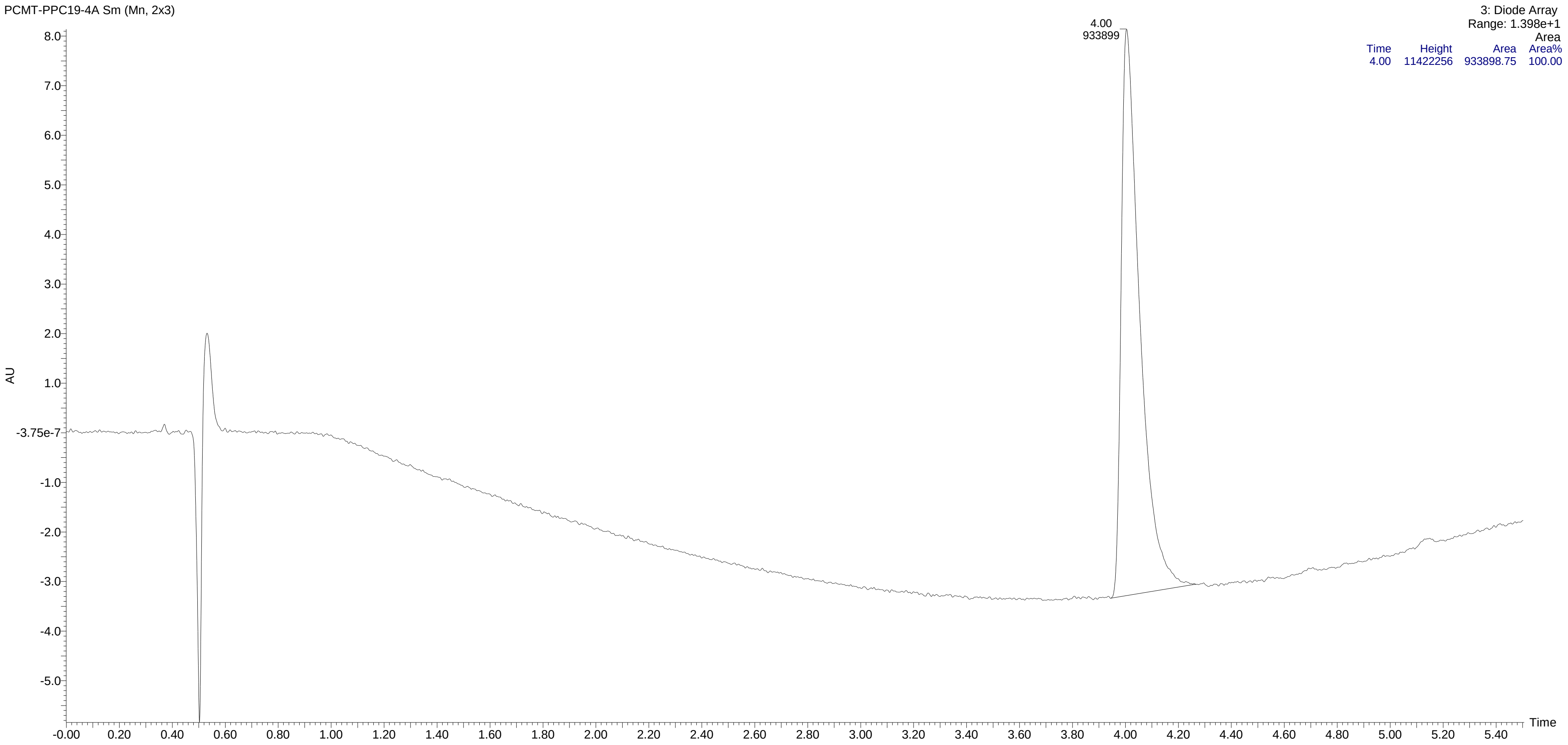

HPLC of compound **14k**

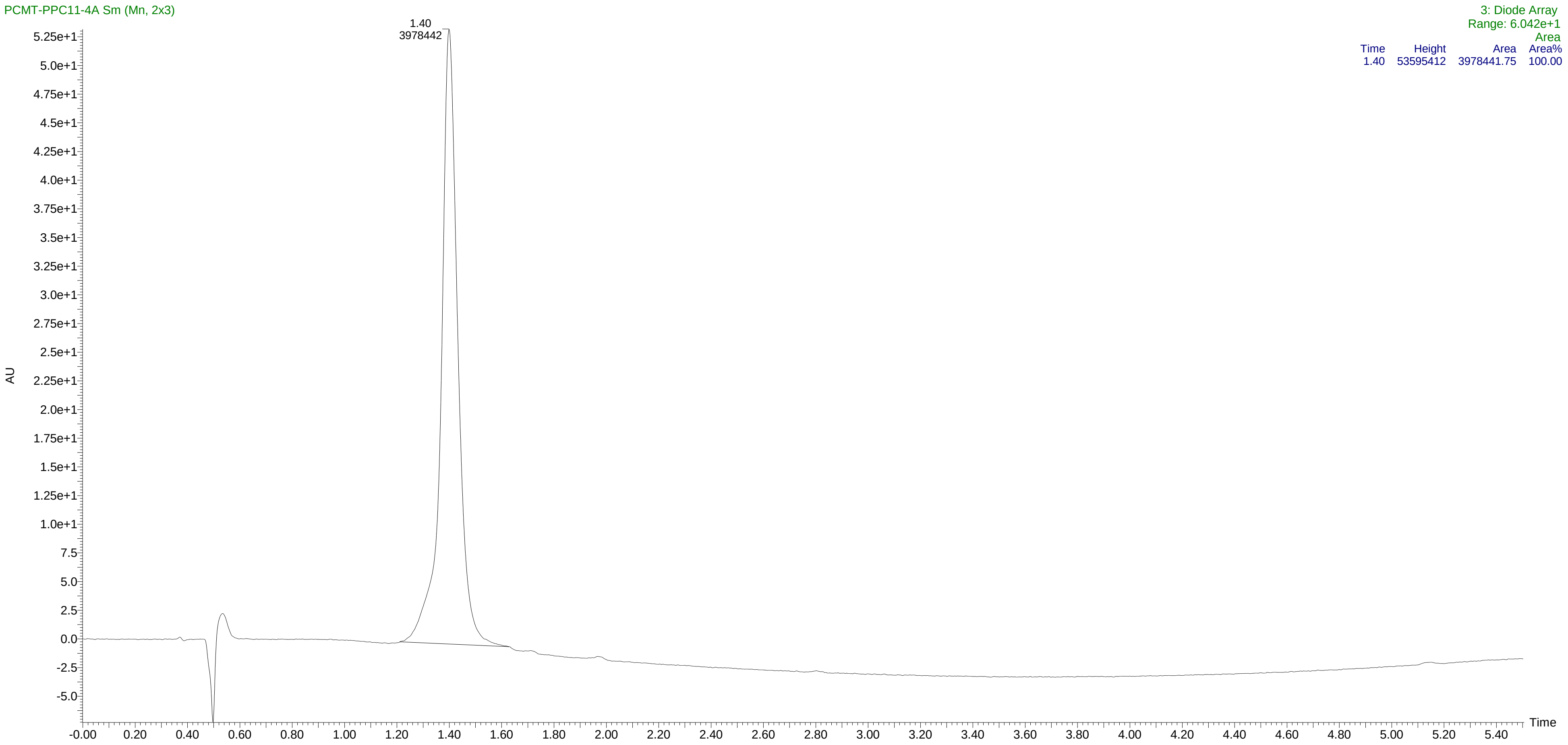

HPLC of compound **14l**

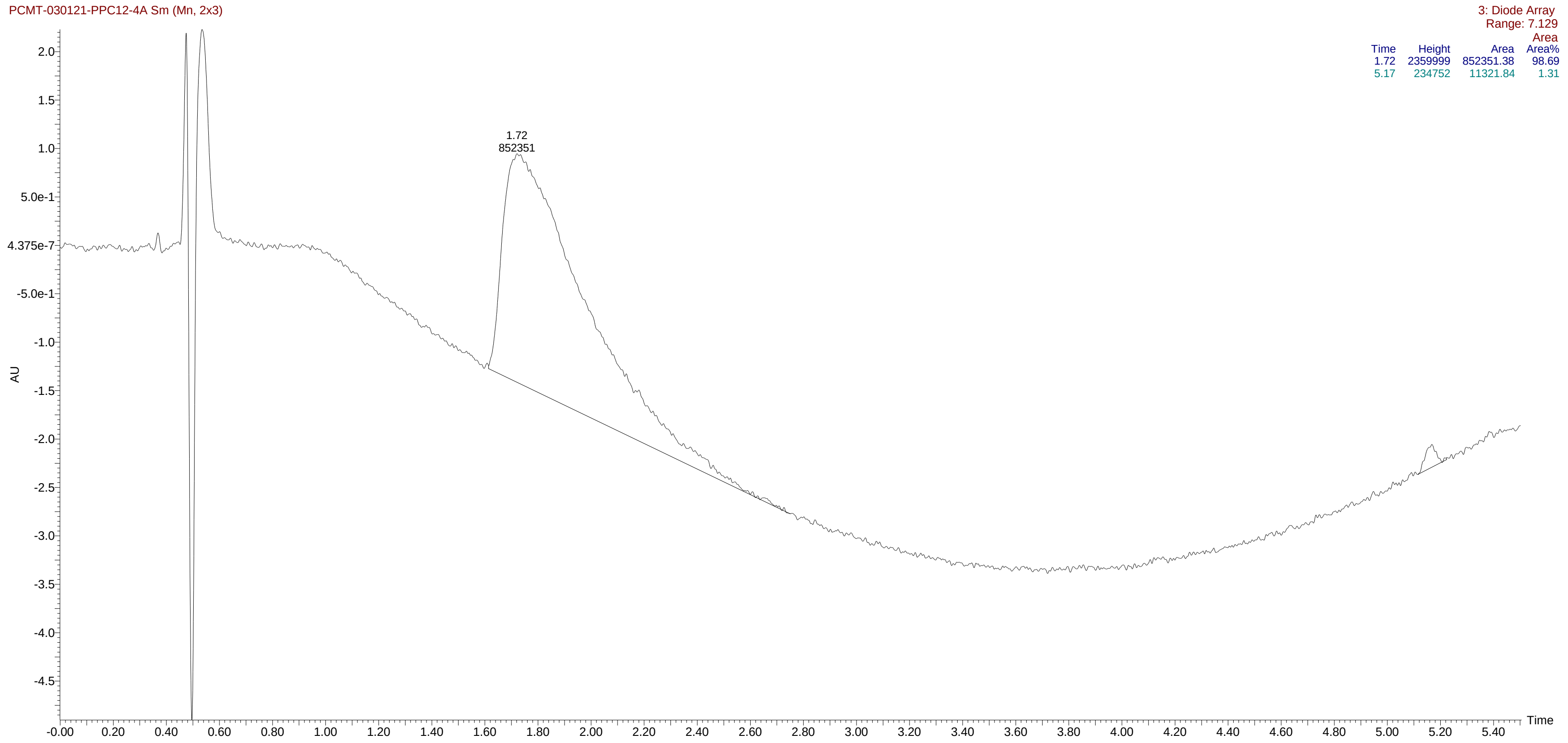

HPLC of compound **14m**

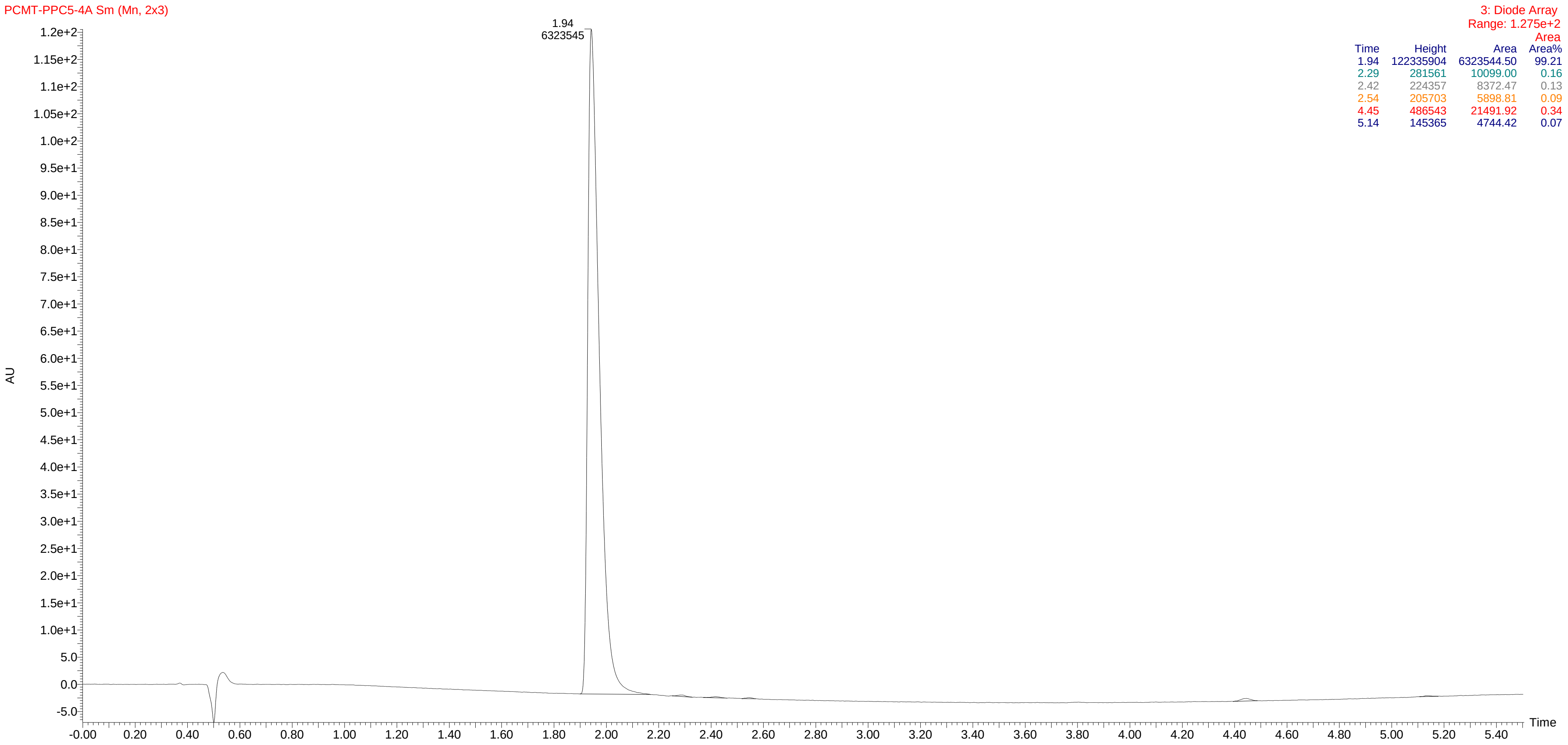


| Time | Height | Area | Area% |
|---|---|---|---|
| 1.94 | 122335904 | 6323544.50 | 99.21 |
| 2.29 | 281561 | 10099.00 | 0.16 |
| 2.42 | 224357 | 8372.47 | 0.13 |
| 2.54 | 205703 | 5898.81 | 0.09 |
| 4.45 | 486543 | 21491.92 | 0.34 |
| 5.14 | 145365 | 4744.42 | 0.07 |

HPLC of compound **14n**

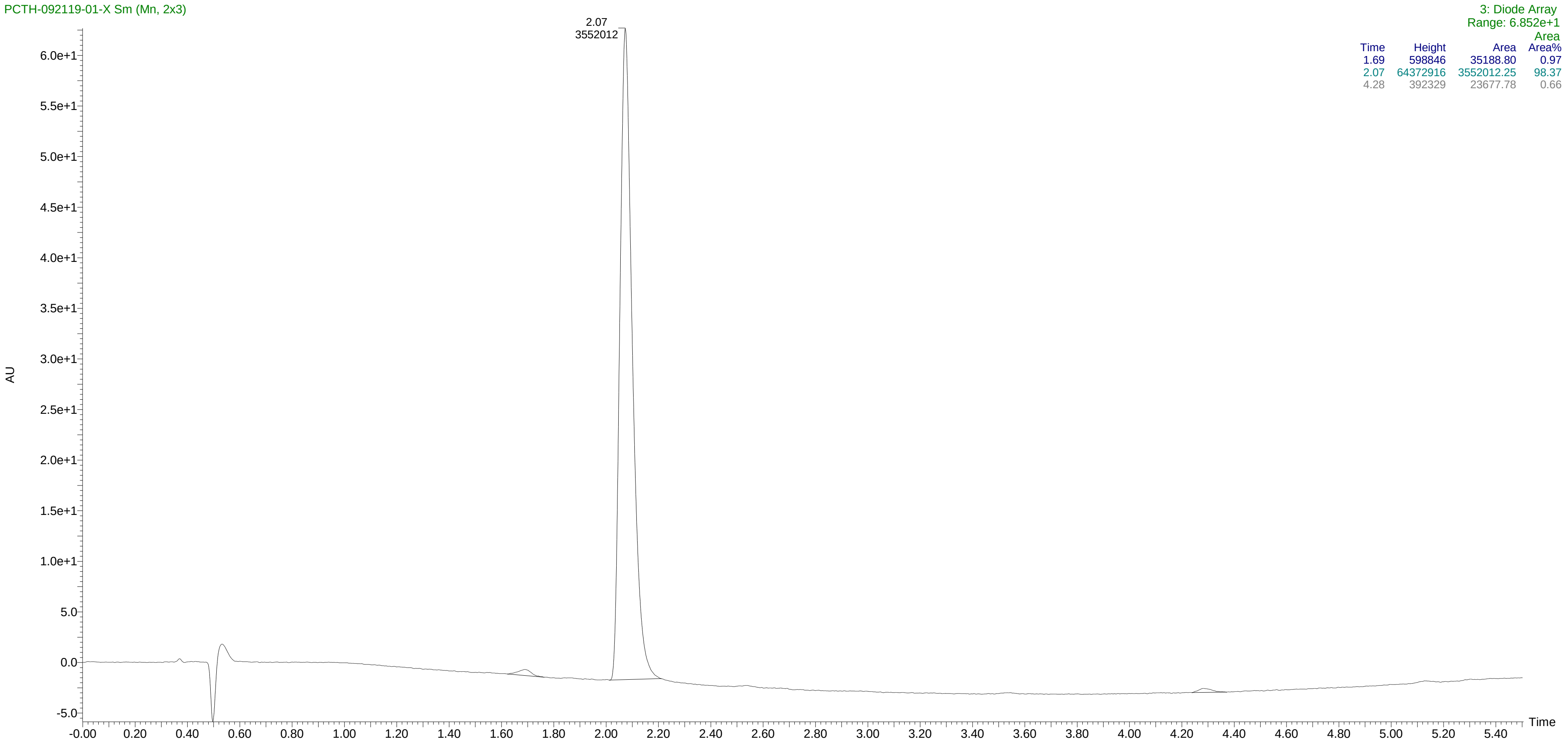

HPLC of compound **14o**

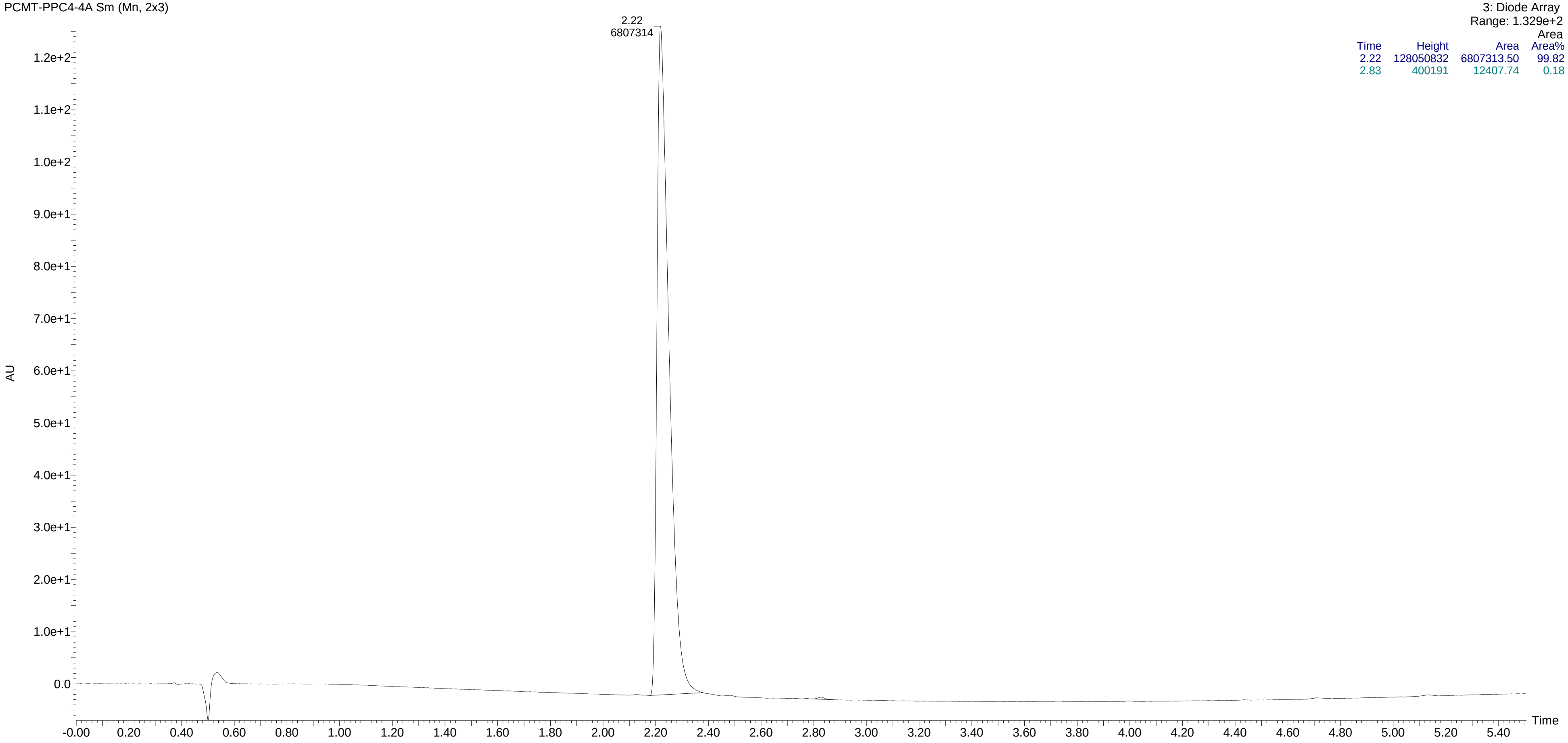


| Time | Height | Area | Area% |
|---|---|---|---|
| 2.22 | 128050832 | 6807313.50 | 99.82 |
| 2.83 | 400191 | 12407.74 | 0.18 |

HPLC of compound **14p**

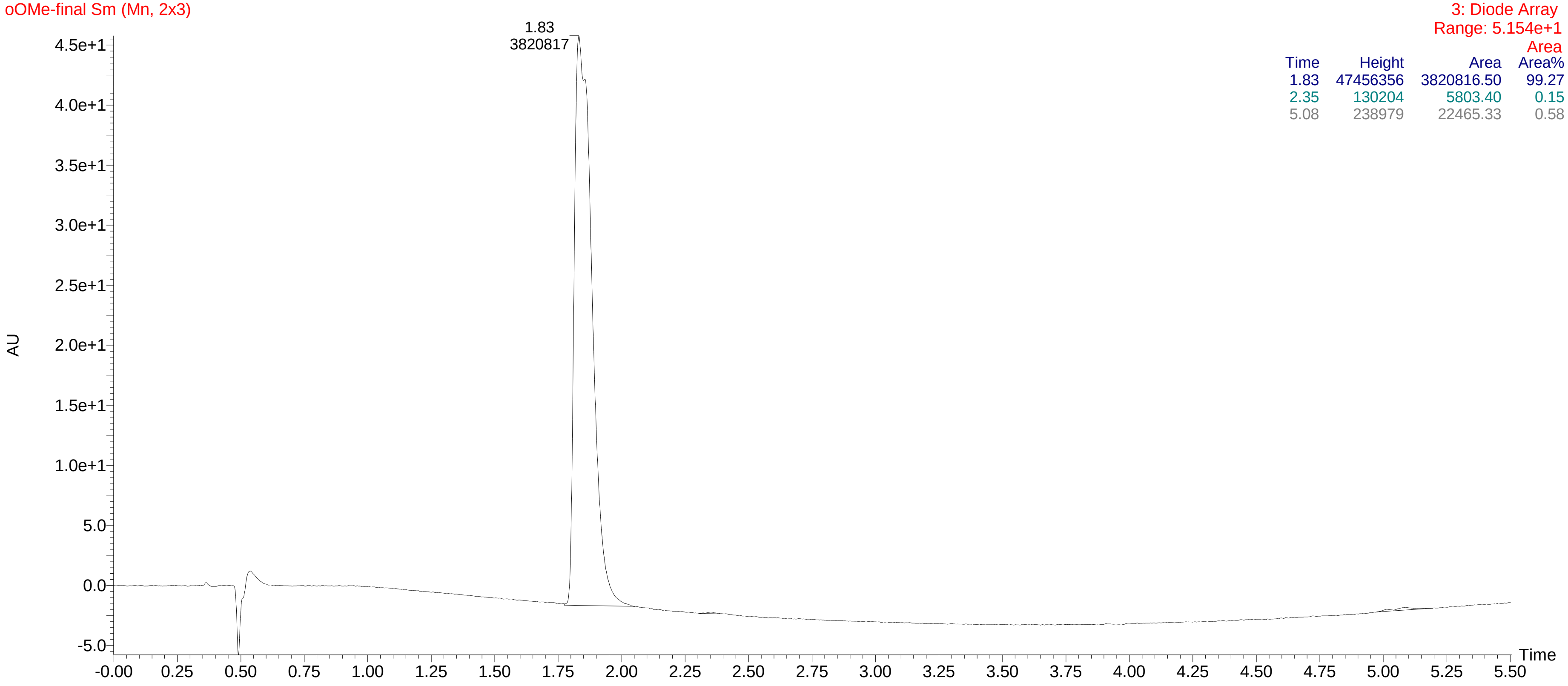


| Time | Height | Area | Area% |
|---|---|---|---|
| 1.83 | 47456356 | 3820816.50 | 99.27 |
| 2.35 | 130204 | 5803.40 | 0.15 |
| 5.08 | 238979 | 22465.33 | 0.58 |

HPLC of compound **14q**

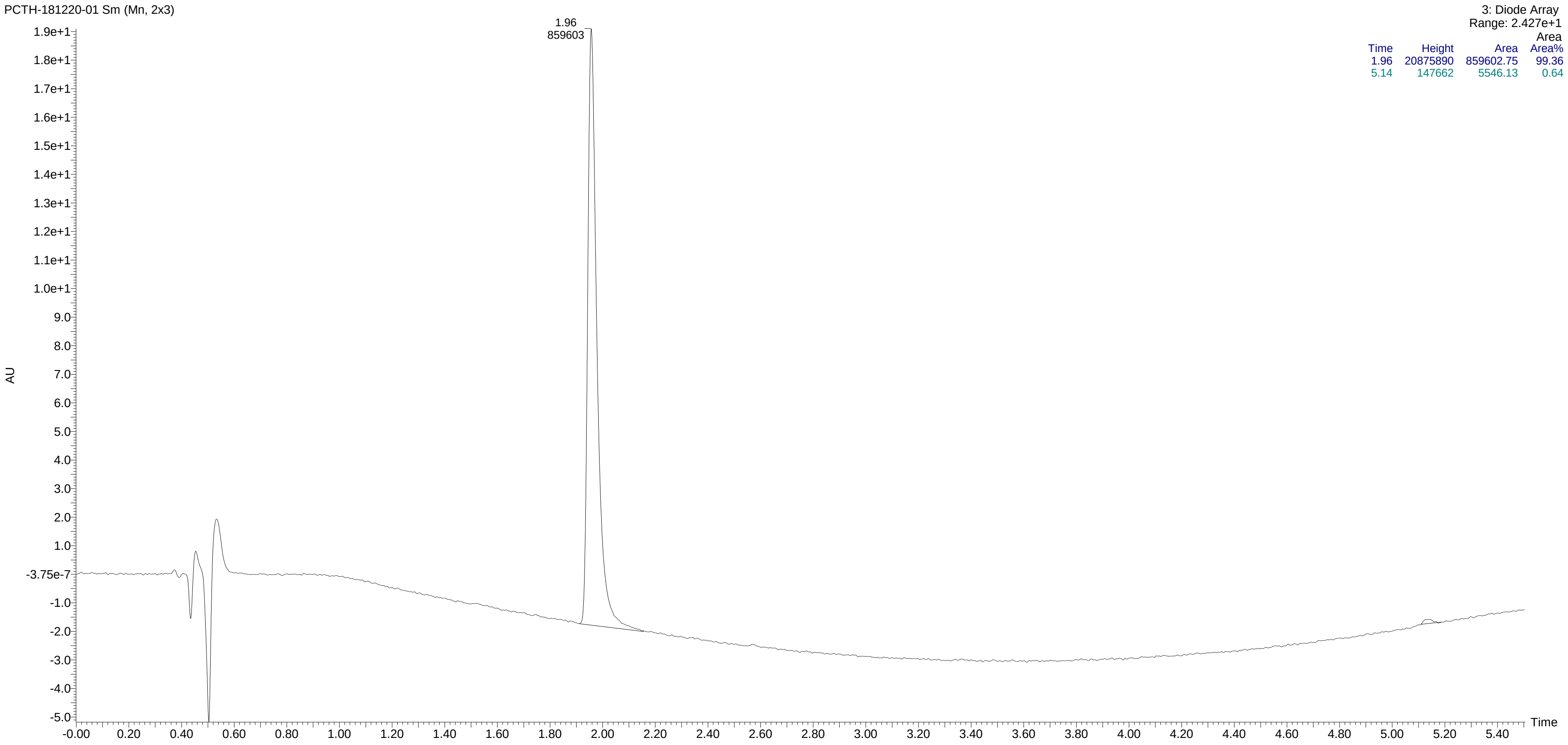


| Time | Height | Area | Area% |
|---|---|---|---|
| 1.96 | 20875890 | 859602.75 | 99.36 |
| 5.14 | 147662 | 5546.13 | 0.64 |

HPLC of compound **14r**

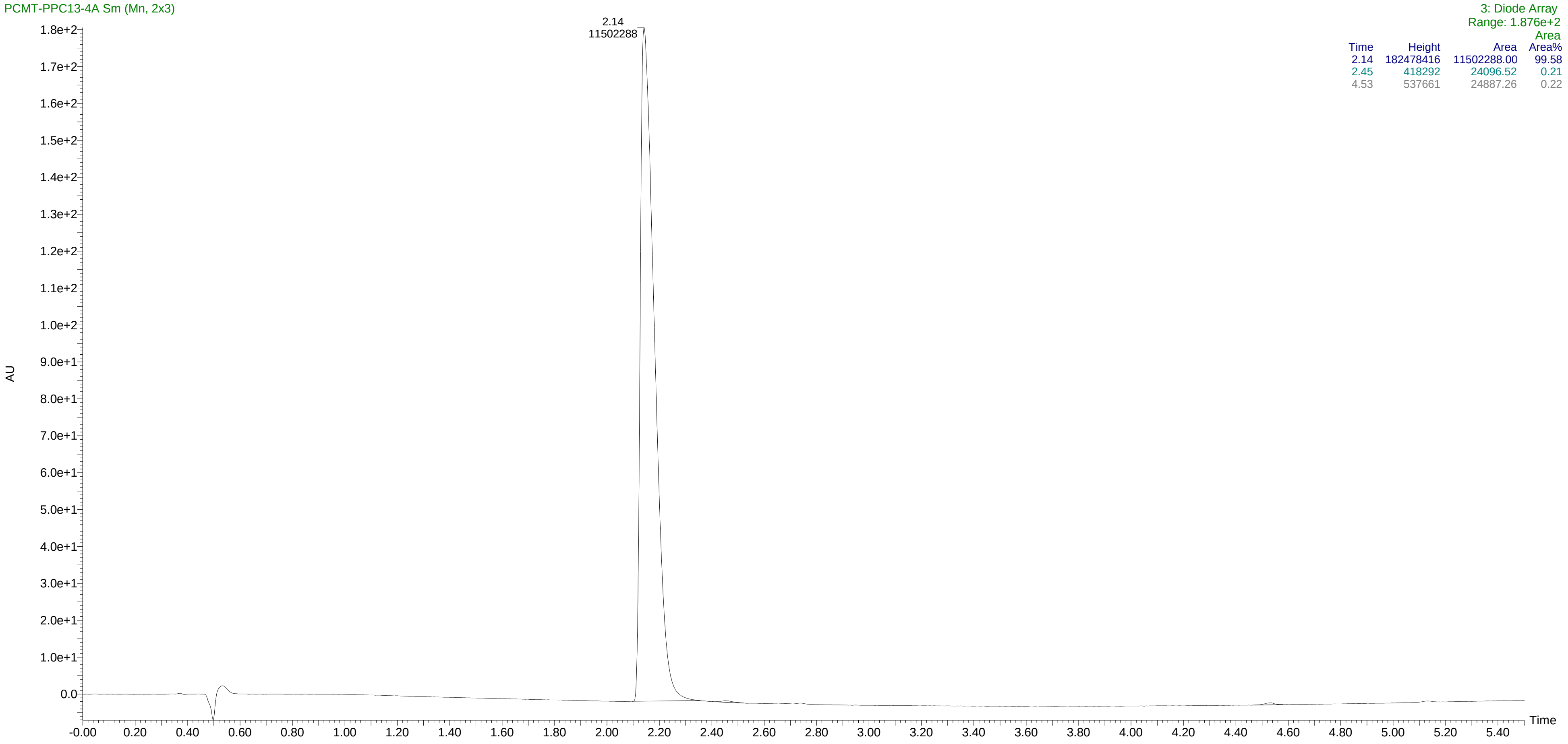


| Time | Height | Area | Area% |
|---|---|---|---|
| 2.14 | 182478416 | 11502288.00 | 99.58 |
| 2.45 | 418292 | 24096.52 | 0.21 |
| 4.53 | 537661 | 24887.26 | 0.22 |

HPLC of compound **14s**

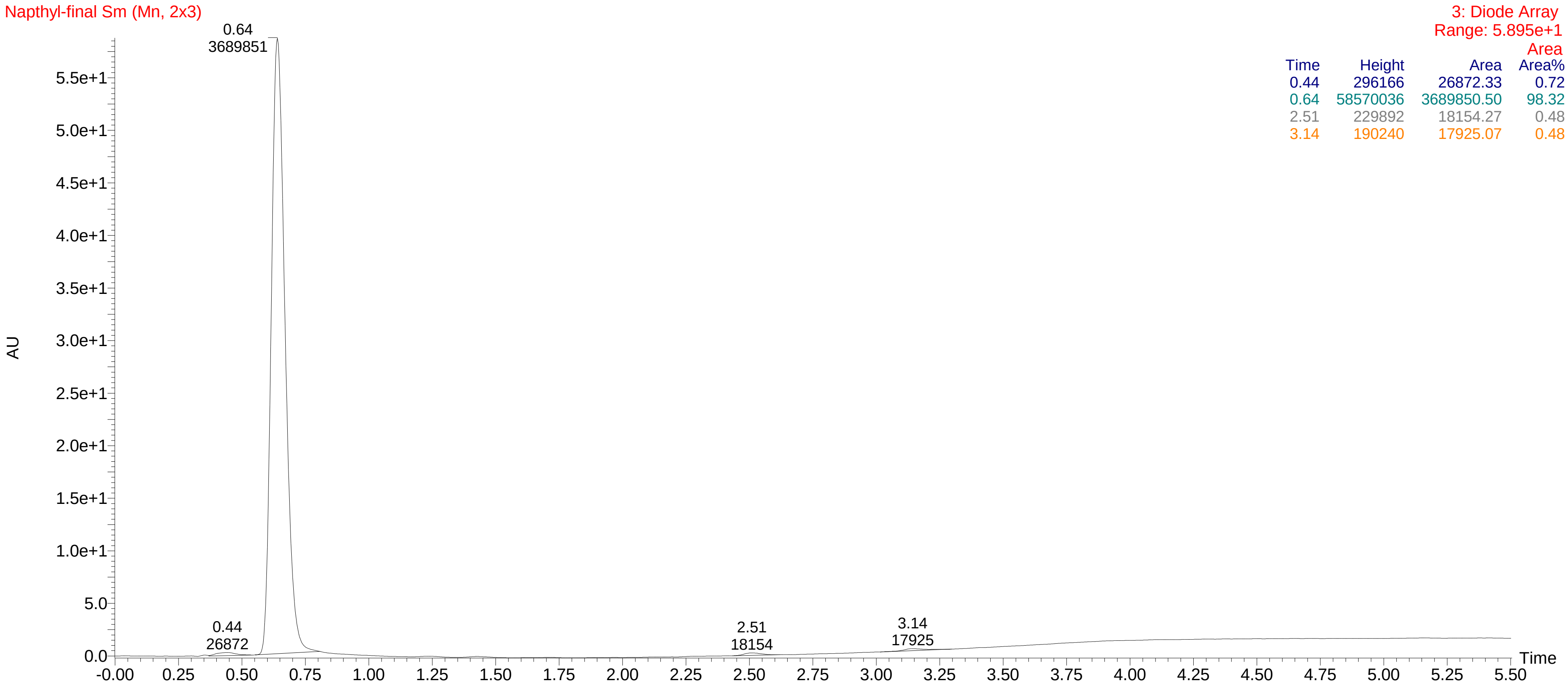


| Time | Height | Area | Area% |
|---|---|---|---|
| 0.44 | 296166 | 26872.33 | 0.72 |
| 0.64 | 58570036 | 3689850.50 | 98.32 |
| 2.51 | 229892 | 18154.27 | 0.48 |
| 3.14 | 190240 | 17925.07 | 0.48 |

HPLC of compound **14t**

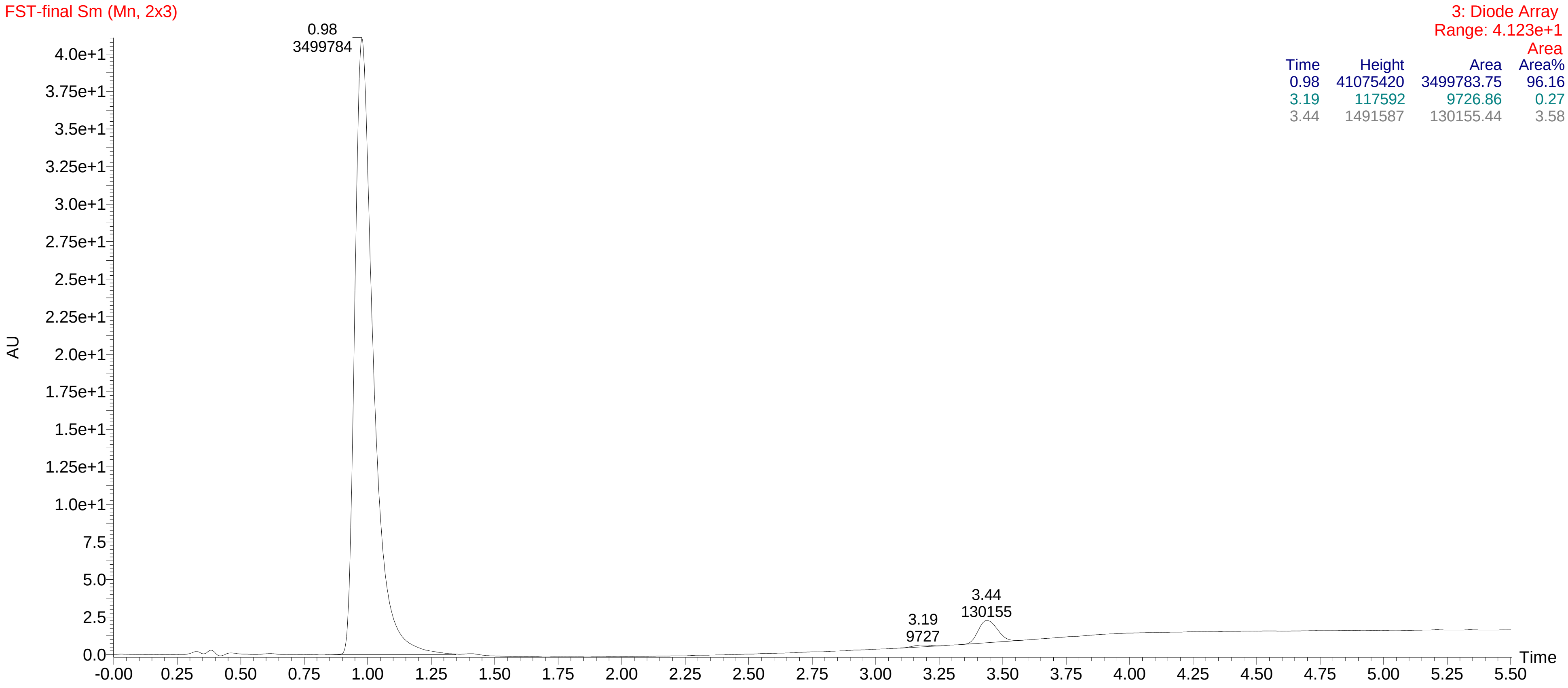


| Time | Height | Area | Area Area% |
|---|---|---|---|
| 0.98 | 41075420 | 3499783.75 | 96.16 |
| 3.19 | 117592 | 9726.86 | 0.27 |
| 3.44 | 1491587 | 130155.44 | 3.58 |

HPLC of compound **15a**

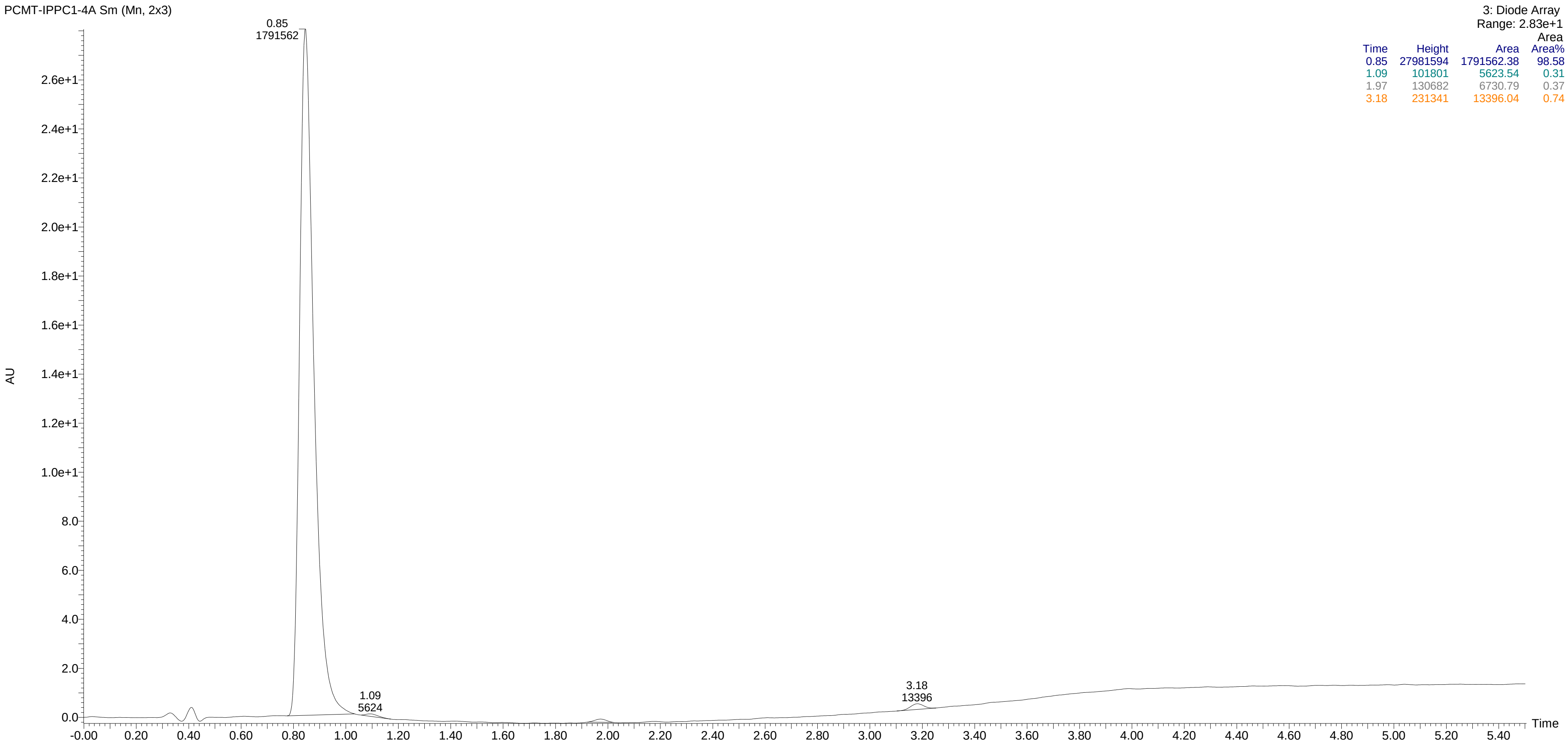


| Time | Height | Area | Area% |
|---|---|---|---|
| 0.85 | 27981594 | 1791562.38 | 98.58 |
| 1.09 | 101801 | 5623.54 | 0.31 |
| 1.97 | 130682 | 6730.79 | 0.37 |
| 3.18 | 231341 | 13396.04 | 0.74 |

HPLC of compound **15b**

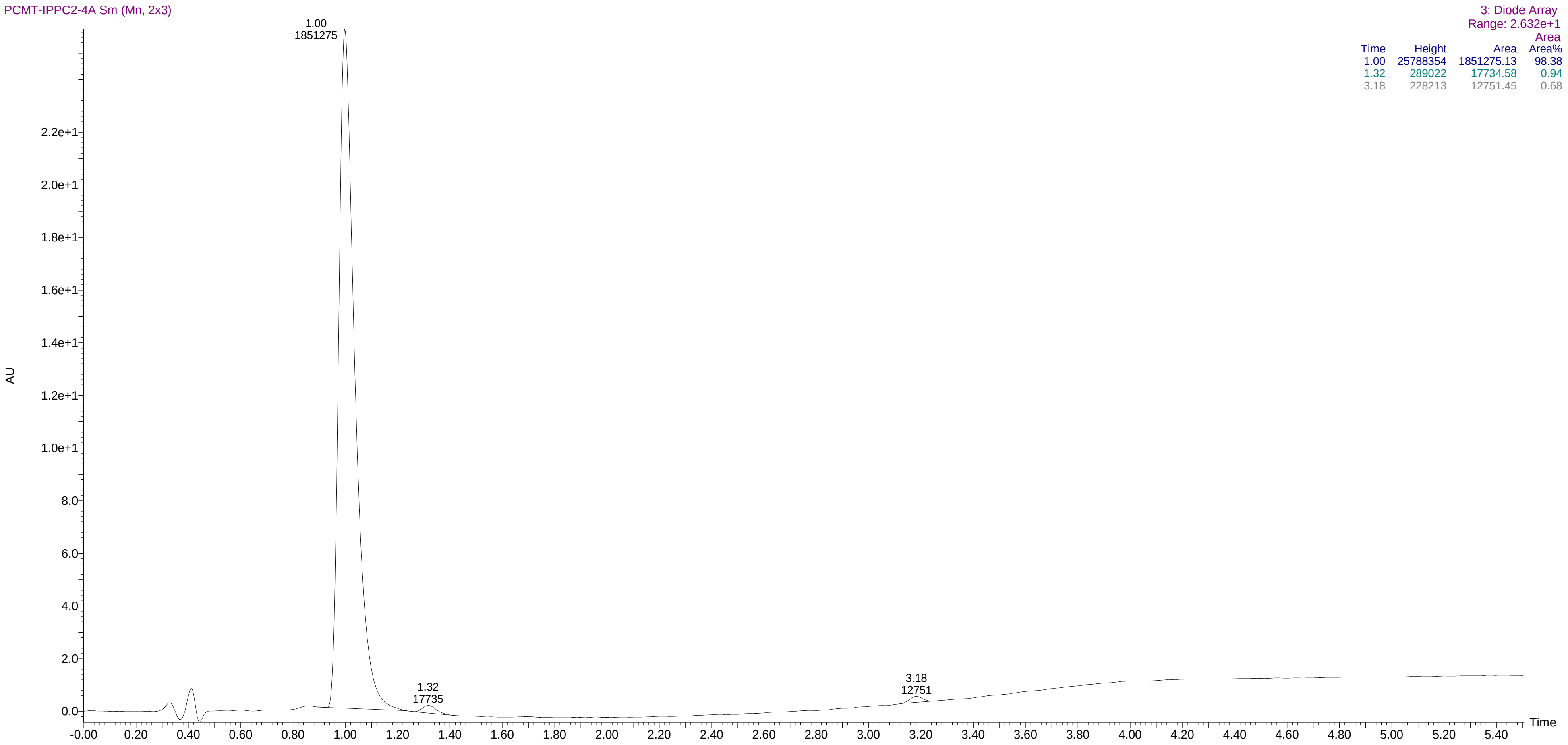


| Time | Height | Area | Area Area% |
|---|---|---|---|
| 1.00 | 25788354 | 1851275.13 | 98.38 |
| 1.32 | 289022 | 17734.58 | 0.94 |
| 3.18 | 228213 | 12751.45 | 0.68 |

HPLC of compound **15c**

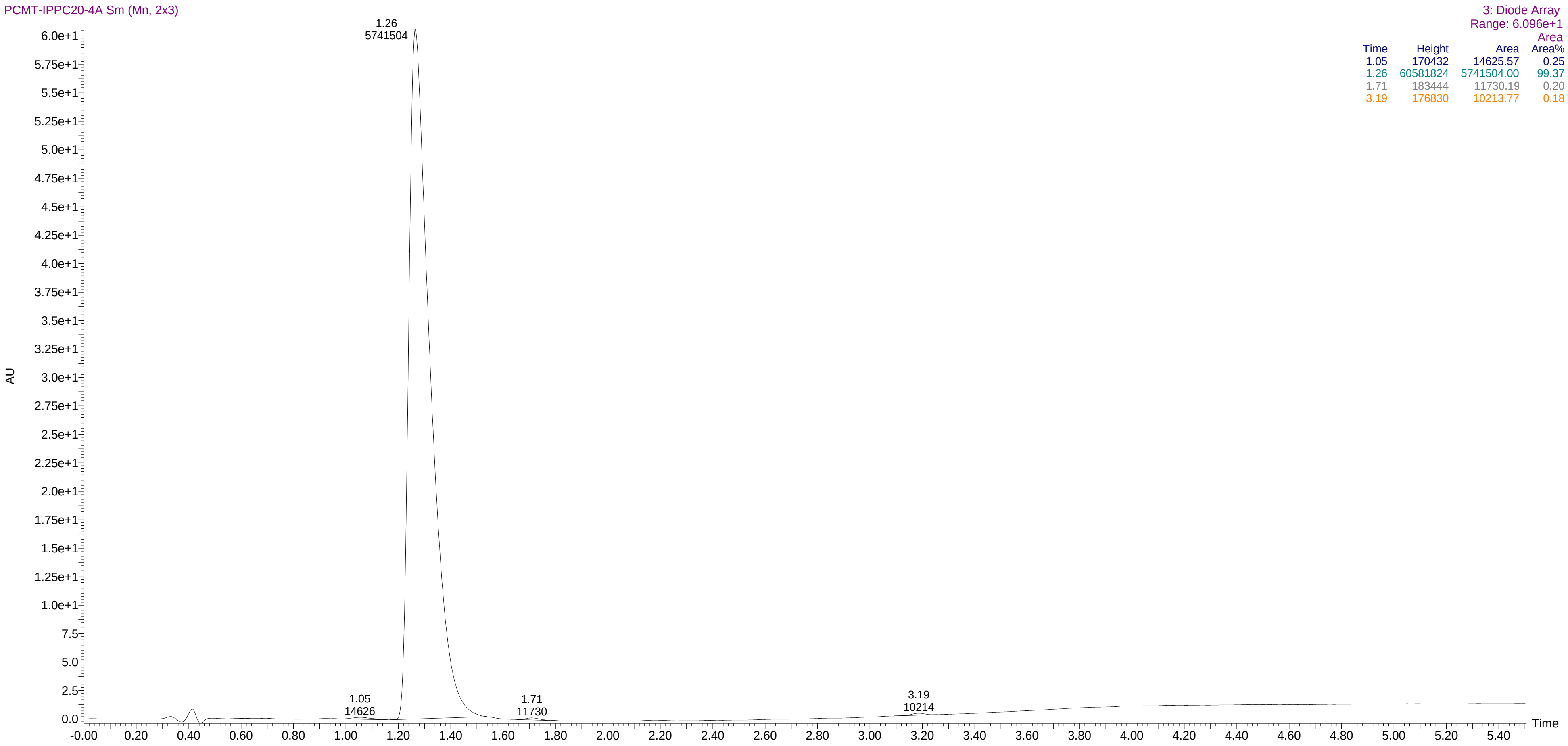


| Time | Height | Area | Area% |
|---|---|---|---|
| 1.05 | 170432 | 14625.57 | 0.25 |
| 1.26 | 60581824 | 5741504.00 | 99.37 |
| 1.71 | 183444 | 11730.19 | 0.20 |
| 3.19 | 176830 | 10213.77 | 0.18 |

HPLC of compound **15d**

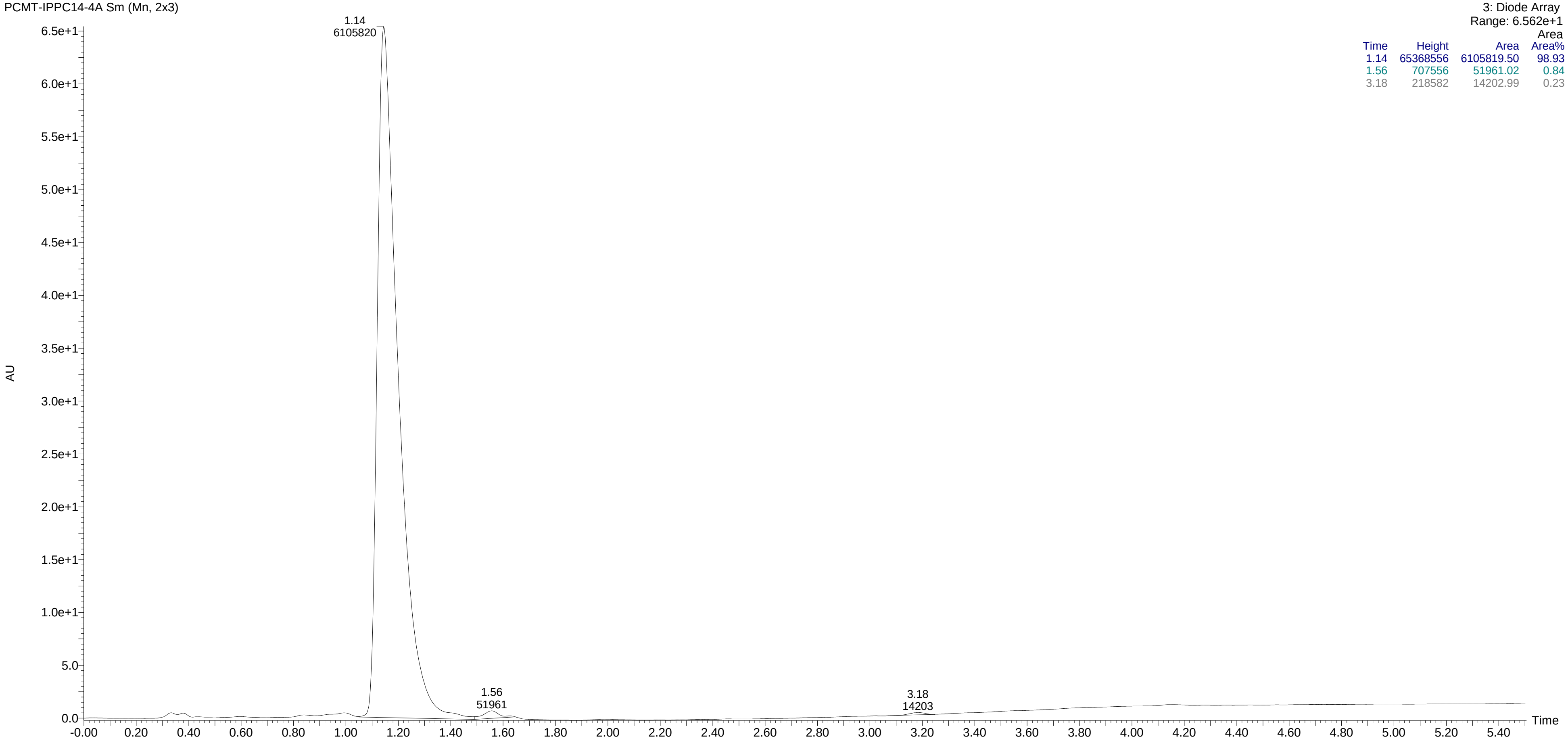


| Time | Height | Area | Area% |
|---|---|---|---|
| 1.14 | 65368556 | 6105819.50 | 98.93 |
| 1.56 | 707556 | 51961.02 | 0.84 |
| 3.18 | 218582 | 14202.99 | 0.23 |
